# Response of barley (*Hordeum vulgare*) leaves to vapour pressure deficit (VPD) involves rapid shifts in transcript pools

**DOI:** 10.64898/2026.11.24.768816

**Authors:** Maitry Paul, Jaakko Tanskanen, Hannes Kollist, Ebe Merilo, Alan H. Schulman

## Abstract

Barley (*Hordeum vulgare* L.) growth and productivity are sensitive to environmental fluctuations, such as changes in air humidity and the resulting vapour pressure deficit (VPD), which impact plant transpiration, stomatal conductance, and hence photosynthesis. We investigated the rapid stomatal and transcriptomic responses of barley to elevated VPD. Using a custom-built gas-exchange platform, stomatal conductance (Gs) was measured, and leaf RNA samples collected, at 10-minute intervals post-VPD increase. RNA-seq analysis was performed to assess changes in transcript abundance and affected pathways and networks. We found a hydropassive increase in Gs immediately following the VPD shift, followed by a sharp decline as an active response within 10 min, and a gradual recovery by 40 min, demonstrating rapid acclimation to VPD in barley. Rapid changes in transcript abundance for differentially expressed genes (DEGs) were seen, with sharp downregulation of some DEGs already 2 min after VPD increase. Key genes, including those encoding receptor-like kinases, an ABCG48 protein, and calcineurin-binding protein, were linked to rapid stress responses. These findings highlight adaptation to short-term VPD changes in barley through coordinated physiological and molecular mechanisms, providing insights into potential genetic targets for improving drought tolerance. This research contributes to strategies for enhancing water-use efficiency and resilience in barley under fluctuating environmental conditions.

## 1 INTRODUCTION

The growth and productivity of barley (*Hordeum vulgare* L.) are sensitive to environmental conditions, particularly changes in water supply and atmospheric evaporative demand (Honsdorf et al. 2014; Paul et al. 2024). Vapor pressure deficit (VPD), the difference between the moisture in the air and its maximum water-holding capacity, affects plant transpiration and stomatal behaviour (McAdam et. al. 2016). In barley, stomata are distributed on both the adaxial (upper) and abaxial (lower) leaf surfaces, playing a crucial role in regulating gas exchange and water loss, with densities varying on both sides depending on environmental fluctuations (Rahim et al. 2021; Samantera et al. 2026). Stomatal adjustments in response to VPD are key for regulating water loss and CO₂ uptake (Burgess & Ugalde, 2024). As the air temperature rises, its moisture-holding capacity and consequently VPD also rises (i.e., the air becomes less humid), leading to higher water loss through transpiration. The stomata respond by reducing their aperture to conserve water and maintain internal hydration, a mechanism regulated by abscisic acid (ABA), which accumulates in the guard cells that flank the stomata (McAdam et al. 2016).

The molecular response of guard cells to elevated VPD in angiosperms involves both passive and active mechanisms (Jalakas et al. 2021). Initially, the rise in VPD causes increased transpiration, which in turn results in lower epidermal turgor and thus less epidermal backpressure on the guard cells, causing an initial wrong-way response, i.e., stomatal opening (Buckley, 2019). To counteract this hydropassive opening and prevent further water loss, plants activate an ABA-dependent pathway. ABA binds to its receptors (PYR1/PYL/RCAR), inhibiting Protein Phosphatase 2C members (PP2C) and activating the protein kinase OST1. OST1 triggers ion efflux through anion channels, including SLAC1 and ALMT12/QUAC1, leading to stomatal closure. Additionally, an ABA-independent pathway also contributes, where OST1 is activated under osmotic stress and low humidity via the SnRK2-box, bypassing the need for ABA signalling. This allows stomatal closure even in the absence of significant ABA level changes (Jalakas et al. 2021). This dual mechanism ensures a fast and efficient stomatal response to VPD changes.

These molecular changes, extending beyond guard cells, affect other tissues, leading to the metabolic adjustments and gene regulation necessary for adaptation to fluctuating environmental conditions, including shifts in VPD, humidity, and temperature. While much is known about the long-term effects of VPD on stomatal conductance from drought and heat experiments (Paul et al. 2023, 2024), the rapid response mechanisms that barley employs to maintain water balance in the short term remain unclear. Here, we aimed to explore the short-term responses to changes in VPD. Specifically, we seek to determine how quickly stomatal conductance adjusts in response to a change in VPD and to characterize changes on the whole-leaf level in the transcript pool within the first minutes after a shift in VPD. By examining the early molecular responses, we hope to gain insights into the targets and mechanisms used to physiologically adapt to changes in atmospheric water demand, which may also prime the plant for long-term stress tolerance and develop a set of potential candidate genes for improving climate tolerance.

In this study, we focused on the Golden Promise, the model cultivar for transformation and gene editing, targeting barley plants aged 18 to 24 days. The goal was to analyse the transcriptional responses of barley leaves at defined time points, both before and during exposure to elevated VPD, which was done in a controlled-environment gas exchange chamber. All plants in the study were kept well-watered to ensure that any changes seen were due to air water demand and not because of decreased soil water supply.

## 2 MATERIALS AND METHODS

### 2.1 Plant Material and Growth Conditions

Barley cv. Golden Promise seeds were sown into 1 L pots (4 : 2 (v/v) peat : vermiculite substrate) and grown in a growth chamber (41AR2cLED; Percival Scientific, Perry, IA, USA) at the University of Tartu, Estonia. The growth chamber maintained a temperature of 23°C/19°C (day/night), relative humidity (RH) of 65%/80% (day/night) and VPD of 0.97/0.43 kPa, with a 12-hour photoperiod (6:00-18:00) providing photosynthetically active radiation (PAR) of 700 µmol m^-^² s^-1^. Experiments were performed to follow a similar diurnal pattern: they began between 10:00 and 11:00 AM with stabilization at low VPD for 2 hr, followed by 40-min high VPD treatment. Strict diurnal timing was necessary to minimize the effects of diurnal time on transcript pools. Barley plants, aged 18-24 days at the 6—to 7-leaf stage, were placed within a four-chamber gas exchange platform (Horak et al. 2017). During gas exchange measurements, sampling duration of reference and measurement streams for H_2_O and CO_2_ concentrations was 60 sec each, which means that cuvettes were sampled successively at 8-min intervals. Leaf samples for RNA extraction were collected in five biological replicates at 2-, 12, 22, and 42 minutes after the VPD was raised. Control plants were kept at low VPD and leaf samples collected as for the experimental plants. In total, 40 plants were sampled, 20 each in the control and treatment groups with five replicates per time point.

### 2.2 Gas Exchange Experiments

Stomatal conductance in barley plants was measured using a custom-built, thermostated, four-chamber, flow-through gas-exchange device (Jyrkki MAX, Plant Invent) at the University of Tartu, (Horak et al. 2017). The device comprised temperature-controlled gas-exchange cuvettes formed of two glass cylinders (i.d. 10.6 cm, height 15.6 cm) with a thermostated water jacket in between. A perforated glass plate accommodated the plant stem, with modelling gum ensuring an airtight seal between the plant shoot and the soil-enclosed roots. Further details regarding the platform specifications are available in Horak et al. (2017).

Stomatal conductance was measured under 70% RH, 800-900 µmol m^-^² s^-1^ of PAR, and 22°C to 23°C air temperature. After stabilization of stomatal conductance, plants were subjected to dry air, reducing the chamber humidity to an average of 32% and raising the VPD. The VPD for all control plants at all time points was 0.8 kPa, whereas high VPD during the 40-min treatment interval was maintained at around 1.9 kPa.

### 2.3 Isolation of RNA, RNA-seq, and gene enrichment analysis

The uppermost leaf from the plant was collected for all treatments and controls at 2-, 12-, 22- and 42-minute intervals (T2, T12, T22, T42), with T2 being the point at which the leaf was frozen, about 2 min after change in the VPD was imposed. For RNA extraction, leaf samples were collected in liquid nitrogen and brought to Helsinki on dry ice for transcriptomic analysis. Total RNA was extracted using a Qiagen kit; the remaining steps followed a previously described method (Paul et al. 2023). The five sets of biological replicates from each time point and their controls at the same time points were used for RNA-seq analysis. Libraries were prepared using the TruSeq Stranded Total RNA kit, which removed cytoplasmic and chloroplast ribosomal RNA (Illumina). TruSeq adaptors were used. Sequences were produced from one lane of an Illumina NovaSeq 6000 S4 200-cycle flow cell at the Institute for Molecular Medicine Finland (FIMM), yielding an mean of 75.5 million (median, 73.2 million) reads of 2 x 100 bp, untrimmed, per biological replicate.

FastQC (Andrews, 2010) was employed for quality control and trimming of the RNA-seq data. Gene-level exploratory and differential expressed gene (DEG) analyses were performed with DESeq2 (Love et al. 2015). Venny 2.1.0 was used to create Venn diagrams from the DEGs in order to compare common genes between treatments and controls (Oliveros, 2007-2015). ShinyGO v0.80 was used for fold-enrichment analysis to identify upregulated and downregulated pathways (Ge et al. 2020). The analysis involved comparing DEGs against the Ensembl ID of *hvgoldenpromise_eg_gene* with taxonomy ID 112509. A significance cutoff of p ≤ 0.05 was applied to highlight the top 10 pathways. For VPD and Gs significance, the one-way ANOVA Tukey Post-Hoc test at a significance level of *p* = 0.05 was carried out using SPSS. In the reported analyses, lines without significant differences share superscript letters. Other figures were created using MS Excel.

## 3 Results

### 3.1 Dynamic response of stomatal conductance to change in VPD

The relationship between VPD and stomatal conductance (Gs) was investigated under both control and high VPD conditions in barley cv. Golden Promise over 40 min, following a 2-hr stabilization, using a custom-made plant gas exchange platform (Figure S1). Gas exchange data was collected at 8-min intervals. Stomatal conductance was measured over the course of a continuous physiological experiment (2 hr stabilization at normal (low) VPD followed by high VPD for 40 min, Figure 1). Separate experiments were run to sample leaves for RNA analysis, terminating the gas exchange measurements at the given time point in the sampled cuvette (Figure 2). Leaves were harvested for RNA at 2, 12, 22, and 42 min after the upward shift in VPD.

**Figure 1.**
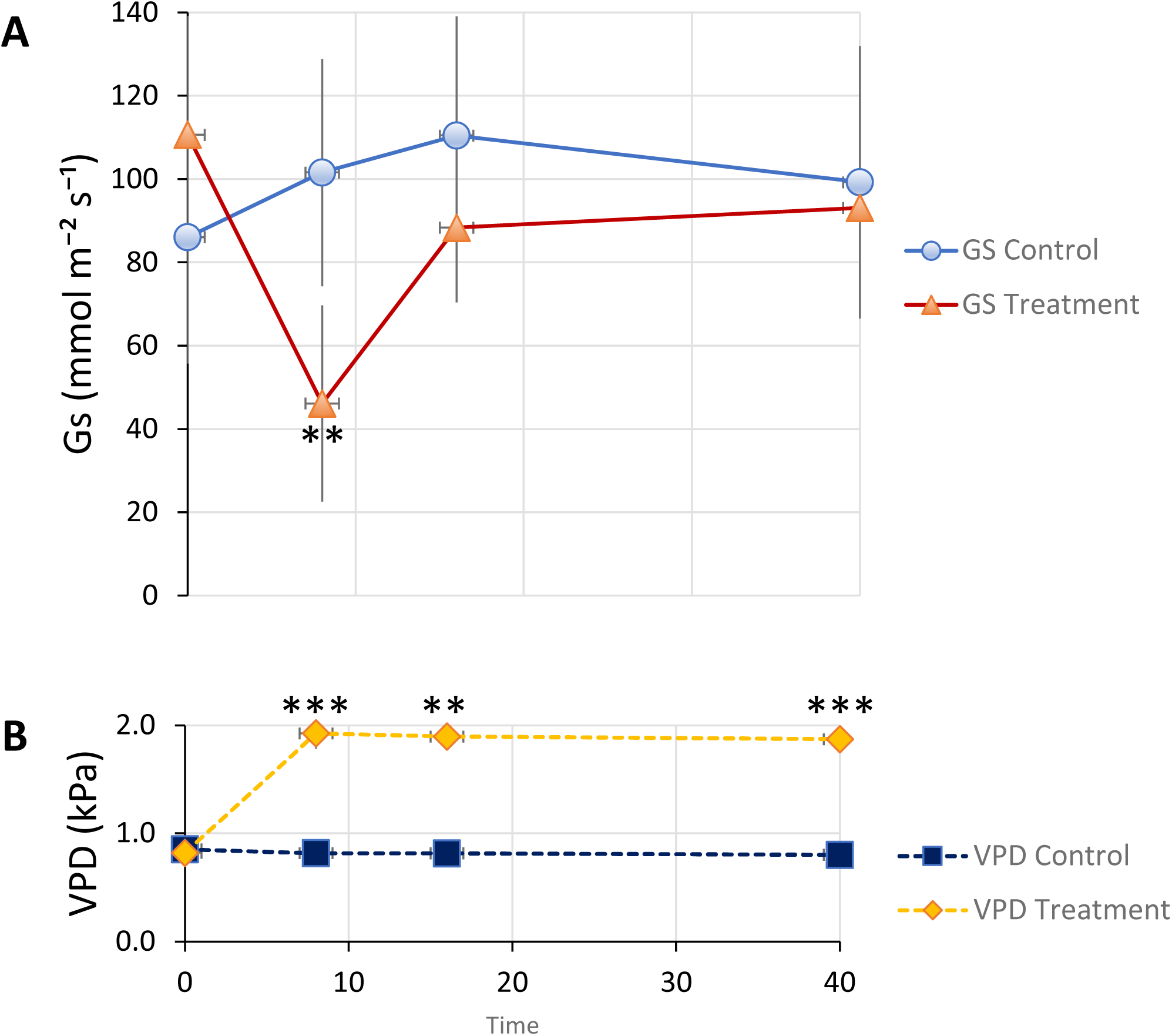
Time course of VPD and stomatal conductance (Gs) for control and treatment groups. (A) Gs: blue line with circle represents controls, red line with triangle represents treatment samples. (B) VPD: blue dashed line with square shows controls (0.8 kPa), while yellow dashed line with diamond shows treatment samples with elevated VPD (1.9 kPa). Each point represents data from five biological replicates. ** indicates p ≤ 0.01 and *** indicates p < 0.001, based on independent-sample t-tests.

**Figure 2.**
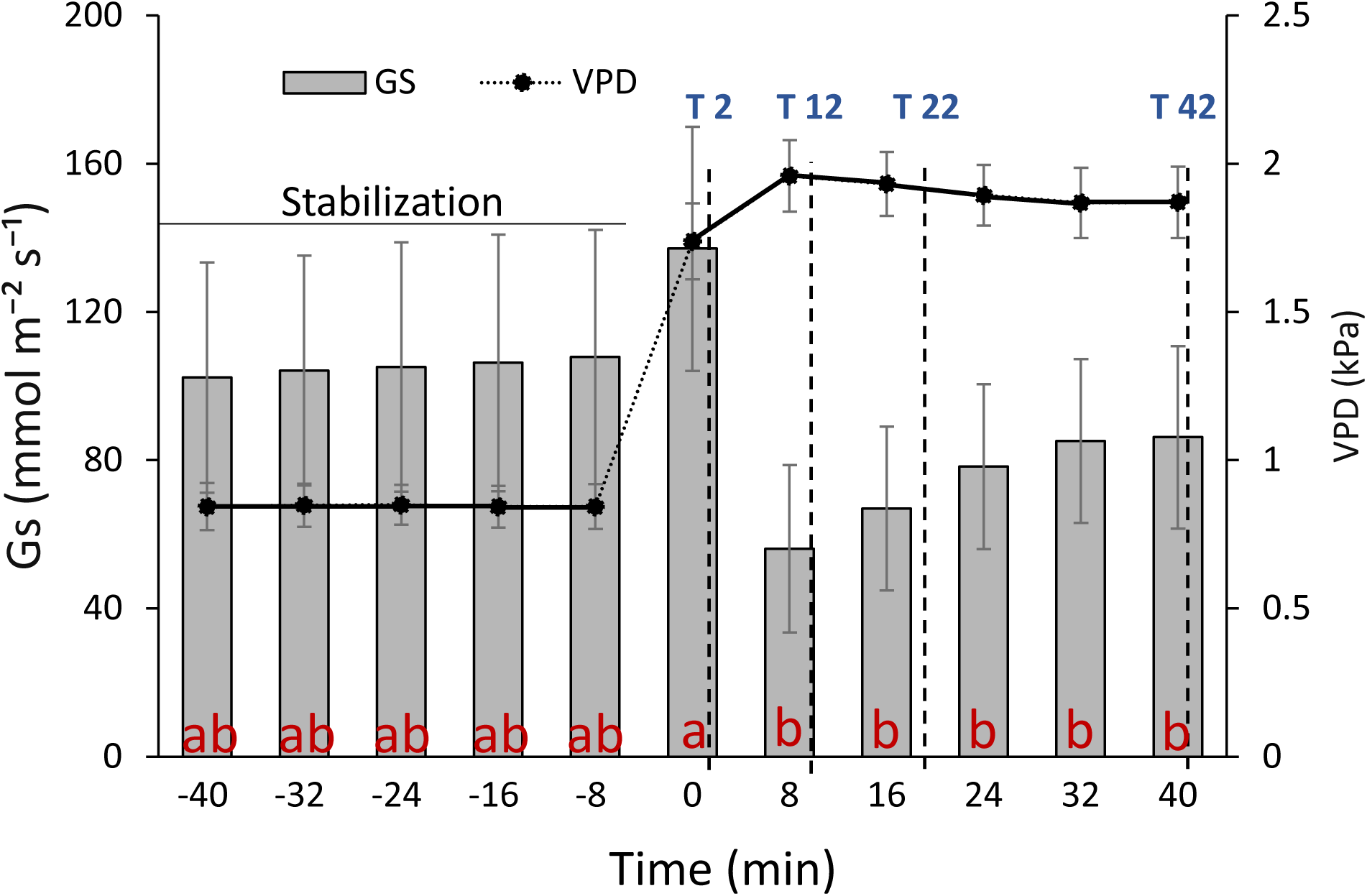
Stomatal conductance (Gs) in response to vapour pressure deficit (VPD) during leaf sampling for RNA. Bars correspond to Gs and line with closed circles to VPD measurement points, whereas the dashed lines indicate leaf sampling points (2, 12, 22, 42 min) for RNA isolation. The data is from five biological replicates. Lowercase letters indicate significance groups for Gs. During high VPD treatment, the VPD was always significantly higher than in under control (low VPD) conditions.

Throughout the stabilization phase, when plants were subjected to a constant VPD of ∼0.8 kPa, Gs was fairly constant, with minor fluctuations (Figures 1, 2). The average stomatal conductance during the stabilization phase was 105 mmol m⁻² s⁻¹. An increase (Figure 2) in Gs was observed at T2, reaching 137 mmol m⁻² s⁻¹ as VPD rose from 0.8 kPa to 1.9 kPa (Figures 1, 2). This could be explained as a hydropassive response of guard cells to decreased epidermal back pressure before active stomatal closure starts. It has been termed the wrong-way response (WWR; Hansen et al. 2014; Zait et al. 2024), as stomata are expected to decrease Gs when VPD increases.

However, just before T12 and under high VPD, Gs experienced a sharp decline compared with the controls at the same time points (Figure 1), dropping by 56 mmol m⁻² s⁻¹ (around 60% reduction compared to T2; Figure 2), by 10 min. After this decline, Gs gradually recovered, reaching 86 mmol m⁻² s⁻¹, which is approximately 80% of control Gs values by 20 min and 94% by 40 min. In both the continuous physiological experiments and in the destructive RNA-sampling experiments, we saw stomatal conductance exhibiting dynamic changes over time, recovering under the constant elevated VPD.

### 3.2 Analysis of rapid changes in RNA transcript pools in leaves

Given the rapid drop in stomatal conductance in response to an increase in VPD, followed by gradual recovery, we looked for parallel rapid changes in transcript pools. We carried out RNA-seq on leaves at 2 min following the VPD increase (T2), 10 min thereafter (T12), and then at T22 and T42, at which point Gs was restored to about 84 % of the initial level. At T2 (Figure 2), the increased VPD likely drives higher Gs due to diffusion physics, when the guard cells have not yet actively responded. The expression of only 36 genes at T2 is greater than two-fold upregulated (Table S1), though none have an adjusted *p*-value (*p*adj) of less than 0.05; one (described below) is 46-fold upregulated with a highly significant *p*-value (9 x 10^-5^), though its *p*adj is not significant. In contrast, there are 125 genes downregulated greater than two-fold at T2, of which three were highly significantly downregulated, with *p*adj ≤ 0.005, from 36-to 7.5-million-fold (Table S2, Figure 3). In general, the proportion of upregulated genes increased from T2 to T42, and that of downregulated genes decreased.

**Figure 3.**
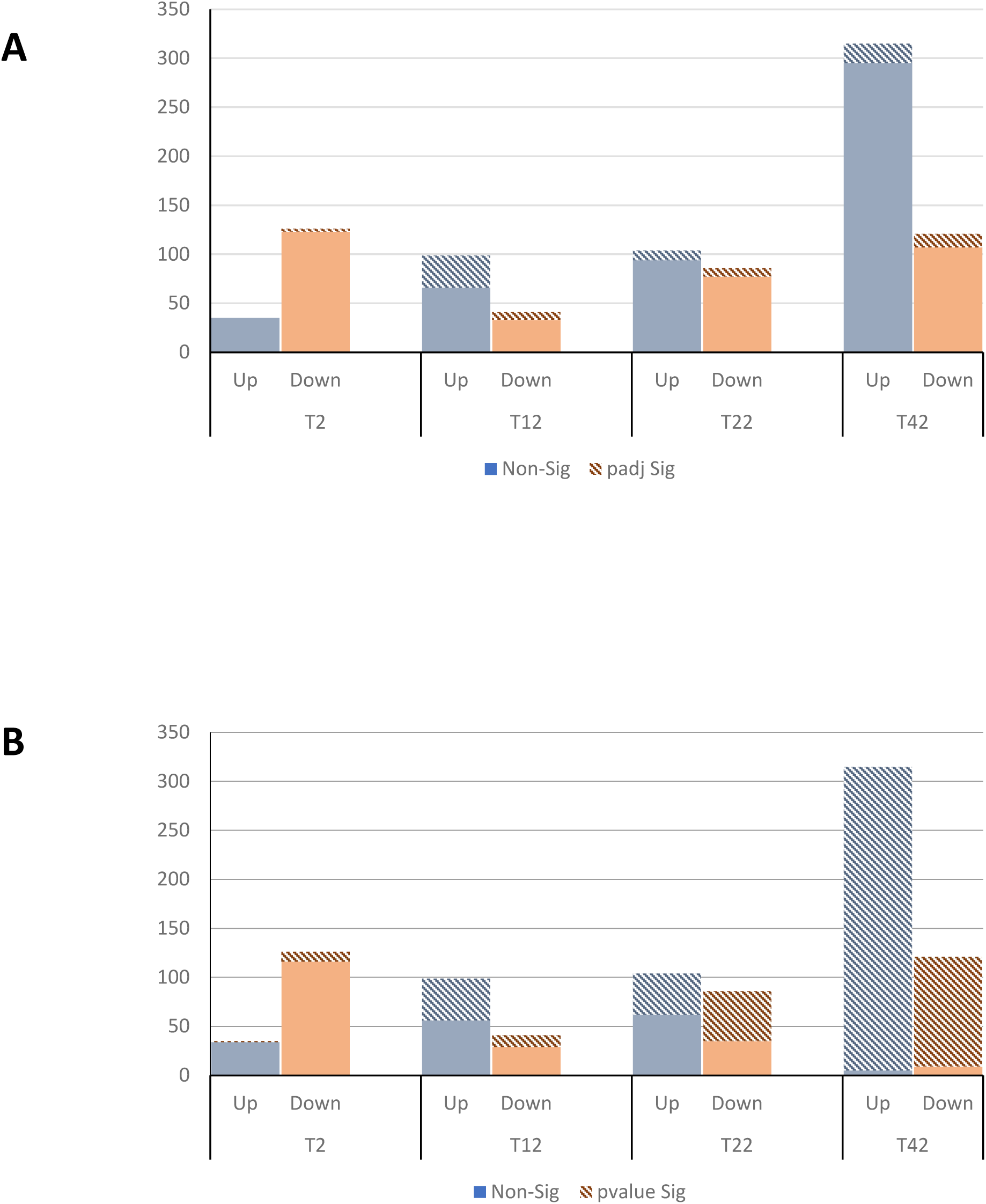
Number of genes differentially expressed genes (DEGs), following an increase in VPD, at greater than twofold compared to the control for each time point. (A) total DEGs (solid) and those with *p*adj < 0.05 (hatched) (B) total DEGs (solid) and those with *p < 0.05* (hatched)

At T12, altogether 99 genes were more than twofold upregulated, of which 33 were significant at *p*adj < 0.05 and had changes ranging from 20 thousand-to over 44 million-fold (Table S3). Only 41 genes were >2-fold downregulated, 12 at *p* < 0.05 and 8 at *p*adj < 0.05 (Table S4). These eight had changes from 7 to 1.7 million-fold. The relative share of up- and downregulated genes showed the same trend at the later time points. At T22, 104 were more than twofold upregulated, 42 significantly at *p* < 0.05 and 10 at *p*adj < 0.05 (Table S5). Among these 10, changes ranged from 15 thousand-to 4.4 million-fold. A total of 86 were more than twofold downregulated, of which 51 were significant at *p* < 0.05, of which 6 had *p*adj < 0.05, with fold changes ranging up to 3 million (Table S6). At T42, 315 genes were more than twofold upregulated, with 310 at *p* < 0.05 and 20 at *p*adj < 0.05 (Table S7). Of these, the degree of upregulation ranged from 8- to 214-fold. By comparison, only 121 genes were more than twofold downregulated at T42 (Table S8), of which 112 were significant at *p* < 0.05 and 14 at *p*adj < 0.05. Of these 14, changes in expression ranged from 8-fold to 24-million-fold.

We examined all genes more than twofold differentially regulated that were in common between the different time points following the VPD shift (Figure 4). Regarding the ones switching the direction of change in their expression, 26 of the downregulated genes at T2 were upregulated already at T12 and 18 at T22, but only 1 at T42. Of the genes upregulated at T2, 2 were downregulated at T12, 6 at T22, and 1 at T42. Considering the genes that kept their direction of expression change at successive timepoints, 10 of the genes upregulated at T2 remain upregulated at T12, 5 at T22, and 2 at T42. Of those downregulated at T2, 12 remained so at T12, 12 at T22, and 5 at T42. As expected, we observed no overlap between upregulated and downregulated genes for any one time point. Taken together, the results suggest that rapid stomatal closure is also associated with a parallel rapid transcriptional response, with particular sets of transcripts increasing or decreasing in abundance in direct or inverse correlation with the direction of change in Gs. Individual genes with significant shifts in expression are discussed below in Section 3.3.

**Figure 4.**
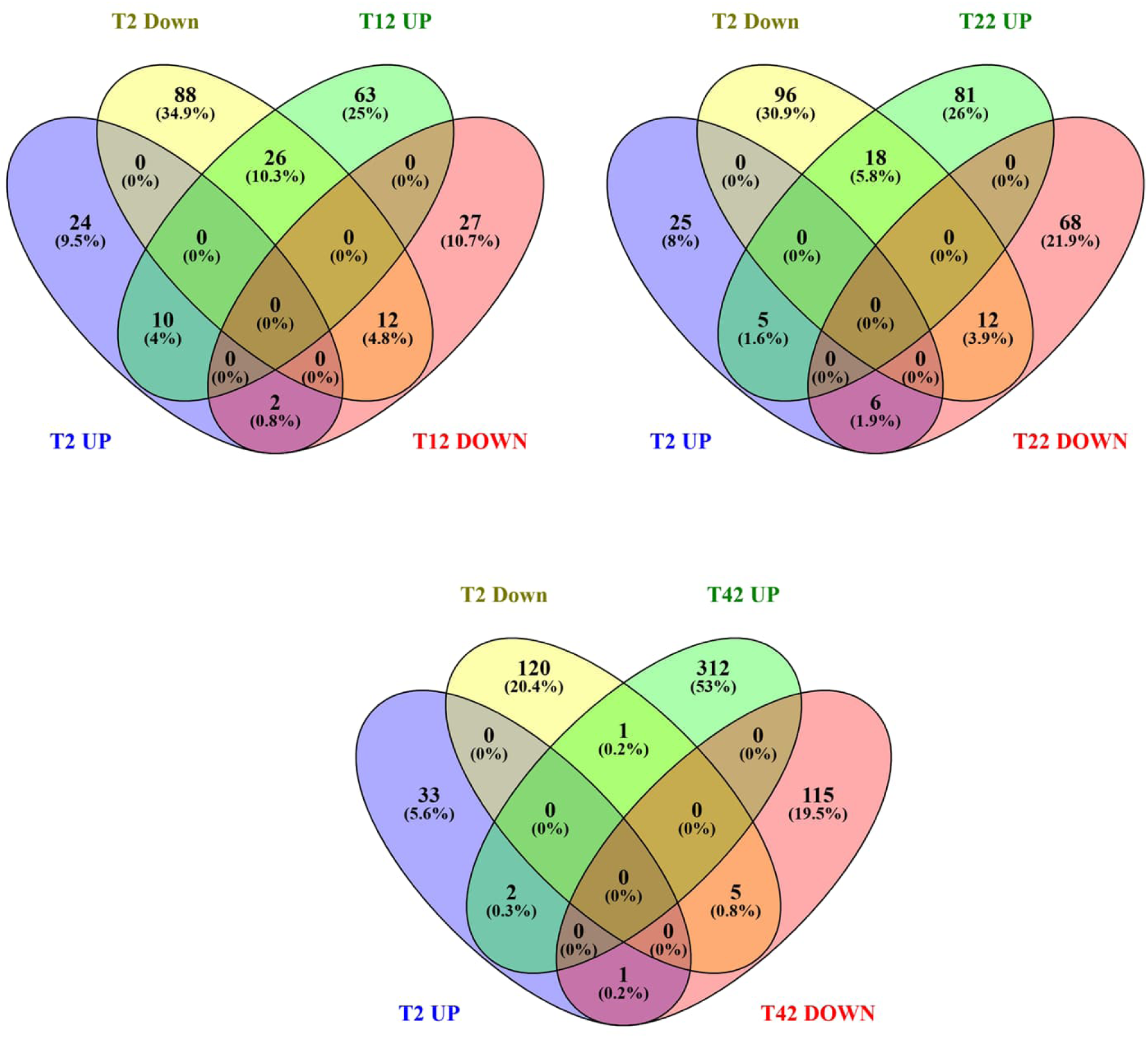
Transcripts shared between T2 and successive timepoints that are more than twofold differentially expressed following an increase in VPD. Upregulation (“UP”) or downregulation (“DOWN”) is indicated in the figures.

### 3.3 Gene-specific abundance changes

The total of 929 differential expression changes that were greater than twofold compared with the controls, pooled for all stages, involved altogether 794 unique genes (Table S16). While pathway analysis of many of the DEGs has been treated above, here we focus on some individual genes among those, for which the transcripts were both relatively abundant and showed strong and statically significant differential expression.

#### Changes after 2 min of high VPD

Over the very short time interval between the change in VPD and the taking of the T2 sample, only four genes (Tables S1, S2) showed significant (adjusted *p*-value, *p*adj < 0.05) changes in abundance, three being down-regulated. Only two of the top forty changes represented increases in abundance. The single significantly upregulated gene (46-fold) at T2 (*p* = 9.22 x 10^-5^; Table S1) is identified in the Barlex v.2 barley assembly as a CACTA transposon protein.

Of the three significantly downregulated genes, HORVU.MOREX.r2.3HG0183610.1 (hereafter, genes in Morex assembly v. 2 will be referred to only by chromosome and gene, e.g. 3HG0183610.1 here), decreased by 7.5-million fold to 1.3 x 10^-7^ of control levels, *p*adj 2.7 x 10^-5^) and is a perfect match for a barley putative receptor-like protein kinase (RLK; E-value 0.0). Alphafold gives high confidence (score 80.75) that it contains a protein-kinase domain, corresponding to UniProt A0A287JXC1. A BLASTp search with this protein gives 89% identity (E = 0.0) to the *T. durum Lr10* RLK (VAH55245) for which, however, there is no information on a role in drought or VPD response. Nevertheless, RLKs are well documented to have a role in response to abiotic stresses (Ye et al. 2017). For example, in rice, the plasma membrane receptor-like kinase leaf panicle 2 (LP2), is down-regulated by drought and acts downstream of the DROUGHT AND SALT TOLERANCE transcription factor to regulate drought sensitivity (Wu, 2015). Overexpression of *LP2* reduces the accumulation of H_2_O_2_ and promotes open stomata. It is directly regulated by the zinc finger transcription factor DST and interacts with the drought-responsive aquaporins OsPIP1;1, OsPIP1;3, and OsPIP2;3. However, LP2 (Os02g0615800 in *Oryza sativa* ssp. *japonica* cv. Nipponbare) shares only 33.8% identity over a 234-residue overlap with A0A287JXC1.

The second most strongly down-regulated gene at T2 (to 0.005 of control levels) is 4HG0295170.1, which corresponds to UPI00162DAA44 and is identified by BLASTp as ribosomal protein S4, having perfect matches in *Hordeum* and *Triticum*. However, there is no literature on its role in stress response. The third, downregulated 36-fold (*padj*= 0.0005), corresponding to UPI0002004F15, is a perfect match to chlorophyll a-b binding protein LHCII type 1 (E-value 0.0). In Arabidopsis, short-term drought stress causes a rapid disassembly of LHCII (Chen et al. 2016b), consistent with its downregulation here.

#### Changes after 12 min of high VPD

By T12, already 42 genes have changed abundance significantly (*p*adj < 0.05); 33 represent increases in expression; of these, only two had a high normalized abundance (base mean) compared to the observed range (0.08 -- 10,482). Of these, 1HG0002570.1 corresponds to Uniprot UPI00162ECF7C and is 97% identical to the receptor-like kinase LRK10. However, a Clustal alignment on the protein level to 3HG0183610.1, which at T2 is downregulated, shows that these are different RLKs, having only 54.3% identity. In Arabidopsis, a gene encoding *AtLRK10L1*, most closely related to wheat *LRK10*, expresses two different transcripts, LRK10L1.1 and LRK10L1.2 (Lim et al. 2015). An *lrk10l1.2* mutant exhibits reduced tolerance to drought stress, compared with wild type, accompanied by increase in stomatal aperture, suggesting that the upregulation seen here is involved in stomatal closure.

The 2HG0127650.1, corresponding to Uniprot UPI00162C68FB, is upregulated over 20-thousand fold. Both BLASTp and Alphafold identify it as cytochrome b6, a di-heme cytochrome. In Arabidopsis, a ROS-producing abiotic stress (SO_2_) leads to *Cytb6f* upregulation (Li & Yi, 2020); it is also upregulated in salt-stressed wheat chloroplasts (Kamal et al. 2012). The third relatively abundant upregulated transcript, three-fold over 10 min (*p*adj 0.067), is from 5HG0432480.1, corresponding to the protein UPI00162EB035. This matches the barley ABC transporter (<u>A</u>TP-<u>b</u>inding <u>c</u>assette) G family member 48-like isoform X1 (ABCG48) fully (E-value 0.0). Arabidopsis ABC transporters of the MULTIDRUG RESISTANCE-ASSOCIATED PROTEIN family, such as AtMRP5, AtMRP4, and AtABCG22, are involved in modulating guard cell responses (Klein et al. 2003; Kuromori et al. 2016). Mutants of AtMRP5 exhibit impaired guard cell function, affecting water use efficiency, while AtMRP4 mutants show decreased stomatal opening, contributing to enhanced drought tolerance (Li et al. 2021).

Recently, AGCG48 (SbSLT2), a plasma membrane protein, was shown in sorghum (*Sorghum bicolor*) to function as a strigolactone (SL) transporter in roots, driving the SL secretion that enables parasitism by *Striga* (Shi et al. 2025). Among their many roles, SLs have been shown to function in drought response. A mutation in gene for the SL-specific receptor in barley makes that line highly sensitive to drought stress and slow in stomatal closure compared to the parental line (Marzec et al. 2020).

Among the down-regulated genes, transcription of 2HG0115250.1 is reduced over 9 million-fold (*p*adj 1.1 x 10^-7^). Corresponding to UPI00162E99E4, it fully matches calcineurin-binding protein 1. Calcineurin B-like proteins (CBLs) in plants perceive calcium signals triggered by abiotic stress and interact with CBL-interacting protein kinases (CIPKs) to form a complex signalling network (Kaya et al. 2024). The Arabidopsis *AtCBL* family contains at least 10 members (Nagae et al. 2003) that are differentially regulated by stress conditions, with one (*AtCBL1*) being implicated in response to drought, cold, and wounding (Shi et. al. 1999). In wheat, the CBL gene family appears to have roles in both ABA-dependent and ABA-independent stress response, indicating a complex interaction with calcium signalling (Pouya et al. 2025).

Gene 2HG0089230.1 is downregulated 1.6-million-fold and is a *VASP*-like homolog; VASP proteins as actin polymerases regulate actin dynamics in animals (Faix & Rottner, 2022) but have no known function in plants and a claim of no homologs (Kollmar et al. 2012). Of the other significant DEGs, one is the transcript for a retrotransposon in superfamily *Copia* (4HG0315140), another (1HG0010570.1) is uncharacterized, a third (7HG0551580.1), 417000-fold reduced, encodes the mitochondrial 26.2 kD heat shock protein, shown to be rapidly induced in perennial rye grass (*Lolium perenne*; Rahman et al. 2022).

Gene 2HG0081740.1 was also downregulated, 218-fold (*p*adj 0.002). BLASTp identifies it as coding for the ATP synthase alpha subunit (also called subunit 1) in various plants, as well as ATP1 in barley, with 100% identity. A proteomics analysis in bermudagrass (*Cynodon dactylon*) found this subunit to be down-regulated in a drought-tolerant line (Zhao et al. 2011), as it was in response to salt stress in wheat (Kamal et al. 2012). While the protein is found in both mitochondria and chloroplasts, here its down-regulation may be associated with decreased photosynthetic capacity as a response to VPD.

Finally, 1HG0004210.1 is both significantly downregulated (*p*adj 0.018, 0.140 control levels) and expressed at substantial levels (177.3 base mean). BLASTp gives 100% alignment to metallothionein-like protein 2C (MT). In plants, MTs are divided into four types; the 17 cysteine residues in this one are diagnostic for an MT4 (Mierek-Adamska, et al. 2018). In oats, the transcriptional response to drought varies with the *MT* type: expression of *AsMT2* and *AsMT3* increases, while AsMT1 decreases (no *MT*4 in oat). Barley has all four MTs, but the response to e.g. water stress for the *MT4* gene has not been characterized earlier (Pourjalali et. al. 2023).

#### Genes with opposite changes at 2 min and 12 min

From T12 to T22 and onward to T42, stomatal conductance gradually recovers despite the continuing high VPD. Changes in gene expression at T2 are presumed to be connected to stomatal closure or response thereto, while changes at T12 may be connected to reopening under high VPD, because they precede the increase in stomatal conductance seen already by 16 min after the VPD change (Fig. 1). To look for genes whose change in expression between T2 and T12 might be connected specifically to stomatal closure and reopening, we examined those that changed in opposite directions at T2 vs T12 (Figure 4; Table S17).

None of the genes showed opposing changes at T2 and T12 where both members were statistically significant. Only two genes were upregulated at T2 but downregulated at T12 with more than two-fold changes compared to respective controls at those timepoints, both lacked statistical significance. One, 1HG0004660.1 (3.6-fold up at T2, 4.2-fold down at T12), corresponds to accession A0A8I6WGQ6 in Uniprot, a 100% match (E=0.0) over 307 residues to momilactone A synthase or CYP701A8, a cytochrome P450 (CYP) monooxygenase enzyme. The second gene that was up at T2 (2.7-fold) but down at T12 (8.9-fold) compared to the controls at each stage is 1HG0001460.1, annotated as a UDP-glucose glycosyl transferase (UGT) and also giving at 100% BLASTx match to this enzyme. An enzyme of this class was shown to be rapidly induced in Arabidopsis under drought stress, glucosidating the auxin indole-3-butyric acid (IBA) (Tognetti et al. 2010).

Of the four genes that were downregulated at T2 but upregulated at T12, two were significantly so at T2. One, 1HG0004990.1 (6-fold down at T2, *p* 0.02; 16-fold up at T12), corresponds to beta-fructofuranosidase (invertase, AVY91565.1) in barley; it is 95% identical on the protein level to a bamboo vacuolar invertase (ABB77251). Invertases have diverse roles in metabolism, development, and, for vacuolar intertases in particular, stress response (Liu et al. 2025). The other, 1HG0004560 (28.5-fold down at T2, *p* = 0.03; 3.2-fold up at T12), is 100% identical to disease resistance protein RGA5 (XP_044954660). Of the other two, 1HG0005910.1 is annotated at low confidence as a En/Spm transposon protein, whereas 1HG0000540.1 matches a gamma hordein gene.

### 3.3 Gene enrichment analysis for biological pathways

We carried out gene enrichment analyses to examine the coordinated regulation of DEGs falling into upregulated and downregulated biological pathways (limited to DEGs at *p* ≤ 0.05 and FDR 0.11) following the VPD increase (Tables S9-S15, Figures 5, S2, S3, S4). At T2, as only one gene was upregulated at *p* ≤ 0.05, no upregulated pathways could be ascertained (Table S1). The 11 downregulated genes at T2 entailed processes related to Photosystems I and II, chloroplasts, and photosynthesis generally, with PS I showing the strongest downregulation, 189-fold (Figure S2A, B).

**Figure 5.**
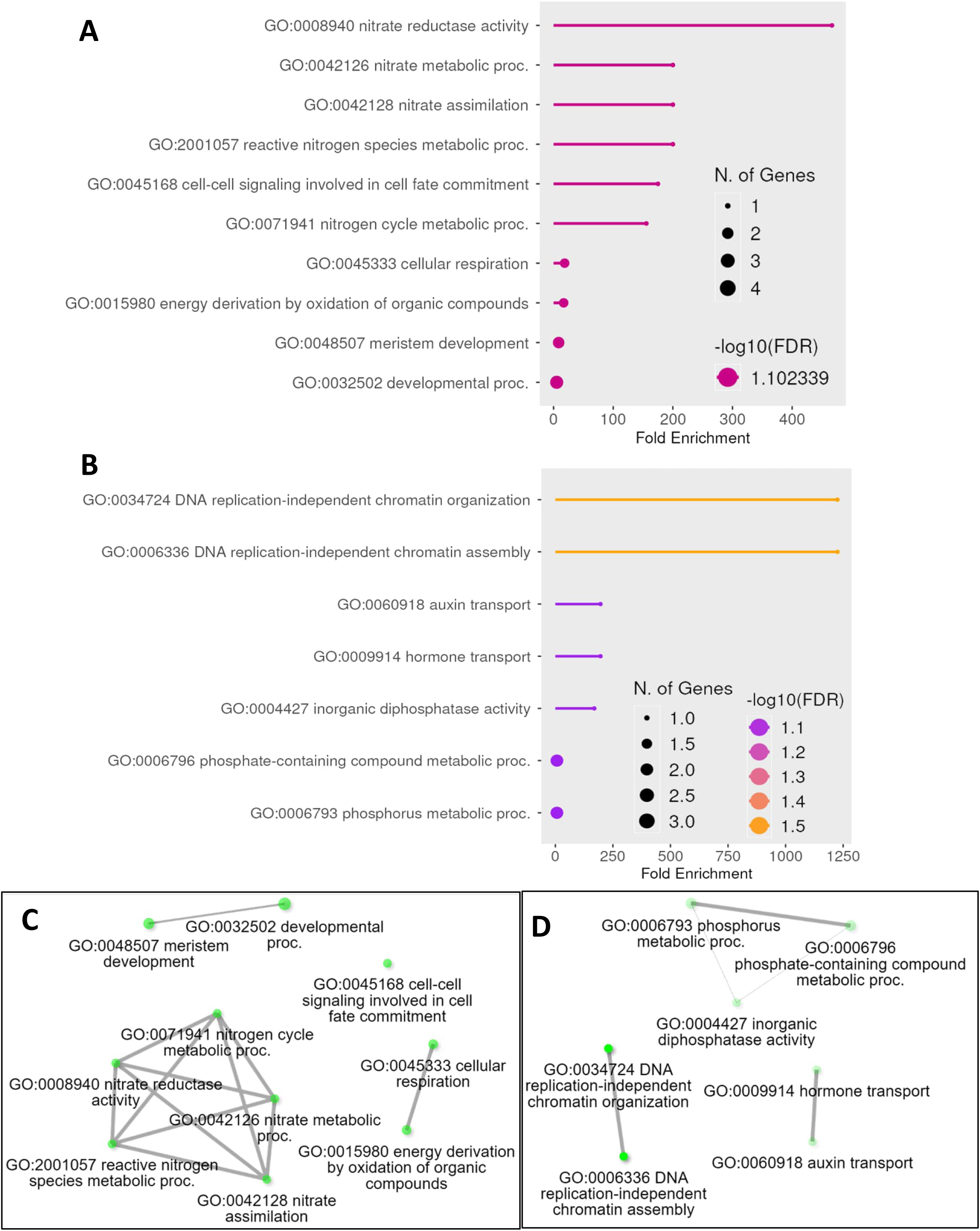
Gene enrichment analysis at T10 vs C10. (A) Line plot of downregulated genes. The x-axis indicates the fold-enrichment compared to controls at this stage, and the y-axis shows the gene ontology (GO) terms for significantly affected pathways. The closed circles connected to each line show gene numbers involved in the pathways. The color scales indicate –log10(FDR) for significance of the pathway expression. (B) Pathway clusters of downregulated genes. Pathways (nodes) are connected with edges to represent overlapped genes. Darker nodes are more significantly enriched gene sets. Larger nodes represent larger gene sets. Thicker edges represent more overlapped genes.

At T12, seven processes forming three functional clusters were downregulated (Figure 5B, D), the most strongly (1225-fold) for DNA replication-independent chromatin assembly and organization, which however comprised only two genes. The other two clusters concern metabolic processes involving phosphorus and related to auxin transport. Altogether 43 genes were upregulated, forming four clusters of pathways, the largest related to nitrogen metabolism (involving five processes), the others to meristem development and energy processes (Figure 5A, C).

At T22, while the stomata are already reopening, the significantly downregulated pathways included a single large cluster of 10 processes and 35 genes, with representation dropping from 16-to 115-fold (Figure S3B, D). The densely interconnected pathways involve cell wall metabolism and biosynthesis, including for the key structural components cellulose and beta-glucans. The upregulated processes at this stage form one large cluster related to nitrogen metabolism (fold-changes ranging 163 to 490), with two GO terms related to nitrate assimilation and nitrate reductase activity (NR) identical to those observed as upregulated at T12 (Figure 5, S3A, S3C). In addition, kinases (four-fold, 10 genes) and glutamate metabolism (approx. 200-fold, two genes) were upregulated.

The last time point, T42, where Gs had returned to near initial conditions, showed the greatest number of significantly differentially regulated genes (668) identified within pathways (Figure S4A, C, Table S14). The upregulated genes formed a single interconnected cluster of processes concerning protein phosphorylation and kinase activity, each involving over 60 genes and displaying eight-fold enrichment.

Notably, a set of interconnected pathways belonging to the photosynthetic machinery are downregulated at T42, even if gas exchange appears to be returning to normal (Figure S4B, D). One downregulated gene (1HG0064430.1, discussed for T2 above), which encodes chlorophyll a-b binding protein, is downregulated both at T2 and T42. Although the change in expression is about the same (36-fold at T2 vs 29-fold at T42), the relative expression is 63 times greater at T42 (a base mean of 9968 vs. 159).

## 4 Discussion

### 4.1 Sensitivity and acclimation of stomatal conductance to varying VPD level

Stomata serve as gateways for transpiration and photosynthetic gas exchange, the guard cells dynamically controlling stomatal aperture. Guard cells are able to do so by altering their shape in response to environmental factors including light, CO₂ concentration, and humidity (Merilo et al., 2013; Merilo et al. 2018). After an increase in VPD, stomata close, but then gradually reopen; we observed approximately 80% recovery of Gs by 12 minutes after VPD increase. A very similar stomatal response to high VPD in barley cv. Golden Promise was seen by Samantara et al. (2026); the authors propose that reopening of stomata under high VPD may be due to a fast active stomatal closure and increase in hydraulic conductance, which has been shown to accompany higher transpiration in well-watered conditions (Simonin et al. 2015). In Arabidopsis, guard cells respond to VPD somewhat more slowly, with a half-response time of six minutes (Merilo et al. 2018; Samantara et al. 2026); a range of response rates have been reported in the literature (Fletcher et al. 2007; Lawson & Vialet-Chabrand, 2019; Tamang et al. 2019).

In field-grown barley, even under well-watered conditions, our laboratory-scale results suggest that there could be continuous waves of stomatal closure and reopening during the course of the day as the air temperature rises and VPD increases. The rate and degree of Gs response and recovery would be expected to affect water use efficiency (WUE) and productivity. Indeed, in addition to VPD-triggered dynamic changes, Gs can also fluctuate under the influence of light variations, such as induced by changes in cloud cover or canopy shading. The response of stomata to these temporary changes can vary from minutes to hours (Liu et al. 2024). Moreover, environmental changes can occur more rapidly than the stomata can respond (Lawson & Vialet-Chabrand, 2019), a mismatch that may lead to temporal variations in water use efficiency. On a global scale, the effects of increasing VPD were shown to depress photosynthesis and transpiration (Li et al. 2023; López et al. 2021). Thus, dissecting rapid transcriptional reprogramming by VPD in crops is of growing importance.

### 4.2 Transcriptional response to VPD

As described above, barley stomata respond rapidly to an increase in VPD. However, we found that the leaf transcriptional response for this crop plant is even more rapid, being evident within two minutes of the VPD shift. It was earlier shown in the model plant Arabidopsis that a dynamic and layered expression of transcriptional regulators in response to abiotic stress may be found on different timescales, within seconds, minutes, or hours (Kollist et al. 2019). Transcriptional upregulation in response to light stress in Arabidopsis was found as early as 20 to 60 sec thereafter, with over 20% linked to H_2_O_2_ and ABA signalling (Suzuki et al. 2015).

Here, we found that among the DEGs, the statistically significant initial responses were predominantly downregulation. Many of the transcript pools showing a significant response to VPD in barley leaves are involved in photosynthesis or in drought regulatory networks for guard cells. At T2, the most strongly downregulated gene, at over 7.5-million-fold, was an *RLK*, *Lr10*. A rice *RLK* gene, *LP2*, was earlier shown to be downregulated by drought (Wu et al. 2015), though not virtually instantaneously as here. Of the other two highly rapid responders, the ribosomal protein S4 is novel to stress response. The third, LHCII, has been shown in Arabidopsis to be rapidly disassembled under drought (Chen et al. 2016b), but downregulation of its transcript is novel. At T12, a *CBL* was over 9-million-fold downregulated; in Arabidopsis, some *CBL* genes perceive calcium signals, responding to abiotic stress (Nagae et al, 2003; Shi et al. 1999; Wang et al. 2024). Likewise, *ATP1* and an *MT* gene were downregulated; in various systems, these are stress-responsive (Zhao et al. 2011; Kamal et al. 2012; Daurova et al. 2026). Gene enrichment analyses to uncover regulated biological pathways pointed to downregulation of photosynthesis and photosystems I and II at T2, and of nitrogen metabolism in particular at T12. Surprisingly, even though Gs recovered by T42, photosynthetic processes were still downregulated at that time point.

Upregulated genes at T12 included an *LRK10,* a mutant of which in wheat inhibited stomatal closure (Lim et al. 2015). *Cytb6f,* a di-heme cytochrome, was strongly upregulated at T12, as it was in Arabidopsis by ROS (Li et al. 2020). This cytochrome plays a vital role in transferring electrons and energy between PS II and PS I in the chloroplast (Hasan et al. 2013). Its upregulation under VPD suggests a role in enhancing the plant’s ability to detoxify ROS and maintain cellular redox homeostasis (Li et al. 2020). Also upregulated was *ABCG48*, a member of the ABC transporters. Some Arabidopsis ABC transporters modulate guard cell responses: mutants of AtMRP5 exhibit impaired guard cell function, affecting water use efficiency, while AtMRP4 mutants show decreased stomatal opening, contributing to enhanced drought tolerance (Li et al. 2021). These transporters influence ion and hormone transport including ABA export and import (Merilo et al., 2015), ensuring efficient stomatal regulation and preventing excessive water loss during abiotic stress, highlighting their importance in plant stress adaptation. The upregulation seen here is consistent with a role for ABA in stomatal closure under VPD (Zait et al. 2024) and for ABA in general leaf VPD response; ABA was shown to strongly increase the transcript levels of *ABCG31/PDR6* in rice leaves (Moons, 2008). The *ABCG* data is also consistent with the downregulation of processes involved in auxin transport at T12; ABA is known to downregulate auxin biosynthetic genes (Lorrai et al. 2018).

### 4.3 Short-term transcriptional dynamics

The majority of changes in the transcript pool at T2 involved small increments, with the median fold-change of the top 500 DEGs being 1.55 (down or up). The strongest decrease in pool size was 7.5-million-fold, while the largest increase (but with *p*adj 0.82) was only 46-fold. Removal of transcripts from the cellular pool by decapping, removal of the poly(A) tail, and then degradation can proceed very much more rapidly than synthesis of new ones (Walley et al. 2010). In wheat, drought stress leads to downregulation of more genes than are upregulated, which is linked as well to various microRNAs (miRNAs) involved in developmental processes (Gomez-Martin et al. 2023). This pattern reflects a broader stress response where mRNA levels are diminished as a first response to stress, often resulting in transcripts being directed to processing bodies (P-bodies) for decay (Huang, 2024). The role of P-bodies or stress granules in stress-responsive RNA turnover has been shown in Arabidopsis (Wang, 2024) and other plants (Huang, 2024).

RNA turnover has a demonstrated role in abiotic stress response in Arabidopsis. Gene-specific destabilisation of transcripts under light stress is followed by their stabilisation during recovery (Smith, 2024), the ten most rapidly turning over transcripts displaying half-lives of four to seven minutes. A prominent feature of the VPD response that we observed was stomatal reopening under high VPD, as evidenced by increasing Gs from T22 onward. Connected to this, we looked for transcripts detectable on the whole-leaf level that showed opposite abundance changes at T2 and T12, as for clues of genes that might be vanguards of stomatal reopening and equilibration following VPD increase.

Of the transcripts upregulated at T2 but downregulated at T12 with more than two-fold changes, one was annotated as encoding a UDP-glucose glycosyl transferase (UGT), which is involved in auxin metabolism. In Arabidopsis, UGT86A1 plays a positive regulatory role in drought resistance, promoting ROS scavenging capacity (Ma et al. 2025). The other (1HG0004660) was annotated as momilactone A synthase, an enzyme that synthesizes diterpenoid phytoalexins. In rice, a cluster of genes for this enzyme are inducible by both biotic and abiotic stresses (Shimura et al. 2007). Intriguingly, one group of products, labdane-related diterpenoid phytoalexins (LRDs), affect drought tolerance in maize (Vaughan et al. 2015). Knockout of LRD biosynthesis in rice inhibited stomatal closure in an ABA-independent manner; LRDs are proposed to serve as a regulatory switch triggering stomatal closure (Zhang et al. 2021). These results suggest a similar mechanism may operate in barley, consistent with the expression profile seen here for 1HG0004660 and with preparation at T12 for stomatal reopening shortly thereafter.

Of the two genes significantly downregulated at T2 but also upregulated at T12, one (1HG0004990) encodes a putative vacuolar invertase. In Arabidopsis, vacuolar invertase activity is comparatively high in guard cells and promotes stomatal opening (Chen et al. 2016a), as its cleavage of sucrose doubles osmotic potential, thereby increasing turgor pressure. Hence, its 6-fold drop at T2 and 16-fold rise at T12 is consistent with a key role in stomatal closure followed by equilibration to increased VPD and stomatal reopening. The other (1HG0004560), 28.5-fold down at T2, and 3.2-fold up at T12, is the CC-NLR immune receptor RGA5. In grapevine, RGA5 is induced not only by pathogens, but also by salicyclic acid (SA), jasmonic acid (JA), ABA, and H_2_O_2_ (Fan et al. 2015). Given the crosstalk between biotic and abiotic stress signalling for JA (Kazan & Manners, 2013), the uptick in the RGA5 pool at T12 may reflect a rebalancing of the role of JA in the MYC2 immunity signalling pathway away from the leaf state at stomatal closure (Ramirez-Prado et al. 2019), in anticipation of stomatal reopening.

### 4.4 VPD as an abiotic stress

Barley responds to decreasing soil water content over a period of days by eventual stomatal closure and transcriptional modulation, independent of the diurnal cycle, at a point depending on the plant variety and its drought response strategy (Paul, 2024). Here, we have demonstrated that barley leaves respond rapidly to a VPD increase with short-term stomatal closure and with changes in the transcript pool that overlap those of other (e.g. drought) stress responses, although the roots of high-VPD-treated plants are well watered. The prevailing view is that guard cells respond not to humidity but rather to the rate of transpirational water loss per se from the leaf; they sense neither the atmospheric water vapour concentration nor the difference in water vapour concentration between the leaf interior and the leaf surface (Monteith, 1995; Mott, 1991). More recently, experiments with ABA-insensitive Arabidopsis mutants have indicated the response to a rapid VPD shift, such as we imposed here, occurs in two phases, the first ABA-independent, the second ABA-dependent (Yaaran, 2019).

The transcriptional response data collected here is consistent with a biphasic VPD response mechanism. As discussed above, several genes with significant alterations in abundance are involved in ABA signalling and metabolism. In addition, we see by pathway enrichment analysis that nitrate reductase activity is 466-fold upregulated at T12, by 490-fold at T22. Moreover, ROS and reactive nitrogen species (RNS) interact closely in plant stress responses, with nitric oxide (NO) playing a pivotal role as a signalling molecule. In plants, NO synthesis involves NR under stress conditions (Hancock, 2019). NO interacts with ROS to modulate cellular processes, including redox homeostasis and signalling (Gogoi, 2024). During abiotic stress, NO regulates stomatal closure by integrating into the ABA signalling pathway, where it works together with ROS to induce guard cell responses. This dynamic interplay between ROS and RNS under stress conditions ensures a balanced response, enhancing the plant’s ability to adapt and survive adverse environmental conditions.

Notably, our transcriptional analysis was on whole leaves rather than guard cells alone. While the proportion of guard cells and the contribution of their RNA to the total leaf pool is unreported for barley, in Arabidopsis guard cells comprise 10.5 % of the total leaf cell number (Tolleter et al. 2024). If the proportion is the same in barley and likewise corresponds to RNA levels, transcriptional changes of less than 13.3-fold that occurred only in the guard cells would have been diluted to below the two-fold analysis threshold that we used. A likely example of a guard cell transcriptional response in our dataset is for 5HG0432480.1, corresponding to *ABCG48*, known in Arabidopsis to modulate guard cell response but showing only 2.8-fold upregulation at T12 (*p*adj 0.07). However, recent work in Arabidopsis indicates that in drought response, especially for mild drought and ABA treatment, whole-leaf and guard cell transcriptomes in any case overlap, with 48% of the upregulated genes and 59% of those downregulated being shared between treatments and samples (van Weringh et al. 2025).

On the whole-leaf level, strikingly, transcriptome changes were detectable within the first few minutes after a VPD increase, which fit well into the context of stomatal closure and consequent leaf photosynthetic and metabolic adjustment. Earlier, using decreased leaf turgor in Arabidopsis as a proxy for a VPD increase, the rapid (5 min) induction of the gene for 9-cis-epoxycarotenoid dioxygenase 3 (NCED3), which is rate-limiting for ABA biosynthesis, was demonstrated (Sussmilch et al. 2017). However, no published earlier results have addressed whole-transcriptome changes in the leaf on the short time scale examined here and not in a major crop such as barley. Taken together, our data suggest that the response to VPD involves large-scale and rapid adjustments extending from the transcriptional level to a multitude of biochemical pathways and physiological processes. Given that VPD is under constant flux over the course of the day, these layers of regulation are highly dynamic and the connections to water use efficiency, crop yield, and stress resilience are complex. Establishment of rapid cell-type-specific transcriptomics in barley using a method such as INTACT will help resolve the VPD response by cell function within the leaf (Debernardi et al. 2023; van Weringh et al. 2025).

## 5 Authors’ contributions

MP carried out the gas exchange experiments and expression assays, analysed the data, and wrote the manuscript; JT was responsible for bioinformatics analyses; EM carried out conceptualization, data curation and methodology; EM, HK, and AS designed the experiments; AS revised the manuscript and was project P.I. All authors contributed to the article.

## 7 Acknowledgements

We acknowledge the DNA Sequencing and Genomics Laboratory, Institute of Biotechnology, and the Genomics unit, Institute for Molecular Medicine Finland (FIMM), both belonging to the University of Helsinki, for the library preps and sequencing, respectively. We thank Anne-Mari Narvanto for her skilled technical assistance, as well as Drs. Triin Vahisalu and Alexey Shapiguzov for valuable insights.

## 8 Data availability statement

The data that support the findings of this study are available from the author upon reasonable request. The RNAseq data are available through the ENA and SRA databases in project ***.

## 9 Generative AI statement

The authors declare that no generative AI was used in the creation of this manuscript.

## 11 Supporting Information

Additional supporting information can be found online in the Supporting Information section.

**Figure S1** Gas exchange platform. The multi-cuvette system is shown during a typical experiment with well-watered barley plants and four collection timepoints being run simultaneously.

**Figure S2** Gene enrichment analysis at T2 vs C2. (A) Line plot of downregulated genes. The x-axis indicates the fold-enrichment compared to controls at this stage, and the y-axis shows the gene ontology (GO) terms for significantly affected pathways. The closed circles connected to each line show gene numbers involved in the pathways. The colour scales indicate –log10(FDR) for significance of the pathway expression. (B) Pathway clusters of downregulated genes. Pathways (nodes) are connected with edges to represent overlapped genes. Darker nodes are more significantly enriched gene sets. Larger nodes represent larger gene sets. Thicker edges represent more overlapped genes.

**Figure S3** Gene enrichment analysis at T22 vs C22. (A) Line plot of downregulated genes. The x-axis indicates the fold-enrichment compared to controls at this stage, and the y-axis shows the gene ontology (GO) terms for significantly affected pathways. The closed circles connected to each line show gene numbers involved in the pathways. The colour scales indicate –log10(FDR) for significance of the pathway expression. (B) Pathway clusters of downregulated genes. Pathways (nodes) are connected with edges to represent overlapped genes. Darker nodes are more significantly enriched gene sets. Larger nodes represent larger gene sets. Thicker edges represent more overlapped genes.

**Figure S4** Gene enrichment analysis at T42 vs C42. (A) Line plot of downregulated genes. The x-axis indicates the fold-enrichment compared to controls at this stage, and the y-axis shows the gene ontology (GO) terms for significantly affected pathways. The closed circles connected to each line show gene numbers involved in the pathways. The colour scales indicate –log10(FDR) for significance of the pathway expression. (B) Pathway clusters of downregulated genes. Pathways (nodes) are connected with edges to represent overlapped genes. Darker nodes are more significantly enriched gene sets. Larger nodes represent larger gene sets. Thicker edges represent more overlapped genes.

**Table S1** Genes upregulated at T2. The table columns include: (A) gene name in the Morex Version 2 reference genome assembly; (B) baseMean, the average of the normalized count values across all samples; (C) log2FoldChange, the effect size between the control and the sample for the gene in this row; (D) the standard error of the log2FC estimate; (E) Wald test statistic, calculated as log2FC/lfcSE, used to compute the *p*-value; (F) raw *p*-value from the Wald test; (G) adjusted *p*-value corrected for multiple hypothesis testing using the Benjamini-Hochberg procedure to control the false discovery rate (FDR).

**Table S2** Genes downregulated at T2. The table columns include: (A) gene name in the Morex Version 2 reference genome assembly; (B) baseMean, the average of the normalized count values across all samples; (C) log2FoldChange, the effect size between the control and the sample for the gene in this row; (D) the standard error of the log2FC estimate; (E) Wald test statistic, calculated as log2FC/lfcSE, used to compute the *p*-value; (F) raw *p*-value from the Wald test; (G) adjusted *p*-value corrected for multiple hypothesis testing using the Benjamini-Hochberg procedure to control the false discovery rate (FDR).

**Table S3** Genes upregulated at T12. The table columns include: (A) gene name in the Morex Version 2 reference genome assembly; (B) baseMean, the average of the normalized count values across all samples; (C) log2FoldChange, the effect size between the control and the sample for the gene in this row; (D) the standard error of the log2FC estimate; (E) Wald test statistic, calculated as log2FC/lfcSE, used to compute the *p*-value; (F) raw *p*-value from the Wald test; (G) adjusted *p*-value corrected for multiple hypothesis testing using the Benjamini-Hochberg procedure to control the false discovery rate (FDR).

**Table S4** Genes downregulated at T12. The table columns include: (A) gene name in the Morex Version 2 reference genome assembly; (B) baseMean, the average of the normalized count values across all samples; (C) log2FoldChange, the effect size between the control and the sample for the gene in this row; (D) the standard error of the log2FC estimate; (E) Wald test statistic, calculated as log2FC/lfcSE, used to compute the *p*-value; (F) raw *p*-value from the Wald test; (G) adjusted *p*-value corrected for multiple hypothesis testing using the Benjamini-Hochberg procedure to control the false discovery rate (FDR).

**Table S5** Genes upregulated at T22. The table columns include: (A) gene name in the Morex Version 2 reference genome assembly; (B) baseMean, the average of the normalized count values across all samples; (C) log2FoldChange, the effect size between the control and the sample for the gene in this row; (D) the standard error of the log2FC estimate; (E) Wald test statistic, calculated as log2FC/lfcSE, used to compute the *p*-value; (F) raw *p*-value from the Wald test; (G) adjusted *p*-value corrected for multiple hypothesis testing using the Benjamini-Hochberg procedure to control the false discovery rate (FDR).

**Table S6** Genes downregulated at T22. The table columns include: (A) gene name in the Morex Version 2 reference genome assembly; (B) baseMean, the average of the normalized count values across all samples; (C) log2FoldChange, the effect size between the control and the sample for the gene in this row; (D) the standard error of the log2FC estimate; (E) Wald test statistic, calculated as log2FC/lfcSE, used to compute the *p*-value; (F) raw *p*-value from the Wald test; (G) adjusted *p*-value corrected for multiple hypothesis testing using the Benjamini-Hochberg procedure to control the false discovery rate (FDR).

**Table S7** Genes upregulated at T42. The table columns include: (A) gene name in the Morex Version 2 reference genome assembly; (B) baseMean, the average of the normalized count values across all samples; (C) log2FoldChange, the effect size between the control and the sample for the gene in this row; (D) the standard error of the log2FC estimate; (E) Wald test statistic, calculated as log2FC/lfcSE, used to compute the *p*-value; (F) raw *p*-value from the Wald test; (G) adjusted *p*-value corrected for multiple hypothesis testing using the Benjamini-Hochberg procedure to control the false discovery rate (FDR).

**Table S8** Genes downregulated at T42. The table columns include: (A) gene name in the Morex Version 2 reference genome assembly; (B) baseMean, the average of the normalized count values across all samples; (C) log2FoldChange, the effect size between the control and the sample for the gene in this row; (D) the standard error of the log2FC estimate; (E) Wald test statistic, calculated as log2FC/lfcSE, used to compute the *p*-value; (F) raw *p*-value from the Wald test; (G) adjusted *p*-value corrected for multiple hypothesis testing using the Benjamini-Hochberg procedure to control the false discovery rate (FDR).

**Table S9** Pathways downregulated at T2 vs the control C2. Columns: (A) Enrichment FDR, False Discovery Rate, the adjusted *p*-value for false-positive results among the significant pathways identified as enriched; (B) number of genes identified as enriched for the pathway; (C) number of genes in the pathway; (D) fold enrichment (E) Pathway, including Gene Ontology ID and process term; (F) gene names in the Morex Version 2 reference genome assembly.

**Table S10** Pathways upregulated at T12 vs the control C12. Columns: (A) Enrichment FDR, False Discovery Rate, the adjusted *p*-value for false-positive results among the significant pathways identified as enriched; (B) number of genes identified as enriched for the pathway; (C) number of genes in the pathway; (D) fold enrichment (E) Pathway, including Gene Ontology ID and process term; (F) gene names in the Morex Version 2 reference genome assembly.

**Table S11** Pathways downregulated at T12 vs the control C12. Columns: (A) Enrichment FDR, False Discovery Rate, the adjusted *p*-value for false-positive results among the significant pathways identified as enriched; (B) number of genes identified as enriched for the pathway; (C) number of genes in the pathway; (D) fold enrichment (E) Pathway, including Gene Ontology ID and process term; (F) gene names in the Morex Version 2 reference genome assembly.

**Table S12** Pathways upregulated at T22 vs the control C12. Columns: (A) Enrichment FDR, False Discovery Rate, the adjusted *p*-value for false-positive results among the significant pathways identified as enriched; (B) number of genes identified as enriched for the pathway; (C) number of genes in the pathway; (D) fold enrichment (E) Pathway, including Gene Ontology ID and process term; (F) gene names in the Morex Version 2 reference genome assembly.

**Table S13** Pathways downregulated at T22 vs the control C22. Columns: (A) Enrichment FDR, False Discovery Rate, the adjusted *p*-value for false-positive results among the significant pathways identified as enriched; (B) number of genes identified as enriched for the pathway; (C) number of genes in the pathway; (D) fold enrichment (E) Pathway, including Gene Ontology ID and process term; (F) gene names in the Morex Version 2 reference genome assembly.

**Table S14** Pathways upregulated at T42 vs the control C42. Columns: (A) Enrichment FDR, False Discovery Rate, the adjusted *p*-value for false-positive results among the significant pathways identified as enriched; (B) number of genes identified as enriched for the pathway; (C) number of genes in the pathway; (D) fold enrichment (E) Pathway, including Gene Ontology ID and process term; (F) gene names in the Morex Version 2 reference genome assembly.

**Table S15** Pathways downregulated at T12 vs the control C12. Columns: (A) Enrichment FDR, False Discovery Rate, the adjusted *p*-value for false-positive results among the significant pathways identified as enriched; (B) number of genes identified as enriched for the pathway; (C) number of genes in the pathway; (D) fold enrichment (E) Pathway, including Gene Ontology ID and process term; (F) gene names in the Morex Version 2 reference genome assembly.

