## Supplementary Figures for "Response of barley (*Hordeum vulgare*) leaves to vapour pressure deficit (VPD) involves rapid shifts in transcript pools"

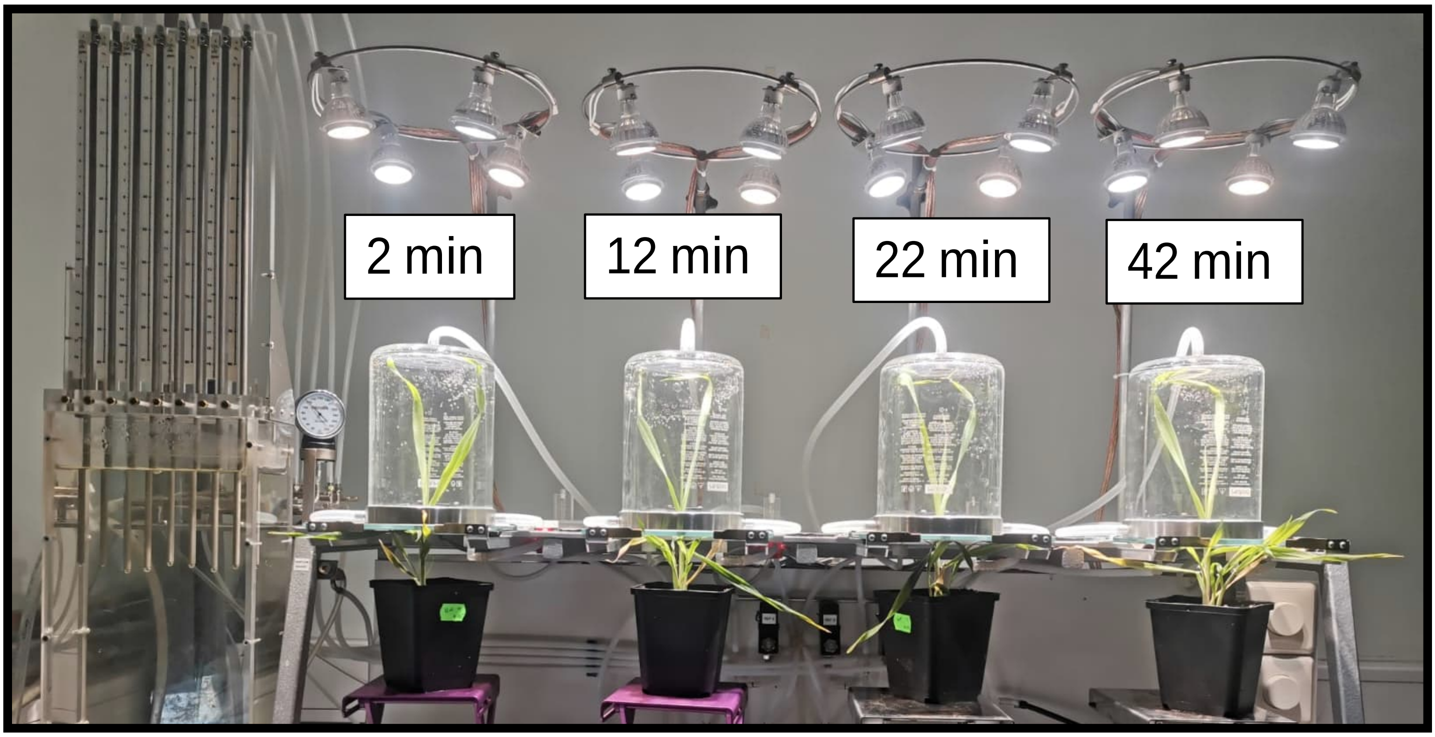


Figure S1. Gas exchange platform. The multi-cuvette system is shown during a typical experiment with well-watered barley plants and four collection timepoints being run simultaneously.

A

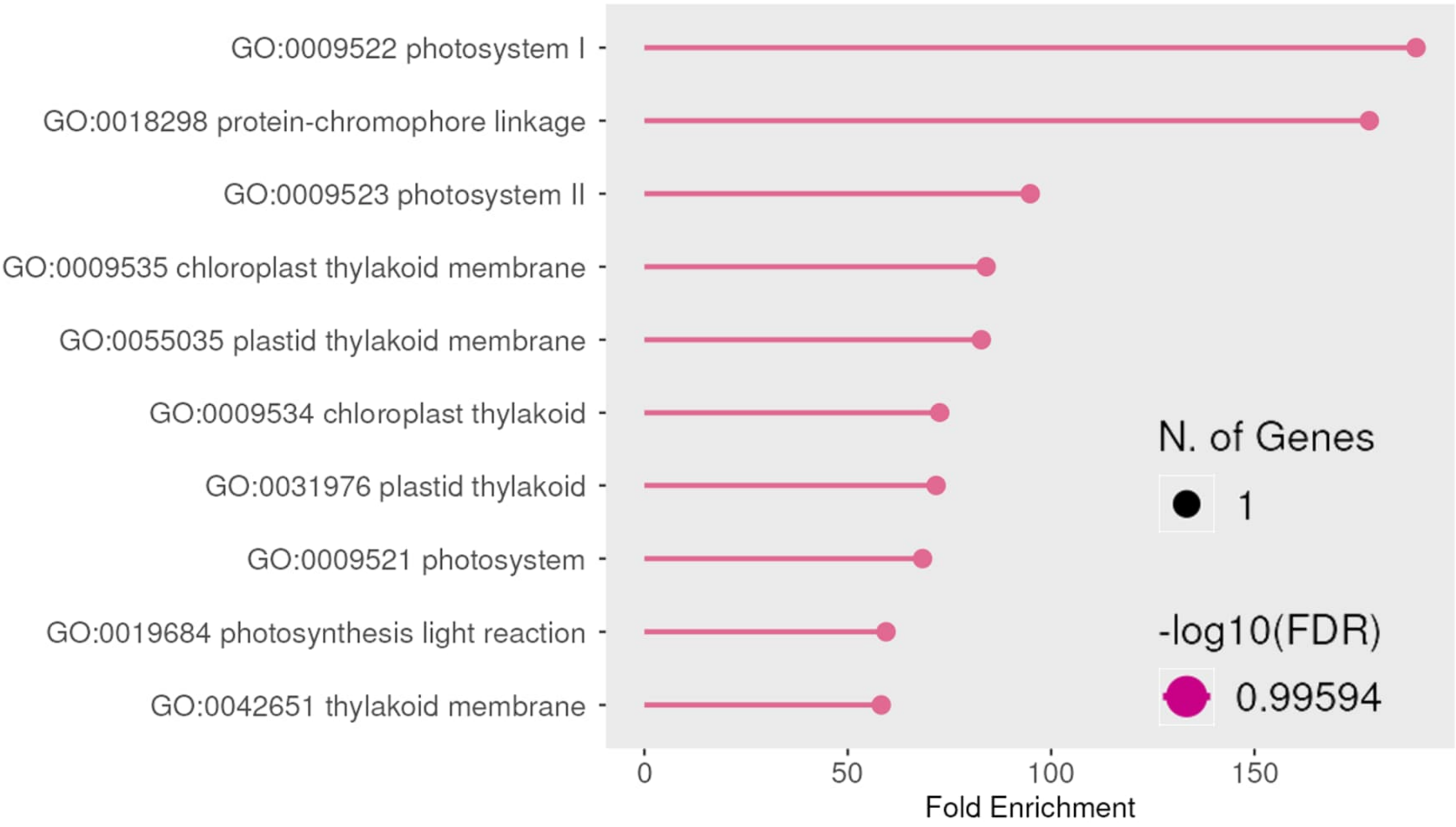


B

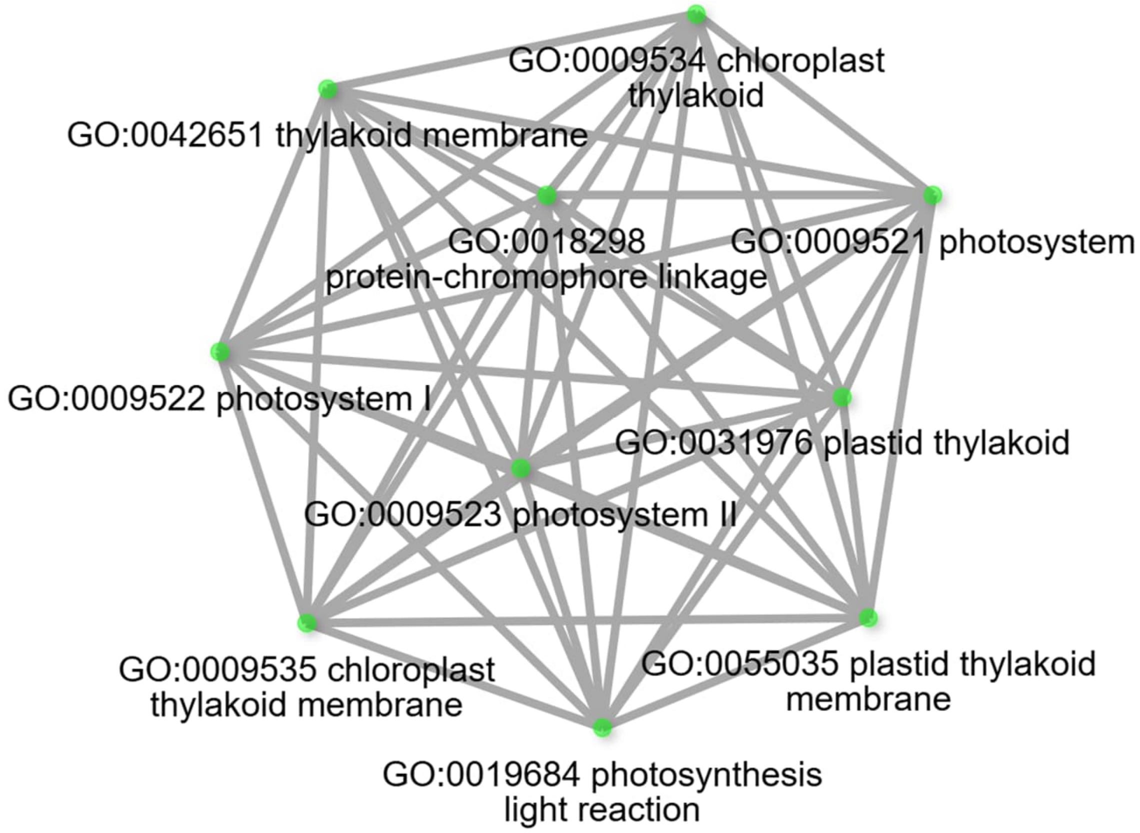


Figure S2. Gene enrichment analysis at T2 vs C2. (A) Line plot of downregulated genes. The x-axis indicates the fold-enrichment compared to controls at this stage, and the y-axis shows the gene ontology (GO) terms for significantly affected pathways. The closed circles connected to each line show gene numbers involved in the pathways. The colour scales indicate –log10(FDR) for significance of the pathway expression. (B) Pathway clusters of downregulated genes. Pathways (nodes) are connected with edges to represent overlapped genes. Darker nodes are more significantly enriched gene sets. Larger nodes represent larger gene sets. Thicker edges represent more overlapped genes.

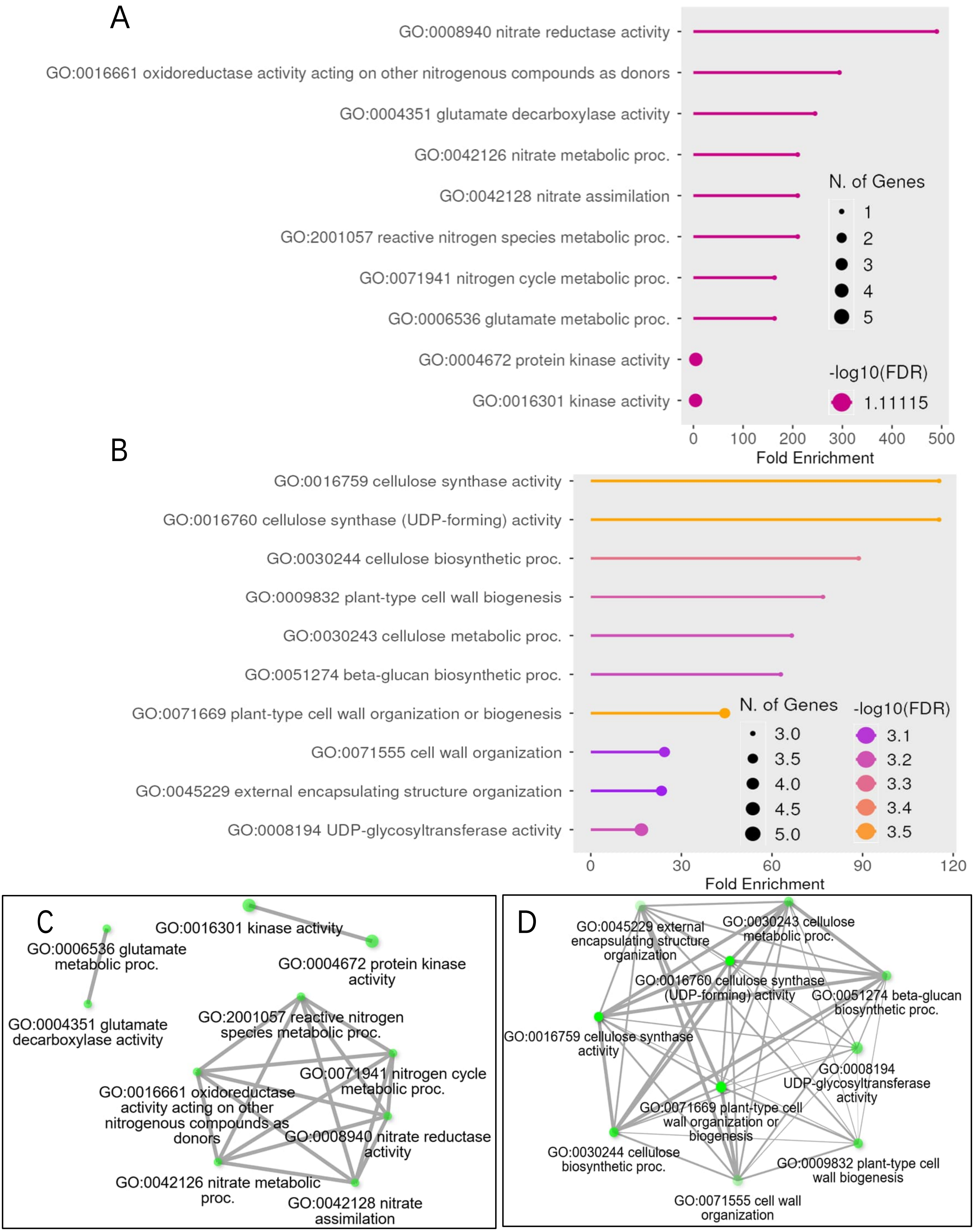


Figure S3. Gene enrichment analysis at T22 vs C22. (A) Line plot of downregulated genes. The x-axis indicates the fold-enrichment compared to controls at this stage, and the y-axis shows the gene ontology (GO) terms for significantly affected pathways. The closed circles connected to each line show gene numbers involved in the pathways. The colour scales indicate –log10(FDR) for significance of the pathway expression. (B) Pathway clusters of downregulated genes. Pathways (nodes) are connected with edges to represent overlapped genes. Darker nodes are more significantly enriched gene sets. Larger nodes represent larger gene sets. Thicker edges represent more overlapped genes.

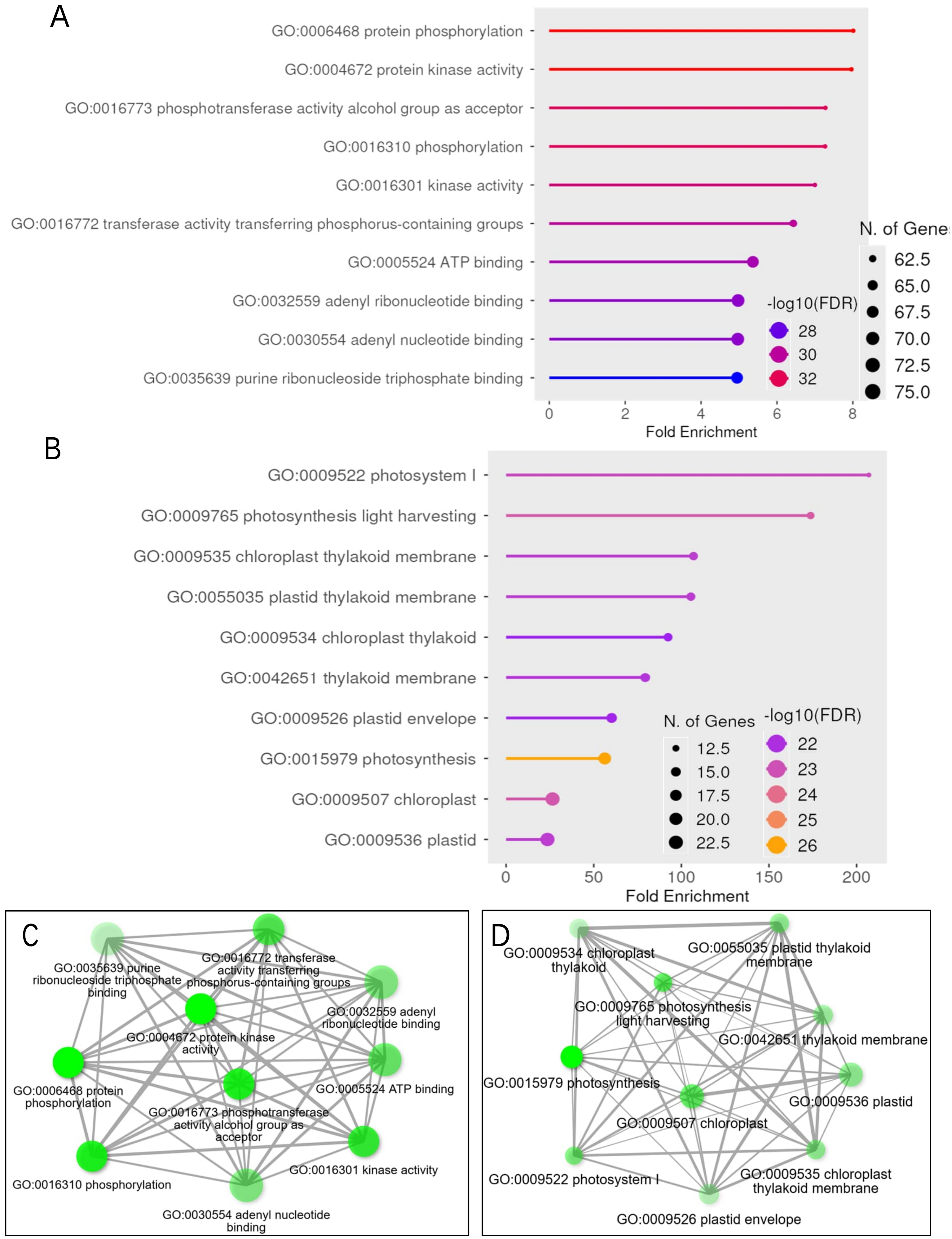


Figure S4. Gene enrichment analysis at T42 vs C42. (A) Line plot of downregulated genes. The x-axis indicates the fold-enrichment compared to controls at this stage, and the y-axis shows the gene ontology (GO) terms for significantly affected pathways. The closed circles connected to each line show gene numbers involved in the pathways. The colour scales indicate –log10(FDR) for significance of the pathway expression. (B) Pathway clusters of downregulated genes. Pathways (nodes) are connected with edges to represent overlapped genes. Darker nodes are more significantly enriched gene sets. Larger nodes represent larger gene sets. Thicker edges represent more overlapped genes.
