## Supplementary Tables for "Response of barley (*Hordeum vulgare*) leaves to vapour pressure deficit (VPD) involves rapid shifts in transcript pools"

Table S1

| Upregulated genes at T2 | baseMean | log2FC | lfcSE | stat | pvalue | padj |
|---|---|---|---|---|---|---|
| HORVU.MOREX.r2.1HG0028700.1 | 9.190 | 5.518 | 1.411 | 3.910 | 0.000 | 0.812 |
| HORVU.MOREX.r2.1HG0003300.1 | 5.987 | 3.010 | 1.719 | 1.751 | 0.080 | 1.000 |
| HORVU.MOREX.r2.1HG0006880.1 | 4.322 | 2.027 | 1.263 | 1.605 | 0.108 | 1.000 |
| HORVU.MOREX.r2.1HG0002460.1 | 10.405 | 1.358 | 0.936 | 1.450 | 0.147 | 1.000 |
| HORVU.MOREX.r2.1HG0007840.1 | 40.549 | 1.849 | 1.333 | 1.387 | 0.165 | 1.000 |
| HORVU.MOREX.r2.1HG0007220.1 | 30.332 | 1.253 | 0.988 | 1.269 | 0.204 | 1.000 |
| HORVU.MOREX.r2.1HG0001870.1 | 616.937 | 0.422 | 0.336 | 1.256 | 0.209 | 1.000 |
| HORVU.MOREX.r2.1HG0003640.1 | 2.926 | 1.984 | 1.582 | 1.254 | 0.210 | 1.000 |
| HORVU.MOREX.r2.1HG0007850.1 | 7.513 | 0.958 | 0.894 | 1.071 | 0.284 | 1.000 |
| HORVU.MOREX.r2.1HG0001430.1 | 223.201 | 0.383 | 0.387 | 0.992 | 0.321 | 1.000 |
| HORVU.MOREX.r2.1HG0006870.1 | 4.584 | 1.553 | 1.629 | 0.954 | 0.340 | 1.000 |
| HORVU.MOREX.r2.1HG0002070.1 | 1.473 | 2.247 | 2.363 | 0.951 | 0.342 | 1.000 |
| HORVU.MOREX.r2.1HG0002140.1 | 89.363 | 0.386 | 0.435 | 0.888 | 0.375 | 1.000 |
| HORVU.MOREX.r2.1HG0004660.1 | 1.613 | 1.840 | 2.078 | 0.885 | 0.376 | 1.000 |
| HORVU.MOREX.r2.1HG0003650.1 | 1.632 | 1.516 | 1.735 | 0.874 | 0.382 | 1.000 |
| HORVU.MOREX.r2.1HG0004250.1 | 163.196 | 0.654 | 0.752 | 0.869 | 0.385 | 1.000 |
| HORVU.MOREX.r2.1HG0002350.1 | 54.474 | 0.394 | 0.467 | 0.844 | 0.399 | 1.000 |
| HORVU.MOREX.r2.1HG0007120.1 | 275.760 | 0.802 | 0.974 | 0.824 | 0.410 | 1.000 |
| HORVU.MOREX.r2.1HG0004680.1 | 207.751 | 0.278 | 0.348 | 0.799 | 0.424 | 1.000 |
| HORVU.MOREX.r2.1HG0000830.1 | 1.524 | 1.682 | 2.132 | 0.789 | 0.430 | 1.000 |
| HORVU.MOREX.r2.1HG0004260.1 | 35.100 | 0.735 | 0.950 | 0.773 | 0.440 | 1.000 |
| HORVU.MOREX.r2.1HG0007050.1 | 5.770 | 0.976 | 1.279 | 0.763 | 0.445 | 1.000 |
| HORVU.MOREX.r2.1HG0005390.1 | 5.021 | 0.810 | 1.061 | 0.763 | 0.445 | 1.000 |
| HORVU.MOREX.r2.1HG0002390.1 | 15.526 | 1.070 | 1.446 | 0.740 | 0.459 | 1.000 |
| HORVU.MOREX.r2.1HG0003400.1 | 63.539 | 0.447 | 0.606 | 0.737 | 0.461 | 1.000 |
| HORVU.MOREX.r2.1HG0006820.1 | 51.230 | 0.436 | 0.598 | 0.730 | 0.466 | 1.000 |
| HORVU.MOREX.r2.1HG0003500.1 | 1198.884 | 0.199 | 0.277 | 0.721 | 0.471 | 1.000 |
| HORVU.MOREX.r2.1HG0004060.1 | 689.873 | 0.192 | 0.271 | 0.709 | 0.478 | 1.000 |
| HORVU.MOREX.r2.1HG0004230.1 | 100.935 | 0.286 | 0.407 | 0.703 | 0.482 | 1.000 |
| HORVU.MOREX.r2.1HG0000210.1 | 0.804 | 2.538 | 3.760 | 0.675 | 0.500 | 1.000 |
| HORVU.MOREX.r2.1HG0005820.1 | 1.355 | 1.796 | 2.841 | 0.632 | 0.527 | 1.000 |
| HORVU.MOREX.r2.1HG0001690.1 | 245.481 | 0.209 | 0.337 | 0.621 | 0.535 | 1.000 |
| HORVU.MOREX.r2.1HG0006520.1 | 1.379 | 2.223 | 3.637 | 0.611 | 0.541 | 1.000 |
| HORVU.MOREX.r2.1HG0001720.1 | 319.702 | 0.158 | 0.264 | 0.598 | 0.550 | 1.000 |
| HORVU.MOREX.r2.1HG0004240.1 | 2.019 | 0.994 | 1.748 | 0.569 | 0.570 | 1.000 |
| HORVU.MOREX.r2.1HG0002020.1 | 6994.222 | 0.219 | 0.386 | 0.566 | 0.571 | 1.000 |
| HORVU.MOREX.r2.1HG0006570.1 | 7.178 | 0.605 | 1.077 | 0.562 | 0.574 | 1.000 |
| HORVU.MOREX.r2.1HG0005360.1 | 135.681 | 0.234 | 0.417 | 0.560 | 0.576 | 1.000 |
| HORVU.MOREX.r2.1HG0002120.1 | 1.624 | 1.103 | 1.973 | 0.559 | 0.576 | 1.000 |
| HORVU.MOREX.r2.1HG0007880.1 | 6.032 | 0.935 | 1.677 | 0.557 | 0.577 | 1.000 |
| HORVU.MOREX.r2.1HG0007370.1 | 4.001 | 0.801 | 1.446 | 0.554 | 0.580 | 1.000 |
| HORVU.MOREX.r2.1HG0007650.1 | 4.745 | 0.937 | 1.697 | 0.552 | 0.581 | 1.000 |
| HORVU.MOREX.r2.1HG0006620.1 | 1499.955 | 0.150 | 0.272 | 0.552 | 0.581 | 1.000 |
| HORVU.MOREX.r2.1HG0002730.1 | 0.732 | 2.001 | 3.730 | 0.537 | 0.592 | 1.000 |
| HORVU.MOREX.r2.1HG0003910.1 | 1182.881 | 0.329 | 0.623 | 0.528 | 0.597 | 1.000 |
| HORVU.MOREX.r2.1HG0001470.1 | 3.628 | 0.646 | 1.235 | 0.523 | 0.601 | 1.000 |
| HORVU.MOREX.r2.1HG0005680.1 | 1.066 | 1.947 | 3.788 | 0.514 | 0.607 | 1.000 |
| HORVU.MOREX.r2.1HG0003210.1 | 307.565 | 0.154 | 0.300 | 0.513 | 0.608 | 1.000 |
| HORVU.MOREX.r2.1HG0000280.1 | 2456.313 | 0.188 | 0.374 | 0.504 | 0.615 | 1.000 |
| HORVU.MOREX.r2.1HG0004430.1 | 30.950 | 0.334 | 0.681 | 0.490 | 0.624 | 1.000 |
| HORVU.MOREX.r2.1HG0004960.1 | 37.301 | 0.256 | 0.560 | 0.457 | 0.648 | 1.000 |
| HORVU.MOREX.r2.1HG0006020.1 | 6.399 | 0.440 | 0.968 | 0.455 | 0.649 | 1.000 |

| | | | | | | |
|---|---|---|---|---|---|---|
| HORVU.MOREX.r2.1HG0000900.1 | 1.488 | 1.373 | 3.074 | 0.447 | 0.655 | 1.000 |
| HORVU.MOREX.r2.1HG0005050.1 | 68.015 | 0.178 | 0.408 | 0.436 | 0.663 | 1.000 |
| HORVU.MOREX.r2.1HG0002750.1 | 386.100 | 0.162 | 0.389 | 0.417 | 0.677 | 1.000 |
| HORVU.MOREX.r2.1HG0007670.1 | 539.946 | 0.170 | 0.408 | 0.416 | 0.678 | 1.000 |
| HORVU.MOREX.r2.1HG0001460.1 | 1.853 | 1.417 | 3.473 | 0.408 | 0.683 | 1.000 |
| HORVU.MOREX.r2.1HG0004410.1 | 25.290 | 0.273 | 0.678 | 0.403 | 0.687 | 1.000 |
| HORVU.MOREX.r2.1HG0007560.1 | 4.864 | 0.683 | 1.698 | 0.402 | 0.688 | 1.000 |
| HORVU.MOREX.r2.1HG0003890.1 | 10.645 | 1.399 | 3.529 | 0.396 | 0.692 | 1.000 |
| HORVU.MOREX.r2.1HG0000050.1 | 211.647 | 0.228 | 0.576 | 0.396 | 0.692 | 1.000 |
| HORVU.MOREX.r2.1HG0003970.1 | 0.516 | 1.478 | 3.763 | 0.393 | 0.694 | 1.000 |
| HORVU.MOREX.r2.1HG0005810.1 | 1264.182 | 0.104 | 0.268 | 0.389 | 0.698 | 1.000 |
| HORVU.MOREX.r2.1HG0000700.1 | 216.979 | 0.115 | 0.296 | 0.387 | 0.699 | 1.000 |
| HORVU.MOREX.r2.1HG0001130.1 | 0.264 | 1.470 | 3.810 | 0.386 | 0.700 | 1.000 |
| HORVU.MOREX.r2.1HG0003820.1 | 0.264 | 1.470 | 3.810 | 0.386 | 0.700 | 1.000 |
| HORVU.MOREX.r2.1HG0005720.1 | 0.264 | 1.470 | 3.810 | 0.386 | 0.700 | 1.000 |
| HORVU.MOREX.r2.1HG0002830.1 | 1.262 | 1.062 | 2.757 | 0.385 | 0.700 | 1.000 |
| HORVU.MOREX.r2.1HG0007540.1 | 5.206 | 0.598 | 1.552 | 0.385 | 0.700 | 1.000 |
| HORVU.MOREX.r2.1HG0004860.1 | 0.362 | 1.464 | 3.810 | 0.384 | 0.701 | 1.000 |
| HORVU.MOREX.r2.1HG0000170.1 | 0.662 | 1.461 | 3.811 | 0.383 | 0.701 | 1.000 |
| HORVU.MOREX.r2.1HG0001020.1 | 116.544 | 0.174 | 0.454 | 0.383 | 0.702 | 1.000 |
| HORVU.MOREX.r2.1HG0006740.1 | 0.716 | 1.417 | 3.716 | 0.381 | 0.703 | 1.000 |
| HORVU.MOREX.r2.1HG0003810.1 | 1.575 | 1.385 | 3.684 | 0.376 | 0.707 | 1.000 |
| HORVU.MOREX.r2.1HG0005450.1 | 0.236 | 1.419 | 3.802 | 0.373 | 0.709 | 1.000 |
| HORVU.MOREX.r2.1HG0001220.1 | 0.236 | 1.418 | 3.802 | 0.373 | 0.709 | 1.000 |
| HORVU.MOREX.r2.1HG0006180.1 | 7.783 | 0.376 | 1.009 | 0.373 | 0.709 | 1.000 |
| HORVU.MOREX.r2.1HG0007240.1 | 0.335 | 1.411 | 3.812 | 0.370 | 0.711 | 1.000 |
| HORVU.MOREX.r2.1HG0003220.1 | 105.459 | 0.137 | 0.375 | 0.367 | 0.714 | 1.000 |
| HORVU.MOREX.r2.1HG0002870.1 | 5.192 | 0.427 | 1.168 | 0.366 | 0.715 | 1.000 |
| HORVU.MOREX.r2.1HG0002410.1 | 137.535 | 0.129 | 0.358 | 0.359 | 0.720 | 1.000 |
| HORVU.MOREX.r2.1HG0002840.1 | 1293.912 | 0.101 | 0.285 | 0.355 | 0.723 | 1.000 |
| HORVU.MOREX.r2.1HG0006900.1 | 4.462 | 0.592 | 1.703 | 0.348 | 0.728 | 1.000 |
| HORVU.MOREX.r2.1HG0003350.1 | 0.995 | 1.151 | 3.349 | 0.344 | 0.731 | 1.000 |
| HORVU.MOREX.r2.1HG0006400.1 | 1837.108 | 0.087 | 0.255 | 0.339 | 0.735 | 1.000 |
| HORVU.MOREX.r2.1HG0002790.1 | 12.719 | 0.240 | 0.715 | 0.335 | 0.738 | 1.000 |
| HORVU.MOREX.r2.1HG0005570.1 | 3521.646 | 0.118 | 0.361 | 0.328 | 0.743 | 1.000 |
| HORVU.MOREX.r2.1HG0003630.1 | 61.815 | 0.231 | 0.722 | 0.320 | 0.749 | 1.000 |
| HORVU.MOREX.r2.1HG0004420.1 | 104.741 | 0.252 | 0.823 | 0.306 | 0.759 | 1.000 |
| HORVU.MOREX.r2.1HG0007530.1 | 132.495 | 0.639 | 2.169 | 0.295 | 0.768 | 1.000 |
| HORVU.MOREX.r2.1HG0003680.1 | 0.193 | 1.114 | 3.817 | 0.292 | 0.770 | 1.000 |
| HORVU.MOREX.r2.1HG0005440.1 | 28.771 | 0.182 | 0.671 | 0.271 | 0.786 | 1.000 |
| HORVU.MOREX.r2.1HG0005030.1 | 3.168 | 0.719 | 2.662 | 0.270 | 0.787 | 1.000 |
| HORVU.MOREX.r2.1HG0003770.1 | 458.532 | 0.086 | 0.327 | 0.262 | 0.794 | 1.000 |
| HORVU.MOREX.r2.1HG0000190.1 | 0.210 | 0.985 | 3.805 | 0.259 | 0.796 | 1.000 |
| HORVU.MOREX.r2.1HG0001100.1 | 0.210 | 0.985 | 3.805 | 0.259 | 0.796 | 1.000 |
| HORVU.MOREX.r2.1HG0005580.1 | 0.210 | 0.985 | 3.805 | 0.259 | 0.796 | 1.000 |
| HORVU.MOREX.r2.1HG0006150.1 | 0.210 | 0.985 | 3.805 | 0.259 | 0.796 | 1.000 |
| HORVU.MOREX.r2.1HG0007210.1 | 0.210 | 0.985 | 3.805 | 0.259 | 0.796 | 1.000 |
| HORVU.MOREX.r2.1HG0000240.1 | 47.283 | 0.850 | 3.325 | 0.256 | 0.798 | 1.000 |
| HORVU.MOREX.r2.1HG0000070.1 | 557.632 | 0.076 | 0.299 | 0.253 | 0.800 | 1.000 |
| HORVU.MOREX.r2.1HG0001900.1 | 0.794 | 0.659 | 2.674 | 0.246 | 0.805 | 1.000 |
| HORVU.MOREX.r2.1HG0005340.1 | 1770.925 | 0.118 | 0.494 | 0.239 | 0.811 | 1.000 |
| HORVU.MOREX.r2.1HG0000610.1 | 0.357 | 0.897 | 3.772 | 0.238 | 0.812 | 1.000 |
| HORVU.MOREX.r2.1HG0002960.1 | 0.425 | 0.894 | 3.778 | 0.237 | 0.813 | 1.000 |

| | | | | | | |
|---|---|---|---|---|---|---|
| HORVU.MOREX.r2.1HG0000890.1 | 0.204 | 0.889 | 3.821 | 0.233 | 0.816 | 1.000 |
| HORVU.MOREX.r2.1HG0001200.1 | 0.413 | 0.872 | 3.778 | 0.231 | 0.817 | 1.000 |
| HORVU.MOREX.r2.1HG0003560.1 | 0.409 | 0.850 | 3.778 | 0.225 | 0.822 | 1.000 |
| HORVU.MOREX.r2.1HG0000090.1 | 1313.276 | 0.076 | 0.350 | 0.218 | 0.828 | 1.000 |
| HORVU.MOREX.r2.1HG0005400.1 | 273.511 | 0.117 | 0.547 | 0.215 | 0.830 | 1.000 |
| HORVU.MOREX.r2.1HG0002190.1 | 694.570 | 0.076 | 0.367 | 0.207 | 0.836 | 1.000 |
| HORVU.MOREX.r2.1HG0003130.1 | 0.908 | 0.736 | 3.725 | 0.198 | 0.843 | 1.000 |
| HORVU.MOREX.r2.1HG0005800.1 | 873.705 | 0.049 | 0.259 | 0.188 | 0.851 | 1.000 |
| HORVU.MOREX.r2.1HG0000040.1 | 528.106 | 0.075 | 0.411 | 0.183 | 0.855 | 1.000 |
| HORVU.MOREX.r2.1HG0000350.1 | 14.288 | 0.141 | 0.776 | 0.181 | 0.856 | 1.000 |
| HORVU.MOREX.r2.1HG0006190.1 | 2.953 | 0.250 | 1.469 | 0.170 | 0.865 | 1.000 |
| HORVU.MOREX.r2.1HG0005850.1 | 1103.994 | 0.103 | 0.604 | 0.170 | 0.865 | 1.000 |
| HORVU.MOREX.r2.1HG0001160.1 | 2590.931 | 0.219 | 1.319 | 0.166 | 0.868 | 1.000 |
| HORVU.MOREX.r2.1HG0000180.1 | 0.105 | 0.628 | 3.821 | 0.164 | 0.869 | 1.000 |
| HORVU.MOREX.r2.1HG0002990.1 | 0.105 | 0.628 | 3.821 | 0.164 | 0.869 | 1.000 |
| HORVU.MOREX.r2.1HG0005280.1 | 0.105 | 0.628 | 3.821 | 0.164 | 0.869 | 1.000 |
| HORVU.MOREX.r2.1HG0007200.1 | 0.105 | 0.628 | 3.821 | 0.164 | 0.869 | 1.000 |
| HORVU.MOREX.r2.1HG0003790.1 | 0.332 | 0.624 | 3.826 | 0.163 | 0.870 | 1.000 |
| HORVU.MOREX.r2.1HG0005980.1 | 0.232 | 0.612 | 3.827 | 0.160 | 0.873 | 1.000 |
| HORVU.MOREX.r2.1HG0000030.1 | 1207.210 | 0.077 | 0.484 | 0.160 | 0.873 | 1.000 |
| HORVU.MOREX.r2.1HG0000430.1 | 0.132 | 0.605 | 3.827 | 0.158 | 0.874 | 1.000 |
| HORVU.MOREX.r2.1HG0001120.1 | 0.136 | 0.605 | 3.827 | 0.158 | 0.874 | 1.000 |
| HORVU.MOREX.r2.1HG0003880.1 | 0.132 | 0.605 | 3.827 | 0.158 | 0.874 | 1.000 |
| HORVU.MOREX.r2.1HG0004020.1 | 0.132 | 0.605 | 3.827 | 0.158 | 0.874 | 1.000 |
| HORVU.MOREX.r2.1HG0005230.1 | 0.132 | 0.605 | 3.827 | 0.158 | 0.874 | 1.000 |
| HORVU.MOREX.r2.1HG0005260.1 | 0.132 | 0.605 | 3.827 | 0.158 | 0.874 | 1.000 |
| HORVU.MOREX.r2.1HG0006330.1 | 0.132 | 0.605 | 3.827 | 0.158 | 0.874 | 1.000 |
| HORVU.MOREX.r2.1HG0001030.1 | 214.566 | 0.078 | 0.505 | 0.154 | 0.878 | 1.000 |
| HORVU.MOREX.r2.1HG0005290.1 | 0.229 | 0.545 | 3.828 | 0.142 | 0.887 | 1.000 |
| HORVU.MOREX.r2.1HG0007000.1 | 0.305 | 0.536 | 3.786 | 0.142 | 0.887 | 1.000 |
| HORVU.MOREX.r2.1HG0003380.1 | 0.053 | 0.505 | 3.829 | 0.132 | 0.895 | 1.000 |
| HORVU.MOREX.r2.1HG0003440.1 | 0.053 | 0.505 | 3.829 | 0.132 | 0.895 | 1.000 |
| HORVU.MOREX.r2.1HG0001050.1 | 0.053 | 0.505 | 3.829 | 0.132 | 0.895 | 1.000 |
| HORVU.MOREX.r2.1HG0002440.1 | 0.053 | 0.505 | 3.829 | 0.132 | 0.895 | 1.000 |
| HORVU.MOREX.r2.1HG0001560.1 | 0.199 | 0.502 | 3.829 | 0.131 | 0.896 | 1.000 |
| HORVU.MOREX.r2.1HG0000450.1 | 49.801 | 0.083 | 0.642 | 0.129 | 0.898 | 1.000 |
| HORVU.MOREX.r2.1HG0002130.1 | 0.154 | 0.487 | 3.828 | 0.127 | 0.899 | 1.000 |
| HORVU.MOREX.r2.1HG0004340.1 | 0.227 | 0.479 | 3.828 | 0.125 | 0.900 | 1.000 |
| HORVU.MOREX.r2.1HG0005300.1 | 382.793 | 0.033 | 0.278 | 0.118 | 0.906 | 1.000 |
| HORVU.MOREX.r2.1HG0005040.1 | 125.067 | 0.057 | 0.490 | 0.117 | 0.907 | 1.000 |
| HORVU.MOREX.r2.1HG0003340.1 | 51.384 | 0.252 | 2.255 | 0.112 | 0.911 | 1.000 |
| HORVU.MOREX.r2.1HG0001590.1 | 21.586 | 0.103 | 0.977 | 0.106 | 0.916 | 1.000 |
| HORVU.MOREX.r2.1HG0004330.1 | 7.285 | 0.138 | 1.328 | 0.104 | 0.918 | 1.000 |
| HORVU.MOREX.r2.1HG0001400.1 | 2947.253 | 0.071 | 0.725 | 0.098 | 0.922 | 1.000 |
| HORVU.MOREX.r2.1HG0003410.1 | 109.350 | 0.050 | 0.519 | 0.096 | 0.924 | 1.000 |
| HORVU.MOREX.r2.1HG0000640.1 | 255.623 | 0.098 | 1.078 | 0.091 | 0.927 | 1.000 |
| HORVU.MOREX.r2.1HG0006590.1 | 0.839 | 0.336 | 3.784 | 0.089 | 0.929 | 1.000 |
| HORVU.MOREX.r2.1HG0004670.1 | 6.357 | 0.107 | 1.206 | 0.089 | 0.929 | 1.000 |
| HORVU.MOREX.r2.1HG0004050.1 | 59.987 | 0.044 | 0.503 | 0.088 | 0.930 | 1.000 |
| HORVU.MOREX.r2.1HG0006240.1 | 1131.280 | 0.037 | 0.444 | 0.083 | 0.933 | 1.000 |
| HORVU.MOREX.r2.1HG0000060.1 | 703.437 | 0.035 | 0.433 | 0.081 | 0.935 | 1.000 |
| HORVU.MOREX.r2.1HG0003940.1 | 0.715 | 0.287 | 3.626 | 0.079 | 0.937 | 1.000 |
| HORVU.MOREX.r2.1HG0003080.1 | 1286.581 | 0.028 | 0.380 | 0.073 | 0.942 | 1.000 |

| | | | | | | |
|---|---|---|---|---|---|---|
| HORVU.MOREX.r2.1HG0004810.1 | 2154.157 | 0.038 | 0.534 | 0.071 | 0.944 | 1.000 |
| HORVU.MOREX.r2.1HG0004490.1 | 2.123 | 0.148 | 2.236 | 0.066 | 0.947 | 1.000 |
| HORVU.MOREX.r2.1HG0002180.1 | 171.612 | 0.022 | 0.340 | 0.064 | 0.949 | 1.000 |
| HORVU.MOREX.r2.1HG0003920.1 | 32.579 | 0.036 | 0.567 | 0.063 | 0.950 | 1.000 |
| HORVU.MOREX.r2.1HG0005670.1 | 0.448 | 0.232 | 3.761 | 0.062 | 0.951 | 1.000 |
| HORVU.MOREX.r2.1HG0000320.1 | 12.717 | 0.047 | 0.757 | 0.062 | 0.951 | 1.000 |
| HORVU.MOREX.r2.1HG0003370.1 | 0.559 | 0.221 | 3.767 | 0.059 | 0.953 | 1.000 |
| HORVU.MOREX.r2.1HG0001790.1 | 0.714 | 0.170 | 3.747 | 0.045 | 0.964 | 1.000 |
| HORVU.MOREX.r2.1HG0006290.1 | 707.053 | 0.014 | 0.318 | 0.044 | 0.965 | 1.000 |
| HORVU.MOREX.r2.1HG0005080.1 | 35.599 | 0.021 | 0.548 | 0.039 | 0.969 | 1.000 |
| HORVU.MOREX.r2.1HG0007010.1 | 15.691 | 0.026 | 0.698 | 0.037 | 0.970 | 1.000 |
| HORVU.MOREX.r2.1HG0001250.1 | 19.282 | 0.029 | 0.806 | 0.036 | 0.971 | 1.000 |
| HORVU.MOREX.r2.1HG0006750.1 | 1.290 | 0.079 | 2.742 | 0.029 | 0.977 | 1.000 |
| HORVU.MOREX.r2.1HG0006840.1 | 0.810 | 0.097 | 3.534 | 0.027 | 0.978 | 1.000 |
| HORVU.MOREX.r2.1HG0000650.1 | 2.755 | 0.052 | 2.106 | 0.025 | 0.980 | 1.000 |
| HORVU.MOREX.r2.1HG0006540.1 | 419.400 | 0.008 | 0.380 | 0.021 | 0.984 | 1.000 |
| HORVU.MOREX.r2.1HG0004760.1 | 1407.106 | 0.006 | 0.422 | 0.015 | 0.988 | 1.000 |
| HORVU.MOREX.r2.1HG0001580.1 | 184.997 | 0.005 | 0.432 | 0.013 | 0.990 | 1.000 |
| HORVU.MOREX.r2.1HG0003050.1 | 191.271 | 0.002 | 0.381 | 0.005 | 0.996 | 1.000 |
| HORVU.MOREX.r2.1HG0002620.1 | 52.093 | 0.001 | 0.486 | 0.003 | 0.998 | 1.000 |
| HORVU.MOREX.r2.1HG0004820.1 | 744.521 | 0.001 | 0.411 | 0.002 | 0.998 | 1.000 |

Table S2

| Downregulated genes at T2 | baseMean | log2FC | lfcSE | stat | pvalue | padj |
|---|---|---|---|---|---|---|
| HORVU.MOREX.r2.3HG0183610.1 | 77.843 | -22.840 | 3.691 | -6.187 | 0.000 | 0.000 |
| HORVU.MOREX.r2.4HG0295170.1 | 21.428 | -7.407 | 1.271 | -5.827 | 0.000 | 0.000 |
| HORVU.MOREX.r2.1HG0064430.1 | 159.091 | -5.170 | 0.938 | -5.512 | 0.000 | 0.001 |
| HORVU.MOREX.r2.7HG0620060.1 | 59.608 | -2.206 | 0.539 | -4.097 | 0.000 | 0.461 |
| HORVU.MOREX.r2.1HG0004990.1 | 8.801 | -2.639 | 1.090 | -2.420 | 0.016 | 1.000 |
| HORVU.MOREX.r2.1HG0004560.1 | 2.016 | -4.831 | 2.201 | -2.195 | 0.028 | 1.000 |
| HORVU.MOREX.r2.1HG0001570.1 | 4.411 | -2.474 | 1.154 | -2.144 | 0.032 | 1.000 |
| HORVU.MOREX.r2.1HG0003200.1 | 2.316 | -3.753 | 1.818 | -2.065 | 0.039 | 1.000 |
| HORVU.MOREX.r2.1HG0005020.1 | 5.009 | -3.055 | 1.481 | -2.063 | 0.039 | 1.000 |
| HORVU.MOREX.r2.1HG0000490.1 | 22.390 | -1.383 | 0.682 | -2.028 | 0.043 | 1.000 |
| HORVU.MOREX.r2.1HG0004210.1 | 144.419 | -1.935 | 1.010 | -1.916 | 0.055 | 1.000 |
| HORVU.MOREX.r2.1HG0004520.1 | 9.186 | -1.924 | 1.085 | -1.773 | 0.076 | 1.000 |
| HORVU.MOREX.r2.1HG0000390.1 | 3.776 | -2.775 | 1.789 | -1.551 | 0.121 | 1.000 |
| HORVU.MOREX.r2.1HG0003140.1 | 12.898 | -5.495 | 3.665 | -1.499 | 0.134 | 1.000 |
| HORVU.MOREX.r2.1HG0002780.1 | 3.755 | -2.137 | 1.490 | -1.434 | 0.152 | 1.000 |
| HORVU.MOREX.r2.1HG0000080.1 | 95.285 | -0.504 | 0.389 | -1.295 | 0.195 | 1.000 |
| HORVU.MOREX.r2.1HG0000540.1 | 1.136 | -3.940 | 3.112 | -1.266 | 0.206 | 1.000 |
| HORVU.MOREX.r2.1HG0005910.1 | 9.320 | -1.169 | 0.988 | -1.182 | 0.237 | 1.000 |
| HORVU.MOREX.r2.1HG0007020.1 | 61.226 | -0.706 | 0.606 | -1.165 | 0.244 | 1.000 |
| HORVU.MOREX.r2.1HG0006680.1 | 6.909 | -1.541 | 1.323 | -1.164 | 0.244 | 1.000 |
| HORVU.MOREX.r2.1HG0002200.1 | 28.484 | -0.678 | 0.606 | -1.118 | 0.264 | 1.000 |
| HORVU.MOREX.r2.1HG0003280.1 | 2.347 | -2.991 | 2.704 | -1.106 | 0.269 | 1.000 |
| HORVU.MOREX.r2.1HG0000100.1 | 1.672 | -2.056 | 1.863 | -1.104 | 0.270 | 1.000 |
| HORVU.MOREX.r2.1HG0001860.1 | 1.039 | -2.536 | 2.335 | -1.086 | 0.277 | 1.000 |
| HORVU.MOREX.r2.1HG0004010.1 | 5.187 | -2.408 | 2.252 | -1.069 | 0.285 | 1.000 |
| HORVU.MOREX.r2.1HG0002210.1 | 5.101 | -1.182 | 1.108 | -1.066 | 0.286 | 1.000 |
| HORVU.MOREX.r2.1HG0004270.1 | 8.417 | -1.538 | 1.459 | -1.054 | 0.292 | 1.000 |
| HORVU.MOREX.r2.1HG0000580.1 | 39.484 | -0.463 | 0.446 | -1.039 | 0.299 | 1.000 |
| HORVU.MOREX.r2.1HG0002230.1 | 1.161 | -3.073 | 3.002 | -1.024 | 0.306 | 1.000 |
| HORVU.MOREX.r2.1HG0000750.1 | 3.108 | -1.557 | 1.531 | -1.017 | 0.309 | 1.000 |
| HORVU.MOREX.r2.1HG0000670.1 | 93.808 | -0.408 | 0.421 | -0.969 | 0.333 | 1.000 |
| HORVU.MOREX.r2.1HG0005900.1 | 1.127 | -3.095 | 3.213 | -0.963 | 0.335 | 1.000 |
| HORVU.MOREX.r2.1HG0007140.1 | 1.513 | -2.314 | 2.408 | -0.961 | 0.337 | 1.000 |
| HORVU.MOREX.r2.1HG0007420.1 | 0.656 | -3.548 | 3.700 | -0.959 | 0.338 | 1.000 |
| HORVU.MOREX.r2.1HG0004190.1 | 0.653 | -3.543 | 3.700 | -0.958 | 0.338 | 1.000 |
| HORVU.MOREX.r2.1HG0003040.1 | 5.892 | -0.976 | 1.029 | -0.948 | 0.343 | 1.000 |
| HORVU.MOREX.r2.1HG0002680.1 | 3.250 | -1.727 | 1.880 | -0.918 | 0.358 | 1.000 |
| HORVU.MOREX.r2.1HG0006510.1 | 97.046 | -2.109 | 2.321 | -0.909 | 0.363 | 1.000 |
| HORVU.MOREX.r2.1HG0003760.1 | 83.085 | -0.353 | 0.394 | -0.896 | 0.370 | 1.000 |
| HORVU.MOREX.r2.1HG0000840.1 | 8.791 | -1.010 | 1.127 | -0.896 | 0.370 | 1.000 |
| HORVU.MOREX.r2.1HG0004700.1 | 12.288 | -0.625 | 0.707 | -0.883 | 0.377 | 1.000 |
| HORVU.MOREX.r2.1HG0007660.1 | 1.009 | -3.261 | 3.714 | -0.878 | 0.380 | 1.000 |
| HORVU.MOREX.r2.1HG0001710.1 | 482.000 | -0.398 | 0.455 | -0.875 | 0.381 | 1.000 |
| HORVU.MOREX.r2.1HG0004770.1 | 0.488 | -3.180 | 3.715 | -0.856 | 0.392 | 1.000 |
| HORVU.MOREX.r2.1HG0003730.1 | 0.761 | -3.113 | 3.719 | -0.837 | 0.403 | 1.000 |
| HORVU.MOREX.r2.1HG0004200.1 | 73.527 | -0.522 | 0.625 | -0.834 | 0.404 | 1.000 |
| HORVU.MOREX.r2.1HG0001410.1 | 469.032 | -0.555 | 0.670 | -0.828 | 0.407 | 1.000 |
| HORVU.MOREX.r2.1HG0000660.1 | 0.824 | -2.682 | 3.301 | -0.813 | 0.416 | 1.000 |
| HORVU.MOREX.r2.1HG0000470.1 | 867.445 | -0.260 | 0.323 | -0.804 | 0.422 | 1.000 |
| HORVU.MOREX.r2.1HG0001350.1 | 0.536 | -2.990 | 3.727 | -0.802 | 0.422 | 1.000 |
| HORVU.MOREX.r2.1HG0003950.1 | 0.428 | -2.982 | 3.724 | -0.801 | 0.423 | 1.000 |
| HORVU.MOREX.r2.1HG0003750.1 | 83.253 | -0.377 | 0.472 | -0.798 | 0.425 | 1.000 |

| | | | | | | |
|---|---|---|---|---|---|---|
| HORVU.MOREX.r2.1HG0001300.1 | 0.613 | -2.946 | 3.727 | -0.790 | 0.429 | 1.000 |
| HORVU.MOREX.r2.1HG0002580.1 | 73.373 | -2.836 | 3.590 | -0.790 | 0.430 | 1.000 |
| HORVU.MOREX.r2.1HG0007130.1 | 0.907 | -1.944 | 2.480 | -0.784 | 0.433 | 1.000 |
| HORVU.MOREX.r2.1HG0002860.1 | 173.806 | -0.347 | 0.445 | -0.780 | 0.435 | 1.000 |
| HORVU.MOREX.r2.1HG0000340.1 | 28.564 | -0.491 | 0.656 | -0.749 | 0.454 | 1.000 |
| HORVU.MOREX.r2.1HG0006010.1 | 99.820 | -0.830 | 1.122 | -0.740 | 0.460 | 1.000 |
| HORVU.MOREX.r2.1HG0003290.1 | 1.663 | -1.602 | 2.187 | -0.733 | 0.464 | 1.000 |
| HORVU.MOREX.r2.1HG0008040.1 | 0.461 | -2.729 | 3.739 | -0.730 | 0.465 | 1.000 |
| HORVU.MOREX.r2.1HG0003580.1 | 0.463 | -2.722 | 3.739 | -0.728 | 0.467 | 1.000 |
| HORVU.MOREX.r2.1HG0002610.1 | 6.414 | -1.244 | 1.715 | -0.726 | 0.468 | 1.000 |
| HORVU.MOREX.r2.1HG0005640.1 | 20.376 | -0.531 | 0.738 | -0.719 | 0.472 | 1.000 |
| HORVU.MOREX.r2.1HG0001290.1 | 0.363 | -2.676 | 3.740 | -0.716 | 0.474 | 1.000 |
| HORVU.MOREX.r2.1HG0001640.1 | 48.197 | -0.856 | 1.197 | -0.715 | 0.474 | 1.000 |
| HORVU.MOREX.r2.1HG0007740.1 | 56.025 | -0.420 | 0.588 | -0.713 | 0.476 | 1.000 |
| HORVU.MOREX.r2.1HG0004040.1 | 6.509 | -0.769 | 1.108 | -0.694 | 0.487 | 1.000 |
| HORVU.MOREX.r2.1HG0004620.1 | 453.087 | -0.322 | 0.471 | -0.685 | 0.493 | 1.000 |
| HORVU.MOREX.r2.1HG0000980.1 | 3.059 | -1.763 | 2.578 | -0.684 | 0.494 | 1.000 |
| HORVU.MOREX.r2.1HG0000930.1 | 4.229 | -1.580 | 2.345 | -0.674 | 0.500 | 1.000 |
| HORVU.MOREX.r2.1HG0005860.1 | 55.904 | -0.320 | 0.478 | -0.669 | 0.503 | 1.000 |
| HORVU.MOREX.r2.1HG0002820.1 | 0.396 | -2.509 | 3.754 | -0.668 | 0.504 | 1.000 |
| HORVU.MOREX.r2.1HG0004590.1 | 0.535 | -2.460 | 3.756 | -0.655 | 0.513 | 1.000 |
| HORVU.MOREX.r2.1HG0006210.1 | 0.334 | -2.437 | 3.756 | -0.649 | 0.516 | 1.000 |
| HORVU.MOREX.r2.1HG0000720.1 | 233.751 | -0.327 | 0.504 | -0.648 | 0.517 | 1.000 |
| HORVU.MOREX.r2.1HG0003310.1 | 1.062 | -1.832 | 2.837 | -0.646 | 0.518 | 1.000 |
| HORVU.MOREX.r2.1HG0002490.1 | 2.524 | -1.204 | 1.870 | -0.644 | 0.520 | 1.000 |
| HORVU.MOREX.r2.1HG0002240.1 | 0.911 | -2.112 | 3.295 | -0.641 | 0.522 | 1.000 |
| HORVU.MOREX.r2.1HG0006040.1 | 0.315 | -2.386 | 3.762 | -0.634 | 0.526 | 1.000 |
| HORVU.MOREX.r2.1HG0001550.1 | 88.721 | -0.274 | 0.435 | -0.631 | 0.528 | 1.000 |
| HORVU.MOREX.r2.1HG0005370.1 | 0.286 | -2.360 | 3.760 | -0.628 | 0.530 | 1.000 |
| HORVU.MOREX.r2.1HG0007890.1 | 94.466 | -0.570 | 0.910 | -0.626 | 0.531 | 1.000 |
| HORVU.MOREX.r2.1HG0003190.1 | 0.768 | -2.143 | 3.448 | -0.621 | 0.534 | 1.000 |
| HORVU.MOREX.r2.1HG0007300.1 | 0.586 | -2.318 | 3.758 | -0.617 | 0.537 | 1.000 |
| HORVU.MOREX.r2.1HG0001170.1 | 0.924 | -2.264 | 3.721 | -0.608 | 0.543 | 1.000 |
| HORVU.MOREX.r2.1HG0001980.1 | 102.668 | -0.208 | 0.352 | -0.591 | 0.555 | 1.000 |
| HORVU.MOREX.r2.1HG0005500.1 | 1.096 | -1.557 | 2.668 | -0.584 | 0.559 | 1.000 |
| HORVU.MOREX.r2.1HG0006430.1 | 0.772 | -1.754 | 3.008 | -0.583 | 0.560 | 1.000 |
| HORVU.MOREX.r2.1HG0007070.1 | 59.404 | -0.268 | 0.460 | -0.582 | 0.561 | 1.000 |
| HORVU.MOREX.r2.1HG0004130.1 | 3.222 | -1.082 | 1.862 | -0.581 | 0.561 | 1.000 |
| HORVU.MOREX.r2.1HG0000110.1 | 52.996 | -0.270 | 0.477 | -0.566 | 0.571 | 1.000 |
| HORVU.MOREX.r2.1HG0000950.1 | 0.229 | -2.125 | 3.774 | -0.563 | 0.573 | 1.000 |
| HORVU.MOREX.r2.1HG0001010.1 | 0.229 | -2.125 | 3.774 | -0.563 | 0.573 | 1.000 |
| HORVU.MOREX.r2.1HG0005940.1 | 0.229 | -2.125 | 3.774 | -0.563 | 0.573 | 1.000 |
| HORVU.MOREX.r2.1HG0004800.1 | 3.636 | -0.729 | 1.295 | -0.563 | 0.573 | 1.000 |
| HORVU.MOREX.r2.1HG0004360.1 | 248.134 | -0.209 | 0.371 | -0.562 | 0.574 | 1.000 |
| HORVU.MOREX.r2.1HG0005930.1 | 0.313 | -2.111 | 3.771 | -0.560 | 0.576 | 1.000 |
| HORVU.MOREX.r2.1HG0004220.1 | 2.362 | -1.208 | 2.183 | -0.554 | 0.580 | 1.000 |
| HORVU.MOREX.r2.1HG0008010.1 | 23.845 | -0.637 | 1.151 | -0.553 | 0.580 | 1.000 |
| HORVU.MOREX.r2.1HG0000440.1 | 127.345 | -0.239 | 0.452 | -0.528 | 0.597 | 1.000 |
| HORVU.MOREX.r2.1HG0004080.1 | 136.041 | -0.427 | 0.813 | -0.525 | 0.599 | 1.000 |
| HORVU.MOREX.r2.1HG0001910.1 | 146.448 | -0.213 | 0.407 | -0.523 | 0.601 | 1.000 |
| HORVU.MOREX.r2.1HG0004550.1 | 36.355 | -0.630 | 1.229 | -0.512 | 0.609 | 1.000 |
| HORVU.MOREX.r2.1HG0001960.1 | 0.832 | -1.452 | 2.889 | -0.503 | 0.615 | 1.000 |
| HORVU.MOREX.r2.1HG0004630.1 | 186.096 | -0.243 | 0.490 | -0.496 | 0.620 | 1.000 |

| | | | | | | |
|---|---|---|---|---|---|---|
| HORVU.MOREX.r2.1HG0003980.1 | 0.172 | -1.855 | 3.788 | -0.490 | 0.624 | 1.000 |
| HORVU.MOREX.r2.1HG0004120.1 | 0.172 | -1.855 | 3.788 | -0.490 | 0.624 | 1.000 |
| HORVU.MOREX.r2.1HG0007480.1 | 1.182 | -1.174 | 2.402 | -0.489 | 0.625 | 1.000 |
| HORVU.MOREX.r2.1HG0004690.1 | 0.944 | -1.351 | 2.828 | -0.478 | 0.633 | 1.000 |
| HORVU.MOREX.r2.1HG0003180.1 | 1.798 | -1.195 | 2.503 | -0.477 | 0.633 | 1.000 |
| HORVU.MOREX.r2.1HG0007760.1 | 786.289 | -0.157 | 0.329 | -0.476 | 0.634 | 1.000 |
| HORVU.MOREX.r2.1HG0002850.1 | 1.691 | -0.769 | 1.621 | -0.475 | 0.635 | 1.000 |
| HORVU.MOREX.r2.1HG0004160.1 | 0.211 | -1.761 | 3.786 | -0.465 | 0.642 | 1.000 |
| HORVU.MOREX.r2.1HG0004510.1 | 0.993 | -1.688 | 3.717 | -0.454 | 0.650 | 1.000 |
| HORVU.MOREX.r2.1HG0000290.1 | 49.787 | -0.285 | 0.633 | -0.450 | 0.653 | 1.000 |
| HORVU.MOREX.r2.1HG0001820.1 | 0.446 | -1.674 | 3.751 | -0.446 | 0.655 | 1.000 |
| HORVU.MOREX.r2.1HG0004900.1 | 47.097 | -0.260 | 0.591 | -0.440 | 0.660 | 1.000 |
| HORVU.MOREX.r2.1HG0003020.1 | 2.089 | -0.826 | 1.879 | -0.439 | 0.660 | 1.000 |
| HORVU.MOREX.r2.1HG0001530.1 | 333.606 | -0.127 | 0.290 | -0.439 | 0.661 | 1.000 |
| HORVU.MOREX.r2.1HG0003620.1 | 72.039 | -0.317 | 0.729 | -0.435 | 0.664 | 1.000 |
| HORVU.MOREX.r2.1HG0000690.1 | 0.848 | -1.458 | 3.392 | -0.430 | 0.667 | 1.000 |
| HORVU.MOREX.r2.1HG0005520.1 | 8.507 | -0.396 | 0.947 | -0.418 | 0.676 | 1.000 |
| HORVU.MOREX.r2.1HG0002090.1 | 22.172 | -0.611 | 1.476 | -0.414 | 0.679 | 1.000 |
| HORVU.MOREX.r2.1HG0000870.1 | 250.196 | -0.269 | 0.651 | -0.413 | 0.680 | 1.000 |
| HORVU.MOREX.r2.1HG0005350.1 | 13.562 | -0.375 | 0.919 | -0.409 | 0.683 | 1.000 |
| HORVU.MOREX.r2.1HG0001660.1 | 0.114 | -1.538 | 3.796 | -0.405 | 0.685 | 1.000 |
| HORVU.MOREX.r2.1HG0003510.1 | 0.114 | -1.538 | 3.796 | -0.405 | 0.685 | 1.000 |
| HORVU.MOREX.r2.1HG0005970.1 | 0.114 | -1.538 | 3.796 | -0.405 | 0.685 | 1.000 |
| HORVU.MOREX.r2.1HG0006920.1 | 0.114 | -1.538 | 3.796 | -0.405 | 0.685 | 1.000 |
| HORVU.MOREX.r2.1HG0007640.1 | 0.114 | -1.538 | 3.796 | -0.405 | 0.685 | 1.000 |
| HORVU.MOREX.r2.1HG0005790.1 | 0.468 | -1.409 | 3.498 | -0.403 | 0.687 | 1.000 |
| HORVU.MOREX.r2.1HG0003070.1 | 145.870 | -0.175 | 0.443 | -0.396 | 0.692 | 1.000 |
| HORVU.MOREX.r2.1HG0000500.1 | 1.030 | -1.469 | 3.740 | -0.393 | 0.694 | 1.000 |
| HORVU.MOREX.r2.1HG0001150.1 | 1.030 | -1.469 | 3.740 | -0.393 | 0.694 | 1.000 |
| HORVU.MOREX.r2.1HG0000820.1 | 1.030 | -1.469 | 3.740 | -0.393 | 0.694 | 1.000 |
| HORVU.MOREX.r2.1HG0002510.1 | 0.495 | -1.457 | 3.741 | -0.389 | 0.697 | 1.000 |
| HORVU.MOREX.r2.1HG0004000.1 | 0.736 | -1.433 | 3.749 | -0.382 | 0.702 | 1.000 |
| HORVU.MOREX.r2.1HG0002100.1 | 0.167 | -1.445 | 3.799 | -0.380 | 0.704 | 1.000 |
| HORVU.MOREX.r2.1HG0002270.1 | 0.167 | -1.445 | 3.799 | -0.380 | 0.704 | 1.000 |
| HORVU.MOREX.r2.1HG0004580.1 | 0.167 | -1.445 | 3.799 | -0.380 | 0.704 | 1.000 |
| HORVU.MOREX.r2.1HG0004880.1 | 0.167 | -1.445 | 3.799 | -0.380 | 0.704 | 1.000 |
| HORVU.MOREX.r2.1HG0006850.1 | 0.167 | -1.445 | 3.799 | -0.380 | 0.704 | 1.000 |
| HORVU.MOREX.r2.1HG0003530.1 | 16.438 | -0.273 | 0.720 | -0.380 | 0.704 | 1.000 |
| HORVU.MOREX.r2.1HG0001850.1 | 1.062 | -1.071 | 2.821 | -0.380 | 0.704 | 1.000 |
| HORVU.MOREX.r2.1HG0001450.1 | 0.633 | -1.394 | 3.726 | -0.374 | 0.708 | 1.000 |
| HORVU.MOREX.r2.1HG0002430.1 | 0.589 | -1.403 | 3.758 | -0.373 | 0.709 | 1.000 |
| HORVU.MOREX.r2.1HG0003060.1 | 0.589 | -1.403 | 3.758 | -0.373 | 0.709 | 1.000 |
| HORVU.MOREX.r2.1HG0007380.1 | 0.589 | -1.403 | 3.758 | -0.373 | 0.709 | 1.000 |
| HORVU.MOREX.r2.1HG0007800.1 | 0.589 | -1.403 | 3.758 | -0.373 | 0.709 | 1.000 |
| HORVU.MOREX.r2.1HG0005960.1 | 13.974 | -0.250 | 0.675 | -0.370 | 0.711 | 1.000 |
| HORVU.MOREX.r2.1HG0000760.1 | 4.403 | -0.537 | 1.451 | -0.370 | 0.711 | 1.000 |
| HORVU.MOREX.r2.1HG0006440.1 | 0.699 | -1.388 | 3.765 | -0.369 | 0.712 | 1.000 |
| HORVU.MOREX.r2.1HG0006350.1 | 2.155 | -0.510 | 1.394 | -0.366 | 0.715 | 1.000 |
| HORVU.MOREX.r2.1HG0002150.1 | 0.399 | -1.375 | 3.770 | -0.365 | 0.715 | 1.000 |
| HORVU.MOREX.r2.1HG0007290.1 | 0.590 | -1.359 | 3.770 | -0.361 | 0.718 | 1.000 |
| HORVU.MOREX.r2.1HG0001180.1 | 0.441 | -1.358 | 3.771 | -0.360 | 0.719 | 1.000 |
| HORVU.MOREX.r2.1HG0001920.1 | 84.646 | -0.175 | 0.486 | -0.359 | 0.720 | 1.000 |
| HORVU.MOREX.r2.1HG0006660.1 | 35.585 | -0.188 | 0.542 | -0.347 | 0.729 | 1.000 |

| | | | | | | |
|---|---|---|---|---|---|---|
| HORVU.MOREX.r2.1HG0003740.1 | 174.518 | -0.143 | 0.416 | -0.344 | 0.731 | 1.000 |
| HORVU.MOREX.r2.1HG0004600.1 | 0.410 | -1.296 | 3.791 | -0.342 | 0.733 | 1.000 |
| HORVU.MOREX.r2.1HG0002880.1 | 0.453 | -1.295 | 3.791 | -0.342 | 0.733 | 1.000 |
| HORVU.MOREX.r2.1HG0003600.1 | 220.153 | -0.158 | 0.462 | -0.342 | 0.733 | 1.000 |
| HORVU.MOREX.r2.1HG0007330.1 | 52.753 | -0.144 | 0.425 | -0.339 | 0.734 | 1.000 |
| HORVU.MOREX.r2.1HG0001260.1 | 0.294 | -1.280 | 3.794 | -0.337 | 0.736 | 1.000 |
| HORVU.MOREX.r2.1HG0002760.1 | 0.294 | -1.280 | 3.794 | -0.337 | 0.736 | 1.000 |
| HORVU.MOREX.r2.1HG0002810.1 | 0.294 | -1.280 | 3.794 | -0.337 | 0.736 | 1.000 |
| HORVU.MOREX.r2.1HG0003840.1 | 0.294 | -1.280 | 3.794 | -0.337 | 0.736 | 1.000 |
| HORVU.MOREX.r2.1HG0004180.1 | 0.294 | -1.280 | 3.794 | -0.337 | 0.736 | 1.000 |
| HORVU.MOREX.r2.1HG0004300.1 | 0.294 | -1.280 | 3.794 | -0.337 | 0.736 | 1.000 |
| HORVU.MOREX.r2.1HG0005180.1 | 0.294 | -1.280 | 3.794 | -0.337 | 0.736 | 1.000 |
| HORVU.MOREX.r2.1HG0005510.1 | 0.294 | -1.280 | 3.794 | -0.337 | 0.736 | 1.000 |
| HORVU.MOREX.r2.1HG0005700.1 | 0.294 | -1.280 | 3.794 | -0.337 | 0.736 | 1.000 |
| HORVU.MOREX.r2.1HG0007580.1 | 0.294 | -1.280 | 3.794 | -0.337 | 0.736 | 1.000 |
| HORVU.MOREX.r2.1HG0004830.1 | 0.773 | -1.178 | 3.532 | -0.334 | 0.739 | 1.000 |
| HORVU.MOREX.r2.1HG0003570.1 | 0.485 | -1.259 | 3.798 | -0.331 | 0.740 | 1.000 |
| HORVU.MOREX.r2.1HG0006060.1 | 5.520 | -0.521 | 1.581 | -0.329 | 0.742 | 1.000 |
| HORVU.MOREX.r2.1HG0003480.1 | 24.872 | -0.336 | 1.035 | -0.324 | 0.746 | 1.000 |
| HORVU.MOREX.r2.1HG0004390.1 | 65.272 | -0.267 | 0.832 | -0.321 | 0.748 | 1.000 |
| HORVU.MOREX.r2.1HG0005770.1 | 58.473 | -0.205 | 0.638 | -0.321 | 0.749 | 1.000 |
| HORVU.MOREX.r2.1HG0003860.1 | 0.537 | -1.212 | 3.789 | -0.320 | 0.749 | 1.000 |
| HORVU.MOREX.r2.1HG0001760.1 | 40.936 | -0.946 | 3.186 | -0.297 | 0.766 | 1.000 |
| HORVU.MOREX.r2.1HG0006050.1 | 1.041 | -0.739 | 2.525 | -0.293 | 0.770 | 1.000 |
| HORVU.MOREX.r2.1HG0004540.1 | 0.342 | -1.071 | 3.786 | -0.283 | 0.777 | 1.000 |
| HORVU.MOREX.r2.1HG0002310.1 | 0.057 | -1.049 | 3.813 | -0.275 | 0.783 | 1.000 |
| HORVU.MOREX.r2.1HG0005540.1 | 0.057 | -1.049 | 3.813 | -0.275 | 0.783 | 1.000 |
| HORVU.MOREX.r2.1HG0006760.1 | 0.057 | -1.049 | 3.813 | -0.275 | 0.783 | 1.000 |
| HORVU.MOREX.r2.1HG0001090.1 | 0.057 | -1.049 | 3.813 | -0.275 | 0.783 | 1.000 |
| HORVU.MOREX.r2.1HG0001840.1 | 0.057 | -1.049 | 3.813 | -0.275 | 0.783 | 1.000 |
| HORVU.MOREX.r2.1HG0002340.1 | 0.057 | -1.049 | 3.813 | -0.275 | 0.783 | 1.000 |
| HORVU.MOREX.r2.1HG0007340.1 | 0.918 | -0.776 | 2.856 | -0.272 | 0.786 | 1.000 |
| HORVU.MOREX.r2.1HG0006390.1 | 21.002 | -0.189 | 0.713 | -0.265 | 0.791 | 1.000 |
| HORVU.MOREX.r2.1HG0002530.1 | 0.342 | -0.986 | 3.828 | -0.258 | 0.797 | 1.000 |
| HORVU.MOREX.r2.1HG0003260.1 | 7.326 | -0.275 | 1.081 | -0.254 | 0.799 | 1.000 |
| HORVU.MOREX.r2.1HG0000770.1 | 0.567 | -0.951 | 3.786 | -0.251 | 0.802 | 1.000 |
| HORVU.MOREX.r2.1HG0001730.1 | 45.018 | -0.106 | 0.422 | -0.251 | 0.802 | 1.000 |
| HORVU.MOREX.r2.1HG0002170.1 | 1142.816 | -0.181 | 0.722 | -0.251 | 0.802 | 1.000 |
| HORVU.MOREX.r2.1HG0003930.1 | 0.656 | -0.929 | 3.707 | -0.251 | 0.802 | 1.000 |
| HORVU.MOREX.r2.1HG0005830.1 | 0.221 | -0.958 | 3.828 | -0.250 | 0.802 | 1.000 |
| HORVU.MOREX.r2.1HG0000200.1 | 0.147 | -0.950 | 3.828 | -0.248 | 0.804 | 1.000 |
| HORVU.MOREX.r2.1HG0000550.1 | 0.147 | -0.950 | 3.828 | -0.248 | 0.804 | 1.000 |
| HORVU.MOREX.r2.1HG0000570.1 | 0.147 | -0.950 | 3.828 | -0.248 | 0.804 | 1.000 |
| HORVU.MOREX.r2.1HG0000600.1 | 0.147 | -0.950 | 3.828 | -0.248 | 0.804 | 1.000 |
| HORVU.MOREX.r2.1HG0000860.1 | 0.147 | -0.950 | 3.828 | -0.248 | 0.804 | 1.000 |
| HORVU.MOREX.r2.1HG0001780.1 | 0.147 | -0.950 | 3.828 | -0.248 | 0.804 | 1.000 |
| HORVU.MOREX.r2.1HG0001950.1 | 0.147 | -0.950 | 3.828 | -0.248 | 0.804 | 1.000 |
| HORVU.MOREX.r2.1HG0002160.1 | 0.147 | -0.950 | 3.828 | -0.248 | 0.804 | 1.000 |
| HORVU.MOREX.r2.1HG0002380.1 | 0.147 | -0.950 | 3.828 | -0.248 | 0.804 | 1.000 |
| HORVU.MOREX.r2.1HG0002650.1 | 0.147 | -0.950 | 3.828 | -0.248 | 0.804 | 1.000 |
| HORVU.MOREX.r2.1HG0002660.1 | 0.147 | -0.950 | 3.828 | -0.248 | 0.804 | 1.000 |
| HORVU.MOREX.r2.1HG0002910.1 | 0.147 | -0.950 | 3.828 | -0.248 | 0.804 | 1.000 |
| HORVU.MOREX.r2.1HG0002980.1 | 0.147 | -0.950 | 3.828 | -0.248 | 0.804 | 1.000 |

| | | | | | | |
|---|---|---|---|---|---|---|
| HORVU.MOREX.r2.1HG0003490.1 | 0.147 | -0.950 | 3.828 | -0.248 | 0.804 | 1.000 |
| HORVU.MOREX.r2.1HG0005070.1 | 0.147 | -0.950 | 3.828 | -0.248 | 0.804 | 1.000 |
| HORVU.MOREX.r2.1HG0005170.1 | 0.147 | -0.950 | 3.828 | -0.248 | 0.804 | 1.000 |
| HORVU.MOREX.r2.1HG0005410.1 | 0.147 | -0.950 | 3.828 | -0.248 | 0.804 | 1.000 |
| HORVU.MOREX.r2.1HG0005430.1 | 0.147 | -0.950 | 3.828 | -0.248 | 0.804 | 1.000 |
| HORVU.MOREX.r2.1HG0005470.1 | 0.147 | -0.950 | 3.828 | -0.248 | 0.804 | 1.000 |
| HORVU.MOREX.r2.1HG0005590.1 | 0.147 | -0.950 | 3.828 | -0.248 | 0.804 | 1.000 |
| HORVU.MOREX.r2.1HG0005870.1 | 0.147 | -0.950 | 3.828 | -0.248 | 0.804 | 1.000 |
| HORVU.MOREX.r2.1HG0006130.1 | 0.147 | -0.950 | 3.828 | -0.248 | 0.804 | 1.000 |
| HORVU.MOREX.r2.1HG0006160.1 | 0.147 | -0.950 | 3.828 | -0.248 | 0.804 | 1.000 |
| HORVU.MOREX.r2.1HG0006310.1 | 0.147 | -0.950 | 3.828 | -0.248 | 0.804 | 1.000 |
| HORVU.MOREX.r2.1HG0006450.1 | 0.147 | -0.950 | 3.828 | -0.248 | 0.804 | 1.000 |
| HORVU.MOREX.r2.1HG0006480.1 | 0.147 | -0.950 | 3.828 | -0.248 | 0.804 | 1.000 |
| HORVU.MOREX.r2.1HG0006500.1 | 0.147 | -0.950 | 3.828 | -0.248 | 0.804 | 1.000 |
| HORVU.MOREX.r2.1HG0006560.1 | 0.147 | -0.950 | 3.828 | -0.248 | 0.804 | 1.000 |
| HORVU.MOREX.r2.1HG0006780.1 | 0.147 | -0.950 | 3.828 | -0.248 | 0.804 | 1.000 |
| HORVU.MOREX.r2.1HG0006810.1 | 0.147 | -0.950 | 3.828 | -0.248 | 0.804 | 1.000 |
| HORVU.MOREX.r2.1HG0006970.1 | 0.147 | -0.950 | 3.828 | -0.248 | 0.804 | 1.000 |
| HORVU.MOREX.r2.1HG0007060.1 | 0.147 | -0.950 | 3.828 | -0.248 | 0.804 | 1.000 |
| HORVU.MOREX.r2.1HG0007460.1 | 0.147 | -0.950 | 3.828 | -0.248 | 0.804 | 1.000 |
| HORVU.MOREX.r2.1HG0007690.1 | 0.147 | -0.950 | 3.828 | -0.248 | 0.804 | 1.000 |
| HORVU.MOREX.r2.1HG0007700.1 | 0.147 | -0.950 | 3.828 | -0.248 | 0.804 | 1.000 |
| HORVU.MOREX.r2.1HG0007720.1 | 0.147 | -0.950 | 3.828 | -0.248 | 0.804 | 1.000 |
| HORVU.MOREX.r2.1HG0007950.1 | 0.147 | -0.950 | 3.828 | -0.248 | 0.804 | 1.000 |
| HORVU.MOREX.r2.1HG0001360.1 | 0.578 | -0.923 | 3.728 | -0.248 | 0.804 | 1.000 |
| HORVU.MOREX.r2.1HG0002800.1 | 0.243 | -0.905 | 3.827 | -0.236 | 0.813 | 1.000 |
| HORVU.MOREX.r2.1HG0000810.1 | 0.951 | -0.661 | 2.824 | -0.234 | 0.815 | 1.000 |
| HORVU.MOREX.r2.1HG0005990.1 | 0.306 | -0.871 | 3.771 | -0.231 | 0.817 | 1.000 |
| HORVU.MOREX.r2.1HG0008030.1 | 0.960 | -0.861 | 3.744 | -0.230 | 0.818 | 1.000 |
| HORVU.MOREX.r2.1HG0008020.1 | 5.770 | -0.267 | 1.183 | -0.226 | 0.821 | 1.000 |
| HORVU.MOREX.r2.1HG0006860.1 | 3.199 | -0.409 | 1.834 | -0.223 | 0.824 | 1.000 |
| HORVU.MOREX.r2.1HG0000730.1 | 919.448 | -0.058 | 0.277 | -0.209 | 0.834 | 1.000 |
| HORVU.MOREX.r2.1HG0007080.1 | 0.299 | -0.735 | 3.779 | -0.194 | 0.846 | 1.000 |
| HORVU.MOREX.r2.1HG0002250.1 | 0.299 | -0.730 | 3.779 | -0.193 | 0.847 | 1.000 |
| HORVU.MOREX.r2.1HG0007090.1 | 24.780 | -0.143 | 0.809 | -0.177 | 0.859 | 1.000 |
| HORVU.MOREX.r2.1HG0001540.1 | 6788.271 | -0.048 | 0.270 | -0.176 | 0.860 | 1.000 |
| HORVU.MOREX.r2.1HG0001700.1 | 2.938 | -0.268 | 1.542 | -0.174 | 0.862 | 1.000 |
| HORVU.MOREX.r2.1HG0004980.1 | 23.924 | -0.108 | 0.630 | -0.171 | 0.864 | 1.000 |
| HORVU.MOREX.r2.1HG0000960.1 | 127.752 | -0.055 | 0.348 | -0.157 | 0.876 | 1.000 |
| HORVU.MOREX.r2.1HG0006200.1 | 36.565 | -0.076 | 0.486 | -0.156 | 0.876 | 1.000 |
| HORVU.MOREX.r2.1HG0000910.1 | 1.247 | -0.496 | 3.197 | -0.155 | 0.877 | 1.000 |
| HORVU.MOREX.r2.1HG0006720.1 | 295.346 | -0.053 | 0.355 | -0.149 | 0.882 | 1.000 |
| HORVU.MOREX.r2.1HG0004380.1 | 0.621 | -0.558 | 3.753 | -0.149 | 0.882 | 1.000 |
| HORVU.MOREX.r2.1HG0000360.1 | 0.250 | -0.548 | 3.778 | -0.145 | 0.885 | 1.000 |
| HORVU.MOREX.r2.1HG0001190.1 | 0.665 | -0.540 | 3.744 | -0.144 | 0.885 | 1.000 |
| HORVU.MOREX.r2.1HG0006250.1 | 890.338 | -0.065 | 0.460 | -0.141 | 0.888 | 1.000 |
| HORVU.MOREX.r2.1HG0007990.1 | 244.464 | -0.110 | 0.788 | -0.140 | 0.889 | 1.000 |
| HORVU.MOREX.r2.1HG0001970.1 | 0.717 | -0.512 | 3.744 | -0.137 | 0.891 | 1.000 |
| HORVU.MOREX.r2.1HG0004070.1 | 172.362 | -0.049 | 0.362 | -0.135 | 0.893 | 1.000 |
| HORVU.MOREX.r2.1HG0003330.1 | 0.501 | -0.497 | 3.754 | -0.132 | 0.895 | 1.000 |
| HORVU.MOREX.r2.1HG0005610.1 | 0.577 | -0.494 | 3.754 | -0.132 | 0.895 | 1.000 |
| HORVU.MOREX.r2.1HG0001680.1 | 0.575 | -0.490 | 3.754 | -0.130 | 0.896 | 1.000 |
| HORVU.MOREX.r2.1HG0002560.1 | 7881.049 | -0.079 | 0.615 | -0.128 | 0.898 | 1.000 |

| | | | | | | |
|---|---|---|---|---|---|---|
| HORVU.MOREX.r2.1HG0005560.1 | 0.489 | -0.479 | 3.754 | -0.128 | 0.898 | 1.000 |
| HORVU.MOREX.r2.1HG0007110.1 | 1535.584 | -0.165 | 1.326 | -0.125 | 0.901 | 1.000 |
| HORVU.MOREX.r2.1HG0004840.1 | 98.596 | -0.046 | 0.372 | -0.123 | 0.902 | 1.000 |
| HORVU.MOREX.r2.1HG0003010.1 | 93.048 | -0.045 | 0.374 | -0.120 | 0.904 | 1.000 |
| HORVU.MOREX.r2.1HG0000590.1 | 1456.678 | -0.088 | 0.733 | -0.120 | 0.904 | 1.000 |
| HORVU.MOREX.r2.1HG0004890.1 | 0.355 | -0.452 | 3.772 | -0.120 | 0.905 | 1.000 |
| HORVU.MOREX.r2.1HG0004440.1 | 20.928 | -0.083 | 0.698 | -0.119 | 0.906 | 1.000 |
| HORVU.MOREX.r2.1HG0007550.1 | 0.430 | -0.445 | 3.773 | -0.118 | 0.906 | 1.000 |
| HORVU.MOREX.r2.1HG0003660.1 | 0.645 | -0.444 | 3.771 | -0.118 | 0.906 | 1.000 |
| HORVU.MOREX.r2.1HG0007280.1 | 0.429 | -0.442 | 3.773 | -0.117 | 0.907 | 1.000 |
| HORVU.MOREX.r2.1HG0006230.1 | 0.394 | -0.433 | 3.764 | -0.115 | 0.908 | 1.000 |
| HORVU.MOREX.r2.1HG0007790.1 | 2473.077 | -0.050 | 0.441 | -0.114 | 0.909 | 1.000 |
| HORVU.MOREX.r2.1HG0007810.1 | 0.425 | -0.429 | 3.773 | -0.114 | 0.909 | 1.000 |
| HORVU.MOREX.r2.1HG0003670.1 | 0.807 | -0.416 | 3.771 | -0.110 | 0.912 | 1.000 |
| HORVU.MOREX.r2.1HG0003270.1 | 16.977 | -0.066 | 0.616 | -0.107 | 0.915 | 1.000 |
| HORVU.MOREX.r2.1HG0003170.1 | 0.614 | -0.407 | 3.802 | -0.107 | 0.915 | 1.000 |
| HORVU.MOREX.r2.1HG0003830.1 | 1.788 | -0.217 | 2.058 | -0.106 | 0.916 | 1.000 |
| HORVU.MOREX.r2.1HG0006370.1 | 86.190 | -0.040 | 0.381 | -0.104 | 0.917 | 1.000 |
| HORVU.MOREX.r2.1HG0003470.1 | 0.432 | -0.357 | 3.753 | -0.095 | 0.924 | 1.000 |
| HORVU.MOREX.r2.1HG0002770.1 | 1.419 | -0.275 | 2.910 | -0.095 | 0.925 | 1.000 |
| HORVU.MOREX.r2.1HG0002050.1 | 0.371 | -0.350 | 3.825 | -0.092 | 0.927 | 1.000 |
| HORVU.MOREX.r2.1HG0000520.1 | 0.204 | -0.325 | 3.822 | -0.085 | 0.932 | 1.000 |
| HORVU.MOREX.r2.1HG0000120.1 | 149.037 | -0.027 | 0.326 | -0.083 | 0.934 | 1.000 |
| HORVU.MOREX.r2.1HG0001210.1 | 0.279 | -0.310 | 3.823 | -0.081 | 0.935 | 1.000 |
| HORVU.MOREX.r2.1HG0007860.1 | 0.909 | -0.269 | 3.393 | -0.079 | 0.937 | 1.000 |
| HORVU.MOREX.r2.1HG0002590.1 | 0.279 | -0.292 | 3.823 | -0.076 | 0.939 | 1.000 |
| HORVU.MOREX.r2.1HG0006140.1 | 0.279 | -0.279 | 3.824 | -0.073 | 0.942 | 1.000 |
| HORVU.MOREX.r2.1HG0003870.1 | 343.681 | -0.023 | 0.339 | -0.068 | 0.946 | 1.000 |
| HORVU.MOREX.r2.1HG0003720.1 | 0.820 | -0.238 | 3.678 | -0.065 | 0.948 | 1.000 |
| HORVU.MOREX.r2.1HG0006280.1 | 653.171 | -0.018 | 0.277 | -0.065 | 0.949 | 1.000 |
| HORVU.MOREX.r2.1HG0005090.1 | 10.179 | -0.049 | 0.780 | -0.063 | 0.950 | 1.000 |
| HORVU.MOREX.r2.1HG0002640.1 | 6.549 | -0.060 | 0.964 | -0.062 | 0.951 | 1.000 |
| HORVU.MOREX.r2.1HG0007260.1 | 2692.288 | -0.048 | 0.859 | -0.056 | 0.956 | 1.000 |
| HORVU.MOREX.r2.1HG0001890.1 | 0.172 | -0.139 | 3.833 | -0.036 | 0.971 | 1.000 |
| HORVU.MOREX.r2.1HG0003150.1 | 0.146 | -0.139 | 3.833 | -0.036 | 0.971 | 1.000 |
| HORVU.MOREX.r2.1HG0005310.1 | 0.146 | -0.139 | 3.833 | -0.036 | 0.971 | 1.000 |
| HORVU.MOREX.r2.1HG0007190.1 | 0.146 | -0.139 | 3.833 | -0.036 | 0.971 | 1.000 |
| HORVU.MOREX.r2.1HG0001480.1 | 0.099 | -0.139 | 3.833 | -0.036 | 0.971 | 1.000 |
| HORVU.MOREX.r2.1HG0000850.1 | 0.073 | -0.139 | 3.833 | -0.036 | 0.971 | 1.000 |
| HORVU.MOREX.r2.1HG0002040.1 | 0.073 | -0.139 | 3.833 | -0.036 | 0.971 | 1.000 |
| HORVU.MOREX.r2.1HG0007100.1 | 0.073 | -0.139 | 3.833 | -0.036 | 0.971 | 1.000 |
| HORVU.MOREX.r2.1HG0007750.1 | 0.073 | -0.139 | 3.833 | -0.036 | 0.971 | 1.000 |
| HORVU.MOREX.r2.1HG0000940.1 | 0.099 | -0.139 | 3.833 | -0.036 | 0.971 | 1.000 |
| HORVU.MOREX.r2.1HG0005660.1 | 0.099 | -0.139 | 3.833 | -0.036 | 0.971 | 1.000 |
| HORVU.MOREX.r2.1HG0005950.1 | 0.099 | -0.139 | 3.833 | -0.036 | 0.971 | 1.000 |
| HORVU.MOREX.r2.1HG0007600.1 | 0.099 | -0.139 | 3.833 | -0.036 | 0.971 | 1.000 |
| HORVU.MOREX.r2.1HG0002970.1 | 0.198 | -0.139 | 3.833 | -0.036 | 0.971 | 1.000 |
| HORVU.MOREX.r2.1HG0006730.1 | 0.297 | -0.139 | 3.833 | -0.036 | 0.971 | 1.000 |
| HORVU.MOREX.r2.1HG0005600.1 | 0.396 | -0.139 | 3.833 | -0.036 | 0.971 | 1.000 |
| HORVU.MOREX.r2.1HG0003710.1 | 0.891 | -0.139 | 3.833 | -0.036 | 0.971 | 1.000 |
| HORVU.MOREX.r2.1HG0007040.1 | 1.584 | -0.139 | 3.833 | -0.036 | 0.971 | 1.000 |
| HORVU.MOREX.r2.1HG0000740.1 | 1235.669 | -0.017 | 0.476 | -0.035 | 0.972 | 1.000 |
| HORVU.MOREX.r2.1HG0005010.1 | 0.251 | -0.125 | 3.783 | -0.033 | 0.974 | 1.000 |

| | | | | | | |
|---|---|---|---|---|---|---|
| HORVU.MOREX.r2.1HG0007780.1 | 0.250 | -0.102 | 3.783 | -0.027 | 0.978 | 1.000 |
| HORVU.MOREX.r2.1HG0002010.1 | 53.587 | -0.018 | 0.768 | -0.023 | 0.982 | 1.000 |
| HORVU.MOREX.r2.1HG0002330.1 | 327.667 | -0.001 | 0.304 | -0.005 | 0.996 | 1.000 |
| HORVU.MOREX.r2.1HG0007770.1 | 439.948 | -0.002 | 0.365 | -0.004 | 0.996 | 1.000 |

Table S3

| Upregulated gene at T12 | baseMean | log2FC | lfcSE | stat | pvalue | padj |
|---|---|---|---|---|---|---|
| HORVU.MOREX.r2.UnG0634070.1 | 442.533 | 25.410 | 3.319 | 7.657 | 0.000 | 0.000 |
| HORVU.MOREX.r2.1HG0002570.1 | 22.166 | 20.267 | 3.430 | 5.908 | 0.000 | 0.000 |
| HORVU.MOREX.r2.3HG0220600.1 | 0.655 | 18.513 | 3.383 | 5.472 | 0.000 | 0.000 |
| HORVU.MOREX.r2.2HG0125490.1 | 0.812 | 17.131 | 3.381 | 5.066 | 0.000 | 0.002 |
| HORVU.MOREX.r2.3HG0195270.1 | 0.649 | 17.027 | 3.383 | 5.034 | 0.000 | 0.002 |
| HORVU.MOREX.r2.5HG0378750.1 | 0.730 | 16.920 | 3.382 | 5.003 | 0.000 | 0.002 |
| HORVU.MOREX.r2.6HG0497350.1 | 1.136 | 16.849 | 3.380 | 4.985 | 0.000 | 0.002 |
| HORVU.MOREX.r2.4HG0316780.1 | 0.568 | 16.678 | 3.383 | 4.929 | 0.000 | 0.002 |
| HORVU.MOREX.r2.1HG0034850.1 | 0.325 | 16.374 | 3.388 | 4.832 | 0.000 | 0.002 |
| HORVU.MOREX.r2.4HG0341920.1 | 0.325 | 16.374 | 3.388 | 4.832 | 0.000 | 0.002 |
| HORVU.MOREX.r2.7HG0562560.1 | 0.325 | 16.374 | 3.388 | 4.832 | 0.000 | 0.002 |
| HORVU.MOREX.r2.1HG0059640.1 | 0.325 | 16.374 | 3.388 | 4.832 | 0.000 | 0.002 |
| HORVU.MOREX.r2.6HG0482090.1 | 0.325 | 16.374 | 3.388 | 4.832 | 0.000 | 0.002 |
| HORVU.MOREX.r2.7HG0572970.1 | 0.325 | 16.374 | 3.388 | 4.832 | 0.000 | 0.002 |
| HORVU.MOREX.r2.5HG0394780.1 | 0.325 | 16.374 | 3.388 | 4.832 | 0.000 | 0.002 |
| HORVU.MOREX.r2.7HG0549100.1 | 0.325 | 16.374 | 3.388 | 4.832 | 0.000 | 0.002 |
| HORVU.MOREX.r2.2HG0131850.1 | 0.325 | 16.374 | 3.388 | 4.832 | 0.000 | 0.002 |
| HORVU.MOREX.r2.2HG0161130.1 | 0.325 | 16.374 | 3.388 | 4.832 | 0.000 | 0.002 |
| HORVU.MOREX.r2.3HG0249080.1 | 0.325 | 16.374 | 3.388 | 4.832 | 0.000 | 0.002 |
| HORVU.MOREX.r2.3HG0259900.1 | 0.325 | 16.374 | 3.388 | 4.832 | 0.000 | 0.002 |
| HORVU.MOREX.r2.4HG0306080.1 | 0.325 | 16.374 | 3.388 | 4.832 | 0.000 | 0.002 |
| HORVU.MOREX.r2.4HG0340840.1 | 0.325 | 16.374 | 3.388 | 4.832 | 0.000 | 0.002 |
| HORVU.MOREX.r2.4HG0331710.1 | 0.325 | 16.374 | 3.388 | 4.832 | 0.000 | 0.002 |
| HORVU.MOREX.r2.4HG0336190.1 | 0.325 | 16.374 | 3.388 | 4.832 | 0.000 | 0.002 |
| HORVU.MOREX.r2.3HG0268080.1 | 0.325 | 16.374 | 3.388 | 4.832 | 0.000 | 0.002 |
| HORVU.MOREX.r2.1HG0069050.1 | 0.325 | 16.374 | 3.388 | 4.832 | 0.000 | 0.002 |
| HORVU.MOREX.r2.1HG0009060.1 | 0.406 | 16.277 | 3.386 | 4.807 | 0.000 | 0.002 |
| HORVU.MOREX.r2.2HG0145620.1 | 0.406 | 16.277 | 3.386 | 4.807 | 0.000 | 0.002 |
| HORVU.MOREX.r2.7HG0584550.1 | 0.406 | 16.277 | 3.386 | 4.807 | 0.000 | 0.002 |
| HORVU.MOREX.r2.5HG0356840.1 | 0.406 | 16.277 | 3.386 | 4.807 | 0.000 | 0.002 |
| HORVU.MOREX.r2.3HG0253910.1 | 0.406 | 16.277 | 3.386 | 4.807 | 0.000 | 0.002 |
| HORVU.MOREX.r2.4HG0296500.1 | 0.406 | 16.277 | 3.386 | 4.807 | 0.000 | 0.002 |
| HORVU.MOREX.r2.2HG0127650.1 | 1010.253 | 14.310 | 3.251 | 4.402 | 0.000 | 0.012 |
| HORVU.MOREX.r2.5HG0432480.1 | 322.467 | 1.462 | 0.366 | 4.000 | 0.000 | 0.067 |
| HORVU.MOREX.r2.4HG0348610.1 | 69.734 | 2.770 | 0.710 | 3.902 | 0.000 | 0.099 |
| HORVU.MOREX.r2.6HG0513160.1 | 86.310 | 2.453 | 0.633 | 3.872 | 0.000 | 0.109 |
| HORVU.MOREX.r2.3HG0274450.1 | 12.911 | 4.256 | 1.166 | 3.652 | 0.000 | 0.257 |
| HORVU.MOREX.r2.1HG0012300.1 | 18.775 | 4.699 | 1.290 | 3.644 | 0.000 | 0.260 |
| HORVU.MOREX.r2.2HG0115620.1 | 374.874 | 1.198 | 0.343 | 3.492 | 0.000 | 0.443 |
| HORVU.MOREX.r2.3HG0237050.1 | 5.869 | 5.604 | 1.645 | 3.406 | 0.001 | 0.576 |
| HORVU.MOREX.r2.UnG0628800.1 | 3416.147 | 1.775 | 0.539 | 3.293 | 0.001 | 0.849 |
| HORVU.MOREX.r2.1HG0003890.1 | 14.538 | 5.481 | 3.130 | 1.751 | 0.080 | 0.999 |
| HORVU.MOREX.r2.1HG0005910.1 | 8.857 | 4.792 | 3.234 | 1.482 | 0.138 | 0.999 |
| HORVU.MOREX.r2.1HG0004990.1 | 7.065 | 4.015 | 3.196 | 1.256 | 0.209 | 0.999 |
| HORVU.MOREX.r2.1HG0004510.1 | 3.097 | 3.319 | 2.290 | 1.449 | 0.147 | 0.999 |
| HORVU.MOREX.r2.1HG0003290.1 | 3.479 | 3.283 | 1.800 | 1.823 | 0.068 | 0.999 |
| HORVU.MOREX.r2.1HG0006750.1 | 3.061 | 3.193 | 1.630 | 1.959 | 0.050 | 0.999 |
| HORVU.MOREX.r2.1HG0002070.1 | 1.771 | 3.164 | 2.081 | 1.520 | 0.128 | 0.999 |
| HORVU.MOREX.r2.1HG0002830.1 | 1.262 | 2.943 | 2.277 | 1.292 | 0.196 | 0.999 |
| HORVU.MOREX.r2.1HG0001850.1 | 1.733 | 2.871 | 2.394 | 1.199 | 0.230 | 0.999 |
| HORVU.MOREX.r2.1HG0007140.1 | 2.254 | 2.791 | 1.911 | 1.461 | 0.144 | 0.999 |
| HORVU.MOREX.r2.1HG0003830.1 | 1.475 | 2.758 | 2.275 | 1.213 | 0.225 | 0.999 |

| | | | | | | |
|---|---|---|---|---|---|---|
| HORVU.MOREX.r2.1HG0003860.1 | 2.476 | 2.678 | 1.871 | 1.431 | 0.152 | 0.999 |
| HORVU.MOREX.r2.1HG0005030.1 | 1.809 | 2.667 | 2.459 | 1.084 | 0.278 | 0.999 |
| HORVU.MOREX.r2.1HG0001260.1 | 0.538 | 2.592 | 3.385 | 0.766 | 0.444 | 0.999 |
| HORVU.MOREX.r2.1HG0006520.1 | 1.026 | 2.451 | 3.388 | 0.724 | 0.469 | 0.999 |
| HORVU.MOREX.r2.1HG0005680.1 | 0.461 | 2.405 | 3.395 | 0.708 | 0.479 | 0.999 |
| HORVU.MOREX.r2.1HG0000690.1 | 1.448 | 2.376 | 2.259 | 1.052 | 0.293 | 0.999 |
| HORVU.MOREX.r2.1HG0005160.1 | 1.418 | 2.369 | 2.840 | 0.834 | 0.404 | 0.999 |
| HORVU.MOREX.r2.1HG0002780.1 | 3.922 | 2.325 | 1.800 | 1.292 | 0.196 | 0.999 |
| HORVU.MOREX.r2.1HG0005580.1 | 1.043 | 2.303 | 2.690 | 0.856 | 0.392 | 0.999 |
| HORVU.MOREX.r2.1HG0005530.1 | 1.189 | 2.269 | 2.626 | 0.864 | 0.388 | 0.999 |
| HORVU.MOREX.r2.1HG0004520.1 | 10.976 | 2.035 | 1.291 | 1.576 | 0.115 | 0.999 |
| HORVU.MOREX.r2.1HG0006430.1 | 1.512 | 2.013 | 2.608 | 0.772 | 0.440 | 0.999 |
| HORVU.MOREX.r2.1HG0006190.1 | 4.069 | 1.935 | 1.147 | 1.687 | 0.092 | 0.999 |
| HORVU.MOREX.r2.1HG0003470.1 | 1.248 | 1.872 | 2.980 | 0.628 | 0.530 | 0.999 |
| HORVU.MOREX.r2.1HG0000570.1 | 0.284 | 1.870 | 3.420 | 0.547 | 0.585 | 0.999 |
| HORVU.MOREX.r2.1HG0003580.1 | 0.880 | 1.855 | 3.436 | 0.540 | 0.589 | 0.999 |
| HORVU.MOREX.r2.1HG0002430.1 | 0.628 | 1.855 | 3.429 | 0.541 | 0.589 | 0.999 |
| HORVU.MOREX.r2.1HG0001350.1 | 1.100 | 1.767 | 2.791 | 0.633 | 0.527 | 0.999 |
| HORVU.MOREX.r2.1HG0000900.1 | 1.638 | 1.718 | 2.629 | 0.653 | 0.514 | 0.999 |
| HORVU.MOREX.r2.1HG0005380.1 | 0.481 | 1.680 | 3.379 | 0.497 | 0.619 | 0.999 |
| HORVU.MOREX.r2.1HG0004560.1 | 1.993 | 1.672 | 2.103 | 0.795 | 0.427 | 0.999 |
| HORVU.MOREX.r2.1HG0005930.1 | 0.501 | 1.660 | 3.434 | 0.483 | 0.629 | 0.999 |
| HORVU.MOREX.r2.1HG0003810.1 | 0.884 | 1.649 | 2.860 | 0.576 | 0.564 | 0.999 |
| HORVU.MOREX.r2.1HG0002680.1 | 3.721 | 1.621 | 1.562 | 1.038 | 0.299 | 0.999 |
| HORVU.MOREX.r2.1HG0002100.1 | 0.706 | 1.614 | 3.374 | 0.479 | 0.632 | 0.999 |
| HORVU.MOREX.r2.1HG0002770.1 | 1.550 | 1.586 | 1.890 | 0.840 | 0.401 | 0.999 |
| HORVU.MOREX.r2.1HG0003190.1 | 1.651 | 1.536 | 2.326 | 0.660 | 0.509 | 0.999 |
| HORVU.MOREX.r2.1HG0006320.1 | 0.492 | 1.513 | 3.434 | 0.441 | 0.659 | 0.999 |
| HORVU.MOREX.r2.1HG0003880.1 | 0.162 | 1.496 | 3.428 | 0.436 | 0.662 | 0.999 |
| HORVU.MOREX.r2.1HG0001840.1 | 0.450 | 1.492 | 3.441 | 0.434 | 0.664 | 0.999 |
| HORVU.MOREX.r2.1HG0001860.1 | 1.423 | 1.476 | 2.334 | 0.632 | 0.527 | 0.999 |
| HORVU.MOREX.r2.1HG0002850.1 | 2.156 | 1.362 | 1.765 | 0.772 | 0.440 | 0.999 |
| HORVU.MOREX.r2.1HG0005720.1 | 0.378 | 1.290 | 3.447 | 0.374 | 0.708 | 0.999 |
| HORVU.MOREX.r2.1HG0002980.1 | 0.377 | 1.290 | 3.447 | 0.374 | 0.708 | 0.999 |
| HORVU.MOREX.r2.1HG0005180.1 | 0.769 | 1.278 | 3.420 | 0.374 | 0.709 | 0.999 |
| HORVU.MOREX.r2.1HG0000650.1 | 0.950 | 1.251 | 3.164 | 0.395 | 0.693 | 0.999 |
| HORVU.MOREX.r2.1HG0003970.1 | 0.824 | 1.228 | 3.310 | 0.371 | 0.711 | 0.999 |
| HORVU.MOREX.r2.1HG0000930.1 | 2.768 | 1.140 | 1.779 | 0.641 | 0.522 | 0.999 |
| HORVU.MOREX.r2.1HG0000830.1 | 2.445 | 1.133 | 1.398 | 0.811 | 0.418 | 0.999 |
| HORVU.MOREX.r2.1HG0004490.1 | 3.193 | 1.126 | 1.530 | 0.736 | 0.462 | 0.999 |
| HORVU.MOREX.r2.1HG0000970.1 | 0.435 | 1.117 | 3.398 | 0.329 | 0.742 | 0.999 |
| HORVU.MOREX.r2.1HG0001470.1 | 3.775 | 1.105 | 1.087 | 1.017 | 0.309 | 0.999 |
| HORVU.MOREX.r2.1HG0000390.1 | 4.989 | 1.092 | 1.312 | 0.833 | 0.405 | 0.999 |
| HORVU.MOREX.r2.1HG0003930.1 | 1.070 | 1.067 | 2.243 | 0.476 | 0.634 | 0.999 |
| HORVU.MOREX.r2.1HG0002820.1 | 0.866 | 1.045 | 2.852 | 0.366 | 0.714 | 0.999 |
| HORVU.MOREX.r2.1HG0002870.1 | 4.458 | 1.016 | 1.069 | 0.951 | 0.342 | 0.999 |
| HORVU.MOREX.r2.1HG0000520.1 | 0.456 | 1.008 | 3.441 | 0.293 | 0.770 | 0.999 |
| HORVU.MOREX.r2.1HG0001560.1 | 0.236 | 0.999 | 3.418 | 0.292 | 0.770 | 0.999 |
| HORVU.MOREX.r2.1HG0004160.1 | 0.293 | 0.993 | 3.425 | 0.290 | 0.772 | 0.999 |
| HORVU.MOREX.r2.1HG0001100.1 | 0.457 | 0.963 | 3.444 | 0.279 | 0.780 | 0.999 |
| HORVU.MOREX.r2.1HG0003370.1 | 0.900 | 0.961 | 3.261 | 0.295 | 0.768 | 0.999 |
| HORVU.MOREX.r2.1HG0003180.1 | 1.097 | 0.942 | 2.641 | 0.357 | 0.721 | 0.999 |
| HORVU.MOREX.r2.1HG0000150.1 | 0.250 | 0.936 | 3.459 | 0.271 | 0.787 | 0.999 |

| | | | | | | |
|---|---|---|---|---|---|---|
| HORVU.MOREX.r2.1HG0007060.1 | 0.250 | 0.935 | 3.459 | 0.270 | 0.787 | 0.999 |
| HORVU.MOREX.r2.1HG0003440.1 | 0.250 | 0.933 | 3.459 | 0.270 | 0.787 | 0.999 |
| HORVU.MOREX.r2.1HG0001960.1 | 0.545 | 0.911 | 3.409 | 0.267 | 0.789 | 0.999 |
| HORVU.MOREX.r2.1HG0000810.1 | 0.900 | 0.897 | 2.548 | 0.352 | 0.725 | 0.999 |
| HORVU.MOREX.r2.1HG0006420.1 | 0.806 | 0.889 | 2.930 | 0.304 | 0.761 | 0.999 |
| HORVU.MOREX.r2.1HG0007050.1 | 2.798 | 0.886 | 2.288 | 0.387 | 0.699 | 0.999 |
| HORVU.MOREX.r2.1HG0000320.1 | 12.970 | 0.881 | 0.667 | 1.320 | 0.187 | 0.999 |
| HORVU.MOREX.r2.1HG0003260.1 | 7.459 | 0.863 | 1.110 | 0.777 | 0.437 | 0.999 |
| HORVU.MOREX.r2.1HG0002130.1 | 0.863 | 0.849 | 2.326 | 0.365 | 0.715 | 0.999 |
| HORVU.MOREX.r2.1HG0002890.1 | 0.256 | 0.842 | 3.449 | 0.244 | 0.807 | 0.999 |
| HORVU.MOREX.r2.1HG0006790.1 | 0.256 | 0.842 | 3.449 | 0.244 | 0.807 | 0.999 |
| HORVU.MOREX.r2.1HG0006920.1 | 0.256 | 0.842 | 3.449 | 0.244 | 0.807 | 0.999 |
| HORVU.MOREX.r2.1HG0003660.1 | 0.574 | 0.826 | 3.377 | 0.245 | 0.807 | 0.999 |
| HORVU.MOREX.r2.1HG0002160.1 | 0.917 | 0.824 | 3.276 | 0.252 | 0.801 | 0.999 |
| HORVU.MOREX.r2.1HG0004980.1 | 28.446 | 0.812 | 0.513 | 1.583 | 0.113 | 0.999 |
| HORVU.MOREX.r2.1HG0005360.1 | 191.964 | 0.793 | 0.382 | 2.078 | 0.038 | 0.999 |
| HORVU.MOREX.r2.1HG0000540.1 | 0.367 | 0.793 | 3.417 | 0.232 | 0.817 | 0.999 |
| HORVU.MOREX.r2.1HG0002230.1 | 0.859 | 0.789 | 3.263 | 0.242 | 0.809 | 0.999 |
| HORVU.MOREX.r2.1HG0005790.1 | 0.352 | 0.788 | 3.419 | 0.231 | 0.818 | 0.999 |
| HORVU.MOREX.r2.1HG0001900.1 | 2.237 | 0.755 | 1.511 | 0.500 | 0.617 | 0.999 |
| HORVU.MOREX.r2.1HG0000950.1 | 0.245 | 0.744 | 3.442 | 0.216 | 0.829 | 0.999 |
| HORVU.MOREX.r2.1HG0001600.1 | 0.245 | 0.744 | 3.442 | 0.216 | 0.829 | 0.999 |
| HORVU.MOREX.r2.1HG0002650.1 | 0.245 | 0.744 | 3.442 | 0.216 | 0.829 | 0.999 |
| HORVU.MOREX.r2.1HG0003230.1 | 0.245 | 0.744 | 3.442 | 0.216 | 0.829 | 0.999 |
| HORVU.MOREX.r2.1HG0004770.1 | 0.245 | 0.744 | 3.442 | 0.216 | 0.829 | 0.999 |
| HORVU.MOREX.r2.1HG0006670.1 | 0.245 | 0.744 | 3.442 | 0.216 | 0.829 | 0.999 |
| HORVU.MOREX.r2.1HG0000400.1 | 0.245 | 0.744 | 3.442 | 0.216 | 0.829 | 0.999 |
| HORVU.MOREX.r2.1HG0000500.1 | 0.245 | 0.744 | 3.442 | 0.216 | 0.829 | 0.999 |
| HORVU.MOREX.r2.1HG0001170.1 | 0.245 | 0.744 | 3.442 | 0.216 | 0.829 | 0.999 |
| HORVU.MOREX.r2.1HG0004090.1 | 0.245 | 0.744 | 3.442 | 0.216 | 0.829 | 0.999 |
| HORVU.MOREX.r2.1HG0004850.1 | 0.245 | 0.744 | 3.442 | 0.216 | 0.829 | 0.999 |
| HORVU.MOREX.r2.1HG0002800.1 | 1.189 | 0.728 | 2.683 | 0.271 | 0.786 | 0.999 |
| HORVU.MOREX.r2.1HG0004550.1 | 44.767 | 0.724 | 0.874 | 0.828 | 0.408 | 0.999 |
| HORVU.MOREX.r2.1HG0003530.1 | 18.462 | 0.684 | 0.651 | 1.051 | 0.293 | 0.999 |
| HORVU.MOREX.r2.1HG0006880.1 | 7.567 | 0.662 | 1.071 | 0.618 | 0.536 | 0.999 |
| HORVU.MOREX.r2.1HG0000980.1 | 1.417 | 0.643 | 3.052 | 0.211 | 0.833 | 0.999 |
| HORVU.MOREX.r2.1HG0003330.1 | 0.777 | 0.633 | 2.694 | 0.235 | 0.814 | 0.999 |
| HORVU.MOREX.r2.1HG0005860.1 | 66.302 | 0.630 | 0.394 | 1.598 | 0.110 | 0.999 |
| HORVU.MOREX.r2.1HG0006150.1 | 0.335 | 0.629 | 3.460 | 0.182 | 0.856 | 0.999 |
| HORVU.MOREX.r2.1HG0003200.1 | 1.511 | 0.611 | 1.819 | 0.336 | 0.737 | 0.999 |
| HORVU.MOREX.r2.1HG0003480.1 | 31.434 | 0.609 | 0.690 | 0.883 | 0.377 | 0.999 |
| HORVU.MOREX.r2.1HG0001570.1 | 4.064 | 0.604 | 1.172 | 0.515 | 0.606 | 0.999 |
| HORVU.MOREX.r2.1HG0003680.1 | 0.178 | 0.598 | 3.444 | 0.174 | 0.862 | 0.999 |
| HORVU.MOREX.r2.1HG0004440.1 | 21.832 | 0.571 | 0.692 | 0.826 | 0.409 | 0.999 |
| HORVU.MOREX.r2.1HG0003950.1 | 0.320 | 0.566 | 3.460 | 0.164 | 0.870 | 0.999 |
| HORVU.MOREX.r2.1HG0002150.1 | 0.329 | 0.564 | 3.399 | 0.166 | 0.868 | 0.999 |
| HORVU.MOREX.r2.1HG0006800.1 | 0.164 | 0.562 | 3.443 | 0.163 | 0.870 | 0.999 |
| HORVU.MOREX.r2.1HG0004050.1 | 71.538 | 0.561 | 0.529 | 1.060 | 0.289 | 0.999 |
| HORVU.MOREX.r2.1HG0005260.1 | 0.333 | 0.559 | 3.461 | 0.162 | 0.872 | 0.999 |
| HORVU.MOREX.r2.1HG0004240.1 | 2.068 | 0.554 | 1.993 | 0.278 | 0.781 | 0.999 |
| HORVU.MOREX.r2.1HG0005900.1 | 1.100 | 0.550 | 2.964 | 0.186 | 0.853 | 0.999 |
| HORVU.MOREX.r2.1HG0003270.1 | 23.718 | 0.535 | 0.513 | 1.042 | 0.298 | 0.999 |
| HORVU.MOREX.r2.1HG0007190.1 | 0.299 | 0.513 | 3.461 | 0.148 | 0.882 | 0.999 |

| | | | | | | |
|---|---|---|---|---|---|---|
| HORVU.MOREX.r2.1HG0003380.1 | 0.339 | 0.504 | 3.450 | 0.146 | 0.884 | 0.999 |
| HORVU.MOREX.r2.1HG0007040.1 | 2.511 | 0.494 | 2.803 | 0.176 | 0.860 | 0.999 |
| HORVU.MOREX.r2.1HG0005040.1 | 139.594 | 0.491 | 0.453 | 1.083 | 0.279 | 0.999 |
| HORVU.MOREX.r2.1HG0002790.1 | 13.763 | 0.490 | 0.692 | 0.708 | 0.479 | 0.999 |
| HORVU.MOREX.r2.1HG0006890.1 | 0.714 | 0.473 | 3.375 | 0.140 | 0.889 | 0.999 |
| HORVU.MOREX.r2.1HG0005550.1 | 0.407 | 0.461 | 3.392 | 0.136 | 0.892 | 0.999 |
| HORVU.MOREX.r2.1HG0005440.1 | 39.273 | 0.460 | 0.448 | 1.028 | 0.304 | 0.999 |
| HORVU.MOREX.r2.1HG0006660.1 | 50.914 | 0.456 | 0.599 | 0.761 | 0.447 | 0.999 |
| HORVU.MOREX.r2.1HG0003400.1 | 91.111 | 0.451 | 0.449 | 1.005 | 0.315 | 0.999 |
| HORVU.MOREX.r2.1HG0006370.1 | 87.105 | 0.448 | 0.358 | 1.251 | 0.211 | 0.999 |
| HORVU.MOREX.r2.1HG0000800.1 | 0.128 | 0.444 | 3.461 | 0.128 | 0.898 | 0.999 |
| HORVU.MOREX.r2.1HG0001120.1 | 0.128 | 0.444 | 3.461 | 0.128 | 0.898 | 0.999 |
| HORVU.MOREX.r2.1HG0003790.1 | 0.128 | 0.444 | 3.461 | 0.128 | 0.898 | 0.999 |
| HORVU.MOREX.r2.1HG0004140.1 | 0.128 | 0.444 | 3.461 | 0.128 | 0.898 | 0.999 |
| HORVU.MOREX.r2.1HG0004300.1 | 0.128 | 0.444 | 3.461 | 0.128 | 0.898 | 0.999 |
| HORVU.MOREX.r2.1HG0006230.1 | 0.128 | 0.444 | 3.461 | 0.128 | 0.898 | 0.999 |
| HORVU.MOREX.r2.1HG0000160.1 | 0.128 | 0.444 | 3.461 | 0.128 | 0.898 | 0.999 |
| HORVU.MOREX.r2.1HG0003560.1 | 0.128 | 0.444 | 3.461 | 0.128 | 0.898 | 0.999 |
| HORVU.MOREX.r2.1HG0003990.1 | 0.128 | 0.444 | 3.461 | 0.128 | 0.898 | 0.999 |
| HORVU.MOREX.r2.1HG0005390.1 | 8.473 | 0.429 | 0.760 | 0.564 | 0.573 | 0.999 |
| HORVU.MOREX.r2.1HG0000210.1 | 0.109 | 0.425 | 3.462 | 0.123 | 0.902 | 0.999 |
| HORVU.MOREX.r2.1HG0000620.1 | 0.122 | 0.425 | 3.462 | 0.123 | 0.902 | 0.999 |
| HORVU.MOREX.r2.1HG0001520.1 | 0.122 | 0.425 | 3.462 | 0.123 | 0.902 | 0.999 |
| HORVU.MOREX.r2.1HG0001660.1 | 0.122 | 0.425 | 3.462 | 0.123 | 0.902 | 0.999 |
| HORVU.MOREX.r2.1HG0001820.1 | 0.122 | 0.425 | 3.462 | 0.123 | 0.902 | 0.999 |
| HORVU.MOREX.r2.1HG0001970.1 | 0.122 | 0.425 | 3.462 | 0.123 | 0.902 | 0.999 |
| HORVU.MOREX.r2.1HG0002030.1 | 0.122 | 0.425 | 3.462 | 0.123 | 0.902 | 0.999 |
| HORVU.MOREX.r2.1HG0002380.1 | 0.122 | 0.425 | 3.462 | 0.123 | 0.902 | 0.999 |
| HORVU.MOREX.r2.1HG0003780.1 | 0.122 | 0.425 | 3.462 | 0.123 | 0.902 | 0.999 |
| HORVU.MOREX.r2.1HG0003980.1 | 0.122 | 0.425 | 3.462 | 0.123 | 0.902 | 0.999 |
| HORVU.MOREX.r2.1HG0004100.1 | 0.122 | 0.425 | 3.462 | 0.123 | 0.902 | 0.999 |
| HORVU.MOREX.r2.1HG0004720.1 | 0.122 | 0.425 | 3.462 | 0.123 | 0.902 | 0.999 |
| HORVU.MOREX.r2.1HG0005190.1 | 0.122 | 0.425 | 3.462 | 0.123 | 0.902 | 0.999 |
| HORVU.MOREX.r2.1HG0005240.1 | 0.122 | 0.425 | 3.462 | 0.123 | 0.902 | 0.999 |
| HORVU.MOREX.r2.1HG0005320.1 | 0.122 | 0.425 | 3.462 | 0.123 | 0.902 | 0.999 |
| HORVU.MOREX.r2.1HG0005450.1 | 0.122 | 0.425 | 3.462 | 0.123 | 0.902 | 0.999 |
| HORVU.MOREX.r2.1HG0005480.1 | 0.122 | 0.425 | 3.462 | 0.123 | 0.902 | 0.999 |
| HORVU.MOREX.r2.1HG0005620.1 | 0.122 | 0.425 | 3.462 | 0.123 | 0.902 | 0.999 |
| HORVU.MOREX.r2.1HG0006260.1 | 0.109 | 0.425 | 3.462 | 0.123 | 0.902 | 0.999 |
| HORVU.MOREX.r2.1HG0004230.1 | 130.284 | 0.398 | 0.300 | 1.325 | 0.185 | 0.999 |
| HORVU.MOREX.r2.1HG0006870.1 | 4.439 | 0.392 | 1.308 | 0.300 | 0.764 | 0.999 |
| HORVU.MOREX.r2.1HG0000290.1 | 46.475 | 0.364 | 0.396 | 0.921 | 0.357 | 0.999 |
| HORVU.MOREX.r2.1HG0003500.1 | 1571.829 | 0.356 | 0.386 | 0.920 | 0.357 | 0.999 |
| HORVU.MOREX.r2.1HG0001450.1 | 0.704 | 0.354 | 3.387 | 0.105 | 0.917 | 0.999 |
| HORVU.MOREX.r2.1HG0003740.1 | 199.075 | 0.347 | 0.322 | 1.078 | 0.281 | 0.999 |
| HORVU.MOREX.r2.1HG0006180.1 | 7.274 | 0.341 | 0.858 | 0.398 | 0.691 | 0.999 |
| HORVU.MOREX.r2.1HG0004410.1 | 45.103 | 0.341 | 0.437 | 0.781 | 0.435 | 0.999 |
| HORVU.MOREX.r2.1HG0002530.1 | 0.570 | 0.334 | 3.394 | 0.098 | 0.922 | 0.999 |
| HORVU.MOREX.r2.1HG0002760.1 | 0.459 | 0.325 | 3.405 | 0.095 | 0.924 | 0.999 |
| HORVU.MOREX.r2.1HG0006060.1 | 3.026 | 0.318 | 1.267 | 0.251 | 0.802 | 0.999 |
| HORVU.MOREX.r2.1HG0000080.1 | 140.805 | 0.317 | 0.509 | 0.623 | 0.533 | 0.999 |
| HORVU.MOREX.r2.1HG0004260.1 | 45.341 | 0.315 | 0.554 | 0.569 | 0.569 | 0.999 |
| HORVU.MOREX.r2.1HG0001010.1 | 0.448 | 0.312 | 3.406 | 0.092 | 0.927 | 0.999 |

| | | | | | | |
|---|---|---|---|---|---|---|
| HORVU.MOREX.r2.1HG0004220.1 | 2.138 | 0.309 | 1.668 | 0.185 | 0.853 | 0.999 |
| HORVU.MOREX.r2.1HG0003630.1 | 84.850 | 0.295 | 0.443 | 0.665 | 0.506 | 0.999 |
| HORVU.MOREX.r2.1HG0003350.1 | 0.318 | 0.287 | 3.430 | 0.084 | 0.933 | 0.999 |
| HORVU.MOREX.r2.1HG0002970.1 | 0.331 | 0.284 | 3.427 | 0.083 | 0.934 | 0.999 |
| HORVU.MOREX.r2.1HG0001290.1 | 0.516 | 0.281 | 3.401 | 0.083 | 0.934 | 0.999 |
| HORVU.MOREX.r2.1HG0002490.1 | 3.850 | 0.271 | 1.109 | 0.245 | 0.807 | 0.999 |
| HORVU.MOREX.r2.1HG0002860.1 | 210.764 | 0.271 | 0.320 | 0.847 | 0.397 | 0.999 |
| HORVU.MOREX.r2.1HG0002050.1 | 0.315 | 0.265 | 3.434 | 0.077 | 0.938 | 0.999 |
| HORVU.MOREX.r2.1HG0005400.1 | 263.994 | 0.263 | 0.423 | 0.623 | 0.533 | 0.999 |
| HORVU.MOREX.r2.1HG0004880.1 | 0.421 | 0.260 | 3.433 | 0.076 | 0.940 | 0.999 |
| HORVU.MOREX.r2.1HG0002120.1 | 2.915 | 0.249 | 1.788 | 0.139 | 0.889 | 0.999 |
| HORVU.MOREX.r2.1HG0004700.1 | 18.187 | 0.234 | 0.620 | 0.378 | 0.706 | 0.999 |
| HORVU.MOREX.r2.1HG0003070.1 | 178.648 | 0.230 | 0.426 | 0.540 | 0.589 | 0.999 |
| HORVU.MOREX.r2.1HG0005670.1 | 0.729 | 0.222 | 3.352 | 0.066 | 0.947 | 0.999 |
| HORVU.MOREX.r2.1HG0006680.1 | 11.546 | 0.218 | 0.962 | 0.226 | 0.821 | 0.999 |
| HORVU.MOREX.r2.1HG0005080.1 | 33.396 | 0.216 | 0.437 | 0.495 | 0.621 | 0.999 |
| HORVU.MOREX.r2.1HG0004430.1 | 36.007 | 0.201 | 0.458 | 0.440 | 0.660 | 0.999 |
| HORVU.MOREX.r2.1HG0003170.1 | 0.545 | 0.198 | 2.967 | 0.067 | 0.947 | 0.999 |
| HORVU.MOREX.r2.1HG0004420.1 | 69.595 | 0.194 | 0.658 | 0.295 | 0.768 | 0.999 |
| HORVU.MOREX.r2.1HG0005340.1 | 1951.770 | 0.179 | 0.334 | 0.537 | 0.591 | 0.999 |
| HORVU.MOREX.r2.1HG0002620.1 | 64.728 | 0.176 | 0.405 | 0.435 | 0.664 | 0.999 |
| HORVU.MOREX.r2.1HG0001050.1 | 0.217 | 0.165 | 3.461 | 0.048 | 0.962 | 0.999 |
| HORVU.MOREX.r2.1HG0001870.1 | 798.170 | 0.161 | 0.273 | 0.592 | 0.554 | 0.999 |
| HORVU.MOREX.r2.1HG0004680.1 | 288.899 | 0.149 | 0.371 | 0.400 | 0.689 | 0.999 |
| HORVU.MOREX.r2.1HG0002840.1 | 1454.863 | 0.146 | 0.241 | 0.606 | 0.544 | 0.999 |
| HORVU.MOREX.r2.1HG0003010.1 | 139.107 | 0.139 | 0.350 | 0.396 | 0.692 | 0.999 |
| HORVU.MOREX.r2.1HG0001910.1 | 176.085 | 0.131 | 0.332 | 0.395 | 0.693 | 0.999 |
| HORVU.MOREX.r2.1HG0000280.1 | 3038.603 | 0.131 | 0.315 | 0.415 | 0.678 | 0.999 |
| HORVU.MOREX.r2.1HG0006250.1 | 1033.642 | 0.129 | 0.315 | 0.410 | 0.682 | 0.999 |
| HORVU.MOREX.r2.1HG0002630.1 | 0.211 | 0.126 | 3.462 | 0.036 | 0.971 | 0.999 |
| HORVU.MOREX.r2.1HG0001130.1 | 0.229 | 0.120 | 3.462 | 0.035 | 0.972 | 0.999 |
| HORVU.MOREX.r2.1HG0006020.1 | 11.634 | 0.118 | 0.825 | 0.144 | 0.886 | 0.999 |
| HORVU.MOREX.r2.1HG0000040.1 | 792.456 | 0.111 | 0.330 | 0.338 | 0.735 | 0.999 |
| HORVU.MOREX.r2.1HG0000840.1 | 17.818 | 0.111 | 0.681 | 0.163 | 0.871 | 0.999 |
| HORVU.MOREX.r2.1HG0003870.1 | 412.104 | 0.110 | 0.290 | 0.381 | 0.703 | 0.999 |
| HORVU.MOREX.r2.1HG0003210.1 | 375.731 | 0.101 | 0.268 | 0.376 | 0.707 | 0.999 |
| HORVU.MOREX.r2.1HG0004810.1 | 2369.651 | 0.094 | 0.269 | 0.349 | 0.727 | 0.999 |
| HORVU.MOREX.r2.1HG0002450.1 | 0.207 | 0.093 | 3.462 | 0.027 | 0.979 | 0.999 |
| HORVU.MOREX.r2.1HG0004010.1 | 5.405 | 0.092 | 1.715 | 0.053 | 0.957 | 0.999 |
| HORVU.MOREX.r2.1HG0004900.1 | 50.527 | 0.088 | 0.369 | 0.238 | 0.812 | 0.999 |
| HORVU.MOREX.r2.1HG0007120.1 | 311.970 | 0.087 | 0.606 | 0.144 | 0.885 | 0.999 |
| HORVU.MOREX.r2.1HG0002720.1 | 0.206 | 0.086 | 3.462 | 0.025 | 0.980 | 0.999 |
| HORVU.MOREX.r2.1HG0003800.1 | 0.206 | 0.085 | 3.462 | 0.025 | 0.980 | 0.999 |
| HORVU.MOREX.r2.1HG0003840.1 | 0.206 | 0.082 | 3.462 | 0.024 | 0.981 | 0.999 |
| HORVU.MOREX.r2.1HG0002990.1 | 0.205 | 0.080 | 3.462 | 0.023 | 0.982 | 0.999 |
| HORVU.MOREX.r2.1HG0001030.1 | 291.067 | 0.079 | 0.441 | 0.180 | 0.857 | 0.999 |
| HORVU.MOREX.r2.1HG0005640.1 | 26.877 | 0.079 | 0.588 | 0.135 | 0.893 | 0.999 |
| HORVU.MOREX.r2.1HG0003310.1 | 0.192 | 0.079 | 3.462 | 0.023 | 0.982 | 0.999 |
| HORVU.MOREX.r2.1HG0005950.1 | 0.205 | 0.079 | 3.462 | 0.023 | 0.982 | 0.999 |
| HORVU.MOREX.r2.1HG0000200.1 | 0.205 | 0.078 | 3.462 | 0.022 | 0.982 | 0.999 |
| HORVU.MOREX.r2.1HG0006130.1 | 0.211 | 0.078 | 3.462 | 0.022 | 0.982 | 0.999 |
| HORVU.MOREX.r2.1HG0005610.1 | 0.205 | 0.077 | 3.462 | 0.022 | 0.982 | 0.999 |
| HORVU.MOREX.r2.1HG0004600.1 | 0.211 | 0.077 | 3.462 | 0.022 | 0.982 | 0.999 |

| | | | | | | |
|---|---|---|---|---|---|---|
| HORVU.MOREX.r2.1HG0004630.1 | 313.095 | 0.076 | 0.430 | 0.178 | 0.859 | 0.999 |
| HORVU.MOREX.r2.1HG0006780.1 | 0.211 | 0.076 | 3.462 | 0.022 | 0.983 | 0.999 |
| HORVU.MOREX.r2.1HG0000410.1 | 0.211 | 0.075 | 3.462 | 0.022 | 0.983 | 0.999 |
| HORVU.MOREX.r2.1HG0001530.1 | 432.020 | 0.074 | 0.352 | 0.211 | 0.833 | 0.999 |
| HORVU.MOREX.r2.1HG0005600.1 | 0.210 | 0.070 | 3.462 | 0.020 | 0.984 | 0.999 |
| HORVU.MOREX.r2.1HG0001310.1 | 0.210 | 0.069 | 3.462 | 0.020 | 0.984 | 0.999 |
| HORVU.MOREX.r2.1HG0003300.1 | 11.740 | 0.068 | 0.699 | 0.098 | 0.922 | 0.999 |
| HORVU.MOREX.r2.1HG0005820.1 | 0.210 | 0.066 | 3.462 | 0.019 | 0.985 | 0.999 |
| HORVU.MOREX.r2.1HG0002610.1 | 2.971 | 0.062 | 2.346 | 0.027 | 0.979 | 0.999 |
| HORVU.MOREX.r2.1HG0004760.1 | 1716.706 | 0.061 | 0.325 | 0.188 | 0.851 | 0.999 |
| HORVU.MOREX.r2.1HG0005050.1 | 101.213 | 0.058 | 0.381 | 0.153 | 0.879 | 0.999 |
| HORVU.MOREX.r2.1HG0005770.1 | 94.442 | 0.057 | 0.578 | 0.099 | 0.921 | 0.999 |
| HORVU.MOREX.r2.1HG0003340.1 | 72.196 | 0.057 | 1.714 | 0.033 | 0.974 | 0.999 |
| HORVU.MOREX.r2.1HG0003220.1 | 126.229 | 0.053 | 0.332 | 0.159 | 0.874 | 0.999 |
| HORVU.MOREX.r2.1HG0002020.1 | 9235.663 | 0.053 | 0.348 | 0.152 | 0.880 | 0.999 |
| HORVU.MOREX.r2.1HG0002460.1 | 15.333 | 0.052 | 0.553 | 0.094 | 0.925 | 0.999 |
| HORVU.MOREX.r2.1HG0006280.1 | 862.033 | 0.052 | 0.281 | 0.185 | 0.853 | 0.999 |
| HORVU.MOREX.r2.1HG0003650.1 | 2.857 | 0.049 | 1.551 | 0.032 | 0.975 | 0.999 |
| HORVU.MOREX.r2.1HG0000120.1 | 194.023 | 0.048 | 0.288 | 0.168 | 0.867 | 0.999 |
| HORVU.MOREX.r2.1HG0006200.1 | 48.583 | 0.044 | 0.534 | 0.082 | 0.935 | 0.999 |
| HORVU.MOREX.r2.1HG0001920.1 | 99.596 | 0.036 | 0.443 | 0.081 | 0.935 | 0.999 |
| HORVU.MOREX.r2.1HG0001020.1 | 123.632 | 0.026 | 0.357 | 0.073 | 0.942 | 0.999 |
| HORVU.MOREX.r2.1HG0005300.1 | 467.002 | 0.026 | 0.256 | 0.101 | 0.920 | 0.999 |
| HORVU.MOREX.r2.1HG0001550.1 | 103.281 | 0.021 | 0.376 | 0.057 | 0.955 | 0.999 |
| HORVU.MOREX.r2.1HG0001540.1 | 7797.215 | 0.014 | 0.271 | 0.050 | 0.960 | 0.999 |
| HORVU.MOREX.r2.1HG0000960.1 | 165.160 | 0.013 | 0.288 | 0.044 | 0.965 | 0.999 |
| HORVU.MOREX.r2.1HG0000700.1 | 266.958 | 0.009 | 0.272 | 0.031 | 0.975 | 0.999 |
| HORVU.MOREX.r2.1HG0001720.1 | 375.058 | 0.004 | 0.253 | 0.015 | 0.988 | 0.999 |

Table S4

| Downregulated gene at T12 | baseMean | log2FC | lfcSE | stat | pvalue | padj |
|---|---|---|---|---|---|---|
| HORVU.MOREX.r2.1HG0028240.1 | 2.559 | -20.711 | 2.778 | -7.456 | 0.000 | 0.000 |
| HORVU.MOREX.r2.2HG0089230.1 | 2.239 | -20.651 | 2.904 | -7.112 | 0.000 | 0.000 |
| HORVU.MOREX.r2.2HG0115250.1 | 11.515 | -23.116 | 3.396 | -6.807 | 0.000 | 0.000 |
| HORVU.MOREX.r2.4HG0315140.1 | 1.919 | -20.233 | 3.060 | -6.612 | 0.000 | 0.000 |
| HORVU.MOREX.r2.1HG0010570.1 | 1.352 | -20.254 | 3.397 | -5.962 | 0.000 | 0.000 |
| HORVU.MOREX.r2.7HG0551580.1 | 0.795 | -18.670 | 3.399 | -5.493 | 0.000 | 0.000 |
| HORVU.MOREX.r2.2HG0081740.1 | 5208.128 | -7.775 | 1.627 | -4.780 | 0.000 | 0.002 |
| HORVU.MOREX.r2.1HG0004210.1 | 177.307 | -2.842 | 0.660 | -4.308 | 0.000 | 0.018 |
| HORVU.MOREX.r2.1HG0002580.1 | 152.317 | -11.500 | 3.272 | -3.514 | 0.000 | 0.418 |
| HORVU.MOREX.r2.1HG0004870.1 | 35.116 | -3.728 | 1.070 | -3.486 | 0.000 | 0.443 |
| HORVU.MOREX.r2.1HG0001760.1 | 41.442 | -7.360 | 2.114 | -3.482 | 0.000 | 0.443 |
| HORVU.MOREX.r2.1HG0004330.1 | 5.799 | -3.855 | 1.523 | -2.531 | 0.011 | 0.999 |
| HORVU.MOREX.r2.1HG0007020.1 | 72.317 | -0.904 | 0.575 | -1.572 | 0.116 | 0.999 |
| HORVU.MOREX.r2.1HG0006390.1 | 36.079 | -0.846 | 0.558 | -1.517 | 0.129 | 0.999 |
| HORVU.MOREX.r2.1HG0001730.1 | 64.846 | -0.558 | 0.392 | -1.422 | 0.155 | 0.999 |
| HORVU.MOREX.r2.1HG0005570.1 | 4066.813 | -0.341 | 0.255 | -1.337 | 0.181 | 0.999 |
| HORVU.MOREX.r2.1HG0004270.1 | 8.345 | -1.041 | 0.846 | -1.230 | 0.219 | 0.999 |
| HORVU.MOREX.r2.1HG0002210.1 | 7.264 | -1.682 | 1.426 | -1.180 | 0.238 | 0.999 |
| HORVU.MOREX.r2.1HG0007110.1 | 1945.905 | -0.668 | 0.580 | -1.151 | 0.250 | 0.999 |
| HORVU.MOREX.r2.1HG0000440.1 | 141.012 | -0.342 | 0.307 | -1.115 | 0.265 | 0.999 |
| HORVU.MOREX.r2.1HG0001700.1 | 3.617 | -1.441 | 1.311 | -1.099 | 0.272 | 0.999 |
| HORVU.MOREX.r2.1HG0004070.1 | 221.430 | -0.310 | 0.286 | -1.082 | 0.279 | 0.999 |
| HORVU.MOREX.r2.1HG0006350.1 | 2.967 | -1.452 | 1.356 | -1.071 | 0.284 | 0.999 |
| HORVU.MOREX.r2.1HG0003410.1 | 175.005 | -0.367 | 0.345 | -1.065 | 0.287 | 0.999 |
| HORVU.MOREX.r2.1HG0001460.1 | 1.032 | -3.154 | 2.972 | -1.062 | 0.288 | 0.999 |
| HORVU.MOREX.r2.1HG0001710.1 | 644.163 | -0.282 | 0.268 | -1.051 | 0.293 | 0.999 |
| HORVU.MOREX.r2.1HG0000580.1 | 55.680 | -0.355 | 0.338 | -1.051 | 0.293 | 0.999 |
| HORVU.MOREX.r2.1HG0003040.1 | 5.950 | -0.959 | 0.921 | -1.041 | 0.298 | 0.999 |
| HORVU.MOREX.r2.1HG0005960.1 | 20.468 | -0.537 | 0.517 | -1.039 | 0.299 | 0.999 |
| HORVU.MOREX.r2.1HG0004670.1 | 4.948 | -1.468 | 1.429 | -1.027 | 0.305 | 0.999 |
| HORVU.MOREX.r2.1HG0001400.1 | 3784.793 | -0.404 | 0.394 | -1.026 | 0.305 | 0.999 |
| HORVU.MOREX.r2.1HG0001980.1 | 126.929 | -0.311 | 0.308 | -1.007 | 0.314 | 0.999 |
| HORVU.MOREX.r2.1HG0006540.1 | 494.178 | -0.324 | 0.322 | -1.005 | 0.315 | 0.999 |
| HORVU.MOREX.r2.1HG0000110.1 | 81.950 | -0.327 | 0.328 | -0.998 | 0.318 | 0.999 |
| HORVU.MOREX.r2.1HG0004840.1 | 129.718 | -0.414 | 0.421 | -0.983 | 0.325 | 0.999 |
| HORVU.MOREX.r2.1HG0004040.1 | 6.814 | -0.965 | 0.983 | -0.982 | 0.326 | 0.999 |
| HORVU.MOREX.r2.1HG0001410.1 | 687.179 | -0.493 | 0.506 | -0.974 | 0.330 | 0.999 |
| HORVU.MOREX.r2.1HG0000730.1 | 1074.273 | -0.209 | 0.221 | -0.949 | 0.343 | 0.999 |
| HORVU.MOREX.r2.1HG0006840.1 | 1.182 | -2.201 | 2.338 | -0.941 | 0.347 | 0.999 |
| HORVU.MOREX.r2.1HG0004660.1 | 1.394 | -2.076 | 2.256 | -0.920 | 0.357 | 0.999 |
| HORVU.MOREX.r2.1HG0000660.1 | 0.895 | -2.451 | 2.710 | -0.904 | 0.366 | 0.999 |
| HORVU.MOREX.r2.1HG0004540.1 | 0.810 | -2.146 | 2.380 | -0.902 | 0.367 | 0.999 |
| HORVU.MOREX.r2.1HG0002090.1 | 23.343 | -0.731 | 0.818 | -0.893 | 0.372 | 0.999 |
| HORVU.MOREX.r2.1HG0004390.1 | 93.269 | -0.340 | 0.390 | -0.872 | 0.383 | 0.999 |
| HORVU.MOREX.r2.1HG0000030.1 | 2132.254 | -0.254 | 0.292 | -0.869 | 0.385 | 0.999 |
| HORVU.MOREX.r2.1HG0006310.1 | 0.921 | -2.751 | 3.169 | -0.868 | 0.385 | 0.999 |
| HORVU.MOREX.r2.1HG0000590.1 | 1615.887 | -0.338 | 0.394 | -0.857 | 0.391 | 0.999 |
| HORVU.MOREX.r2.1HG0002560.1 | 10481.774 | -0.328 | 0.386 | -0.852 | 0.394 | 0.999 |
| HORVU.MOREX.r2.1HG0003140.1 | 7.540 | -2.809 | 3.301 | -0.851 | 0.395 | 0.999 |
| HORVU.MOREX.r2.1HG0004200.1 | 104.280 | -0.404 | 0.488 | -0.828 | 0.408 | 0.999 |
| HORVU.MOREX.r2.1HG0004830.1 | 1.294 | -1.651 | 2.020 | -0.817 | 0.414 | 0.999 |
| HORVU.MOREX.r2.1HG0003620.1 | 182.864 | -0.893 | 1.095 | -0.815 | 0.415 | 0.999 |

| | | | | | | |
|---|---|---|---|---|---|---|
| HORVU.MOREX.r2.1HG0000360.1 | 0.727 | -2.712 | 3.392 | -0.800 | 0.424 | 0.999 |
| HORVU.MOREX.r2.1HG0006240.1 | 1287.263 | -0.244 | 0.306 | -0.799 | 0.424 | 0.999 |
| HORVU.MOREX.r2.1HG0003720.1 | 0.772 | -2.273 | 2.920 | -0.779 | 0.436 | 0.999 |
| HORVU.MOREX.r2.1HG0006720.1 | 508.014 | -0.273 | 0.353 | -0.775 | 0.439 | 0.999 |
| HORVU.MOREX.r2.1HG0003600.1 | 300.865 | -0.292 | 0.390 | -0.749 | 0.454 | 0.999 |
| HORVU.MOREX.r2.1HG0004080.1 | 202.278 | -0.405 | 0.551 | -0.734 | 0.463 | 0.999 |
| HORVU.MOREX.r2.1HG0000100.1 | 2.205 | -1.293 | 1.775 | -0.728 | 0.466 | 0.999 |
| HORVU.MOREX.r2.1HG0002410.1 | 256.030 | -0.336 | 0.466 | -0.720 | 0.471 | 0.999 |
| HORVU.MOREX.r2.1HG0002170.1 | 1353.051 | -0.328 | 0.456 | -0.718 | 0.473 | 0.999 |
| HORVU.MOREX.r2.1HG0001160.1 | 3363.669 | -0.308 | 0.432 | -0.713 | 0.476 | 0.999 |
| HORVU.MOREX.r2.1HG0006290.1 | 920.059 | -0.171 | 0.249 | -0.688 | 0.491 | 0.999 |
| HORVU.MOREX.r2.1HG0004380.1 | 0.607 | -2.311 | 3.400 | -0.680 | 0.497 | 0.999 |
| HORVU.MOREX.r2.1HG0005520.1 | 13.538 | -0.453 | 0.668 | -0.678 | 0.498 | 0.999 |
| HORVU.MOREX.r2.1HG0002010.1 | 61.326 | -0.524 | 0.775 | -0.676 | 0.499 | 0.999 |
| HORVU.MOREX.r2.1HG0001430.1 | 247.442 | -0.218 | 0.328 | -0.666 | 0.505 | 0.999 |
| HORVU.MOREX.r2.1HG0001580.1 | 302.045 | -0.236 | 0.366 | -0.645 | 0.519 | 0.999 |
| HORVU.MOREX.r2.1HG0002640.1 | 7.954 | -0.468 | 0.742 | -0.631 | 0.528 | 0.999 |
| HORVU.MOREX.r2.1HG0007130.1 | 0.786 | -1.493 | 2.370 | -0.630 | 0.529 | 0.999 |
| HORVU.MOREX.r2.1HG0000910.1 | 0.557 | -2.104 | 3.408 | -0.617 | 0.537 | 0.999 |
| HORVU.MOREX.r2.1HG0001480.1 | 0.683 | -2.074 | 3.379 | -0.614 | 0.539 | 0.999 |
| HORVU.MOREX.r2.1HG0006000.1 | 0.417 | -2.056 | 3.407 | -0.603 | 0.546 | 0.999 |
| HORVU.MOREX.r2.1HG0004960.1 | 57.219 | -0.345 | 0.577 | -0.598 | 0.550 | 0.999 |
| HORVU.MOREX.r2.1HG0005990.1 | 0.521 | -2.042 | 3.417 | -0.598 | 0.550 | 0.999 |
| HORVU.MOREX.r2.1HG0006900.1 | 8.256 | -0.525 | 0.906 | -0.580 | 0.562 | 0.999 |
| HORVU.MOREX.r2.1HG0000070.1 | 724.809 | -0.164 | 0.283 | -0.577 | 0.564 | 0.999 |
| HORVU.MOREX.r2.1HG0001640.1 | 71.390 | -0.281 | 0.494 | -0.568 | 0.570 | 0.999 |
| HORVU.MOREX.r2.1HG0004820.1 | 1224.590 | -0.215 | 0.385 | -0.559 | 0.576 | 0.999 |
| HORVU.MOREX.r2.1HG0002040.1 | 0.610 | -1.873 | 3.391 | -0.552 | 0.581 | 0.999 |
| HORVU.MOREX.r2.1HG0006820.1 | 55.824 | -0.281 | 0.511 | -0.551 | 0.581 | 0.999 |
| HORVU.MOREX.r2.1HG0000640.1 | 451.330 | -0.533 | 0.969 | -0.550 | 0.583 | 0.999 |
| HORVU.MOREX.r2.1HG0007070.1 | 67.070 | -0.244 | 0.455 | -0.538 | 0.591 | 0.999 |
| HORVU.MOREX.r2.1HG0000750.1 | 1.955 | -0.989 | 1.843 | -0.537 | 0.591 | 0.999 |
| HORVU.MOREX.r2.1HG0003750.1 | 98.457 | -0.197 | 0.368 | -0.535 | 0.593 | 0.999 |
| HORVU.MOREX.r2.1HG0007090.1 | 37.643 | -0.254 | 0.485 | -0.524 | 0.600 | 0.999 |
| HORVU.MOREX.r2.1HG0000740.1 | 1296.137 | -0.232 | 0.449 | -0.516 | 0.606 | 0.999 |
| HORVU.MOREX.r2.1HG0004340.1 | 0.884 | -1.153 | 2.382 | -0.484 | 0.628 | 0.999 |
| HORVU.MOREX.r2.1HG0002190.1 | 766.183 | -0.172 | 0.365 | -0.472 | 0.637 | 0.999 |
| HORVU.MOREX.r2.1HG0000870.1 | 325.639 | -0.155 | 0.331 | -0.467 | 0.640 | 0.999 |
| HORVU.MOREX.r2.1HG0000470.1 | 1023.671 | -0.113 | 0.248 | -0.453 | 0.650 | 0.999 |
| HORVU.MOREX.r2.1HG0000340.1 | 39.986 | -0.274 | 0.610 | -0.449 | 0.654 | 0.999 |
| HORVU.MOREX.r2.1HG0000720.1 | 263.204 | -0.152 | 0.347 | -0.437 | 0.662 | 0.999 |
| HORVU.MOREX.r2.1HG0000060.1 | 845.020 | -0.156 | 0.384 | -0.407 | 0.684 | 0.999 |
| HORVU.MOREX.r2.1HG0000350.1 | 18.771 | -0.274 | 0.692 | -0.396 | 0.692 | 0.999 |
| HORVU.MOREX.r2.1HG0005850.1 | 1436.424 | -0.164 | 0.427 | -0.385 | 0.700 | 0.999 |
| HORVU.MOREX.r2.1HG0003770.1 | 668.894 | -0.126 | 0.338 | -0.373 | 0.709 | 0.999 |
| HORVU.MOREX.r2.1HG0005800.1 | 1046.238 | -0.089 | 0.249 | -0.359 | 0.719 | 0.999 |
| HORVU.MOREX.r2.1HG0001790.1 | 0.747 | -1.197 | 3.398 | -0.352 | 0.725 | 0.999 |
| HORVU.MOREX.r2.1HG0003570.1 | 0.500 | -1.180 | 3.387 | -0.348 | 0.728 | 0.999 |
| HORVU.MOREX.r2.1HG0005540.1 | 0.664 | -1.185 | 3.404 | -0.348 | 0.728 | 0.999 |
| HORVU.MOREX.r2.1HG0003910.1 | 1614.449 | -0.145 | 0.420 | -0.345 | 0.730 | 0.999 |
| HORVU.MOREX.r2.1HG0006400.1 | 2224.910 | -0.086 | 0.253 | -0.338 | 0.735 | 0.999 |
| HORVU.MOREX.r2.1HG0005430.1 | 0.440 | -1.135 | 3.429 | -0.331 | 0.741 | 0.999 |
| HORVU.MOREX.r2.1HG0000450.1 | 49.108 | -0.174 | 0.529 | -0.329 | 0.742 | 0.999 |

| | | | | | | |
|---|---|---|---|---|---|---|
| HORVU.MOREX.r2.1HG0006810.1 | 0.415 | -1.127 | 3.433 | -0.328 | 0.743 | 0.999 |
| HORVU.MOREX.r2.1HG0006050.1 | 0.812 | -0.855 | 2.644 | -0.323 | 0.746 | 0.999 |
| HORVU.MOREX.r2.1HG0002330.1 | 430.225 | -0.152 | 0.496 | -0.307 | 0.759 | 0.999 |
| HORVU.MOREX.r2.1HG0002730.1 | 4.883 | -0.402 | 1.316 | -0.306 | 0.760 | 0.999 |
| HORVU.MOREX.r2.1HG0002180.1 | 204.441 | -0.082 | 0.272 | -0.303 | 0.762 | 0.999 |
| HORVU.MOREX.r2.1HG0006620.1 | 1830.998 | -0.071 | 0.236 | -0.301 | 0.763 | 0.999 |
| HORVU.MOREX.r2.1HG0003080.1 | 1535.624 | -0.079 | 0.271 | -0.291 | 0.771 | 0.999 |
| HORVU.MOREX.r2.1HG0007010.1 | 20.032 | -0.231 | 0.819 | -0.282 | 0.778 | 0.999 |
| HORVU.MOREX.r2.1HG0000090.1 | 1888.311 | -0.105 | 0.373 | -0.281 | 0.779 | 0.999 |
| HORVU.MOREX.r2.1HG0001690.1 | 302.634 | -0.067 | 0.243 | -0.276 | 0.783 | 0.999 |
| HORVU.MOREX.r2.1HG0005290.1 | 0.261 | -0.949 | 3.453 | -0.275 | 0.783 | 0.999 |
| HORVU.MOREX.r2.1HG0000180.1 | 0.249 | -0.927 | 3.453 | -0.268 | 0.788 | 0.999 |
| HORVU.MOREX.r2.1HG0001810.1 | 0.249 | -0.927 | 3.453 | -0.268 | 0.788 | 0.999 |
| HORVU.MOREX.r2.1HG0002310.1 | 0.249 | -0.927 | 3.453 | -0.268 | 0.788 | 0.999 |
| HORVU.MOREX.r2.1HG0005310.1 | 0.249 | -0.927 | 3.453 | -0.268 | 0.788 | 0.999 |
| HORVU.MOREX.r2.1HG0005510.1 | 0.249 | -0.927 | 3.453 | -0.268 | 0.788 | 0.999 |
| HORVU.MOREX.r2.1HG0005830.1 | 0.249 | -0.927 | 3.453 | -0.268 | 0.788 | 0.999 |
| HORVU.MOREX.r2.1HG0006140.1 | 0.249 | -0.927 | 3.453 | -0.268 | 0.788 | 0.999 |
| HORVU.MOREX.r2.1HG0000560.1 | 0.247 | -0.922 | 3.453 | -0.267 | 0.790 | 0.999 |
| HORVU.MOREX.r2.1HG0004130.1 | 5.002 | -0.244 | 0.987 | -0.247 | 0.805 | 0.999 |
| HORVU.MOREX.r2.1HG0004620.1 | 466.396 | -0.107 | 0.461 | -0.233 | 0.816 | 0.999 |
| HORVU.MOREX.r2.1HG0002200.1 | 33.196 | -0.106 | 0.481 | -0.220 | 0.826 | 0.999 |
| HORVU.MOREX.r2.1HG0005090.1 | 13.570 | -0.136 | 0.634 | -0.215 | 0.830 | 0.999 |
| HORVU.MOREX.r2.1HG0003670.1 | 0.180 | -0.741 | 3.459 | -0.214 | 0.830 | 0.999 |
| HORVU.MOREX.r2.1HG0006010.1 | 93.399 | -0.170 | 0.806 | -0.211 | 0.833 | 0.999 |
| HORVU.MOREX.r2.1HG0006590.1 | 0.173 | -0.724 | 3.460 | -0.209 | 0.834 | 0.999 |
| HORVU.MOREX.r2.1HG0000050.1 | 339.383 | -0.095 | 0.463 | -0.206 | 0.837 | 0.999 |
| HORVU.MOREX.r2.1HG0000140.1 | 0.166 | -0.710 | 3.460 | -0.205 | 0.837 | 0.999 |
| HORVU.MOREX.r2.1HG0000770.1 | 0.166 | -0.710 | 3.460 | -0.205 | 0.837 | 0.999 |
| HORVU.MOREX.r2.1HG0000890.1 | 0.166 | -0.710 | 3.460 | -0.205 | 0.837 | 0.999 |
| HORVU.MOREX.r2.1HG0001890.1 | 0.166 | -0.710 | 3.460 | -0.205 | 0.837 | 0.999 |
| HORVU.MOREX.r2.1HG0002000.1 | 0.166 | -0.710 | 3.460 | -0.205 | 0.837 | 0.999 |
| HORVU.MOREX.r2.1HG0003240.1 | 0.166 | -0.710 | 3.460 | -0.205 | 0.837 | 0.999 |
| HORVU.MOREX.r2.1HG0004580.1 | 0.166 | -0.710 | 3.460 | -0.205 | 0.837 | 0.999 |
| HORVU.MOREX.r2.1HG0004710.1 | 0.166 | -0.710 | 3.460 | -0.205 | 0.837 | 0.999 |
| HORVU.MOREX.r2.1HG0005880.1 | 0.166 | -0.710 | 3.460 | -0.205 | 0.837 | 0.999 |
| HORVU.MOREX.r2.1HG0006210.1 | 0.166 | -0.710 | 3.460 | -0.205 | 0.837 | 0.999 |
| HORVU.MOREX.r2.1HG0006550.1 | 0.166 | -0.710 | 3.460 | -0.205 | 0.837 | 0.999 |
| HORVU.MOREX.r2.1HG0007160.1 | 0.166 | -0.710 | 3.460 | -0.205 | 0.837 | 0.999 |
| HORVU.MOREX.r2.1HG0006170.1 | 0.155 | -0.683 | 3.461 | -0.197 | 0.844 | 0.999 |
| HORVU.MOREX.r2.1HG0002750.1 | 456.550 | -0.063 | 0.323 | -0.196 | 0.845 | 0.999 |
| HORVU.MOREX.r2.1HG0003940.1 | 0.605 | -0.644 | 3.386 | -0.190 | 0.849 | 0.999 |
| HORVU.MOREX.r2.1HG0005020.1 | 3.815 | -0.275 | 1.459 | -0.189 | 0.850 | 0.999 |
| HORVU.MOREX.r2.1HG0004860.1 | 0.284 | -0.633 | 3.448 | -0.183 | 0.854 | 0.999 |
| HORVU.MOREX.r2.1HG0006040.1 | 0.529 | -0.620 | 3.399 | -0.182 | 0.855 | 0.999 |
| HORVU.MOREX.r2.1HG0001250.1 | 25.858 | -0.091 | 0.511 | -0.178 | 0.859 | 0.999 |
| HORVU.MOREX.r2.1HG0001150.1 | 1.303 | -0.499 | 2.890 | -0.173 | 0.863 | 0.999 |
| HORVU.MOREX.r2.1HG0000170.1 | 0.576 | -0.563 | 3.301 | -0.171 | 0.864 | 0.999 |
| HORVU.MOREX.r2.1HG0002140.1 | 129.336 | -0.097 | 0.600 | -0.162 | 0.871 | 0.999 |
| HORVU.MOREX.r2.1HG0001590.1 | 20.184 | -0.109 | 0.680 | -0.161 | 0.872 | 0.999 |
| HORVU.MOREX.r2.1HG0006860.1 | 4.664 | -0.152 | 0.977 | -0.156 | 0.876 | 0.999 |
| HORVU.MOREX.r2.1HG0001300.1 | 0.383 | -0.529 | 3.432 | -0.154 | 0.877 | 0.999 |
| HORVU.MOREX.r2.1HG0000760.1 | 4.681 | -0.183 | 1.189 | -0.154 | 0.878 | 0.999 |

| | | | | | | |
|---|---|---|---|---|---|---|
| HORVU.MOREX.r2.1HG0006600.1 | 0.372 | -0.524 | 3.435 | -0.153 | 0.879 | 0.999 |
| HORVU.MOREX.r2.1HG0001200.1 | 0.714 | -0.502 | 3.391 | -0.148 | 0.882 | 0.999 |
| HORVU.MOREX.r2.1HG0000600.1 | 0.634 | -0.493 | 3.398 | -0.145 | 0.885 | 0.999 |
| HORVU.MOREX.r2.1HG0002350.1 | 77.703 | -0.064 | 0.459 | -0.139 | 0.890 | 0.999 |
| HORVU.MOREX.r2.1HG0003640.1 | 2.905 | -0.172 | 1.268 | -0.136 | 0.892 | 0.999 |
| HORVU.MOREX.r2.1HG0003130.1 | 0.444 | -0.456 | 3.418 | -0.134 | 0.894 | 0.999 |
| HORVU.MOREX.r2.1HG0001220.1 | 0.465 | -0.451 | 3.420 | -0.132 | 0.895 | 0.999 |
| HORVU.MOREX.r2.1HG0004000.1 | 0.782 | -0.437 | 3.342 | -0.131 | 0.896 | 0.999 |
| HORVU.MOREX.r2.1HG0003250.1 | 0.861 | -0.427 | 3.303 | -0.129 | 0.897 | 0.999 |
| HORVU.MOREX.r2.1HG0002880.1 | 0.626 | -0.425 | 3.365 | -0.126 | 0.899 | 0.999 |
| HORVU.MOREX.r2.1HG0007100.1 | 0.537 | -0.366 | 2.974 | -0.123 | 0.902 | 0.999 |
| HORVU.MOREX.r2.1HG0003760.1 | 107.350 | -0.039 | 0.323 | -0.120 | 0.904 | 0.999 |
| HORVU.MOREX.r2.1HG0001190.1 | 0.213 | -0.396 | 3.455 | -0.115 | 0.909 | 0.999 |
| HORVU.MOREX.r2.1HG0000130.1 | 0.494 | -0.382 | 3.392 | -0.113 | 0.910 | 0.999 |
| HORVU.MOREX.r2.1HG0007150.1 | 0.097 | -0.383 | 3.462 | -0.111 | 0.912 | 0.999 |
| HORVU.MOREX.r2.1HG0001180.1 | 0.083 | -0.383 | 3.462 | -0.111 | 0.912 | 0.999 |
| HORVU.MOREX.r2.1HG0001230.1 | 0.083 | -0.383 | 3.462 | -0.111 | 0.912 | 0.999 |
| HORVU.MOREX.r2.1HG0001340.1 | 0.083 | -0.383 | 3.462 | -0.111 | 0.912 | 0.999 |
| HORVU.MOREX.r2.1HG0002250.1 | 0.083 | -0.383 | 3.462 | -0.111 | 0.912 | 0.999 |
| HORVU.MOREX.r2.1HG0002320.1 | 0.083 | -0.383 | 3.462 | -0.111 | 0.912 | 0.999 |
| HORVU.MOREX.r2.1HG0002590.1 | 0.083 | -0.383 | 3.462 | -0.111 | 0.912 | 0.999 |
| HORVU.MOREX.r2.1HG0002930.1 | 0.083 | -0.383 | 3.462 | -0.111 | 0.912 | 0.999 |
| HORVU.MOREX.r2.1HG0003510.1 | 0.083 | -0.383 | 3.462 | -0.111 | 0.912 | 0.999 |
| HORVU.MOREX.r2.1HG0004650.1 | 0.083 | -0.383 | 3.462 | -0.111 | 0.912 | 0.999 |
| HORVU.MOREX.r2.1HG0005710.1 | 0.083 | -0.383 | 3.462 | -0.111 | 0.912 | 0.999 |
| HORVU.MOREX.r2.1HG0006300.1 | 0.083 | -0.383 | 3.462 | -0.111 | 0.912 | 0.999 |
| HORVU.MOREX.r2.1HG0006360.1 | 0.083 | -0.383 | 3.462 | -0.111 | 0.912 | 0.999 |
| HORVU.MOREX.r2.1HG0006450.1 | 0.083 | -0.383 | 3.462 | -0.111 | 0.912 | 0.999 |
| HORVU.MOREX.r2.1HG0006490.1 | 0.083 | -0.383 | 3.462 | -0.111 | 0.912 | 0.999 |
| HORVU.MOREX.r2.1HG0006760.1 | 0.083 | -0.383 | 3.462 | -0.111 | 0.912 | 0.999 |
| HORVU.MOREX.r2.1HG0006910.1 | 0.083 | -0.383 | 3.462 | -0.111 | 0.912 | 0.999 |
| HORVU.MOREX.r2.1HG0004060.1 | 805.566 | -0.029 | 0.281 | -0.103 | 0.918 | 0.999 |
| HORVU.MOREX.r2.1HG0000820.1 | 0.202 | -0.324 | 3.429 | -0.094 | 0.925 | 0.999 |
| HORVU.MOREX.r2.1HG0000850.1 | 1.136 | -0.203 | 2.202 | -0.092 | 0.926 | 0.999 |
| HORVU.MOREX.r2.1HG0004360.1 | 319.344 | -0.026 | 0.279 | -0.092 | 0.927 | 0.999 |
| HORVU.MOREX.r2.1HG0005350.1 | 26.041 | -0.050 | 0.547 | -0.091 | 0.927 | 0.999 |
| HORVU.MOREX.r2.1HG0000610.1 | 0.288 | -0.282 | 3.457 | -0.082 | 0.935 | 0.999 |
| HORVU.MOREX.r2.1HG0004250.1 | 200.563 | -0.046 | 0.580 | -0.080 | 0.937 | 0.999 |
| HORVU.MOREX.r2.1HG0004800.1 | 6.996 | -0.069 | 0.894 | -0.077 | 0.938 | 0.999 |
| HORVU.MOREX.r2.1HG0001950.1 | 0.294 | -0.265 | 3.461 | -0.077 | 0.939 | 0.999 |
| HORVU.MOREX.r2.1HG0005170.1 | 0.294 | -0.253 | 3.461 | -0.073 | 0.942 | 0.999 |
| HORVU.MOREX.r2.1HG0000670.1 | 133.307 | -0.025 | 0.347 | -0.073 | 0.942 | 0.999 |
| HORVU.MOREX.r2.1HG0002390.1 | 4.011 | -0.119 | 1.636 | -0.073 | 0.942 | 0.999 |
| HORVU.MOREX.r2.1HG0005500.1 | 0.410 | -0.202 | 3.415 | -0.059 | 0.953 | 0.999 |
| HORVU.MOREX.r2.1HG0005970.1 | 0.292 | -0.179 | 3.461 | -0.052 | 0.959 | 0.999 |
| HORVU.MOREX.r2.1HG0006570.1 | 8.426 | -0.040 | 0.881 | -0.045 | 0.964 | 0.999 |
| HORVU.MOREX.r2.1HG0004690.1 | 0.538 | -0.153 | 3.396 | -0.045 | 0.964 | 0.999 |
| HORVU.MOREX.r2.1HG0003280.1 | 1.307 | -0.078 | 2.293 | -0.034 | 0.973 | 0.999 |
| HORVU.MOREX.r2.1HG0003020.1 | 2.281 | -0.045 | 1.606 | -0.028 | 0.978 | 0.999 |
| HORVU.MOREX.r2.1HG0003050.1 | 232.253 | -0.008 | 0.291 | -0.027 | 0.979 | 0.999 |

Table S5

| Upregulated gene at T22 | baseMean | log2FC | lfcSE | stat | pvalue | padj |
|---|---|---|---|---|---|---|
| HORVU.MOREX.r2.2HG0120870.1 | 3.107 | 20.537 | 2.618 | 7.845 | 0.000 | 0.000 |
| HORVU.MOREX.r2.7HG0570690.1 | 11.346 | 22.081 | 3.215 | 6.868 | 0.000 | 0.000 |
| HORVU.MOREX.r2.1HG0000900.1 | 41.794 | 19.548 | 3.214 | 6.082 | 0.000 | 0.000 |
| HORVU.MOREX.r2.3HG0256040.1 | 1.216 | 19.203 | 3.218 | 5.967 | 0.000 | 0.000 |
| HORVU.MOREX.r2.5HG0437200.1 | 1.216 | 19.203 | 3.218 | 5.967 | 0.000 | 0.000 |
| HORVU.MOREX.r2.5HG0414890.1 | 0.945 | 18.856 | 3.219 | 5.857 | 0.000 | 0.000 |
| HORVU.MOREX.r2.6HG0492550.1 | 0.945 | 18.856 | 3.219 | 5.857 | 0.000 | 0.000 |
| HORVU.MOREX.r2.4HG0310140.1 | 0.945 | 18.856 | 3.219 | 5.857 | 0.000 | 0.000 |
| HORVU.MOREX.r2.3HG0207730.1 | 98.232 | 4.332 | 0.775 | 5.589 | 0.000 | 0.000 |
| HORVU.MOREX.r2.6HG0513160.1 | 158.888 | 3.939 | 0.724 | 5.441 | 0.000 | 0.000 |
| HORVU.MOREX.r2.2HG0141510.1 | 11.837 | 3.923 | 0.983 | 3.990 | 0.000 | 0.098 |
| HORVU.MOREX.r2.1HG0051960.1 | 74.282 | 2.804 | 0.703 | 3.988 | 0.000 | 0.098 |
| HORVU.MOREX.r2.2HG0080830.1 | 51.360 | 2.003 | 0.503 | 3.982 | 0.000 | 0.098 |
| HORVU.MOREX.r2.4HG0327490.1 | 99.861 | 1.713 | 0.432 | 3.968 | 0.000 | 0.099 |
| HORVU.MOREX.r2.5HG0399800.1 | 10.470 | 3.656 | 0.941 | 3.884 | 0.000 | 0.121 |
| HORVU.MOREX.r2.2HG0176560.1 | 192.332 | 12.134 | 3.126 | 3.881 | 0.000 | 0.121 |
| HORVU.MOREX.r2.4HG0329030.1 | 13.877 | 3.988 | 1.102 | 3.620 | 0.000 | 0.276 |
| HORVU.MOREX.r2.4HG0341130.1 | 97.537 | 1.925 | 0.532 | 3.618 | 0.000 | 0.276 |
| HORVU.MOREX.r2.3HG0183300.1 | 6263.035 | 1.949 | 0.564 | 3.454 | 0.001 | 0.458 |
| HORVU.MOREX.r2.3HG0183280.1 | 269.894 | 1.898 | 0.563 | 3.368 | 0.001 | 0.570 |
| HORVU.MOREX.r2.2HG0091930.1 | 89.220 | 2.228 | 0.662 | 3.367 | 0.001 | 0.570 |
| HORVU.MOREX.r2.5HG0429110.1 | 1509.937 | 1.829 | 0.552 | 3.312 | 0.001 | 0.646 |
| HORVU.MOREX.r2.6HG0455100.1 | 208.753 | 1.378 | 0.417 | 3.300 | 0.001 | 0.648 |
| HORVU.MOREX.r2.7HG0533680.1 | 77.992 | 2.025 | 0.623 | 3.253 | 0.001 | 0.694 |
| HORVU.MOREX.r2.3HG0265550.1 | 61.360 | 10.115 | 3.150 | 3.211 | 0.001 | 0.707 |
| HORVU.MOREX.r2.2HG0115620.1 | 479.670 | 1.945 | 0.618 | 3.148 | 0.002 | 0.820 |
| HORVU.MOREX.r2.4HG0323620.1 | 228.415 | 1.802 | 0.573 | 3.146 | 0.002 | 0.820 |
| HORVU.MOREX.r2.7HG0547790.1 | 1637.896 | 1.685 | 0.536 | 3.144 | 0.002 | 0.820 |
| HORVU.MOREX.r2.1HG0060460.1 | 1095.150 | 1.455 | 0.466 | 3.120 | 0.002 | 0.864 |
| HORVU.MOREX.r2.5HG0392490.1 | 119.536 | 1.846 | 0.593 | 3.114 | 0.002 | 0.868 |
| HORVU.MOREX.r2.5HG0405180.1 | 155.864 | 2.128 | 0.684 | 3.110 | 0.002 | 0.868 |
| HORVU.MOREX.r2.4HG0315810.1 | 356.566 | 1.598 | 0.517 | 3.093 | 0.002 | 0.892 |
| HORVU.MOREX.r2.1HG0001590.1 | 31.041 | 1.962 | 0.639 | 3.072 | 0.002 | 0.929 |
| HORVU.MOREX.r2.7HG0546260.1 | 73.660 | 1.706 | 0.557 | 3.064 | 0.002 | 0.929 |
| HORVU.MOREX.r2.5HG0399580.1 | 52.958 | 2.249 | 0.738 | 3.048 | 0.002 | 0.932 |
| HORVU.MOREX.r2.1HG0060420.1 | 342.663 | 1.822 | 0.599 | 3.043 | 0.002 | 0.936 |
| HORVU.MOREX.r2.7HG0569500.1 | 11.549 | 3.833 | 1.267 | 3.025 | 0.002 | 0.956 |
| HORVU.MOREX.r2.1HG0004080.1 | 172.903 | 1.375 | 0.499 | 2.757 | 0.006 | 1.000 |
| HORVU.MOREX.r2.1HG0005440.1 | 45.593 | 1.408 | 0.522 | 2.699 | 0.007 | 1.000 |
| HORVU.MOREX.r2.1HG0006820.1 | 72.837 | 1.170 | 0.545 | 2.146 | 0.032 | 1.000 |
| HORVU.MOREX.r2.1HG0002580.1 | 173.023 | 6.067 | 2.844 | 2.133 | 0.033 | 1.000 |
| HORVU.MOREX.r2.1HG0001760.1 | 26.728 | 1.764 | 0.900 | 1.961 | 0.050 | 1.000 |
| HORVU.MOREX.r2.1HG0007090.1 | 45.585 | 1.114 | 0.608 | 1.831 | 0.067 | 1.000 |
| HORVU.MOREX.r2.1HG0007840.1 | 45.091 | 1.562 | 0.874 | 1.787 | 0.074 | 1.000 |
| HORVU.MOREX.r2.1HG0000640.1 | 340.814 | 1.106 | 0.638 | 1.734 | 0.083 | 1.000 |
| HORVU.MOREX.r2.1HG0001570.1 | 5.436 | 2.009 | 1.170 | 1.717 | 0.086 | 1.000 |
| HORVU.MOREX.r2.1HG0000390.1 | 5.267 | 2.784 | 1.632 | 1.706 | 0.088 | 1.000 |
| HORVU.MOREX.r2.1HG0007220.1 | 48.794 | 0.929 | 0.550 | 1.690 | 0.091 | 1.000 |
| HORVU.MOREX.r2.1HG0005910.1 | 8.941 | 2.545 | 1.512 | 1.683 | 0.092 | 1.000 |
| HORVU.MOREX.r2.1HG0008920.1 | 18.433 | 1.058 | 0.632 | 1.674 | 0.094 | 1.000 |
| HORVU.MOREX.r2.1HG0000890.1 | 4.818 | 3.501 | 2.119 | 1.652 | 0.099 | 1.000 |
| HORVU.MOREX.r2.1HG0007120.1 | 270.650 | 0.774 | 0.480 | 1.615 | 0.106 | 1.000 |

| | | | | | | |
|---|---|---|---|---|---|---|
| HORVU.MOREX.r2.1HG0006680.1 | 10.154 | 1.250 | 0.818 | 1.529 | 0.126 | 1.000 |
| HORVU.MOREX.r2.1HG0007880.1 | 6.613 | 2.140 | 1.406 | 1.522 | 0.128 | 1.000 |
| HORVU.MOREX.r2.1HG0002870.1 | 6.625 | 1.465 | 0.998 | 1.468 | 0.142 | 1.000 |
| HORVU.MOREX.r2.1HG0006390.1 | 45.386 | 0.876 | 0.598 | 1.466 | 0.143 | 1.000 |
| HORVU.MOREX.r2.1HG0007110.1 | 1273.715 | 0.670 | 0.469 | 1.428 | 0.153 | 1.000 |
| HORVU.MOREX.r2.1HG0003500.1 | 1531.538 | 0.450 | 0.336 | 1.338 | 0.181 | 1.000 |
| HORVU.MOREX.r2.1HG0000930.1 | 103.508 | 3.001 | 2.554 | 1.175 | 0.240 | 1.000 |
| HORVU.MOREX.r2.1HG0005350.1 | 23.727 | 0.666 | 0.575 | 1.159 | 0.246 | 1.000 |
| HORVU.MOREX.r2.1HG0009120.1 | 22.224 | 0.632 | 0.555 | 1.139 | 0.255 | 1.000 |
| HORVU.MOREX.r2.1HG0009190.1 | 553.394 | 0.460 | 0.425 | 1.082 | 0.279 | 1.000 |
| HORVU.MOREX.r2.1HG0008430.1 | 454.130 | 0.311 | 0.287 | 1.082 | 0.279 | 1.000 |
| HORVU.MOREX.r2.1HG0003600.1 | 322.646 | 0.463 | 0.429 | 1.079 | 0.281 | 1.000 |
| HORVU.MOREX.r2.1HG0008790.1 | 4.879 | 1.460 | 1.355 | 1.078 | 0.281 | 1.000 |
| HORVU.MOREX.r2.1HG0003400.1 | 95.141 | 0.459 | 0.432 | 1.064 | 0.288 | 1.000 |
| HORVU.MOREX.r2.1HG0001580.1 | 294.736 | 0.334 | 0.316 | 1.057 | 0.291 | 1.000 |
| HORVU.MOREX.r2.1HG0000840.1 | 13.192 | 0.997 | 0.967 | 1.031 | 0.302 | 1.000 |
| HORVU.MOREX.r2.1HG0005820.1 | 1.121 | 2.921 | 2.919 | 1.001 | 0.317 | 1.000 |
| HORVU.MOREX.r2.1HG0001430.1 | 274.554 | 0.381 | 0.384 | 0.992 | 0.321 | 1.000 |
| HORVU.MOREX.r2.1HG0003010.1 | 124.499 | 0.353 | 0.356 | 0.992 | 0.321 | 1.000 |
| HORVU.MOREX.r2.1HG0006450.1 | 1.209 | 2.848 | 2.884 | 0.987 | 0.323 | 1.000 |
| HORVU.MOREX.r2.1HG0003410.1 | 179.549 | 0.372 | 0.378 | 0.984 | 0.325 | 1.000 |
| HORVU.MOREX.r2.1HG0008050.1 | 82.159 | 0.564 | 0.576 | 0.979 | 0.327 | 1.000 |
| HORVU.MOREX.r2.1HG0003530.1 | 19.599 | 0.578 | 0.593 | 0.975 | 0.330 | 1.000 |
| HORVU.MOREX.r2.1HG0002120.1 | 2.552 | 1.712 | 1.777 | 0.963 | 0.335 | 1.000 |
| HORVU.MOREX.r2.1HG0004490.1 | 4.102 | 1.439 | 1.506 | 0.956 | 0.339 | 1.000 |
| HORVU.MOREX.r2.1HG0008740.1 | 6.961 | 0.977 | 1.036 | 0.943 | 0.346 | 1.000 |
| HORVU.MOREX.r2.1HG0006020.1 | 13.527 | 0.643 | 0.683 | 0.941 | 0.347 | 1.000 |
| HORVU.MOREX.r2.1HG0001730.1 | 55.623 | 0.502 | 0.542 | 0.926 | 0.355 | 1.000 |
| HORVU.MOREX.r2.1HG0005310.1 | 1.037 | 2.511 | 2.758 | 0.910 | 0.363 | 1.000 |
| HORVU.MOREX.r2.1HG0000980.1 | 20.677 | 2.792 | 3.128 | 0.893 | 0.372 | 1.000 |
| HORVU.MOREX.r2.1HG0007540.1 | 6.379 | 0.912 | 1.036 | 0.880 | 0.379 | 1.000 |
| HORVU.MOREX.r2.1HG0003300.1 | 10.275 | 0.617 | 0.734 | 0.841 | 0.401 | 1.000 |
| HORVU.MOREX.r2.1HG0009320.1 | 76.001 | 0.311 | 0.371 | 0.839 | 0.401 | 1.000 |
| HORVU.MOREX.r2.1HG0006190.1 | 4.480 | 1.020 | 1.243 | 0.820 | 0.412 | 1.000 |
| HORVU.MOREX.r2.1HG0005090.1 | 13.636 | 0.579 | 0.721 | 0.803 | 0.422 | 1.000 |
| HORVU.MOREX.r2.1HG0003770.1 | 656.121 | 0.302 | 0.379 | 0.797 | 0.425 | 1.000 |
| HORVU.MOREX.r2.1HG0005230.1 | 0.744 | 2.478 | 3.224 | 0.769 | 0.442 | 1.000 |
| HORVU.MOREX.r2.1HG0004390.1 | 78.092 | 0.343 | 0.447 | 0.768 | 0.443 | 1.000 |
| HORVU.MOREX.r2.1HG0005570.1 | 3604.234 | 0.236 | 0.310 | 0.762 | 0.446 | 1.000 |
| HORVU.MOREX.r2.1HG0007850.1 | 9.797 | 0.548 | 0.725 | 0.755 | 0.450 | 1.000 |
| HORVU.MOREX.r2.1HG0002830.1 | 2.778 | 1.127 | 1.503 | 0.750 | 0.453 | 1.000 |
| HORVU.MOREX.r2.1HG0000500.1 | 0.535 | 2.418 | 3.224 | 0.750 | 0.453 | 1.000 |
| HORVU.MOREX.r2.1HG0002680.1 | 2.616 | 1.251 | 1.675 | 0.747 | 0.455 | 1.000 |
| HORVU.MOREX.r2.1HG0002790.1 | 14.089 | 0.486 | 0.660 | 0.737 | 0.461 | 1.000 |
| HORVU.MOREX.r2.1HG0001250.1 | 35.924 | 0.341 | 0.469 | 0.726 | 0.468 | 1.000 |
| HORVU.MOREX.r2.1HG0001290.1 | 0.703 | 2.339 | 3.229 | 0.724 | 0.469 | 1.000 |
| HORVU.MOREX.r2.1HG0007050.1 | 2.044 | 1.389 | 1.919 | 0.724 | 0.469 | 1.000 |
| HORVU.MOREX.r2.1HG0008750.1 | 3.210 | 1.256 | 1.754 | 0.716 | 0.474 | 1.000 |
| HORVU.MOREX.r2.1HG0003630.1 | 78.729 | 0.302 | 0.432 | 0.699 | 0.485 | 1.000 |
| HORVU.MOREX.r2.1HG0000110.1 | 103.999 | 0.409 | 0.586 | 0.698 | 0.485 | 1.000 |
| HORVU.MOREX.r2.1HG0009450.1 | 1.053 | 2.155 | 3.140 | 0.686 | 0.492 | 1.000 |
| HORVU.MOREX.r2.1HG0009910.1 | 233.778 | 0.256 | 0.380 | 0.673 | 0.501 | 1.000 |
| HORVU.MOREX.r2.1HG0002410.1 | 198.886 | 0.327 | 0.490 | 0.669 | 0.504 | 1.000 |

| | | | | | | |
|---|---|---|---|---|---|---|
| HORVU.MOREX.r2.1HG0007760.1 | 897.962 | 0.212 | 0.317 | 0.666 | 0.505 | 1.000 |
| HORVU.MOREX.r2.1HG0002860.1 | 215.948 | 0.261 | 0.405 | 0.645 | 0.519 | 1.000 |
| HORVU.MOREX.r2.1HG0005360.1 | 179.487 | 0.286 | 0.447 | 0.641 | 0.521 | 1.000 |
| HORVU.MOREX.r2.1HG0009110.1 | 196.354 | 0.202 | 0.319 | 0.634 | 0.526 | 1.000 |
| HORVU.MOREX.r2.1HG0009880.1 | 0.665 | 2.036 | 3.238 | 0.629 | 0.529 | 1.000 |
| HORVU.MOREX.r2.1HG0005930.1 | 0.719 | 2.029 | 3.237 | 0.627 | 0.531 | 1.000 |
| HORVU.MOREX.r2.1HG0002310.1 | 0.630 | 2.022 | 3.246 | 0.623 | 0.533 | 1.000 |
| HORVU.MOREX.r2.1HG0002620.1 | 78.970 | 0.347 | 0.557 | 0.623 | 0.534 | 1.000 |
| HORVU.MOREX.r2.1HG0006810.1 | 0.641 | 2.019 | 3.246 | 0.622 | 0.534 | 1.000 |
| HORVU.MOREX.r2.1HG0005520.1 | 12.200 | 0.481 | 0.784 | 0.613 | 0.540 | 1.000 |
| HORVU.MOREX.r2.1HG0003200.1 | 1.276 | 1.177 | 1.954 | 0.602 | 0.547 | 1.000 |
| HORVU.MOREX.r2.1HG0000130.1 | 0.439 | 1.936 | 3.247 | 0.596 | 0.551 | 1.000 |
| HORVU.MOREX.r2.1HG0001980.1 | 117.076 | 0.225 | 0.379 | 0.595 | 0.552 | 1.000 |
| HORVU.MOREX.r2.1HG0006980.1 | 0.437 | 1.926 | 3.246 | 0.593 | 0.553 | 1.000 |
| HORVU.MOREX.r2.1HG0006060.1 | 3.097 | 0.815 | 1.378 | 0.592 | 0.554 | 1.000 |
| HORVU.MOREX.r2.1HG0008310.1 | 1.815 | 1.017 | 1.723 | 0.590 | 0.555 | 1.000 |
| HORVU.MOREX.r2.1HG0003130.1 | 1.184 | 1.418 | 2.431 | 0.583 | 0.560 | 1.000 |
| HORVU.MOREX.r2.1HG0005960.1 | 21.249 | 0.312 | 0.550 | 0.567 | 0.570 | 1.000 |
| HORVU.MOREX.r2.1HG0002330.1 | 452.718 | 0.286 | 0.512 | 0.559 | 0.576 | 1.000 |
| HORVU.MOREX.r2.1HG0005600.1 | 0.405 | 1.765 | 3.239 | 0.545 | 0.586 | 1.000 |
| HORVU.MOREX.r2.1HG0007470.1 | 0.405 | 1.765 | 3.239 | 0.545 | 0.586 | 1.000 |
| HORVU.MOREX.r2.1HG0008570.1 | 0.405 | 1.765 | 3.239 | 0.545 | 0.586 | 1.000 |
| HORVU.MOREX.r2.1HG0006840.1 | 1.154 | 1.302 | 2.448 | 0.532 | 0.595 | 1.000 |
| HORVU.MOREX.r2.1HG0008100.1 | 14.736 | 0.440 | 0.834 | 0.528 | 0.597 | 1.000 |
| HORVU.MOREX.r2.1HG0002970.1 | 0.654 | 1.703 | 3.229 | 0.527 | 0.598 | 1.000 |
| HORVU.MOREX.r2.1HG0002590.1 | 0.585 | 1.682 | 3.211 | 0.524 | 0.600 | 1.000 |
| HORVU.MOREX.r2.1HG0004060.1 | 720.324 | 0.170 | 0.324 | 0.524 | 0.601 | 1.000 |
| HORVU.MOREX.r2.1HG0005400.1 | 273.827 | 0.201 | 0.392 | 0.515 | 0.607 | 1.000 |
| HORVU.MOREX.r2.1HG0008650.1 | 60.326 | 0.248 | 0.494 | 0.502 | 0.615 | 1.000 |
| HORVU.MOREX.r2.1HG0005300.1 | 455.844 | 0.148 | 0.297 | 0.500 | 0.617 | 1.000 |
| HORVU.MOREX.r2.1HG0006860.1 | 2.542 | 0.805 | 1.612 | 0.499 | 0.618 | 1.000 |
| HORVU.MOREX.r2.1HG0004230.1 | 125.302 | 0.161 | 0.332 | 0.486 | 0.627 | 1.000 |
| HORVU.MOREX.r2.1HG0007370.1 | 6.458 | 0.512 | 1.061 | 0.482 | 0.630 | 1.000 |
| HORVU.MOREX.r2.1HG0003080.1 | 1397.103 | 0.141 | 0.294 | 0.479 | 0.632 | 1.000 |
| HORVU.MOREX.r2.1HG0003180.1 | 0.625 | 1.530 | 3.218 | 0.476 | 0.634 | 1.000 |
| HORVU.MOREX.r2.1HG0008330.1 | 1481.753 | 0.137 | 0.291 | 0.470 | 0.638 | 1.000 |
| HORVU.MOREX.r2.1HG0004210.1 | 151.594 | 0.537 | 1.146 | 0.469 | 0.639 | 1.000 |
| HORVU.MOREX.r2.1HG0000030.1 | 2126.407 | 0.181 | 0.391 | 0.463 | 0.644 | 1.000 |
| HORVU.MOREX.r2.1HG0005390.1 | 6.972 | 0.444 | 0.980 | 0.453 | 0.651 | 1.000 |
| HORVU.MOREX.r2.1HG0009250.1 | 1361.862 | 0.153 | 0.341 | 0.450 | 0.653 | 1.000 |
| HORVU.MOREX.r2.1HG0009710.1 | 0.867 | 1.091 | 2.440 | 0.447 | 0.655 | 1.000 |
| HORVU.MOREX.r2.1HG0002140.1 | 94.971 | 0.177 | 0.405 | 0.438 | 0.662 | 1.000 |
| HORVU.MOREX.r2.1HG0002610.1 | 6.024 | 0.607 | 1.392 | 0.436 | 0.663 | 1.000 |
| HORVU.MOREX.r2.1HG0002190.1 | 731.340 | 0.170 | 0.391 | 0.434 | 0.664 | 1.000 |
| HORVU.MOREX.r2.1HG0000690.1 | 0.505 | 1.387 | 3.220 | 0.431 | 0.667 | 1.000 |
| HORVU.MOREX.r2.1HG0003870.1 | 369.352 | 0.138 | 0.326 | 0.425 | 0.671 | 1.000 |
| HORVU.MOREX.r2.1HG0000810.1 | 0.526 | 1.357 | 3.238 | 0.419 | 0.675 | 1.000 |
| HORVU.MOREX.r2.1HG0009060.1 | 0.392 | 1.339 | 3.233 | 0.414 | 0.679 | 1.000 |
| HORVU.MOREX.r2.1HG0008380.1 | 1016.986 | 0.141 | 0.342 | 0.412 | 0.680 | 1.000 |
| HORVU.MOREX.r2.1HG0003050.1 | 255.557 | 0.146 | 0.358 | 0.409 | 0.682 | 1.000 |
| HORVU.MOREX.r2.1HG0008840.1 | 112.739 | 0.154 | 0.382 | 0.402 | 0.688 | 1.000 |
| HORVU.MOREX.r2.1HG0005030.1 | 0.540 | 1.282 | 3.227 | 0.397 | 0.691 | 1.000 |
| HORVU.MOREX.r2.1HG0007550.1 | 0.787 | 1.211 | 3.066 | 0.395 | 0.693 | 1.000 |

| | | | | | | |
|---|---|---|---|---|---|---|
| HORVU.MOREX.r2.1HG0005050.1 | 111.298 | 0.145 | 0.372 | 0.391 | 0.696 | 1.000 |
| HORVU.MOREX.r2.1HG0004700.1 | 18.000 | 0.259 | 0.669 | 0.387 | 0.699 | 1.000 |
| HORVU.MOREX.r2.1HG0000060.1 | 761.307 | 0.145 | 0.382 | 0.381 | 0.704 | 1.000 |
| HORVU.MOREX.r2.1HG0008680.1 | 0.860 | 1.013 | 2.677 | 0.379 | 0.705 | 1.000 |
| HORVU.MOREX.r2.1HG0008400.1 | 0.501 | 1.177 | 3.229 | 0.365 | 0.715 | 1.000 |
| HORVU.MOREX.r2.1HG0009530.1 | 7.289 | 0.536 | 1.573 | 0.340 | 0.734 | 1.000 |
| HORVU.MOREX.r2.1HG0001170.1 | 0.448 | 1.098 | 3.259 | 0.337 | 0.736 | 1.000 |
| HORVU.MOREX.r2.1HG0000940.1 | 0.845 | 1.081 | 3.212 | 0.336 | 0.737 | 1.000 |
| HORVU.MOREX.r2.1HG0008420.1 | 267.788 | 0.118 | 0.353 | 0.334 | 0.738 | 1.000 |
| HORVU.MOREX.r2.1HG0009300.1 | 0.676 | 1.067 | 3.219 | 0.331 | 0.740 | 1.000 |
| HORVU.MOREX.r2.1HG0002240.1 | 0.582 | 1.056 | 3.224 | 0.328 | 0.743 | 1.000 |
| HORVU.MOREX.r2.1HG0008410.1 | 2152.090 | 0.120 | 0.367 | 0.326 | 0.744 | 1.000 |
| HORVU.MOREX.r2.1HG0007860.1 | 0.546 | 1.052 | 3.226 | 0.326 | 0.744 | 1.000 |
| HORVU.MOREX.r2.1HG0002990.1 | 0.496 | 1.043 | 3.218 | 0.324 | 0.746 | 1.000 |
| HORVU.MOREX.r2.1HG0000190.1 | 0.507 | 1.044 | 3.230 | 0.323 | 0.746 | 1.000 |
| HORVU.MOREX.r2.1HG0003560.1 | 0.507 | 1.044 | 3.230 | 0.323 | 0.746 | 1.000 |
| HORVU.MOREX.r2.1HG0007100.1 | 0.507 | 1.044 | 3.230 | 0.323 | 0.746 | 1.000 |
| HORVU.MOREX.r2.1HG0004510.1 | 0.586 | 1.038 | 3.223 | 0.322 | 0.747 | 1.000 |
| HORVU.MOREX.r2.1HG0000100.1 | 1.333 | 0.727 | 2.280 | 0.319 | 0.750 | 1.000 |
| HORVU.MOREX.r2.1HG0008850.1 | 29.332 | 0.151 | 0.479 | 0.317 | 0.752 | 1.000 |
| HORVU.MOREX.r2.1HG0002350.1 | 73.214 | 0.136 | 0.434 | 0.313 | 0.754 | 1.000 |
| HORVU.MOREX.r2.1HG0008670.1 | 0.534 | 1.000 | 3.194 | 0.313 | 0.754 | 1.000 |
| HORVU.MOREX.r2.1HG0001920.1 | 103.655 | 0.124 | 0.402 | 0.309 | 0.757 | 1.000 |
| HORVU.MOREX.r2.1HG0004520.1 | 6.315 | 0.462 | 1.551 | 0.298 | 0.766 | 1.000 |
| HORVU.MOREX.r2.1HG0001160.1 | 2812.191 | 0.147 | 0.501 | 0.294 | 0.769 | 1.000 |
| HORVU.MOREX.r2.1HG0003750.1 | 108.420 | 0.111 | 0.387 | 0.288 | 0.773 | 1.000 |
| HORVU.MOREX.r2.1HG0001860.1 | 0.545 | 0.868 | 3.096 | 0.281 | 0.779 | 1.000 |
| HORVU.MOREX.r2.1HG0000470.1 | 1141.970 | 0.099 | 0.356 | 0.278 | 0.781 | 1.000 |
| HORVU.MOREX.r2.1HG0004660.1 | 1.702 | 0.600 | 2.231 | 0.269 | 0.788 | 1.000 |
| HORVU.MOREX.r2.1HG0004680.1 | 316.337 | 0.104 | 0.389 | 0.267 | 0.789 | 1.000 |
| HORVU.MOREX.r2.1HG0007130.1 | 1.259 | 0.624 | 2.380 | 0.262 | 0.793 | 1.000 |
| HORVU.MOREX.r2.1HG0002730.1 | 2.410 | 0.557 | 2.128 | 0.262 | 0.793 | 1.000 |
| HORVU.MOREX.r2.1HG0000320.1 | 11.583 | 0.259 | 1.012 | 0.256 | 0.798 | 1.000 |
| HORVU.MOREX.r2.1HG0008320.1 | 54.642 | 0.175 | 0.690 | 0.254 | 0.799 | 1.000 |
| HORVU.MOREX.r2.1HG0004620.1 | 445.578 | 0.100 | 0.422 | 0.237 | 0.813 | 1.000 |
| HORVU.MOREX.r2.1HG0004250.1 | 177.331 | 0.109 | 0.460 | 0.237 | 0.813 | 1.000 |
| HORVU.MOREX.r2.1HG0001700.1 | 2.150 | 0.440 | 1.904 | 0.231 | 0.817 | 1.000 |
| HORVU.MOREX.r2.1HG0009560.1 | 0.625 | 0.679 | 2.937 | 0.231 | 0.817 | 1.000 |
| HORVU.MOREX.r2.1HG0004760.1 | 1904.094 | 0.086 | 0.373 | 0.230 | 0.818 | 1.000 |
| HORVU.MOREX.r2.1HG0006280.1 | 836.916 | 0.071 | 0.318 | 0.223 | 0.824 | 1.000 |
| HORVU.MOREX.r2.1HG0007770.1 | 505.980 | 0.081 | 0.363 | 0.222 | 0.824 | 1.000 |
| HORVU.MOREX.r2.1HG0000740.1 | 1096.492 | 0.094 | 0.430 | 0.219 | 0.827 | 1.000 |
| HORVU.MOREX.r2.1HG0000040.1 | 794.858 | 0.067 | 0.308 | 0.219 | 0.827 | 1.000 |
| HORVU.MOREX.r2.1HG0006310.1 | 0.414 | 0.700 | 3.223 | 0.217 | 0.828 | 1.000 |
| HORVU.MOREX.r2.1HG0001150.1 | 0.597 | 0.650 | 3.202 | 0.203 | 0.839 | 1.000 |
| HORVU.MOREX.r2.1HG0006400.1 | 2052.565 | 0.057 | 0.283 | 0.203 | 0.839 | 1.000 |
| HORVU.MOREX.r2.1HG0006200.1 | 50.211 | 0.090 | 0.452 | 0.199 | 0.842 | 1.000 |
| HORVU.MOREX.r2.1HG0003640.1 | 3.167 | 0.247 | 1.267 | 0.195 | 0.845 | 1.000 |
| HORVU.MOREX.r2.1HG0001850.1 | 1.483 | 0.490 | 2.514 | 0.195 | 0.846 | 1.000 |
| HORVU.MOREX.r2.1HG0001870.1 | 732.849 | 0.070 | 0.362 | 0.194 | 0.846 | 1.000 |
| HORVU.MOREX.r2.1HG0007340.1 | 0.727 | 0.581 | 3.043 | 0.191 | 0.849 | 1.000 |
| HORVU.MOREX.r2.1HG0007140.1 | 0.731 | 0.606 | 3.200 | 0.190 | 0.850 | 1.000 |
| HORVU.MOREX.r2.1HG0001550.1 | 112.355 | 0.080 | 0.421 | 0.189 | 0.850 | 1.000 |

| | | | | | | |
|---|---|---|---|---|---|---|
| HORVU.MOREX.r2.1HG0007010.1 | 18.774 | 0.130 | 0.694 | 0.188 | 0.851 | 1.000 |
| HORVU.MOREX.r2.1HG0006900.1 | 5.133 | 0.190 | 1.041 | 0.183 | 0.855 | 1.000 |
| HORVU.MOREX.r2.1HG0007740.1 | 56.391 | 0.104 | 0.572 | 0.182 | 0.855 | 1.000 |
| HORVU.MOREX.r2.1HG0000960.1 | 159.080 | 0.062 | 0.362 | 0.173 | 0.863 | 1.000 |
| HORVU.MOREX.r2.1HG0003580.1 | 0.850 | 0.545 | 3.170 | 0.172 | 0.864 | 1.000 |
| HORVU.MOREX.r2.1HG0003650.1 | 2.456 | 0.232 | 1.370 | 0.170 | 0.865 | 1.000 |
| HORVU.MOREX.r2.1HG0002490.1 | 3.354 | 0.257 | 1.549 | 0.166 | 0.869 | 1.000 |
| HORVU.MOREX.r2.1HG0005770.1 | 81.745 | 0.103 | 0.624 | 0.165 | 0.869 | 1.000 |
| HORVU.MOREX.r2.1HG0002800.1 | 0.672 | 0.503 | 3.214 | 0.156 | 0.876 | 1.000 |
| HORVU.MOREX.r2.1HG0008860.1 | 0.445 | 0.475 | 3.237 | 0.147 | 0.883 | 1.000 |
| HORVU.MOREX.r2.1HG0006880.1 | 8.872 | 0.145 | 1.002 | 0.145 | 0.885 | 1.000 |
| HORVU.MOREX.r2.1HG0008980.1 | 0.654 | 0.446 | 3.206 | 0.139 | 0.889 | 1.000 |
| HORVU.MOREX.r2.1HG0001400.1 | 3169.159 | 0.059 | 0.431 | 0.137 | 0.891 | 1.000 |
| HORVU.MOREX.r2.1HG0000730.1 | 1080.077 | 0.037 | 0.276 | 0.134 | 0.894 | 1.000 |
| HORVU.MOREX.r2.1HG0007670.1 | 656.606 | 0.050 | 0.382 | 0.132 | 0.895 | 1.000 |
| HORVU.MOREX.r2.1HG0008710.1 | 2.728 | 0.202 | 1.587 | 0.128 | 0.899 | 1.000 |
| HORVU.MOREX.r2.1HG0001130.1 | 0.475 | 0.407 | 3.218 | 0.127 | 0.899 | 1.000 |
| HORVU.MOREX.r2.1HG0006290.1 | 876.161 | 0.032 | 0.271 | 0.120 | 0.905 | 1.000 |
| HORVU.MOREX.r2.1HG0002130.1 | 0.389 | 0.366 | 3.224 | 0.114 | 0.910 | 1.000 |
| HORVU.MOREX.r2.1HG0001190.1 | 0.470 | 0.359 | 3.223 | 0.111 | 0.911 | 1.000 |
| HORVU.MOREX.r2.1HG0006530.1 | 0.809 | 0.337 | 3.201 | 0.105 | 0.916 | 1.000 |
| HORVU.MOREX.r2.1HG0006220.1 | 0.561 | 0.331 | 3.193 | 0.104 | 0.917 | 1.000 |
| HORVU.MOREX.r2.1HG0000280.1 | 2834.758 | 0.038 | 0.383 | 0.100 | 0.920 | 1.000 |
| HORVU.MOREX.r2.1HG0002460.1 | 17.625 | 0.068 | 0.679 | 0.100 | 0.921 | 1.000 |
| HORVU.MOREX.r2.1HG0003840.1 | 0.649 | 0.312 | 3.212 | 0.097 | 0.923 | 1.000 |
| HORVU.MOREX.r2.1HG0008830.1 | 3.466 | 0.127 | 1.385 | 0.092 | 0.927 | 1.000 |
| HORVU.MOREX.r2.1HG0000240.1 | 69.587 | 0.189 | 2.256 | 0.084 | 0.933 | 1.000 |
| HORVU.MOREX.r2.1HG0005080.1 | 34.361 | 0.043 | 0.517 | 0.083 | 0.934 | 1.000 |
| HORVU.MOREX.r2.1HG0008660.1 | 3.619 | 0.099 | 1.224 | 0.081 | 0.936 | 1.000 |
| HORVU.MOREX.r2.1HG0000910.1 | 25.679 | 0.186 | 2.313 | 0.080 | 0.936 | 1.000 |
| HORVU.MOREX.r2.1HG0005800.1 | 1034.295 | 0.022 | 0.312 | 0.072 | 0.943 | 1.000 |
| HORVU.MOREX.r2.1HG0004550.1 | 32.062 | 0.034 | 0.579 | 0.059 | 0.953 | 1.000 |
| HORVU.MOREX.r2.1HG0000120.1 | 178.276 | 0.020 | 0.344 | 0.059 | 0.953 | 1.000 |
| HORVU.MOREX.r2.1HG0004800.1 | 8.132 | 0.063 | 1.208 | 0.052 | 0.958 | 1.000 |
| HORVU.MOREX.r2.1HG0003480.1 | 30.366 | 0.038 | 0.740 | 0.051 | 0.959 | 1.000 |
| HORVU.MOREX.r2.1HG0006660.1 | 42.390 | 0.027 | 0.550 | 0.049 | 0.961 | 1.000 |
| HORVU.MOREX.r2.1HG0009630.1 | 144.330 | 0.021 | 0.435 | 0.048 | 0.961 | 1.000 |
| HORVU.MOREX.r2.1HG0001900.1 | 1.615 | 0.088 | 1.842 | 0.048 | 0.962 | 1.000 |
| HORVU.MOREX.r2.1HG0003260.1 | 5.495 | 0.059 | 1.289 | 0.046 | 0.964 | 1.000 |
| HORVU.MOREX.r2.1HG0004580.1 | 0.629 | 0.137 | 3.217 | 0.042 | 0.966 | 1.000 |
| HORVU.MOREX.r2.1HG0000050.1 | 292.947 | 0.022 | 0.528 | 0.041 | 0.967 | 1.000 |
| HORVU.MOREX.r2.1HG0007750.1 | 1.187 | 0.084 | 2.168 | 0.039 | 0.969 | 1.000 |
| HORVU.MOREX.r2.1HG0000210.1 | 0.838 | 0.122 | 3.174 | 0.038 | 0.969 | 1.000 |
| HORVU.MOREX.r2.1HG0009180.1 | 296.999 | 0.012 | 0.325 | 0.038 | 0.970 | 1.000 |
| HORVU.MOREX.r2.1HG0004160.1 | 0.683 | 0.121 | 3.206 | 0.038 | 0.970 | 1.000 |
| HORVU.MOREX.r2.1HG0001030.1 | 235.563 | 0.018 | 0.492 | 0.037 | 0.971 | 1.000 |
| HORVU.MOREX.r2.1HG0000610.1 | 0.734 | 0.089 | 3.214 | 0.028 | 0.978 | 1.000 |
| HORVU.MOREX.r2.1HG0002850.1 | 1.626 | 0.040 | 2.022 | 0.020 | 0.984 | 1.000 |
| HORVU.MOREX.r2.1HG0008080.1 | 6168.381 | 0.006 | 0.380 | 0.017 | 0.986 | 1.000 |
| HORVU.MOREX.r2.1HG0004200.1 | 118.568 | 0.008 | 0.555 | 0.014 | 0.989 | 1.000 |
| HORVU.MOREX.r2.1HG0004430.1 | 34.651 | 0.007 | 0.510 | 0.014 | 0.989 | 1.000 |
| HORVU.MOREX.r2.1HG0000490.1 | 12.467 | 0.014 | 1.041 | 0.013 | 0.989 | 1.000 |
| HORVU.MOREX.r2.1HG0002770.1 | 1.750 | 0.021 | 1.768 | 0.012 | 0.990 | 1.000 |

| | | | | | | |
|---|---|---|---|---|---|---|
| HORVU.MOREX.r2.1HG0002020.1 | 8590.754 | 0.005 | 0.414 | 0.011 | 0.991 | 1.000 |
| HORVU.MOREX.r2.1HG0008270.1 | 222.406 | 0.003 | 0.307 | 0.008 | 0.993 | 1.000 |
| HORVU.MOREX.r2.1HG0001690.1 | 308.456 | 0.002 | 0.282 | 0.006 | 0.995 | 1.000 |
| HORVU.MOREX.r2.1HG0000750.1 | 5.634 | 0.007 | 1.258 | 0.005 | 0.996 | 1.000 |
| HORVU.MOREX.r2.1HG0008070.1 | 1045.183 | 0.002 | 0.407 | 0.005 | 0.996 | 1.000 |
| HORVU.MOREX.r2.1HG0001790.1 | 0.494 | 0.014 | 3.224 | 0.004 | 0.997 | 1.000 |
| HORVU.MOREX.r2.1HG0004260.1 | 37.278 | 0.001 | 0.482 | 0.003 | 0.998 | 1.000 |
| HORVU.MOREX.r2.1HG0006370.1 | 93.985 | 0.001 | 0.413 | 0.002 | 0.998 | 1.000 |
| HORVU.MOREX.r2.1HG0005500.1 | 0.884 | 0.004 | 2.978 | 0.001 | 0.999 | 1.000 |

Table S6

| Downregulated gene at T22 | baseMean | log2FC | lfcSE | stat | pvalue | padj |
|---|---|---|---|---|---|---|
| HORVU.MOREX.r2.7HG0595970.1 | 2.613 | -20.359 | 2.724 | -7.474 | 0.000 | 0.000 |
| HORVU.MOREX.r2.2HG0177340.1 | 2.090 | -19.733 | 2.950 | -6.689 | 0.000 | 0.000 |
| HORVU.MOREX.r2.UnG0629660.1 | 6.359 | -21.500 | 3.215 | -6.687 | 0.000 | 0.000 |
| HORVU.MOREX.r2.3HG0266490.1 | 169.819 | -2.568 | 0.529 | -4.857 | 0.000 | 0.003 |
| HORVU.MOREX.r2.3HG0266310.1 | 11.257 | -4.331 | 0.954 | -4.540 | 0.000 | 0.012 |
| HORVU.MOREX.r2.7HG0579230.1 | 190.814 | -2.952 | 0.683 | -4.321 | 0.000 | 0.031 |
| HORVU.MOREX.r2.2HG0113870.1 | 171.666 | -2.995 | 0.718 | -4.168 | 0.000 | 0.057 |
| HORVU.MOREX.r2.7HG0551730.1 | 85.392 | -2.012 | 0.485 | -4.151 | 0.000 | 0.058 |
| HORVU.MOREX.r2.2HG0083880.1 | 50.164 | -2.261 | 0.555 | -4.078 | 0.000 | 0.075 |
| HORVU.MOREX.r2.6HG0456290.1 | 673.022 | -2.598 | 0.661 | -3.933 | 0.000 | 0.110 |
| HORVU.MOREX.r2.7HG0604840.1 | 164.170 | -1.413 | 0.361 | -3.920 | 0.000 | 0.112 |
| HORVU.MOREX.r2.2HG0149690.1 | 60.485 | -1.806 | 0.478 | -3.780 | 0.000 | 0.176 |
| HORVU.MOREX.r2.2HG0172430.1 | 85.801 | -2.666 | 0.711 | -3.747 | 0.000 | 0.194 |
| HORVU.MOREX.r2.4HG0283150.1 | 20.566 | -2.866 | 0.772 | -3.714 | 0.000 | 0.215 |
| HORVU.MOREX.r2.6HG0504080.1 | 6.478 | -5.201 | 1.436 | -3.623 | 0.000 | 0.276 |
| HORVU.MOREX.r2.5HG0446960.1 | 20.225 | -3.897 | 1.076 | -3.620 | 0.000 | 0.276 |
| HORVU.MOREX.r2.7HG0560100.1 | 51.623 | -2.193 | 0.615 | -3.567 | 0.000 | 0.325 |
| HORVU.MOREX.r2.5HG0359930.1 | 2673.547 | -1.075 | 0.303 | -3.551 | 0.000 | 0.337 |
| HORVU.MOREX.r2.7HG0614860.1 | 424.370 | -1.913 | 0.550 | -3.478 | 0.001 | 0.431 |
| HORVU.MOREX.r2.5HG0446540.1 | 24.573 | -2.393 | 0.695 | -3.442 | 0.001 | 0.467 |
| HORVU.MOREX.r2.2HG0081130.1 | 43.367 | -3.267 | 0.959 | -3.406 | 0.001 | 0.520 |
| HORVU.MOREX.r2.3HG0192770.1 | 345.321 | -1.343 | 0.401 | -3.349 | 0.001 | 0.595 |
| HORVU.MOREX.r2.5HG0369430.1 | 93.116 | -2.654 | 0.800 | -3.318 | 0.001 | 0.646 |
| HORVU.MOREX.r2.6HG0523360.1 | 195.181 | -5.557 | 1.680 | -3.307 | 0.001 | 0.646 |
| HORVU.MOREX.r2.5HG0428780.1 | 159.576 | -1.926 | 0.585 | -3.293 | 0.001 | 0.651 |
| HORVU.MOREX.r2.3HG0194300.1 | 162.095 | -1.356 | 0.416 | -3.258 | 0.001 | 0.694 |
| HORVU.MOREX.r2.7HG0527490.1 | 25.349 | -2.262 | 0.694 | -3.258 | 0.001 | 0.694 |
| HORVU.MOREX.r2.3HG0243630.1 | 65.246 | -1.667 | 0.514 | -3.246 | 0.001 | 0.694 |
| HORVU.MOREX.r2.7HG0611250.1 | 18.109 | -3.521 | 1.087 | -3.240 | 0.001 | 0.694 |
| HORVU.MOREX.r2.4HG0298730.1 | 113.259 | -1.689 | 0.521 | -3.240 | 0.001 | 0.694 |
| HORVU.MOREX.r2.3HG0192380.1 | 204.105 | -1.596 | 0.493 | -3.236 | 0.001 | 0.694 |
| HORVU.MOREX.r2.2HG0179350.1 | 11.277 | -2.965 | 0.918 | -3.229 | 0.001 | 0.694 |
| HORVU.MOREX.r2.5HG0400090.1 | 425.083 | -2.228 | 0.690 | -3.226 | 0.001 | 0.694 |
| HORVU.MOREX.r2.3HG0189450.1 | 15.386 | -2.905 | 0.903 | -3.218 | 0.001 | 0.701 |
| HORVU.MOREX.r2.5HG0373710.1 | 57.382 | -1.944 | 0.608 | -3.196 | 0.001 | 0.733 |
| HORVU.MOREX.r2.1HG0026880.1 | 29.695 | -3.255 | 1.031 | -3.158 | 0.002 | 0.820 |
| HORVU.MOREX.r2.1HG0039200.1 | 70.055 | -1.545 | 0.495 | -3.123 | 0.002 | 0.864 |
| HORVU.MOREX.r2.3HG0246070.1 | 3070.150 | -1.532 | 0.495 | -3.093 | 0.002 | 0.892 |
| HORVU.MOREX.r2.1HG0003890.1 | 19.103 | -9.557 | 3.101 | -3.082 | 0.002 | 0.913 |
| HORVU.MOREX.r2.7HG0598650.1 | 8.844 | -3.522 | 1.148 | -3.067 | 0.002 | 0.929 |
| HORVU.MOREX.r2.3HG0199730.1 | 9.581 | -2.858 | 0.935 | -3.059 | 0.002 | 0.929 |
| HORVU.MOREX.r2.2HG0115650.1 | 53.641 | -2.082 | 0.681 | -3.057 | 0.002 | 0.929 |
| HORVU.MOREX.r2.4HG0290380.1 | 28.126 | -1.909 | 0.626 | -3.050 | 0.002 | 0.932 |
| HORVU.MOREX.r2.5HG0399170.1 | 58.365 | -1.838 | 0.605 | -3.037 | 0.002 | 0.940 |
| HORVU.MOREX.r2.5HG0351950.1 | 28.349 | -2.743 | 0.906 | -3.027 | 0.002 | 0.956 |
| HORVU.MOREX.r2.1HG0006010.1 | 141.348 | -1.710 | 0.586 | -2.919 | 0.004 | 1.000 |
| HORVU.MOREX.r2.1HG0002210.1 | 5.669 | -3.301 | 1.331 | -2.480 | 0.013 | 1.000 |
| HORVU.MOREX.r2.1HG0007890.1 | 90.755 | -1.838 | 0.788 | -2.331 | 0.020 | 1.000 |
| HORVU.MOREX.r2.1HG0004870.1 | 92.442 | -3.292 | 1.567 | -2.101 | 0.036 | 1.000 |
| HORVU.MOREX.r2.1HG0007990.1 | 206.782 | -1.064 | 0.522 | -2.039 | 0.041 | 1.000 |
| HORVU.MOREX.r2.1HG0002010.1 | 43.186 | -1.464 | 0.731 | -2.002 | 0.045 | 1.000 |
| HORVU.MOREX.r2.1HG0008340.1 | 54.756 | -1.156 | 0.619 | -1.865 | 0.062 | 1.000 |

| | | | | | | |
|---|---|---|---|---|---|---|
| HORVU.MOREX.r2.1HG0007020.1 | 69.838 | -0.778 | 0.495 | -1.572 | 0.116 | 1.000 |
| HORVU.MOREX.r2.1HG0002780.1 | 3.477 | -2.473 | 1.634 | -1.514 | 0.130 | 1.000 |
| HORVU.MOREX.r2.1HG0003620.1 | 168.919 | -0.779 | 0.625 | -1.246 | 0.213 | 1.000 |
| HORVU.MOREX.r2.1HG0009240.1 | 44.913 | -0.703 | 0.572 | -1.229 | 0.219 | 1.000 |
| HORVU.MOREX.r2.1HG0007260.1 | 2666.724 | -1.626 | 1.328 | -1.224 | 0.221 | 1.000 |
| HORVU.MOREX.r2.1HG0008010.1 | 16.760 | -0.894 | 0.762 | -1.173 | 0.241 | 1.000 |
| HORVU.MOREX.r2.1HG0002390.1 | 10.091 | -1.216 | 1.036 | -1.173 | 0.241 | 1.000 |
| HORVU.MOREX.r2.1HG0009310.1 | 8.755 | -0.886 | 0.805 | -1.100 | 0.271 | 1.000 |
| HORVU.MOREX.r2.1HG0004130.1 | 4.270 | -1.414 | 1.315 | -1.075 | 0.282 | 1.000 |
| HORVU.MOREX.r2.1HG0000290.1 | 47.957 | -0.495 | 0.463 | -1.070 | 0.284 | 1.000 |
| HORVU.MOREX.r2.1HG0009050.1 | 3341.300 | -0.678 | 0.636 | -1.066 | 0.286 | 1.000 |
| HORVU.MOREX.r2.1HG0005640.1 | 20.589 | -0.594 | 0.588 | -1.010 | 0.312 | 1.000 |
| HORVU.MOREX.r2.1HG0001640.1 | 55.571 | -1.987 | 1.970 | -1.009 | 0.313 | 1.000 |
| HORVU.MOREX.r2.1HG0004420.1 | 122.455 | -1.148 | 1.195 | -0.960 | 0.337 | 1.000 |
| HORVU.MOREX.r2.1HG0004010.1 | 4.877 | -1.335 | 1.413 | -0.945 | 0.345 | 1.000 |
| HORVU.MOREX.r2.1HG0006430.1 | 1.296 | -2.434 | 2.602 | -0.935 | 0.350 | 1.000 |
| HORVU.MOREX.r2.1HG0008540.1 | 16.411 | -0.810 | 0.874 | -0.927 | 0.354 | 1.000 |
| HORVU.MOREX.r2.1HG0003720.1 | 0.968 | -2.863 | 3.102 | -0.923 | 0.356 | 1.000 |
| HORVU.MOREX.r2.1HG0001540.1 | 7549.488 | -0.255 | 0.278 | -0.917 | 0.359 | 1.000 |
| HORVU.MOREX.r2.1HG0008590.1 | 1.912 | -1.451 | 1.584 | -0.916 | 0.360 | 1.000 |
| HORVU.MOREX.r2.1HG0001410.1 | 627.239 | -0.494 | 0.540 | -0.916 | 0.360 | 1.000 |
| HORVU.MOREX.r2.1HG0004840.1 | 125.328 | -0.346 | 0.380 | -0.909 | 0.364 | 1.000 |
| HORVU.MOREX.r2.1HG0000450.1 | 63.314 | -0.483 | 0.536 | -0.901 | 0.368 | 1.000 |
| HORVU.MOREX.r2.1HG0006590.1 | 1.445 | -2.288 | 2.662 | -0.860 | 0.390 | 1.000 |
| HORVU.MOREX.r2.1HG0007650.1 | 8.261 | -1.235 | 1.461 | -0.845 | 0.398 | 1.000 |
| HORVU.MOREX.r2.1HG0000580.1 | 55.391 | -0.359 | 0.427 | -0.840 | 0.401 | 1.000 |
| HORVU.MOREX.r2.1HG0004960.1 | 62.618 | -0.390 | 0.466 | -0.835 | 0.404 | 1.000 |
| HORVU.MOREX.r2.1HG0003140.1 | 12.265 | -2.566 | 3.098 | -0.828 | 0.407 | 1.000 |
| HORVU.MOREX.r2.1HG0003340.1 | 76.895 | -1.508 | 1.827 | -0.825 | 0.409 | 1.000 |
| HORVU.MOREX.r2.1HG0003910.1 | 1645.966 | -0.354 | 0.434 | -0.817 | 0.414 | 1.000 |
| HORVU.MOREX.r2.1HG0009540.1 | 239.990 | -0.582 | 0.714 | -0.815 | 0.415 | 1.000 |
| HORVU.MOREX.r2.1HG0003760.1 | 110.056 | -0.265 | 0.328 | -0.809 | 0.419 | 1.000 |
| HORVU.MOREX.r2.1HG0009860.1 | 0.861 | -2.267 | 2.944 | -0.770 | 0.441 | 1.000 |
| HORVU.MOREX.r2.1HG0003270.1 | 23.205 | -0.441 | 0.576 | -0.765 | 0.444 | 1.000 |
| HORVU.MOREX.r2.1HG0008180.1 | 92.666 | -0.518 | 0.683 | -0.757 | 0.449 | 1.000 |
| HORVU.MOREX.r2.1HG0008870.1 | 22.016 | -0.429 | 0.568 | -0.755 | 0.450 | 1.000 |
| HORVU.MOREX.r2.1HG0000670.1 | 134.531 | -0.223 | 0.307 | -0.726 | 0.468 | 1.000 |
| HORVU.MOREX.r2.1HG0006510.1 | 89.463 | -1.244 | 1.742 | -0.714 | 0.475 | 1.000 |
| HORVU.MOREX.r2.1HG0001530.1 | 427.761 | -0.246 | 0.346 | -0.712 | 0.477 | 1.000 |
| HORVU.MOREX.r2.1HG0003740.1 | 217.291 | -0.289 | 0.412 | -0.702 | 0.483 | 1.000 |
| HORVU.MOREX.r2.1HG0000340.1 | 43.061 | -0.467 | 0.676 | -0.691 | 0.489 | 1.000 |
| HORVU.MOREX.r2.1HG0004220.1 | 2.156 | -1.129 | 1.639 | -0.689 | 0.491 | 1.000 |
| HORVU.MOREX.r2.1HG0008760.1 | 4.874 | -0.820 | 1.192 | -0.688 | 0.492 | 1.000 |
| HORVU.MOREX.r2.1HG0002070.1 | 2.318 | -1.252 | 1.846 | -0.678 | 0.498 | 1.000 |
| HORVU.MOREX.r2.1HG0009580.1 | 20.632 | -0.437 | 0.658 | -0.664 | 0.507 | 1.000 |
| HORVU.MOREX.r2.1HG0006250.1 | 1043.152 | -0.243 | 0.376 | -0.647 | 0.518 | 1.000 |
| HORVU.MOREX.r2.1HG0000440.1 | 147.037 | -0.285 | 0.447 | -0.637 | 0.524 | 1.000 |
| HORVU.MOREX.r2.1HG0003220.1 | 109.916 | -0.243 | 0.382 | -0.636 | 0.525 | 1.000 |
| HORVU.MOREX.r2.1HG0008020.1 | 12.650 | -0.498 | 0.802 | -0.621 | 0.535 | 1.000 |
| HORVU.MOREX.r2.1HG0007790.1 | 2292.678 | -0.256 | 0.413 | -0.619 | 0.536 | 1.000 |
| HORVU.MOREX.r2.1HG0006240.1 | 1186.402 | -0.214 | 0.362 | -0.591 | 0.554 | 1.000 |
| HORVU.MOREX.r2.1HG0002050.1 | 0.899 | -1.660 | 2.901 | -0.572 | 0.567 | 1.000 |
| HORVU.MOREX.r2.1HG0003860.1 | 1.517 | -1.068 | 1.868 | -0.571 | 0.568 | 1.000 |

| | | | | | | |
|---|---|---|---|---|---|---|
| HORVU.MOREX.r2.1HG0005340.1 | 1895.022 | -0.181 | 0.322 | -0.564 | 0.573 | 1.000 |
| HORVU.MOREX.r2.1HG0008950.1 | 483.736 | -0.205 | 0.363 | -0.564 | 0.573 | 1.000 |
| HORVU.MOREX.r2.1HG0000660.1 | 0.747 | -1.748 | 3.117 | -0.561 | 0.575 | 1.000 |
| HORVU.MOREX.r2.1HG0004070.1 | 200.699 | -0.191 | 0.350 | -0.545 | 0.586 | 1.000 |
| HORVU.MOREX.r2.1HG0006050.1 | 0.775 | -1.489 | 2.758 | -0.540 | 0.589 | 1.000 |
| HORVU.MOREX.r2.1HG0001710.1 | 559.953 | -0.177 | 0.329 | -0.537 | 0.591 | 1.000 |
| HORVU.MOREX.r2.1HG0000700.1 | 273.613 | -0.167 | 0.313 | -0.534 | 0.594 | 1.000 |
| HORVU.MOREX.r2.1HG0002180.1 | 195.753 | -0.185 | 0.349 | -0.531 | 0.596 | 1.000 |
| HORVU.MOREX.r2.1HG0003330.1 | 0.615 | -1.681 | 3.207 | -0.524 | 0.600 | 1.000 |
| HORVU.MOREX.r2.1HG0008730.1 | 36.202 | -0.284 | 0.544 | -0.521 | 0.602 | 1.000 |
| HORVU.MOREX.r2.1HG0001460.1 | 4.955 | -1.603 | 3.117 | -0.514 | 0.607 | 1.000 |
| HORVU.MOREX.r2.1HG0001100.1 | 0.537 | -1.614 | 3.232 | -0.499 | 0.617 | 1.000 |
| HORVU.MOREX.r2.1HG0001910.1 | 173.025 | -0.170 | 0.343 | -0.496 | 0.620 | 1.000 |
| HORVU.MOREX.r2.1HG0005850.1 | 1380.679 | -0.198 | 0.412 | -0.481 | 0.631 | 1.000 |
| HORVU.MOREX.r2.1HG0002170.1 | 1398.422 | -0.192 | 0.400 | -0.480 | 0.631 | 1.000 |
| HORVU.MOREX.r2.1HG0003040.1 | 4.902 | -0.639 | 1.334 | -0.479 | 0.632 | 1.000 |
| HORVU.MOREX.r2.1HG0001720.1 | 379.506 | -0.133 | 0.280 | -0.476 | 0.634 | 1.000 |
| HORVU.MOREX.r2.1HG0008580.1 | 9.787 | -0.377 | 0.808 | -0.467 | 0.641 | 1.000 |
| HORVU.MOREX.r2.1HG0003940.1 | 0.679 | -1.411 | 3.191 | -0.442 | 0.658 | 1.000 |
| HORVU.MOREX.r2.1HG0004050.1 | 72.846 | -0.230 | 0.525 | -0.438 | 0.661 | 1.000 |
| HORVU.MOREX.r2.1HG0005680.1 | 1.007 | -1.182 | 2.712 | -0.436 | 0.663 | 1.000 |
| HORVU.MOREX.r2.1HG0001470.1 | 5.445 | -0.526 | 1.219 | -0.431 | 0.666 | 1.000 |
| HORVU.MOREX.r2.1HG0004990.1 | 7.121 | -0.554 | 1.286 | -0.431 | 0.667 | 1.000 |
| HORVU.MOREX.r2.1HG0002910.1 | 0.387 | -1.361 | 3.236 | -0.421 | 0.674 | 1.000 |
| HORVU.MOREX.r2.1HG0003810.1 | 0.473 | -1.356 | 3.245 | -0.418 | 0.676 | 1.000 |
| HORVU.MOREX.r2.1HG0002090.1 | 146.641 | -0.568 | 1.368 | -0.415 | 0.678 | 1.000 |
| HORVU.MOREX.r2.1HG0006620.1 | 1737.928 | -0.102 | 0.262 | -0.389 | 0.697 | 1.000 |
| HORVU.MOREX.r2.1HG0002820.1 | 0.640 | -1.188 | 3.214 | -0.370 | 0.712 | 1.000 |
| HORVU.MOREX.r2.1HG0004670.1 | 5.093 | -0.375 | 1.034 | -0.362 | 0.717 | 1.000 |
| HORVU.MOREX.r2.1HG0000870.1 | 397.783 | -0.170 | 0.486 | -0.350 | 0.726 | 1.000 |
| HORVU.MOREX.r2.1HG0005020.1 | 1.634 | -0.675 | 1.929 | -0.350 | 0.726 | 1.000 |
| HORVU.MOREX.r2.1HG0001840.1 | 0.568 | -1.111 | 3.223 | -0.345 | 0.730 | 1.000 |
| HORVU.MOREX.r2.1HG0009620.1 | 0.549 | -1.092 | 3.222 | -0.339 | 0.735 | 1.000 |
| HORVU.MOREX.r2.1HG0006180.1 | 11.080 | -0.276 | 0.816 | -0.339 | 0.735 | 1.000 |
| HORVU.MOREX.r2.1HG0008890.1 | 0.389 | -1.078 | 3.227 | -0.334 | 0.738 | 1.000 |
| HORVU.MOREX.r2.1HG0004330.1 | 4.445 | -0.385 | 1.159 | -0.332 | 0.740 | 1.000 |
| HORVU.MOREX.r2.1HG0004340.1 | 0.387 | -1.065 | 3.231 | -0.330 | 0.742 | 1.000 |
| HORVU.MOREX.r2.1HG0009920.1 | 18.619 | -0.201 | 0.626 | -0.322 | 0.748 | 1.000 |
| HORVU.MOREX.r2.1HG0003070.1 | 195.964 | -0.135 | 0.423 | -0.319 | 0.750 | 1.000 |
| HORVU.MOREX.r2.1HG0003830.1 | 1.405 | -0.918 | 2.880 | -0.319 | 0.750 | 1.000 |
| HORVU.MOREX.r2.1HG0004270.1 | 8.056 | -0.276 | 0.884 | -0.313 | 0.755 | 1.000 |
| HORVU.MOREX.r2.1HG0009280.1 | 0.453 | -0.996 | 3.227 | -0.309 | 0.758 | 1.000 |
| HORVU.MOREX.r2.1HG0005860.1 | 74.933 | -0.142 | 0.463 | -0.307 | 0.759 | 1.000 |
| HORVU.MOREX.r2.1HG0004980.1 | 44.220 | -0.160 | 0.528 | -0.304 | 0.761 | 1.000 |
| HORVU.MOREX.r2.1HG0004630.1 | 271.421 | -0.120 | 0.400 | -0.301 | 0.764 | 1.000 |
| HORVU.MOREX.r2.1HG0000070.1 | 749.883 | -0.093 | 0.318 | -0.293 | 0.770 | 1.000 |
| HORVU.MOREX.r2.1HG0004040.1 | 1.839 | -0.501 | 1.750 | -0.286 | 0.775 | 1.000 |
| HORVU.MOREX.r2.1HG0006890.1 | 0.451 | -0.880 | 3.223 | -0.273 | 0.785 | 1.000 |
| HORVU.MOREX.r2.1HG0006350.1 | 2.456 | -0.391 | 1.435 | -0.273 | 0.785 | 1.000 |
| HORVU.MOREX.r2.1HG0000590.1 | 1523.403 | -0.127 | 0.465 | -0.272 | 0.785 | 1.000 |
| HORVU.MOREX.r2.1HG0007530.1 | 90.476 | -0.354 | 1.303 | -0.272 | 0.786 | 1.000 |
| HORVU.MOREX.r2.1HG0007330.1 | 64.496 | -0.109 | 0.402 | -0.272 | 0.786 | 1.000 |
| HORVU.MOREX.r2.1HG0003930.1 | 0.567 | -0.872 | 3.212 | -0.271 | 0.786 | 1.000 |

| | | | | | | |
|---|---|---|---|---|---|---|
| HORVU.MOREX.r2.1HG0008940.1 | 94.897 | -0.109 | 0.408 | -0.267 | 0.789 | 1.000 |
| HORVU.MOREX.r2.1HG0007040.1 | 0.995 | -0.851 | 3.210 | -0.265 | 0.791 | 1.000 |
| HORVU.MOREX.r2.1HG0004360.1 | 289.077 | -0.077 | 0.294 | -0.263 | 0.792 | 1.000 |
| HORVU.MOREX.r2.1HG0003190.1 | 0.956 | -0.616 | 2.393 | -0.257 | 0.797 | 1.000 |
| HORVU.MOREX.r2.1HG0002200.1 | 28.862 | -0.144 | 0.584 | -0.247 | 0.805 | 1.000 |
| HORVU.MOREX.r2.1HG0005670.1 | 0.419 | -0.797 | 3.241 | -0.246 | 0.806 | 1.000 |
| HORVU.MOREX.r2.1HG0003020.1 | 2.105 | -0.385 | 1.602 | -0.240 | 0.810 | 1.000 |
| HORVU.MOREX.r2.1HG0003370.1 | 0.398 | -0.770 | 3.245 | -0.237 | 0.812 | 1.000 |
| HORVU.MOREX.r2.1HG0000090.1 | 1656.078 | -0.089 | 0.381 | -0.233 | 0.816 | 1.000 |
| HORVU.MOREX.r2.1HG0007560.1 | 3.340 | -0.320 | 1.382 | -0.231 | 0.817 | 1.000 |
| HORVU.MOREX.r2.1HG0004440.1 | 24.184 | -0.142 | 0.625 | -0.226 | 0.821 | 1.000 |
| HORVU.MOREX.r2.1HG0007290.1 | 0.780 | -0.690 | 3.101 | -0.223 | 0.824 | 1.000 |
| HORVU.MOREX.r2.1HG0002840.1 | 1487.641 | -0.061 | 0.275 | -0.222 | 0.824 | 1.000 |
| HORVU.MOREX.r2.1HG0001960.1 | 0.404 | -0.706 | 3.240 | -0.218 | 0.828 | 1.000 |
| HORVU.MOREX.r2.1HG0005040.1 | 161.540 | -0.084 | 0.410 | -0.206 | 0.837 | 1.000 |
| HORVU.MOREX.r2.1HG0008470.1 | 0.383 | -0.587 | 3.224 | -0.182 | 0.855 | 1.000 |
| HORVU.MOREX.r2.1HG0001020.1 | 106.662 | -0.067 | 0.372 | -0.179 | 0.858 | 1.000 |
| HORVU.MOREX.r2.1HG0008090.1 | 0.466 | -0.540 | 3.223 | -0.168 | 0.867 | 1.000 |
| HORVU.MOREX.r2.1HG0009290.1 | 0.712 | -0.531 | 3.196 | -0.166 | 0.868 | 1.000 |
| HORVU.MOREX.r2.1HG0007480.1 | 0.848 | -0.477 | 2.916 | -0.164 | 0.870 | 1.000 |
| HORVU.MOREX.r2.1HG0001750.1 | 0.475 | -0.523 | 3.222 | -0.162 | 0.871 | 1.000 |
| HORVU.MOREX.r2.1HG0000350.1 | 20.742 | -0.109 | 0.687 | -0.159 | 0.874 | 1.000 |
| HORVU.MOREX.r2.1HG0002960.1 | 0.380 | -0.512 | 3.226 | -0.159 | 0.874 | 1.000 |
| HORVU.MOREX.r2.1HG0007940.1 | 0.380 | -0.512 | 3.226 | -0.159 | 0.874 | 1.000 |
| HORVU.MOREX.r2.1HG0008810.1 | 0.380 | -0.512 | 3.226 | -0.159 | 0.874 | 1.000 |
| HORVU.MOREX.r2.1HG0007300.1 | 0.531 | -0.503 | 3.213 | -0.156 | 0.876 | 1.000 |
| HORVU.MOREX.r2.1HG0003210.1 | 351.316 | -0.045 | 0.290 | -0.155 | 0.877 | 1.000 |
| HORVU.MOREX.r2.1HG0006570.1 | 6.982 | -0.124 | 0.835 | -0.149 | 0.882 | 1.000 |
| HORVU.MOREX.r2.1HG0004830.1 | 0.389 | -0.477 | 3.225 | -0.148 | 0.883 | 1.000 |
| HORVU.MOREX.r2.1HG0008110.1 | 94.881 | -0.062 | 0.422 | -0.147 | 0.883 | 1.000 |
| HORVU.MOREX.r2.1HG0004560.1 | 0.855 | -0.380 | 2.668 | -0.142 | 0.887 | 1.000 |
| HORVU.MOREX.r2.1HG0003920.1 | 37.114 | -0.067 | 0.486 | -0.138 | 0.890 | 1.000 |
| HORVU.MOREX.r2.1HG0000170.1 | 0.413 | -0.442 | 3.235 | -0.137 | 0.891 | 1.000 |
| HORVU.MOREX.r2.1HG0007070.1 | 69.715 | -0.062 | 0.463 | -0.134 | 0.894 | 1.000 |
| HORVU.MOREX.r2.1HG0009480.1 | 0.492 | -0.410 | 3.219 | -0.127 | 0.899 | 1.000 |
| HORVU.MOREX.r2.1HG0003970.1 | 0.870 | -0.400 | 3.168 | -0.126 | 0.899 | 1.000 |
| HORVU.MOREX.r2.1HG0002100.1 | 0.394 | -0.402 | 3.222 | -0.125 | 0.901 | 1.000 |
| HORVU.MOREX.r2.1HG0003380.1 | 0.394 | -0.402 | 3.222 | -0.125 | 0.901 | 1.000 |
| HORVU.MOREX.r2.1HG0006780.1 | 0.394 | -0.402 | 3.222 | -0.125 | 0.901 | 1.000 |
| HORVU.MOREX.r2.1HG0006750.1 | 3.524 | -0.172 | 1.459 | -0.118 | 0.906 | 1.000 |
| HORVU.MOREX.r2.1HG0004820.1 | 1019.659 | -0.051 | 0.432 | -0.118 | 0.906 | 1.000 |
| HORVU.MOREX.r2.1HG0000760.1 | 4.053 | -0.130 | 1.132 | -0.114 | 0.909 | 1.000 |
| HORVU.MOREX.r2.1HG0004900.1 | 59.707 | -0.066 | 0.589 | -0.112 | 0.911 | 1.000 |
| HORVU.MOREX.r2.1HG0004810.1 | 2509.755 | -0.032 | 0.303 | -0.105 | 0.916 | 1.000 |
| HORVU.MOREX.r2.1HG0004240.1 | 1.417 | -0.270 | 2.593 | -0.104 | 0.917 | 1.000 |
| HORVU.MOREX.r2.1HG0004410.1 | 42.600 | -0.057 | 0.565 | -0.102 | 0.919 | 1.000 |
| HORVU.MOREX.r2.1HG0009650.1 | 0.545 | -0.322 | 3.219 | -0.100 | 0.920 | 1.000 |
| HORVU.MOREX.r2.1HG0002750.1 | 449.184 | -0.032 | 0.348 | -0.091 | 0.927 | 1.000 |
| HORVU.MOREX.r2.1HG0002160.1 | 1.330 | -0.227 | 2.509 | -0.091 | 0.928 | 1.000 |
| HORVU.MOREX.r2.1HG0000360.1 | 0.416 | -0.282 | 3.238 | -0.087 | 0.931 | 1.000 |
| HORVU.MOREX.r2.1HG0009740.1 | 1.857 | -0.132 | 1.618 | -0.082 | 0.935 | 1.000 |
| HORVU.MOREX.r2.1HG0002560.1 | 8977.240 | -0.026 | 0.396 | -0.067 | 0.947 | 1.000 |
| HORVU.MOREX.r2.1HG0003280.1 | 0.494 | -0.214 | 3.217 | -0.067 | 0.947 | 1.000 |

| | | | | | | |
|---|---|---|---|---|---|---|
| HORVU.MOREX.r2.1HG0007810.1 | 0.852 | -0.178 | 3.175 | -0.056 | 0.955 | 1.000 |
| HORVU.MOREX.r2.1HG0009590.1 | 6.417 | -0.062 | 1.162 | -0.053 | 0.958 | 1.000 |
| HORVU.MOREX.r2.1HG0000830.1 | 1.549 | -0.080 | 1.838 | -0.044 | 0.965 | 1.000 |
| HORVU.MOREX.r2.1HG0002640.1 | 8.508 | -0.042 | 1.032 | -0.041 | 0.968 | 1.000 |
| HORVU.MOREX.r2.1HG0006540.1 | 454.276 | -0.014 | 0.361 | -0.040 | 0.968 | 1.000 |
| HORVU.MOREX.r2.1HG0009030.1 | 10.471 | -0.030 | 0.886 | -0.034 | 0.973 | 1.000 |
| HORVU.MOREX.r2.1HG0008450.1 | 30.533 | -0.014 | 0.506 | -0.027 | 0.979 | 1.000 |
| HORVU.MOREX.r2.1HG0002230.1 | 0.396 | -0.086 | 3.217 | -0.027 | 0.979 | 1.000 |
| HORVU.MOREX.r2.1HG0006720.1 | 485.880 | -0.009 | 0.367 | -0.024 | 0.981 | 1.000 |
| HORVU.MOREX.r2.1HG0009330.1 | 2.622 | -0.030 | 1.329 | -0.023 | 0.982 | 1.000 |
| HORVU.MOREX.r2.1HG0003290.1 | 1.394 | -0.045 | 2.647 | -0.017 | 0.986 | 1.000 |
| HORVU.MOREX.r2.1HG0000720.1 | 306.917 | -0.006 | 0.466 | -0.014 | 0.989 | 1.000 |
| HORVU.MOREX.r2.1HG0006870.1 | 2.898 | -0.011 | 1.291 | -0.009 | 0.993 | 1.000 |
| HORVU.MOREX.r2.1HG0000080.1 | 132.977 | -0.003 | 0.414 | -0.007 | 0.994 | 1.000 |
| HORVU.MOREX.r2.1HG0007420.1 | 0.553 | -0.008 | 3.216 | -0.003 | 0.998 | 1.000 |
| HORVU.MOREX.r2.1HG0005810.1 | 1442.400 | -0.001 | 0.266 | -0.002 | 0.998 | 1.000 |

Table S7

| Upregulated genes at T42 | baseMean | log2FC | lfcSE | stat | pvalue | padj |
|---|---|---|---|---|---|---|
| HORVU.MOREX.r2.5HG0423840.1 | 155.575 | 3.339 | 0.704 | 4.747 | 0.000 | 0.004 |
| HORVU.MOREX.r2.4HG0343710.1 | 828.647 | 2.978 | 0.629 | 4.734 | 0.000 | 0.004 |
| HORVU.MOREX.r2.1HG0026860.1 | 38.963 | 7.341 | 1.567 | 4.684 | 0.000 | 0.005 |
| HORVU.MOREX.r2.5HG0392950.1 | 1220.573 | 4.206 | 0.924 | 4.550 | 0.000 | 0.007 |
| HORVU.MOREX.r2.4HG0328900.1 | 33.255 | 7.744 | 1.708 | 4.534 | 0.000 | 0.008 |
| HORVU.MOREX.r2.2HG0080580.1 | 704.138 | 3.793 | 0.848 | 4.475 | 0.000 | 0.009 |
| HORVU.MOREX.r2.5HG0439680.1 | 1028.911 | 3.819 | 0.854 | 4.471 | 0.000 | 0.009 |
| HORVU.MOREX.r2.6HG0524260.1 | 310.264 | 4.047 | 0.923 | 4.386 | 0.000 | 0.012 |
| HORVU.MOREX.r2.7HG0624010.1 | 1269.439 | 4.567 | 1.042 | 4.383 | 0.000 | 0.012 |
| HORVU.MOREX.r2.6HG0511060.1 | 139.335 | 3.496 | 0.804 | 4.349 | 0.000 | 0.014 |
| HORVU.MOREX.r2.3HG0183300.1 | 3700.996 | 3.445 | 0.801 | 4.299 | 0.000 | 0.016 |
| HORVU.MOREX.r2.2HG0091890.1 | 306.863 | 5.434 | 1.270 | 4.280 | 0.000 | 0.017 |
| HORVU.MOREX.r2.5HG0412080.1 | 302.873 | 4.128 | 0.965 | 4.276 | 0.000 | 0.017 |
| HORVU.MOREX.r2.7HG0610830.1 | 29.363 | 7.080 | 1.698 | 4.170 | 0.000 | 0.024 |
| HORVU.MOREX.r2.5HG0431600.1 | 649.356 | 3.503 | 0.841 | 4.166 | 0.000 | 0.024 |
| HORVU.MOREX.r2.2HG0081780.1 | 67.077 | 6.917 | 1.685 | 4.106 | 0.000 | 0.030 |
| HORVU.MOREX.r2.1HG0026880.1 | 18.092 | 7.595 | 1.853 | 4.098 | 0.000 | 0.030 |
| HORVU.MOREX.r2.5HG0399370.1 | 176.253 | 3.188 | 0.781 | 4.080 | 0.000 | 0.030 |
| HORVU.MOREX.r2.6HG0504970.1 | 151.303 | 3.780 | 0.927 | 4.079 | 0.000 | 0.030 |
| HORVU.MOREX.r2.5HG0367910.1 | 89.828 | 4.075 | 1.016 | 4.010 | 0.000 | 0.037 |
| HORVU.MOREX.r2.2HG0164800.1 | 26.982 | 7.278 | 1.855 | 3.924 | 0.000 | 0.051 |
| HORVU.MOREX.r2.2HG0112790.1 | 71.250 | 3.940 | 1.010 | 3.901 | 0.000 | 0.052 |
| HORVU.MOREX.r2.3HG0183930.1 | 228.177 | 3.683 | 0.945 | 3.898 | 0.000 | 0.052 |
| HORVU.MOREX.r2.2HG0101740.1 | 199.607 | 2.677 | 0.689 | 3.888 | 0.000 | 0.053 |
| HORVU.MOREX.r2.2HG0084440.1 | 51.591 | 4.117 | 1.076 | 3.825 | 0.000 | 0.065 |
| HORVU.MOREX.r2.6HG0458460.1 | 105.821 | 3.511 | 0.919 | 3.820 | 0.000 | 0.065 |
| HORVU.MOREX.r2.5HG0445920.1 | 37.692 | 6.428 | 1.696 | 3.791 | 0.000 | 0.071 |
| HORVU.MOREX.r2.1HG0015680.1 | 308.349 | 2.946 | 0.788 | 3.737 | 0.000 | 0.085 |
| HORVU.MOREX.r2.2HG0093850.1 | 41.421 | 3.800 | 1.018 | 3.732 | 0.000 | 0.085 |
| HORVU.MOREX.r2.6HG0513160.1 | 71.831 | 4.344 | 1.165 | 3.729 | 0.000 | 0.085 |
| HORVU.MOREX.r2.5HG0394350.1 | 269.211 | 3.423 | 0.920 | 3.719 | 0.000 | 0.085 |
| HORVU.MOREX.r2.5HG0412430.1 | 297.728 | 3.371 | 0.914 | 3.689 | 0.000 | 0.092 |
| HORVU.MOREX.r2.5HG0397720.1 | 45.458 | 3.703 | 1.007 | 3.678 | 0.000 | 0.093 |
| HORVU.MOREX.r2.4HG0346110.1 | 124.713 | 2.258 | 0.617 | 3.661 | 0.000 | 0.097 |
| HORVU.MOREX.r2.7HG0531330.1 | 1868.906 | 3.620 | 0.994 | 3.640 | 0.000 | 0.103 |
| HORVU.MOREX.r2.2HG0169080.1 | 2381.195 | 2.260 | 0.633 | 3.572 | 0.000 | 0.129 |
| HORVU.MOREX.r2.5HG0400750.1 | 560.307 | 3.605 | 1.014 | 3.553 | 0.000 | 0.134 |
| HORVU.MOREX.r2.4HG0341030.1 | 38.869 | 3.563 | 1.008 | 3.536 | 0.000 | 0.134 |
| HORVU.MOREX.r2.7HG0614380.1 | 20.829 | 6.034 | 1.708 | 3.532 | 0.000 | 0.134 |
| HORVU.MOREX.r2.7HG0547790.1 | 1185.164 | 3.257 | 0.923 | 3.529 | 0.000 | 0.134 |
| HORVU.MOREX.r2.1HG0051960.1 | 41.527 | 4.933 | 1.413 | 3.491 | 0.000 | 0.152 |
| HORVU.MOREX.r2.7HG0558860.1 | 257.179 | 2.448 | 0.705 | 3.471 | 0.001 | 0.161 |
| HORVU.MOREX.r2.4HG0346270.1 | 286.434 | 2.674 | 0.771 | 3.467 | 0.001 | 0.161 |
| HORVU.MOREX.r2.3HG0195300.1 | 24.339 | 6.338 | 1.845 | 3.436 | 0.001 | 0.177 |
| HORVU.MOREX.r2.5HG0392490.1 | 122.805 | 5.707 | 1.663 | 3.431 | 0.001 | 0.177 |
| HORVU.MOREX.r2.1HG0001160.1 | 1667.343 | 3.160 | 0.923 | 3.426 | 0.001 | 0.177 |
| HORVU.MOREX.r2.5HG0414090.1 | 434.671 | 3.139 | 0.920 | 3.411 | 0.001 | 0.178 |
| HORVU.MOREX.r2.1HG0068480.1 | 171.821 | 2.573 | 0.754 | 3.411 | 0.001 | 0.178 |
| HORVU.MOREX.r2.2HG0109510.1 | 74.691 | 3.276 | 0.967 | 3.388 | 0.001 | 0.184 |
| HORVU.MOREX.r2.6HG0518840.1 | 303.132 | 2.719 | 0.803 | 3.385 | 0.001 | 0.184 |
| HORVU.MOREX.r2.7HG0619340.1 | 69.813 | 2.721 | 0.804 | 3.382 | 0.001 | 0.184 |
| HORVU.MOREX.r2.3HG0186650.1 | 103.622 | 3.687 | 1.091 | 3.381 | 0.001 | 0.184 |

| | | | | | | |
|---|---|---|---|---|---|---|
| HORVU.MOREX.r2.5HG0408180.1 | 244.517 | 3.467 | 1.029 | 3.369 | 0.001 | 0.188 |
| HORVU.MOREX.r2.5HG0436730.1 | 1225.776 | 1.987 | 0.590 | 3.368 | 0.001 | 0.188 |
| HORVU.MOREX.r2.3HG0182560.1 | 7835.086 | 3.114 | 0.926 | 3.363 | 0.001 | 0.189 |
| HORVU.MOREX.r2.3HG0199680.1 | 190.403 | 2.213 | 0.661 | 3.347 | 0.001 | 0.195 |
| HORVU.MOREX.r2.7HG0623560.1 | 54.743 | 3.280 | 0.980 | 3.346 | 0.001 | 0.195 |
| HORVU.MOREX.r2.5HG0392720.1 | 130.508 | 3.198 | 0.956 | 3.345 | 0.001 | 0.195 |
| HORVU.MOREX.r2.6HG0474930.1 | 532.865 | 2.703 | 0.812 | 3.329 | 0.001 | 0.200 |
| HORVU.MOREX.r2.7HG0529150.1 | 95.384 | 3.813 | 1.149 | 3.318 | 0.001 | 0.200 |
| HORVU.MOREX.r2.UnG0630000.1 | 284.209 | 2.902 | 0.875 | 3.318 | 0.001 | 0.200 |
| HORVU.MOREX.r2.2HG0091860.1 | 323.256 | 4.060 | 1.224 | 3.317 | 0.001 | 0.200 |
| HORVU.MOREX.r2.5HG0367920.1 | 137.662 | 3.300 | 1.002 | 3.292 | 0.001 | 0.212 |
| HORVU.MOREX.r2.6HG0448250.1 | 2070.904 | 3.265 | 0.994 | 3.284 | 0.001 | 0.215 |
| HORVU.MOREX.r2.7HG0583590.1 | 1697.094 | 2.210 | 0.673 | 3.282 | 0.001 | 0.215 |
| HORVU.MOREX.r2.2HG0173420.1 | 932.557 | 3.197 | 0.975 | 3.278 | 0.001 | 0.216 |
| HORVU.MOREX.r2.5HG0400740.1 | 169.990 | 4.730 | 1.447 | 3.268 | 0.001 | 0.219 |
| HORVU.MOREX.r2.1HG0055490.1 | 2388.068 | 2.496 | 0.765 | 3.264 | 0.001 | 0.221 |
| HORVU.MOREX.r2.5HG0434900.1 | 22.490 | 5.575 | 1.715 | 3.251 | 0.001 | 0.228 |
| HORVU.MOREX.r2.4HG0282400.1 | 54.740 | 2.974 | 0.915 | 3.249 | 0.001 | 0.228 |
| HORVU.MOREX.r2.1HG0040480.1 | 99.037 | 3.437 | 1.061 | 3.239 | 0.001 | 0.230 |
| HORVU.MOREX.r2.7HG0587270.1 | 633.980 | 2.529 | 0.781 | 3.239 | 0.001 | 0.230 |
| HORVU.MOREX.r2.5HG0392230.1 | 778.357 | 3.320 | 1.025 | 3.238 | 0.001 | 0.230 |
| HORVU.MOREX.r2.5HG0418820.1 | 173.441 | 2.142 | 0.665 | 3.219 | 0.001 | 0.241 |
| HORVU.MOREX.r2.3HG0267820.1 | 1211.838 | 2.677 | 0.836 | 3.203 | 0.001 | 0.254 |
| HORVU.MOREX.r2.3HG0195530.1 | 22.814 | 5.860 | 1.835 | 3.193 | 0.001 | 0.260 |
| HORVU.MOREX.r2.7HG0603690.1 | 13.056 | 5.949 | 1.866 | 3.188 | 0.001 | 0.262 |
| HORVU.MOREX.r2.6HG0458780.1 | 2584.564 | 3.239 | 1.018 | 3.182 | 0.001 | 0.265 |
| HORVU.MOREX.r2.1HG0060470.1 | 201.263 | 1.989 | 0.635 | 3.132 | 0.002 | 0.309 |
| HORVU.MOREX.r2.1HG0078280.1 | 679.588 | 1.694 | 0.544 | 3.113 | 0.002 | 0.328 |
| HORVU.MOREX.r2.3HG0255980.1 | 257.731 | 2.826 | 0.911 | 3.102 | 0.002 | 0.334 |
| HORVU.MOREX.r2.2HG0166600.1 | 1464.884 | 2.639 | 0.851 | 3.100 | 0.002 | 0.334 |
| HORVU.MOREX.r2.2HG0091380.1 | 92.389 | 2.691 | 0.869 | 3.097 | 0.002 | 0.334 |
| HORVU.MOREX.r2.3HG0195780.1 | 38.358 | 2.732 | 0.886 | 3.082 | 0.002 | 0.346 |
| HORVU.MOREX.r2.7HG0533680.1 | 94.365 | 3.265 | 1.060 | 3.082 | 0.002 | 0.346 |
| HORVU.MOREX.r2.6HG0448330.1 | 376.352 | 5.513 | 1.793 | 3.075 | 0.002 | 0.349 |
| HORVU.MOREX.r2.2HG0167340.1 | 283.362 | 1.918 | 0.625 | 3.071 | 0.002 | 0.349 |
| HORVU.MOREX.r2.3HG0224130.1 | 25.872 | 3.096 | 1.008 | 3.070 | 0.002 | 0.349 |
| HORVU.MOREX.r2.5HG0431830.1 | 113.715 | 1.955 | 0.638 | 3.064 | 0.002 | 0.353 |
| HORVU.MOREX.r2.1HG0069660.1 | 56.206 | 3.595 | 1.175 | 3.061 | 0.002 | 0.354 |
| HORVU.MOREX.r2.5HG0446820.1 | 552.900 | 3.005 | 0.983 | 3.057 | 0.002 | 0.354 |
| HORVU.MOREX.r2.5HG0409600.1 | 107.363 | 3.648 | 1.193 | 3.057 | 0.002 | 0.354 |
| HORVU.MOREX.r2.2HG0179580.1 | 5434.524 | 2.798 | 0.916 | 3.054 | 0.002 | 0.354 |
| HORVU.MOREX.r2.3HG0254090.1 | 41.767 | 3.202 | 1.051 | 3.046 | 0.002 | 0.354 |
| HORVU.MOREX.r2.7HG0546260.1 | 43.922 | 2.988 | 0.982 | 3.044 | 0.002 | 0.354 |
| HORVU.MOREX.r2.4HG0336220.1 | 445.702 | 2.423 | 0.796 | 3.043 | 0.002 | 0.354 |
| HORVU.MOREX.r2.5HG0399450.1 | 317.308 | 2.874 | 0.944 | 3.043 | 0.002 | 0.354 |
| HORVU.MOREX.r2.7HG0562990.1 | 162.199 | 2.612 | 0.860 | 3.037 | 0.002 | 0.359 |
| HORVU.MOREX.r2.5HG0424170.1 | 121.710 | 2.483 | 0.821 | 3.024 | 0.002 | 0.371 |
| HORVU.MOREX.r2.1HG0017190.1 | 67.370 | 3.862 | 1.279 | 3.019 | 0.003 | 0.375 |
| HORVU.MOREX.r2.2HG0172730.1 | 113.351 | 3.083 | 1.025 | 3.009 | 0.003 | 0.376 |
| HORVU.MOREX.r2.3HG0259270.1 | 42.819 | 3.885 | 1.291 | 3.008 | 0.003 | 0.376 |
| HORVU.MOREX.r2.3HG0226770.1 | 9.907 | 5.645 | 1.879 | 3.004 | 0.003 | 0.376 |
| HORVU.MOREX.r2.5HG0447820.1 | 2099.535 | 2.559 | 0.852 | 3.004 | 0.003 | 0.376 |
| HORVU.MOREX.r2.3HG0266030.1 | 710.267 | 2.740 | 0.917 | 2.988 | 0.003 | 0.389 |

| | | | | | | |
|---|---|---|---|---|---|---|
| HORVU.MOREX.r2.3HG0227960.1 | 277.756 | 1.980 | 0.663 | 2.986 | 0.003 | 0.389 |
| HORVU.MOREX.r2.2HG0175030.1 | 375.448 | 1.956 | 0.655 | 2.985 | 0.003 | 0.389 |
| HORVU.MOREX.r2.4HG0308090.1 | 118.594 | 1.858 | 0.624 | 2.979 | 0.003 | 0.393 |
| HORVU.MOREX.r2.5HG0392250.1 | 191.301 | 2.795 | 0.939 | 2.976 | 0.003 | 0.393 |
| HORVU.MOREX.r2.7HG0558900.1 | 304.955 | 2.481 | 0.835 | 2.972 | 0.003 | 0.394 |
| HORVU.MOREX.r2.5HG0438030.1 | 42.407 | 3.098 | 1.042 | 2.971 | 0.003 | 0.394 |
| HORVU.MOREX.r2.5HG0422250.1 | 294.978 | 2.162 | 0.729 | 2.963 | 0.003 | 0.398 |
| HORVU.MOREX.r2.5HG0417820.1 | 362.796 | 1.665 | 0.562 | 2.963 | 0.003 | 0.398 |
| HORVU.MOREX.r2.5HG0434830.1 | 9691.713 | 1.962 | 0.662 | 2.962 | 0.003 | 0.398 |
| HORVU.MOREX.r2.7HG0547720.1 | 533.314 | 2.992 | 1.011 | 2.960 | 0.003 | 0.398 |
| HORVU.MOREX.r2.7HG0581730.1 | 38.464 | 2.556 | 0.864 | 2.958 | 0.003 | 0.399 |
| HORVU.MOREX.r2.7HG0558810.1 | 31.406 | 3.098 | 1.048 | 2.956 | 0.003 | 0.399 |
| HORVU.MOREX.r2.UnG0628380.1 | 15.063 | 5.480 | 1.863 | 2.942 | 0.003 | 0.415 |
| HORVU.MOREX.r2.3HG0195400.1 | 170.193 | 2.445 | 0.835 | 2.930 | 0.003 | 0.429 |
| HORVU.MOREX.r2.3HG0184570.1 | 106.273 | 2.004 | 0.685 | 2.924 | 0.003 | 0.431 |
| HORVU.MOREX.r2.3HG0234060.1 | 10.507 | 5.394 | 1.844 | 2.924 | 0.003 | 0.431 |
| HORVU.MOREX.r2.1HG0011540.1 | 824.326 | 3.205 | 1.098 | 2.918 | 0.004 | 0.433 |
| HORVU.MOREX.r2.5HG0393010.1 | 228.269 | 2.476 | 0.851 | 2.911 | 0.004 | 0.440 |
| HORVU.MOREX.r2.5HG0385430.1 | 10.108 | 5.535 | 1.904 | 2.907 | 0.004 | 0.443 |
| HORVU.MOREX.r2.1HG0042010.1 | 59.146 | 2.528 | 0.871 | 2.901 | 0.004 | 0.446 |
| HORVU.MOREX.r2.4HG0278920.1 | 367.871 | 2.291 | 0.792 | 2.893 | 0.004 | 0.446 |
| HORVU.MOREX.r2.2HG0083080.1 | 294.206 | 3.264 | 1.128 | 2.893 | 0.004 | 0.446 |
| HORVU.MOREX.r2.2HG0089430.1 | 362.436 | 2.372 | 0.820 | 2.892 | 0.004 | 0.446 |
| HORVU.MOREX.r2.7HG0539640.1 | 118.232 | 2.947 | 1.021 | 2.888 | 0.004 | 0.449 |
| HORVU.MOREX.r2.7HG0615970.1 | 403.845 | 2.527 | 0.877 | 2.881 | 0.004 | 0.457 |
| HORVU.MOREX.r2.3HG0246160.1 | 214.212 | 1.635 | 0.568 | 2.877 | 0.004 | 0.458 |
| HORVU.MOREX.r2.1HG0051760.1 | 62.303 | 3.145 | 1.094 | 2.874 | 0.004 | 0.458 |
| HORVU.MOREX.r2.1HG0076100.1 | 34.464 | 2.939 | 1.023 | 2.873 | 0.004 | 0.458 |
| HORVU.MOREX.r2.5HG0432010.1 | 133.022 | 2.093 | 0.731 | 2.862 | 0.004 | 0.470 |
| HORVU.MOREX.r2.7HG0613890.1 | 119.847 | 2.241 | 0.784 | 2.860 | 0.004 | 0.470 |
| HORVU.MOREX.r2.6HG0476120.1 | 115.895 | 2.608 | 0.916 | 2.846 | 0.004 | 0.488 |
| HORVU.MOREX.r2.4HG0318840.1 | 178.888 | 1.977 | 0.695 | 2.843 | 0.004 | 0.488 |
| HORVU.MOREX.r2.2HG0080620.1 | 10808.117 | 2.643 | 0.930 | 2.843 | 0.004 | 0.488 |
| HORVU.MOREX.r2.UnG0633590.1 | 240.454 | 3.582 | 1.266 | 2.829 | 0.005 | 0.504 |
| HORVU.MOREX.r2.3HG0191060.1 | 26.155 | 4.122 | 1.458 | 2.828 | 0.005 | 0.504 |
| HORVU.MOREX.r2.7HG0605030.1 | 82.713 | 7.004 | 2.481 | 2.823 | 0.005 | 0.509 |
| HORVU.MOREX.r2.2HG0163110.1 | 496.565 | 1.467 | 0.522 | 2.812 | 0.005 | 0.521 |
| HORVU.MOREX.r2.7HG0527370.1 | 3290.454 | 2.725 | 0.969 | 2.812 | 0.005 | 0.521 |
| HORVU.MOREX.r2.7HG0604430.1 | 201.184 | 2.710 | 0.964 | 2.810 | 0.005 | 0.521 |
| HORVU.MOREX.r2.6HG0455440.1 | 34.029 | 3.429 | 1.221 | 2.807 | 0.005 | 0.523 |
| HORVU.MOREX.r2.7HG0591390.1 | 280.164 | 2.645 | 0.943 | 2.805 | 0.005 | 0.524 |
| HORVU.MOREX.r2.1HG0027260.1 | 12.296 | 5.792 | 2.069 | 2.799 | 0.005 | 0.529 |
| HORVU.MOREX.r2.1HG0060880.1 | 269.810 | 2.330 | 0.838 | 2.782 | 0.005 | 0.542 |
| HORVU.MOREX.r2.1HG0007120.1 | 179.249 | 2.628 | 0.945 | 2.781 | 0.005 | 0.542 |
| HORVU.MOREX.r2.2HG0106420.1 | 48.223 | 2.229 | 0.801 | 2.781 | 0.005 | 0.542 |
| HORVU.MOREX.r2.5HG0401270.1 | 14823.205 | 2.360 | 0.849 | 2.781 | 0.005 | 0.542 |
| HORVU.MOREX.r2.1HG0007090.1 | 23.923 | 3.161 | 1.137 | 2.781 | 0.005 | 0.542 |
| HORVU.MOREX.r2.4HG0327010.1 | 34.118 | 3.199 | 1.151 | 2.780 | 0.005 | 0.542 |
| HORVU.MOREX.r2.3HG0192020.1 | 144.668 | 2.660 | 0.957 | 2.779 | 0.005 | 0.542 |
| HORVU.MOREX.r2.7HG0531360.1 | 222.809 | 2.878 | 1.037 | 2.776 | 0.006 | 0.542 |
| HORVU.MOREX.r2.3HG0273000.1 | 9.815 | 5.632 | 2.029 | 2.776 | 0.006 | 0.542 |
| HORVU.MOREX.r2.1HG0007110.1 | 1040.323 | 2.382 | 0.859 | 2.775 | 0.006 | 0.542 |
| HORVU.MOREX.r2.5HG0399360.1 | 755.535 | 1.760 | 0.635 | 2.774 | 0.006 | 0.542 |

| | | | | | | |
|---|---|---|---|---|---|---|
| HORVU.MOREX.r2.2HG0108210.1 | 6105.842 | 1.893 | 0.687 | 2.756 | 0.006 | 0.567 |
| HORVU.MOREX.r2.2HG0088540.1 | 47.962 | 3.074 | 1.116 | 2.753 | 0.006 | 0.567 |
| HORVU.MOREX.r2.7HG0607230.1 | 22.898 | 3.557 | 1.292 | 2.753 | 0.006 | 0.567 |
| HORVU.MOREX.r2.5HG0433970.1 | 301.214 | 2.449 | 0.890 | 2.750 | 0.006 | 0.567 |
| HORVU.MOREX.r2.5HG0446810.1 | 263.983 | 2.697 | 0.981 | 2.750 | 0.006 | 0.567 |
| HORVU.MOREX.r2.1HG0037520.1 | 1045.393 | 2.244 | 0.817 | 2.746 | 0.006 | 0.569 |
| HORVU.MOREX.r2.5HG0438820.1 | 61.621 | 2.718 | 0.992 | 2.741 | 0.006 | 0.573 |
| HORVU.MOREX.r2.2HG0144690.1 | 90.939 | 2.798 | 1.022 | 2.737 | 0.006 | 0.576 |
| HORVU.MOREX.r2.5HG0399590.1 | 198.432 | 2.682 | 0.982 | 2.732 | 0.006 | 0.584 |
| HORVU.MOREX.r2.6HG0495720.1 | 11.869 | 4.837 | 1.775 | 2.724 | 0.006 | 0.588 |
| HORVU.MOREX.r2.2HG0154140.1 | 1068.317 | 2.166 | 0.795 | 2.724 | 0.006 | 0.588 |
| HORVU.MOREX.r2.7HG0568910.1 | 8.119 | 5.843 | 2.147 | 2.722 | 0.006 | 0.588 |
| HORVU.MOREX.r2.5HG0349880.1 | 442.830 | 2.663 | 0.979 | 2.721 | 0.007 | 0.588 |
| HORVU.MOREX.r2.7HG0531380.1 | 47.433 | 4.624 | 1.703 | 2.716 | 0.007 | 0.594 |
| HORVU.MOREX.r2.2HG0108680.1 | 29.770 | 4.395 | 1.621 | 2.712 | 0.007 | 0.598 |
| HORVU.MOREX.r2.2HG0144150.1 | 978.968 | 2.433 | 0.897 | 2.711 | 0.007 | 0.598 |
| HORVU.MOREX.r2.4HG0287960.1 | 267.789 | 2.788 | 1.032 | 2.702 | 0.007 | 0.611 |
| HORVU.MOREX.r2.5HG0413910.1 | 142.236 | 2.289 | 0.849 | 2.698 | 0.007 | 0.615 |
| HORVU.MOREX.r2.6HG0508040.1 | 491.223 | 1.823 | 0.678 | 2.688 | 0.007 | 0.630 |
| HORVU.MOREX.r2.6HG0451160.1 | 91.246 | 3.061 | 1.140 | 2.686 | 0.007 | 0.630 |
| HORVU.MOREX.r2.6HG0504490.1 | 91.221 | 2.269 | 0.846 | 2.682 | 0.007 | 0.633 |
| HORVU.MOREX.r2.5HG0417090.1 | 6.752 | 5.490 | 2.052 | 2.676 | 0.007 | 0.637 |
| HORVU.MOREX.r2.3HG0238970.1 | 629.839 | 1.756 | 0.657 | 2.672 | 0.008 | 0.637 |
| HORVU.MOREX.r2.3HG0263870.1 | 379.249 | 2.382 | 0.892 | 2.670 | 0.008 | 0.637 |
| HORVU.MOREX.r2.6HG0524700.1 | 241.339 | 2.320 | 0.869 | 2.669 | 0.008 | 0.637 |
| HORVU.MOREX.r2.1HG0001400.1 | 2156.376 | 5.608 | 2.105 | 2.664 | 0.008 | 0.644 |
| HORVU.MOREX.r2.6HG0514460.1 | 37.664 | 2.636 | 0.992 | 2.657 | 0.008 | 0.654 |
| HORVU.MOREX.r2.3HG0226850.1 | 38.742 | 2.742 | 1.034 | 2.652 | 0.008 | 0.660 |
| HORVU.MOREX.r2.4HG0289280.1 | 60.141 | 4.535 | 1.714 | 2.646 | 0.008 | 0.669 |
| HORVU.MOREX.r2.7HG0532400.1 | 969.589 | 2.341 | 0.886 | 2.643 | 0.008 | 0.673 |
| HORVU.MOREX.r2.1HG0067240.1 | 606.841 | 2.137 | 0.812 | 2.633 | 0.008 | 0.687 |
| HORVU.MOREX.r2.4HG0276540.1 | 4196.299 | 2.091 | 0.795 | 2.630 | 0.009 | 0.688 |
| HORVU.MOREX.r2.4HG0339550.1 | 107.884 | 2.846 | 1.082 | 2.630 | 0.009 | 0.688 |
| HORVU.MOREX.r2.5HG0440710.1 | 25.352 | 2.484 | 0.946 | 2.626 | 0.009 | 0.688 |
| HORVU.MOREX.r2.1HG0029500.1 | 19.609 | 3.167 | 1.206 | 2.625 | 0.009 | 0.688 |
| HORVU.MOREX.r2.2HG0158630.1 | 709.010 | 2.144 | 0.818 | 2.622 | 0.009 | 0.691 |
| HORVU.MOREX.r2.6HG0519760.1 | 123.885 | 1.714 | 0.655 | 2.618 | 0.009 | 0.696 |
| HORVU.MOREX.r2.1HG0058210.1 | 232.905 | 1.810 | 0.693 | 2.614 | 0.009 | 0.702 |
| HORVU.MOREX.r2.1HG0000640.1 | 146.993 | 2.298 | 0.882 | 2.606 | 0.009 | 0.708 |
| HORVU.MOREX.r2.5HG0414110.1 | 7560.274 | 1.425 | 0.548 | 2.599 | 0.009 | 0.716 |
| HORVU.MOREX.r2.6HG0453490.1 | 7240.758 | 2.168 | 0.834 | 2.598 | 0.009 | 0.716 |
| HORVU.MOREX.r2.6HG0520370.1 | 7.462 | 5.277 | 2.033 | 2.596 | 0.009 | 0.716 |
| HORVU.MOREX.r2.5HG0398010.1 | 2352.930 | 1.955 | 0.754 | 2.595 | 0.009 | 0.716 |
| HORVU.MOREX.r2.UnG0635440.1 | 80.087 | 2.909 | 1.122 | 2.593 | 0.010 | 0.716 |
| HORVU.MOREX.r2.2HG0144950.1 | 19.326 | 3.000 | 1.160 | 2.586 | 0.010 | 0.724 |
| HORVU.MOREX.r2.4HG0346360.1 | 169.991 | 2.658 | 1.028 | 2.586 | 0.010 | 0.724 |
| HORVU.MOREX.r2.7HG0549680.1 | 79.812 | 1.744 | 0.675 | 2.582 | 0.010 | 0.729 |
| HORVU.MOREX.r2.1HG0015090.1 | 12.715 | 4.762 | 1.847 | 2.578 | 0.010 | 0.734 |
| HORVU.MOREX.r2.2HG0100050.1 | 70.259 | 3.475 | 1.349 | 2.576 | 0.010 | 0.736 |
| HORVU.MOREX.r2.4HG0276520.1 | 222.439 | 1.793 | 0.697 | 2.573 | 0.010 | 0.739 |
| HORVU.MOREX.r2.UnG0631950.1 | 1062.732 | 2.415 | 0.939 | 2.572 | 0.010 | 0.739 |
| HORVU.MOREX.r2.2HG0176920.1 | 7.405 | 5.565 | 2.166 | 2.569 | 0.010 | 0.740 |
| HORVU.MOREX.r2.4HG0319270.1 | 144.034 | 2.213 | 0.863 | 2.565 | 0.010 | 0.748 |

| | | | | | | |
|---|---|---|---|---|---|---|
| HORVU.MOREX.r2.3HG0224440.1 | 1491.314 | 1.426 | 0.556 | 2.563 | 0.010 | 0.749 |
| HORVU.MOREX.r2.6HG0516450.1 | 97.950 | 1.772 | 0.692 | 2.562 | 0.010 | 0.749 |
| HORVU.MOREX.r2.2HG0109440.1 | 32.284 | 3.353 | 1.311 | 2.557 | 0.011 | 0.751 |
| HORVU.MOREX.r2.2HG0085880.1 | 113.437 | 2.085 | 0.817 | 2.554 | 0.011 | 0.753 |
| HORVU.MOREX.r2.2HG0097180.1 | 118.830 | 1.628 | 0.638 | 2.553 | 0.011 | 0.753 |
| HORVU.MOREX.r2.1HG0007220.1 | 31.228 | 3.547 | 1.392 | 2.549 | 0.011 | 0.759 |
| HORVU.MOREX.r2.1HG0010820.1 | 5.274 | 5.319 | 2.090 | 2.545 | 0.011 | 0.765 |
| HORVU.MOREX.r2.2HG0090920.1 | 59.219 | 2.470 | 0.971 | 2.543 | 0.011 | 0.766 |
| HORVU.MOREX.r2.2HG0131160.1 | 308.101 | 1.843 | 0.725 | 2.541 | 0.011 | 0.766 |
| HORVU.MOREX.r2.1HG0064720.1 | 16.492 | 3.032 | 1.193 | 2.541 | 0.011 | 0.766 |
| HORVU.MOREX.r2.4HG0344960.1 | 271.701 | 2.353 | 0.928 | 2.537 | 0.011 | 0.771 |
| HORVU.MOREX.r2.7HG0599230.1 | 154.921 | 2.030 | 0.801 | 2.536 | 0.011 | 0.771 |
| HORVU.MOREX.r2.7HG0545660.1 | 160.731 | 2.574 | 1.016 | 2.534 | 0.011 | 0.771 |
| HORVU.MOREX.r2.6HG0511700.1 | 337.503 | 2.170 | 0.857 | 2.533 | 0.011 | 0.771 |
| HORVU.MOREX.r2.7HG0542390.1 | 19.271 | 2.556 | 1.011 | 2.528 | 0.011 | 0.777 |
| HORVU.MOREX.r2.5HG0364350.1 | 1929.454 | 2.237 | 0.886 | 2.526 | 0.012 | 0.778 |
| HORVU.MOREX.r2.3HG0214750.1 | 98.636 | 1.644 | 0.651 | 2.524 | 0.012 | 0.780 |
| HORVU.MOREX.r2.3HG0244560.1 | 91.826 | 2.987 | 1.185 | 2.522 | 0.012 | 0.781 |
| HORVU.MOREX.r2.UnG0628620.1 | 987.485 | 2.658 | 1.056 | 2.517 | 0.012 | 0.784 |
| HORVU.MOREX.r2.2HG0087900.1 | 1045.283 | 2.323 | 0.923 | 2.516 | 0.012 | 0.784 |
| HORVU.MOREX.r2.5HG0349820.1 | 2534.616 | 1.294 | 0.515 | 2.512 | 0.012 | 0.789 |
| HORVU.MOREX.r2.5HG0353860.1 | 19.835 | 3.415 | 1.361 | 2.510 | 0.012 | 0.791 |
| HORVU.MOREX.r2.2HG0091840.1 | 41.108 | 4.534 | 1.807 | 2.509 | 0.012 | 0.791 |
| HORVU.MOREX.r2.1HG0040660.1 | 344.503 | 2.349 | 0.938 | 2.504 | 0.012 | 0.801 |
| HORVU.MOREX.r2.5HG0416570.1 | 101.982 | 1.494 | 0.599 | 2.494 | 0.013 | 0.817 |
| HORVU.MOREX.r2.6HG0450240.1 | 322.088 | 1.615 | 0.647 | 2.494 | 0.013 | 0.817 |
| HORVU.MOREX.r2.7HG0534210.1 | 304.400 | 1.859 | 0.746 | 2.492 | 0.013 | 0.820 |
| HORVU.MOREX.r2.2HG0132080.1 | 192.146 | 2.068 | 0.831 | 2.490 | 0.013 | 0.820 |
| HORVU.MOREX.r2.1HG0044080.1 | 6.642 | 5.166 | 2.076 | 2.488 | 0.013 | 0.821 |
| HORVU.MOREX.r2.3HG0222170.1 | 247.131 | 2.111 | 0.850 | 2.484 | 0.013 | 0.827 |
| HORVU.MOREX.r2.6HG0508620.1 | 320.218 | 1.932 | 0.781 | 2.475 | 0.013 | 0.843 |
| HORVU.MOREX.r2.7HG0618620.1 | 330.532 | 2.504 | 1.016 | 2.465 | 0.014 | 0.849 |
| HORVU.MOREX.r2.3HG0195710.1 | 277.694 | 2.322 | 0.942 | 2.464 | 0.014 | 0.849 |
| HORVU.MOREX.r2.1HG0038140.1 | 196.417 | 2.234 | 0.907 | 2.464 | 0.014 | 0.849 |
| HORVU.MOREX.r2.3HG0185420.1 | 407.694 | 2.366 | 0.960 | 2.463 | 0.014 | 0.849 |
| HORVU.MOREX.r2.2HG0162840.1 | 19.344 | 3.186 | 1.294 | 2.463 | 0.014 | 0.849 |
| HORVU.MOREX.r2.3HG0250370.1 | 6.757 | 4.946 | 2.014 | 2.456 | 0.014 | 0.863 |
| HORVU.MOREX.r2.2HG0177000.1 | 195.003 | 2.623 | 1.069 | 2.454 | 0.014 | 0.863 |
| HORVU.MOREX.r2.6HG0451270.1 | 2893.347 | 2.385 | 0.974 | 2.449 | 0.014 | 0.866 |
| HORVU.MOREX.r2.2HG0166390.1 | 16.051 | 2.932 | 1.198 | 2.448 | 0.014 | 0.866 |
| HORVU.MOREX.r2.3HG0183280.1 | 173.922 | 2.088 | 0.853 | 2.447 | 0.014 | 0.866 |
| HORVU.MOREX.r2.3HG0269080.1 | 130.198 | 2.646 | 1.081 | 2.447 | 0.014 | 0.866 |
| HORVU.MOREX.r2.5HG0421990.1 | 19.171 | 2.925 | 1.197 | 2.444 | 0.015 | 0.868 |
| HORVU.MOREX.r2.2HG0176310.1 | 117.474 | 1.895 | 0.776 | 2.443 | 0.015 | 0.869 |
| HORVU.MOREX.r2.3HG0187780.1 | 543.607 | 1.788 | 0.732 | 2.442 | 0.015 | 0.869 |
| HORVU.MOREX.r2.7HG0529390.1 | 171.473 | 2.411 | 0.989 | 2.439 | 0.015 | 0.873 |
| HORVU.MOREX.r2.1HG0057680.1 | 70.189 | 1.828 | 0.750 | 2.438 | 0.015 | 0.873 |
| HORVU.MOREX.r2.3HG0183290.1 | 40.671 | 2.593 | 1.064 | 2.437 | 0.015 | 0.873 |
| HORVU.MOREX.r2.7HG0537810.1 | 63.337 | 1.986 | 0.817 | 2.430 | 0.015 | 0.883 |
| HORVU.MOREX.r2.2HG0178030.1 | 425.612 | 1.731 | 0.712 | 2.429 | 0.015 | 0.883 |
| HORVU.MOREX.r2.UnG0627300.1 | 2370.339 | 2.113 | 0.870 | 2.428 | 0.015 | 0.884 |
| HORVU.MOREX.r2.5HG0403460.1 | 12.129 | 3.917 | 1.614 | 2.427 | 0.015 | 0.885 |
| HORVU.MOREX.r2.2HG0160620.1 | 119.716 | 1.575 | 0.650 | 2.424 | 0.015 | 0.889 |

| | | | | | | |
|---|---|---|---|---|---|---|
| HORVU.MOREX.r2.3HG0239750.1 | 5266.325 | 2.380 | 0.983 | 2.422 | 0.015 | 0.889 |
| HORVU.MOREX.r2.7HG0593370.1 | 47.316 | 2.181 | 0.901 | 2.421 | 0.015 | 0.889 |
| HORVU.MOREX.r2.4HG0297540.1 | 718.121 | 1.526 | 0.630 | 2.421 | 0.015 | 0.889 |
| HORVU.MOREX.r2.1HG0067870.1 | 43.061 | 2.897 | 1.198 | 2.419 | 0.016 | 0.889 |
| HORVU.MOREX.r2.1HG0010770.1 | 204.150 | 2.396 | 0.991 | 2.418 | 0.016 | 0.889 |
| HORVU.MOREX.r2.3HG0260840.1 | 294.257 | 2.373 | 0.982 | 2.415 | 0.016 | 0.889 |
| HORVU.MOREX.r2.2HG0080670.1 | 9.939 | 3.887 | 1.610 | 2.414 | 0.016 | 0.889 |
| HORVU.MOREX.r2.4HG0326520.1 | 377.386 | 2.048 | 0.849 | 2.412 | 0.016 | 0.889 |
| HORVU.MOREX.r2.2HG0169700.1 | 481.058 | 2.320 | 0.962 | 2.412 | 0.016 | 0.889 |
| HORVU.MOREX.r2.7HG0619610.1 | 17.690 | 2.838 | 1.177 | 2.410 | 0.016 | 0.889 |
| HORVU.MOREX.r2.4HG0328050.1 | 12.743 | 4.326 | 1.797 | 2.408 | 0.016 | 0.892 |
| HORVU.MOREX.r2.7HG0538290.1 | 908.523 | 2.173 | 0.904 | 2.405 | 0.016 | 0.894 |
| HORVU.MOREX.r2.7HG0619780.1 | 36.620 | 1.924 | 0.801 | 2.402 | 0.016 | 0.900 |
| HORVU.MOREX.r2.1HG0050960.1 | 39.358 | 2.501 | 1.042 | 2.400 | 0.016 | 0.902 |
| HORVU.MOREX.r2.7HG0554570.1 | 33.335 | 2.216 | 0.924 | 2.399 | 0.016 | 0.902 |
| HORVU.MOREX.r2.3HG0231980.1 | 2593.949 | 1.815 | 0.757 | 2.398 | 0.016 | 0.902 |
| HORVU.MOREX.r2.4HG0281600.1 | 21.641 | 4.350 | 1.815 | 2.397 | 0.017 | 0.902 |
| HORVU.MOREX.r2.3HG0252970.1 | 110.715 | 2.433 | 1.016 | 2.394 | 0.017 | 0.906 |
| HORVU.MOREX.r2.2HG0166750.1 | 18.531 | 2.923 | 1.222 | 2.393 | 0.017 | 0.906 |
| HORVU.MOREX.r2.1HG0059450.1 | 13.901 | 3.280 | 1.372 | 2.391 | 0.017 | 0.907 |
| HORVU.MOREX.r2.4HG0299180.1 | 320.057 | 4.439 | 1.859 | 2.388 | 0.017 | 0.911 |
| HORVU.MOREX.r2.5HG0438640.1 | 96.812 | 1.826 | 0.767 | 2.380 | 0.017 | 0.928 |
| HORVU.MOREX.r2.1HG0074860.1 | 19.672 | 3.157 | 1.328 | 2.378 | 0.017 | 0.928 |
| HORVU.MOREX.r2.2HG0145800.1 | 118.925 | 1.619 | 0.681 | 2.377 | 0.017 | 0.928 |
| HORVU.MOREX.r2.6HG0516400.1 | 1261.840 | 2.016 | 0.848 | 2.376 | 0.017 | 0.928 |
| HORVU.MOREX.r2.5HG0379710.1 | 7.770 | 4.277 | 1.807 | 2.367 | 0.018 | 0.943 |
| HORVU.MOREX.r2.5HG0412750.1 | 17.126 | 2.379 | 1.006 | 2.366 | 0.018 | 0.943 |
| HORVU.MOREX.r2.5HG0442410.1 | 13.219 | 3.126 | 1.321 | 2.366 | 0.018 | 0.943 |
| HORVU.MOREX.r2.6HG0524710.1 | 670.388 | 1.312 | 0.559 | 2.349 | 0.019 | 0.979 |
| HORVU.MOREX.r2.3HG0209060.1 | 500.268 | 1.599 | 0.681 | 2.347 | 0.019 | 0.982 |
| HORVU.MOREX.r2.1HG0063800.1 | 27.336 | 2.433 | 1.037 | 2.346 | 0.019 | 0.983 |
| HORVU.MOREX.r2.4HG0341040.1 | 50.452 | 2.545 | 1.086 | 2.344 | 0.019 | 0.984 |
| HORVU.MOREX.r2.2HG0127510.1 | 56.308 | 2.466 | 1.053 | 2.341 | 0.019 | 0.987 |
| HORVU.MOREX.r2.7HG0607550.1 | 257.135 | 1.948 | 0.832 | 2.340 | 0.019 | 0.988 |
| HORVU.MOREX.r2.5HG0404510.1 | 11234.259 | 2.247 | 0.961 | 2.338 | 0.019 | 0.988 |
| HORVU.MOREX.r2.1HG0060090.1 | 265.295 | 1.690 | 0.723 | 2.338 | 0.019 | 0.988 |
| HORVU.MOREX.r2.3HG0245260.1 | 305.863 | 2.051 | 0.878 | 2.337 | 0.019 | 0.988 |
| HORVU.MOREX.r2.5HG0419050.1 | 59.997 | 1.883 | 0.806 | 2.335 | 0.020 | 0.993 |
| HORVU.MOREX.r2.5HG0444860.1 | 13790.821 | 2.166 | 0.929 | 2.331 | 0.020 | 0.995 |
| HORVU.MOREX.r2.7HG0608800.1 | 703.046 | 1.249 | 0.536 | 2.331 | 0.020 | 0.995 |
| HORVU.MOREX.r2.3HG0200100.1 | 99.370 | 2.650 | 1.138 | 2.329 | 0.020 | 0.995 |
| HORVU.MOREX.r2.7HG0561520.1 | 590.808 | 1.443 | 0.620 | 2.326 | 0.020 | 0.996 |
| HORVU.MOREX.r2.2HG0173530.1 | 41.277 | 2.349 | 1.010 | 2.326 | 0.020 | 0.996 |
| HORVU.MOREX.r2.4HG0339440.1 | 325.605 | 1.264 | 0.543 | 2.326 | 0.020 | 0.996 |
| HORVU.MOREX.r2.7HG0574240.1 | 69.395 | 1.971 | 0.848 | 2.325 | 0.020 | 0.996 |
| HORVU.MOREX.r2.1HG0002140.1 | 54.744 | 1.732 | 0.858 | 2.019 | 0.043 | 1.000 |
| HORVU.MOREX.r2.1HG0002090.1 | 27.453 | 2.118 | 1.340 | 1.581 | 0.114 | 1.000 |
| HORVU.MOREX.r2.1HG0002870.1 | 6.731 | 1.866 | 1.638 | 1.139 | 0.255 | 1.000 |
| HORVU.MOREX.r2.1HG0001590.1 | 15.271 | 1.233 | 1.152 | 1.070 | 0.285 | 1.000 |
| HORVU.MOREX.r2.1HG0002560.1 | 6277.594 | 0.726 | 0.703 | 1.033 | 0.302 | 1.000 |
| HORVU.MOREX.r2.1HG0001870.1 | 449.835 | 0.520 | 0.545 | 0.953 | 0.340 | 1.000 |
| HORVU.MOREX.r2.1HG0002010.1 | 39.620 | 0.846 | 0.895 | 0.945 | 0.345 | 1.000 |
| HORVU.MOREX.r2.1HG0000900.1 | 5.197 | 2.088 | 2.246 | 0.930 | 0.352 | 1.000 |

| | | | | | | |
|---|---|---|---|---|---|---|
| HORVU.MOREX.r2.1HG0002210.1 | 5.961 | 1.662 | 1.801 | 0.923 | 0.356 | 1.000 |
| HORVU.MOREX.r2.1HG0000740.1 | 841.537 | 0.469 | 0.676 | 0.694 | 0.488 | 1.000 |
| HORVU.MOREX.r2.1HG0003010.1 | 104.738 | 0.404 | 0.601 | 0.673 | 0.501 | 1.000 |
| HORVU.MOREX.r2.1HG0000290.1 | 33.297 | 0.481 | 0.774 | 0.621 | 0.534 | 1.000 |
| HORVU.MOREX.r2.1HG0000030.1 | 1778.666 | 0.360 | 0.591 | 0.610 | 0.542 | 1.000 |
| HORVU.MOREX.r2.1HG0001430.1 | 147.055 | 0.357 | 0.664 | 0.538 | 0.591 | 1.000 |
| HORVU.MOREX.r2.1HG0002410.1 | 116.515 | 0.342 | 0.658 | 0.519 | 0.603 | 1.000 |
| HORVU.MOREX.r2.1HG0002750.1 | 353.690 | 0.264 | 0.529 | 0.499 | 0.618 | 1.000 |
| HORVU.MOREX.r2.1HG0001030.1 | 187.202 | 0.297 | 0.675 | 0.440 | 0.660 | 1.000 |
| HORVU.MOREX.r2.1HG0003070.1 | 142.714 | 0.274 | 0.647 | 0.423 | 0.672 | 1.000 |
| HORVU.MOREX.r2.1HG0002200.1 | 20.589 | 0.352 | 0.932 | 0.378 | 0.706 | 1.000 |
| HORVU.MOREX.r2.1HG0003080.1 | 893.778 | 0.179 | 0.521 | 0.343 | 0.731 | 1.000 |
| HORVU.MOREX.r2.1HG0000050.1 | 190.290 | 0.300 | 0.876 | 0.342 | 0.732 | 1.000 |
| HORVU.MOREX.r2.1HG0000960.1 | 130.330 | 0.201 | 0.614 | 0.328 | 0.743 | 1.000 |
| HORVU.MOREX.r2.1HG0000590.1 | 687.025 | 0.270 | 0.845 | 0.320 | 0.749 | 1.000 |
| HORVU.MOREX.r2.1HG0000060.1 | 709.803 | 0.180 | 0.658 | 0.274 | 0.784 | 1.000 |
| HORVU.MOREX.r2.1HG0002790.1 | 7.946 | 0.306 | 1.396 | 0.219 | 0.826 | 1.000 |
| HORVU.MOREX.r2.1HG0002330.1 | 235.637 | 0.111 | 0.552 | 0.201 | 0.841 | 1.000 |
| HORVU.MOREX.r2.1HG0002840.1 | 865.277 | 0.111 | 0.614 | 0.180 | 0.857 | 1.000 |
| HORVU.MOREX.r2.1HG0000120.1 | 123.273 | 0.074 | 0.578 | 0.128 | 0.898 | 1.000 |
| HORVU.MOREX.r2.1HG0001690.1 | 224.315 | 0.052 | 0.557 | 0.093 | 0.926 | 1.000 |
| HORVU.MOREX.r2.1HG0002680.1 | 4.873 | 0.206 | 2.261 | 0.091 | 0.927 | 1.000 |
| HORVU.MOREX.r2.1HG0001730.1 | 32.948 | 0.070 | 0.827 | 0.085 | 0.932 | 1.000 |
| HORVU.MOREX.r2.1HG0000700.1 | 179.028 | 0.045 | 0.529 | 0.085 | 0.933 | 1.000 |
| HORVU.MOREX.r2.1HG0001540.1 | 4960.580 | 0.039 | 0.510 | 0.077 | 0.938 | 1.000 |
| HORVU.MOREX.r2.1HG0000320.1 | 5.724 | 0.099 | 1.784 | 0.055 | 0.956 | 1.000 |
| HORVU.MOREX.r2.1HG0002610.1 | 4.739 | 0.109 | 2.219 | 0.049 | 0.961 | 1.000 |
| HORVU.MOREX.r2.1HG0000670.1 | 93.072 | 0.020 | 0.612 | 0.033 | 0.974 | 1.000 |
| HORVU.MOREX.r2.1HG0000720.1 | 172.694 | 0.025 | 0.827 | 0.031 | 0.975 | 1.000 |

Table S8

| Downregulated genes at T42 | baseMean | log2FC | lfcSE | stat | pvalue | padj |
|---|---|---|---|---|---|---|
| HORVU.MOREX.r2.6HG0523360.1 | 3199.432 | -8.821 | 1.023 | -8.626 | 0.000 | 0.000 |
| HORVU.MOREX.r2.2HG0155090.1 | 729.301 | -5.455 | 0.671 | -8.126 | 0.000 | 0.000 |
| HORVU.MOREX.r2.7HG0554640.1 | 88.693 | -7.050 | 0.994 | -7.091 | 0.000 | 0.000 |
| HORVU.MOREX.r2.1HG0064430.1 | 9968.387 | -4.839 | 0.684 | -7.071 | 0.000 | 0.000 |
| HORVU.MOREX.r2.7HG0554650.1 | 1170.820 | -5.433 | 0.969 | -5.609 | 0.000 | 0.000 |
| HORVU.MOREX.r2.7HG0541520.1 | 75.775 | -24.575 | 4.470 | -5.498 | 0.000 | 0.000 |
| HORVU.MOREX.r2.1HG0057310.1 | 356.737 | -5.436 | 1.053 | -5.161 | 0.000 | 0.001 |
| HORVU.MOREX.r2.2HG0086010.1 | 15169.918 | -3.883 | 0.757 | -5.129 | 0.000 | 0.001 |
| HORVU.MOREX.r2.3HG0228770.1 | 148.799 | -3.423 | 0.688 | -4.978 | 0.000 | 0.001 |
| HORVU.MOREX.r2.2HG0086020.1 | 14575.148 | -3.733 | 0.787 | -4.744 | 0.000 | 0.004 |
| HORVU.MOREX.r2.1HG0073480.1 | 3574.748 | -4.324 | 0.950 | -4.551 | 0.000 | 0.007 |
| HORVU.MOREX.r2.1HG0073470.1 | 522.396 | -4.234 | 1.000 | -4.233 | 0.000 | 0.019 |
| HORVU.MOREX.r2.5HG0390230.1 | 89.258 | -4.680 | 1.158 | -4.041 | 0.000 | 0.035 |
| HORVU.MOREX.r2.1HG0073500.1 | 9384.527 | -3.091 | 0.768 | -4.026 | 0.000 | 0.036 |
| HORVU.MOREX.r2.3HG0247980.1 | 87.904 | -4.006 | 1.021 | -3.922 | 0.000 | 0.051 |
| HORVU.MOREX.r2.1HG0044650.1 | 109.119 | -3.185 | 0.815 | -3.906 | 0.000 | 0.052 |
| HORVU.MOREX.r2.6HG0517650.1 | 101.994 | -3.984 | 1.043 | -3.819 | 0.000 | 0.065 |
| HORVU.MOREX.r2.7HG0542080.1 | 41.292 | -3.138 | 0.844 | -3.719 | 0.000 | 0.085 |
| HORVU.MOREX.r2.7HG0565820.1 | 38.577 | -4.371 | 1.183 | -3.695 | 0.000 | 0.092 |
| HORVU.MOREX.r2.UnG0630910.1 | 20.148 | -5.401 | 1.467 | -3.681 | 0.000 | 0.093 |
| HORVU.MOREX.r2.7HG0533270.1 | 91.700 | -2.573 | 0.709 | -3.631 | 0.000 | 0.105 |
| HORVU.MOREX.r2.4HG0279880.1 | 19.726 | -4.733 | 1.331 | -3.555 | 0.000 | 0.134 |
| HORVU.MOREX.r2.3HG0254990.1 | 513.135 | -2.553 | 0.720 | -3.546 | 0.000 | 0.134 |
| HORVU.MOREX.r2.1HG0029020.1 | 23366.702 | -2.309 | 0.653 | -3.537 | 0.000 | 0.134 |
| HORVU.MOREX.r2.6HG0456290.1 | 504.750 | -2.556 | 0.723 | -3.534 | 0.000 | 0.134 |
| HORVU.MOREX.r2.6HG0460180.1 | 101.089 | -9.465 | 2.759 | -3.431 | 0.001 | 0.177 |
| HORVU.MOREX.r2.1HG0039200.1 | 142.848 | -2.396 | 0.700 | -3.422 | 0.001 | 0.177 |
| HORVU.MOREX.r2.1HG0037430.1 | 4157.696 | -2.138 | 0.627 | -3.412 | 0.001 | 0.178 |
| HORVU.MOREX.r2.3HG0255670.1 | 130.438 | -3.177 | 0.933 | -3.407 | 0.001 | 0.178 |
| HORVU.MOREX.r2.3HG0189830.1 | 70.490 | -3.506 | 1.034 | -3.391 | 0.001 | 0.184 |
| HORVU.MOREX.r2.6HG0507730.1 | 537.227 | -2.389 | 0.716 | -3.338 | 0.001 | 0.197 |
| HORVU.MOREX.r2.4HG0316490.1 | 14.050 | -4.912 | 1.474 | -3.333 | 0.001 | 0.199 |
| HORVU.MOREX.r2.7HG0544880.1 | 12.890 | -5.071 | 1.528 | -3.319 | 0.001 | 0.200 |
| HORVU.MOREX.r2.4HG0293270.1 | 97.215 | -2.628 | 0.793 | -3.314 | 0.001 | 0.200 |
| HORVU.MOREX.r2.2HG0086030.1 | 37458.759 | -2.586 | 0.783 | -3.303 | 0.001 | 0.206 |
| HORVU.MOREX.r2.6HG0461000.1 | 62.190 | -3.800 | 1.162 | -3.272 | 0.001 | 0.218 |
| HORVU.MOREX.r2.4HG0340230.1 | 193.484 | -2.626 | 0.815 | -3.221 | 0.001 | 0.241 |
| HORVU.MOREX.r2.5HG0438530.1 | 8.400 | -5.691 | 1.814 | -3.137 | 0.002 | 0.307 |
| HORVU.MOREX.r2.2HG0147310.1 | 700.406 | -2.320 | 0.746 | -3.109 | 0.002 | 0.328 |
| HORVU.MOREX.r2.1HG0047170.1 | 2941.363 | -2.544 | 0.828 | -3.072 | 0.002 | 0.349 |
| HORVU.MOREX.r2.7HG0589710.1 | 253.260 | -1.964 | 0.644 | -3.051 | 0.002 | 0.354 |
| HORVU.MOREX.r2.2HG0101220.1 | 43.063 | -3.750 | 1.244 | -3.014 | 0.003 | 0.376 |
| HORVU.MOREX.r2.4HG0344750.1 | 65.111 | -2.740 | 0.910 | -3.011 | 0.003 | 0.376 |
| HORVU.MOREX.r2.5HG0434090.1 | 9584.924 | -5.010 | 1.668 | -3.004 | 0.003 | 0.376 |
| HORVU.MOREX.r2.4HG0330890.1 | 626.909 | -2.634 | 0.881 | -2.990 | 0.003 | 0.389 |
| HORVU.MOREX.r2.5HG0407120.1 | 11.200 | -4.593 | 1.542 | -2.978 | 0.003 | 0.393 |
| HORVU.MOREX.r2.2HG0146770.1 | 11.168 | -5.343 | 1.828 | -2.923 | 0.003 | 0.431 |
| HORVU.MOREX.r2.3HG0259610.1 | 3600.665 | -2.138 | 0.732 | -2.919 | 0.004 | 0.433 |
| HORVU.MOREX.r2.4HG0302780.1 | 8.399 | -4.384 | 1.509 | -2.905 | 0.004 | 0.443 |
| HORVU.MOREX.r2.5HG0392520.1 | 2415.429 | -1.899 | 0.655 | -2.897 | 0.004 | 0.446 |
| HORVU.MOREX.r2.2HG0142880.1 | 17653.326 | -2.044 | 0.707 | -2.892 | 0.004 | 0.446 |
| HORVU.MOREX.r2.5HG0434050.1 | 15980.128 | -2.226 | 0.775 | -2.873 | 0.004 | 0.458 |

| | | | | | | |
|---|---|---|---|---|---|---|
| HORVU.MOREX.r2.4HG0332690.1 | 10.294 | -4.591 | 1.601 | -2.867 | 0.004 | 0.464 |
| HORVU.MOREX.r2.5HG0418360.1 | 40.903 | -3.310 | 1.168 | -2.833 | 0.005 | 0.501 |
| HORVU.MOREX.r2.UnG0632950.1 | 12424.714 | -2.299 | 0.820 | -2.802 | 0.005 | 0.527 |
| HORVU.MOREX.r2.2HG0113870.1 | 147.447 | -2.650 | 0.958 | -2.768 | 0.006 | 0.550 |
| HORVU.MOREX.r2.2HG0111260.1 | 37662.535 | -1.884 | 0.686 | -2.746 | 0.006 | 0.569 |
| HORVU.MOREX.r2.5HG0426190.1 | 2432.047 | -2.003 | 0.730 | -2.742 | 0.006 | 0.573 |
| HORVU.MOREX.r2.6HG0448850.1 | 6.257 | -5.401 | 1.984 | -2.722 | 0.006 | 0.588 |
| HORVU.MOREX.r2.2HG0086050.1 | 26072.389 | -2.193 | 0.806 | -2.721 | 0.007 | 0.588 |
| HORVU.MOREX.r2.4HG0325830.1 | 30.274 | -2.080 | 0.771 | -2.699 | 0.007 | 0.615 |
| HORVU.MOREX.r2.2HG0081740.1 | 8488.327 | -6.040 | 2.249 | -2.685 | 0.007 | 0.630 |
| HORVU.MOREX.r2.3HG0222900.1 | 48.642 | -2.778 | 1.038 | -2.676 | 0.007 | 0.637 |
| HORVU.MOREX.r2.2HG0096630.1 | 74291.740 | -2.102 | 0.786 | -2.675 | 0.007 | 0.637 |
| HORVU.MOREX.r2.1HG0055260.1 | 25098.951 | -1.836 | 0.687 | -2.673 | 0.008 | 0.637 |
| HORVU.MOREX.r2.2HG0128210.1 | 4.565 | -5.435 | 2.035 | -2.671 | 0.008 | 0.637 |
| HORVU.MOREX.r2.7HG0604280.1 | 5.171 | -5.295 | 1.993 | -2.656 | 0.008 | 0.654 |
| HORVU.MOREX.r2.6HG0456520.1 | 612.711 | -1.786 | 0.677 | -2.639 | 0.008 | 0.677 |
| HORVU.MOREX.r2.1HG0016640.1 | 6.750 | -3.935 | 1.499 | -2.626 | 0.009 | 0.688 |
| HORVU.MOREX.r2.5HG0385770.1 | 378.932 | -2.091 | 0.797 | -2.625 | 0.009 | 0.688 |
| HORVU.MOREX.r2.5HG0446960.1 | 15.653 | -4.343 | 1.662 | -2.613 | 0.009 | 0.702 |
| HORVU.MOREX.r2.2HG0127750.1 | 1618.351 | -1.540 | 0.590 | -2.610 | 0.009 | 0.705 |
| HORVU.MOREX.r2.UnG0628370.1 | 546.342 | -1.694 | 0.650 | -2.608 | 0.009 | 0.707 |
| HORVU.MOREX.r2.2HG0133590.1 | 7.803 | -4.486 | 1.724 | -2.603 | 0.009 | 0.711 |
| HORVU.MOREX.r2.1HG0074060.1 | 158.732 | -2.197 | 0.847 | -2.593 | 0.010 | 0.716 |
| HORVU.MOREX.r2.1HG0049650.1 | 1426.341 | -1.376 | 0.532 | -2.587 | 0.010 | 0.724 |
| HORVU.MOREX.r2.3HG0237280.1 | 165.993 | -1.532 | 0.596 | -2.570 | 0.010 | 0.740 |
| HORVU.MOREX.r2.5HG0403890.1 | 9.731 | -3.521 | 1.375 | -2.560 | 0.010 | 0.751 |
| HORVU.MOREX.r2.5HG0412070.1 | 258.925 | -2.195 | 0.858 | -2.557 | 0.011 | 0.751 |
| HORVU.MOREX.r2.1HG0036930.1 | 14.139 | -3.797 | 1.487 | -2.553 | 0.011 | 0.753 |
| HORVU.MOREX.r2.1HG0072930.1 | 295.240 | -2.166 | 0.855 | -2.535 | 0.011 | 0.771 |
| HORVU.MOREX.r2.4HG0342410.1 | 91.147 | -2.390 | 0.945 | -2.530 | 0.011 | 0.774 |
| HORVU.MOREX.r2.5HG0418240.1 | 18932.355 | -1.524 | 0.604 | -2.522 | 0.012 | 0.781 |
| HORVU.MOREX.r2.3HG0214090.1 | 22193.575 | -1.779 | 0.707 | -2.518 | 0.012 | 0.784 |
| HORVU.MOREX.r2.4HG0302790.1 | 9.623 | -3.561 | 1.416 | -2.516 | 0.012 | 0.784 |
| HORVU.MOREX.r2.2HG0109810.1 | 267.894 | -1.840 | 0.739 | -2.491 | 0.013 | 0.820 |
| HORVU.MOREX.r2.1HG0052860.1 | 127.954 | -2.170 | 0.874 | -2.484 | 0.013 | 0.827 |
| HORVU.MOREX.r2.5HG0435270.1 | 13.866 | -3.281 | 1.325 | -2.476 | 0.013 | 0.843 |
| HORVU.MOREX.r2.1HG0035960.1 | 4978.580 | -1.630 | 0.661 | -2.467 | 0.014 | 0.849 |
| HORVU.MOREX.r2.3HG0239800.1 | 573.219 | -2.097 | 0.851 | -2.466 | 0.014 | 0.849 |
| HORVU.MOREX.r2.1HG0044610.1 | 341.048 | -5.659 | 2.296 | -2.465 | 0.014 | 0.849 |
| HORVU.MOREX.r2.2HG0160340.1 | 3257.600 | -1.866 | 0.760 | -2.455 | 0.014 | 0.863 |
| HORVU.MOREX.r2.4HG0298730.1 | 78.357 | -1.928 | 0.786 | -2.453 | 0.014 | 0.863 |
| HORVU.MOREX.r2.2HG0176640.1 | 287.493 | -1.858 | 0.758 | -2.452 | 0.014 | 0.863 |
| HORVU.MOREX.r2.5HG0407340.1 | 1398.927 | -1.898 | 0.776 | -2.444 | 0.015 | 0.868 |
| HORVU.MOREX.r2.4HG0336620.1 | 21.471 | -3.537 | 1.453 | -2.435 | 0.015 | 0.876 |
| HORVU.MOREX.r2.4HG0335810.1 | 205.739 | -1.475 | 0.610 | -2.417 | 0.016 | 0.889 |
| HORVU.MOREX.r2.5HG0384080.1 | 309.483 | -1.695 | 0.703 | -2.410 | 0.016 | 0.889 |
| HORVU.MOREX.r2.7HG0553370.1 | 99.154 | -1.867 | 0.775 | -2.410 | 0.016 | 0.889 |
| HORVU.MOREX.r2.3HG0192770.1 | 355.975 | -1.441 | 0.598 | -2.410 | 0.016 | 0.889 |
| HORVU.MOREX.r2.2HG0137850.1 | 5311.704 | -1.438 | 0.597 | -2.407 | 0.016 | 0.892 |
| HORVU.MOREX.r2.1HG0009050.1 | 3931.879 | -1.613 | 0.674 | -2.393 | 0.017 | 0.906 |
| HORVU.MOREX.r2.4HG0286480.1 | 1259.704 | -1.464 | 0.613 | -2.389 | 0.017 | 0.909 |
| HORVU.MOREX.r2.3HG0248650.1 | 10.518 | -2.905 | 1.223 | -2.376 | 0.017 | 0.928 |
| HORVU.MOREX.r2.UnG0635730.1 | 25017.352 | -1.908 | 0.803 | -2.375 | 0.018 | 0.928 |

| | | | | | | |
|---|---|---|---|---|---|---|
| HORVU.MOREX.r2.4HG0321470.1 | 348.677 | -1.458 | 0.615 | -2.370 | 0.018 | 0.940 |
| HORVU.MOREX.r2.5HG0405770.1 | 7871.735 | -1.448 | 0.615 | -2.353 | 0.019 | 0.973 |
| HORVU.MOREX.r2.2HG0127820.1 | 2009.311 | -1.881 | 0.800 | -2.350 | 0.019 | 0.978 |
| HORVU.MOREX.r2.6HG0459300.1 | 601.816 | -1.619 | 0.691 | -2.344 | 0.019 | 0.984 |
| HORVU.MOREX.r2.3HG0270810.1 | 9511.133 | -1.475 | 0.633 | -2.330 | 0.020 | 0.995 |
| HORVU.MOREX.r2.7HG0567810.1 | 79.727 | -1.733 | 0.744 | -2.330 | 0.020 | 0.995 |
| HORVU.MOREX.r2.1HG0000870.1 | 437.687 | -1.632 | 0.742 | -2.200 | 0.028 | 1.000 |
| HORVU.MOREX.r2.1HG0001760.1 | 37.889 | -6.139 | 3.294 | -1.864 | 0.062 | 1.000 |
| HORVU.MOREX.r2.1HG0002390.1 | 5.742 | -2.988 | 1.734 | -1.723 | 0.085 | 1.000 |
| HORVU.MOREX.r2.1HG0000470.1 | 1009.488 | -1.007 | 0.641 | -1.571 | 0.116 | 1.000 |
| HORVU.MOREX.r2.1HG0000110.1 | 158.819 | -1.371 | 0.895 | -1.531 | 0.126 | 1.000 |
| HORVU.MOREX.r2.1HG0000340.1 | 22.775 | -1.344 | 0.970 | -1.385 | 0.166 | 1.000 |
| HORVU.MOREX.r2.1HG0000450.1 | 39.328 | -0.960 | 0.713 | -1.346 | 0.178 | 1.000 |
| HORVU.MOREX.r2.1HG0000490.1 | 17.708 | -1.761 | 1.363 | -1.292 | 0.196 | 1.000 |
| HORVU.MOREX.r2.1HG0001570.1 | 5.383 | -2.159 | 1.675 | -1.289 | 0.198 | 1.000 |
| HORVU.MOREX.r2.1HG0000930.1 | 7.657 | -2.482 | 1.989 | -1.248 | 0.212 | 1.000 |
| HORVU.MOREX.r2.1HG0000080.1 | 111.377 | -0.821 | 0.691 | -1.188 | 0.235 | 1.000 |
| HORVU.MOREX.r2.1HG0000580.1 | 43.013 | -0.841 | 0.727 | -1.158 | 0.247 | 1.000 |
| HORVU.MOREX.r2.1HG0000730.1 | 744.723 | -0.541 | 0.507 | -1.067 | 0.286 | 1.000 |
| HORVU.MOREX.r2.1HG0003050.1 | 206.378 | -0.559 | 0.562 | -0.996 | 0.319 | 1.000 |
| HORVU.MOREX.r2.1HG0002170.1 | 1889.994 | -0.574 | 0.621 | -0.925 | 0.355 | 1.000 |
| HORVU.MOREX.r2.1HG0001580.1 | 257.578 | -0.494 | 0.536 | -0.923 | 0.356 | 1.000 |
| HORVU.MOREX.r2.1HG0002350.1 | 51.959 | -0.615 | 0.747 | -0.823 | 0.410 | 1.000 |
| HORVU.MOREX.r2.1HG0001550.1 | 84.037 | -0.439 | 0.605 | -0.726 | 0.468 | 1.000 |
| HORVU.MOREX.r2.1HG0000440.1 | 155.189 | -0.427 | 0.606 | -0.705 | 0.481 | 1.000 |
| HORVU.MOREX.r2.1HG0002620.1 | 30.897 | -0.667 | 0.954 | -0.700 | 0.484 | 1.000 |
| HORVU.MOREX.r2.1HG0001710.1 | 512.690 | -0.328 | 0.524 | -0.625 | 0.532 | 1.000 |
| HORVU.MOREX.r2.1HG0001640.1 | 70.137 | -0.517 | 0.847 | -0.610 | 0.542 | 1.000 |
| HORVU.MOREX.r2.1HG0001910.1 | 164.047 | -0.340 | 0.640 | -0.532 | 0.595 | 1.000 |
| HORVU.MOREX.r2.1HG0001410.1 | 372.801 | -0.339 | 0.654 | -0.518 | 0.605 | 1.000 |
| HORVU.MOREX.r2.1HG0001720.1 | 271.693 | -0.260 | 0.504 | -0.515 | 0.606 | 1.000 |
| HORVU.MOREX.r2.1HG0000090.1 | 1436.850 | -0.304 | 0.614 | -0.496 | 0.620 | 1.000 |
| HORVU.MOREX.r2.1HG0001250.1 | 19.507 | -0.343 | 0.923 | -0.372 | 0.710 | 1.000 |
| HORVU.MOREX.r2.1HG0001530.1 | 261.353 | -0.228 | 0.631 | -0.361 | 0.718 | 1.000 |
| HORVU.MOREX.r2.1HG0001920.1 | 103.560 | -0.234 | 0.696 | -0.335 | 0.737 | 1.000 |
| HORVU.MOREX.r2.1HG0000070.1 | 524.120 | -0.163 | 0.520 | -0.313 | 0.754 | 1.000 |
| HORVU.MOREX.r2.1HG0002580.1 | 93.522 | -1.079 | 4.226 | -0.255 | 0.799 | 1.000 |
| HORVU.MOREX.r2.1HG0000040.1 | 542.555 | -0.134 | 0.646 | -0.207 | 0.836 | 1.000 |
| HORVU.MOREX.r2.1HG0002190.1 | 534.923 | -0.109 | 0.545 | -0.200 | 0.841 | 1.000 |
| HORVU.MOREX.r2.1HG0001980.1 | 112.704 | -0.111 | 0.635 | -0.175 | 0.861 | 1.000 |
| HORVU.MOREX.r2.1HG0002180.1 | 124.769 | -0.090 | 0.573 | -0.156 | 0.876 | 1.000 |
| HORVU.MOREX.r2.1HG0002860.1 | 113.175 | -0.063 | 0.604 | -0.104 | 0.917 | 1.000 |
| HORVU.MOREX.r2.1HG0002460.1 | 12.195 | -0.107 | 1.060 | -0.101 | 0.919 | 1.000 |
| HORVU.MOREX.r2.1HG0000840.1 | 6.441 | -0.172 | 1.726 | -0.099 | 0.921 | 1.000 |
| HORVU.MOREX.r2.1HG0000350.1 | 10.472 | -0.105 | 1.122 | -0.094 | 0.925 | 1.000 |
| HORVU.MOREX.r2.1HG0000240.1 | 9.149 | -0.183 | 4.627 | -0.040 | 0.968 | 1.000 |
| HORVU.MOREX.r2.1HG0001020.1 | 121.349 | -0.020 | 0.633 | -0.031 | 0.975 | 1.000 |
| HORVU.MOREX.r2.1HG0000280.1 | 1723.857 | -0.016 | 0.611 | -0.026 | 0.979 | 1.000 |
| HORVU.MOREX.r2.1HG0002020.1 | 5832.087 | -0.008 | 0.631 | -0.012 | 0.990 | 1.000 |

Table S9

Pathways downregulated at T2 vs C2

| Enrichment FDR | nGenes | Pathway Genes | Fold Enrichment | Pathway | Genes |
|---|---|---|---|---|---|
| 0.101 | 1.000 | 31.000 | 189.726 | GO:0009522 photosystem I | HORVU.MOREX.R2.1HG0064430.1 |
| 0.101 | 1.000 | 33.000 | 178.227 | GO:0018298 protein-chromophore linkage | HORVU.MOREX.R2.1HG0064430.1 |
| 0.101 | 1.000 | 62.000 | 94.863 | GO:0009523 photosystem II | HORVU.MOREX.R2.1HG0064430.1 |
| 0.101 | 1.000 | 70.000 | 84.021 | GO:0009535 chloroplast thylakoid membrane | HORVU.MOREX.R2.1HG0064430.1 |
| 0.101 | 1.000 | 71.000 | 82.838 | GO:0055035 plastid thylakoid membrane | HORVU.MOREX.R2.1HG0064430.1 |
| 0.101 | 1.000 | 81.000 | 72.611 | GO:0009534 chloroplast thylakoid | HORVU.MOREX.R2.1HG0064430.1 |
| 0.101 | 1.000 | 82.000 | 71.726 | GO:0031976 plastid thylakoid | HORVU.MOREX.R2.1HG0064430.1 |
| 0.101 | 1.000 | 86.000 | 68.390 | GO:0009521 photosystem | HORVU.MOREX.R2.1HG0064430.1 |
| 0.101 | 1.000 | 99.000 | 59.409 | GO:0019684 photosynthesis light reaction | HORVU.MOREX.R2.1HG0064430.1 |
| 0.101 | 1.000 | 101.000 | 58.233 | GO:0042651 thylakoid membrane | HORVU.MOREX.R2.1HG0064430.1 |

Table S10

Pathways upregulated in T12 vs C12

| Enrichment FDR | nGenes | Pathway Genes | Fold Enrichment | Pathway | Genes |
|---|---|---|---|---|---|
| 0.079 | 1.000 | 3.000 | 466.786 | GO:0008940 nitrate reductase activity | HORVU.MOREX.R2.6HG0513160.1 |
| 0.079 | 1.000 | 7.000 | 200.051 | GO:0042126 nitrate metabolic proc. | HORVU.MOREX.R2.6HG0513160.1 |
| 0.079 | 1.000 | 7.000 | 200.051 | GO:0042128 nitrate assimilation | HORVU.MOREX.R2.6HG0513160.1 |
| 0.079 | 1.000 | 7.000 | 200.051 | GO:2001057 reactive nitrogen species metabolic proc. | HORVU.MOREX.R2.6HG0513160.1 |
| 0.079 | 1.000 | 8.000 | 175.045 | GO:0045168 cell-cell signaling involved in cell fate commitment | HORVU.MOREX.R2.4HG0341920.1 |
| 0.079 | 1.000 | 9.000 | 155.595 | GO:0071941 nitrogen cycle metabolic proc. | HORVU.MOREX.R2.6HG0513160.1 |
| 0.079 | 2.000 | 151.000 | 18.548 | GO:0045333 cellular respiration | HORVU.MOREX.R2.2HG0127650.1<br>HORVU.MOREX.R2.UNG0634070.1 |
| 0.079 | 2.000 | 165.000 | 16.974 | GO:0015980 energy derivation by oxidation of organic compounds | HORVU.MOREX.R2.2HG0127650.1<br>HORVU.MOREX.R2.UNG0634070.1 |
| 0.079 | 3.000 | 494.000 | 8.504 | GO:0048507 meristem development | HORVU.MOREX.R2.2HG0125490.1<br>HORVU.MOREX.R2.3HG0237050.1<br>HORVU.MOREX.R2.5HG0378750.1 |
| 0.079 | 4.000 | 1076.000 | 5.206 | GO:0032502 developmental proc. | HORVU.MOREX.R2.4HG0341920.1<br>HORVU.MOREX.R2.2HG0125490.1<br>HORVU.MOREX.R2.3HG0237050.1<br>HORVU.MOREX.R2.5HG0378750.1 |

Table S11

Pathways downregulated in T12 vs C12

| Enrichment FDR | nGenes | Pathway Genes | Fold Enrichment | Pathway | Genes |
|---|---|---|---|---|---|
| 0.0302 | 1.0000 | 4.0000 | 1225.3125 | GO:0034724 DNA replication-independent | HORVU.MOREX.R2.2HG0115250.1 |
| 0.0302 | 1.0000 | 4.0000 | 1225.3125 | GO:0006336 DNA replication-independent chromatin assembly | HORVU.MOREX.R2.2HG0115250.1 |
| 0.0873 | 1.0000 | 25.0000 | 196.0500 | GO:0060918 auxin transport | HORVU.MOREX.R2.1HG0028240.1 |
| 0.0873 | 1.0000 | 25.0000 | 196.0500 | GO:0009914 hormone transport | HORVU.MOREX.R2.1HG0028240.1 |
| 0.0873 | 1.0000 | 29.0000 | 169.0086 | GO:0004427 inorganic diphosphatase activity | HORVU.MOREX.R2.1HG0004870.1 |
| 0.0874 | 3.0000 | 2119.0000 | 6.9390 | GO:0006796 phosphate-containing compound metabolic proc. | HORVU.MOREX.R2.1HG0001760.1 HORVU.MOREX.R2.1HG0002580.1 HORVU.MOREX.R2.1HG0004870.1 |
| 0.0874 | 3.0000 | 2140.0000 | 6.8709 | GO:0006793 phosphorus metabolic proc. | HORVU.MOREX.R2.1HG0001760.1 HORVU.MOREX.R2.1HG0002580.1 HORVU.MOREX.R2.1HG0004870.1 |

Table S12

Pathways upregulated in T22 vs C22

| Enrichment FDR | nGenes | Pathway Genes | Fold Enrichment | Pathway | Genes |
|---|---|---|---|---|---|
| 0.077 | 1.000 | 3.000 | 490.125 | GO:0008940 nitrate reductase activity | HORVU.MOREX.R2.6HG0513160.1 |
| 0.077 | 1.000 | 5.000 | 294.075 | GO:0016661 oxidoreductase activity acting on other nitrogenous compounds as donors | HORVU.MOREX.R2.6HG0513160.1 |
| 0.077 | 1.000 | 6.000 | 245.063 | GO:0004351 glutamate decarboxylase activity | HORVU.MOREX.R2.4HG0327490.1 |
| 0.077 | 1.000 | 7.000 | 210.054 | GO:0042126 nitrate metabolic proc. | HORVU.MOREX.R2.6HG0513160.1 |
| 0.077 | 1.000 | 7.000 | 210.054 | GO:0042128 nitrate assimilation | HORVU.MOREX.R2.6HG0513160.1 |
| 0.077 | 1.000 | 7.000 | 210.054 | GO:2001057 reactive nitrogen species metabolic proc. | HORVU.MOREX.R2.6HG0513160.1 |
| 0.077 | 1.000 | 9.000 | 163.375 | GO:0071941 nitrogen cycle metabolic proc. | HORVU.MOREX.R2.6HG0513160.1 |
| 0.077 | 1.000 | 9.000 | 163.375 | GO:0006536 glutamate metabolic proc. | HORVU.MOREX.R2.4HG0327490.1 |
| 0.077 | 5.000 | 1520.000 | 4.837 | GO:0004672 protein kinase activity | HORVU.MOREX.R2.1HG0001760.1<br>HORVU.MOREX.R2.1HG0002580.1<br>HORVU.MOREX.R2.3HG0256040.1<br>HORVU.MOREX.R2.4HG0341130.1<br>HORVU.MOREX.R2.7HG0569500.1 |
| 0.077 | 5.000 | 1729.000 | 4.252 | GO:0016301 kinase activity | HORVU.MOREX.R2.1HG0001760.1<br>HORVU.MOREX.R2.1HG0002580.1<br>HORVU.MOREX.R2.3HG0256040.1<br>HORVU.MOREX.R2.4HG0341130.1<br>HORVU.MOREX.R2.7HG0569500.1 |

Table S13

Pathways downregulated in T22 vs C22

| Enrichment FDR | nGenes | Pathway Genes | Fold Enrichment | Pathway | Genes |
|---|---|---|---|---|---|
| 0.000 | 3.000 | 30.000 | 115.324 | GO:0016759 cellulose synthase activity | HORVU.MOREX.R2.3HG0194300.1 HORVU.MOREX.R2.5HG0400090.1 HORVU.MOREX.R2.7HG0579230.1 |
| 0.000 | 3.000 | 30.000 | 115.324 | GO:0016760 cellulose synthase (UDP-forming) activity | HORVU.MOREX.R2.3HG0194300.1 HORVU.MOREX.R2.5HG0400090.1 HORVU.MOREX.R2.7HG0579230.1 |
| 0.000 | 3.000 | 39.000 | 88.710 | GO:0030244 cellulose biosynthetic proc. | HORVU.MOREX.R2.3HG0194300.1 HORVU.MOREX.R2.5HG0400090.1 HORVU.MOREX.R2.7HG0579230.1 |
| 0.001 | 3.000 | 45.000 | 76.882 | GO:0009832 plant-type cell wall biogenesis | HORVU.MOREX.R2.3HG0194300.1 HORVU.MOREX.R2.5HG0369430.1 HORVU.MOREX.R2.5HG0400090.1 |
| 0.001 | 3.000 | 52.000 | 66.533 | GO:0030243 cellulose metabolic proc. | HORVU.MOREX.R2.3HG0194300.1 HORVU.MOREX.R2.5HG0400090.1 HORVU.MOREX.R2.7HG0579230.1 |
| 0.001 | 3.000 | 55.000 | 62.904 | GO:0051274 beta-glucan biosynthetic proc. | HORVU.MOREX.R2.3HG0194300.1 HORVU.MOREX.R2.5HG0400090.1 HORVU.MOREX.R2.7HG0579230.1 |
| 0.000 | 4.000 | 104.000 | 44.355 | GO:0071669 plant-type cell wall organization or biogenesis | HORVU.MOREX.R2.3HG0194300.1 HORVU.MOREX.R2.5HG0369430.1 HORVU.MOREX.R2.5HG0400090.1 HORVU.MOREX.R2.7HG0579230.1 |
| 0.001 | 4.000 | 189.000 | 24.407 | GO:0071555 cell wall organization | HORVU.MOREX.R2.3HG0194300.1 HORVU.MOREX.R2.5HG0369430.1 HORVU.MOREX.R2.5HG0400090.1 HORVU.MOREX.R2.7HG0579230.1 |
| 0.001 | 4.000 | 197.000 | 23.416 | GO:0045229 external encapsulating structure organization | HORVU.MOREX.R2.3HG0194300.1 HORVU.MOREX.R2.5HG0369430.1 HORVU.MOREX.R2.5HG0400090.1 HORVU.MOREX.R2.7HG0579230.1 |
| 0.001 | 5.000 | 344.000 | 16.762 | GO:0008194 UDP-glycosyltransferase activity | HORVU.MOREX.R2.1HG0006010.1 HORVU.MOREX.R2.3HG0194300.1 HORVU.MOREX.R2.3HG0199730.1 HORVU.MOREX.R2.5HG0400090.1 HORVU.MOREX.R2.7HG0579230.1 |

### Table S14

Pathways upregulated in T42 vs C42

| Enrichment FDR | nGenes | Pathway Genes | Fold Enrichment | Pathway | Genes |
|---|---|---|---|---|---|
| 1.75E-34 | 62 | 1511 | 8.018 | GO:0006468 protein phosphorylation | HORVU.MOREX.R2.2HG0112790.1 HORVU.MOREX.R2.2HG0169700.1 HORVU.MOREX.R2.3HG0182560.1 HORVU.MOREX.R2.6HG0504490.1 HORVU.MOREX.R2.1HG0000640.1 HORVU.MOREX.R2.1HG0001160.1 HORVU.MOREX.R2.1HG0001400.1 HORVU.MOREX.R2.1HG0002140.1 HORVU.MOREX.R2.1HG0007120.1 HORVU.MOREX.R2.1HG0060880.1 HORVU.MOREX.R2.1HG0064720.1 HORVU.MOREX.R2.2HG0080580.1 HORVU.MOREX.R2.2HG0080620.1 HORVU.MOREX.R2.2HG0083080.1 HORVU.MOREX.R2.2HG0085880.1 HORVU.MOREX.R2.2HG0089430.1 HORVU.MOREX.R2.2HG0090920.1 HORVU.MOREX.R2.2HG0108680.1 HORVU.MOREX.R2.2HG0131160.1 HORVU.MOREX.R2.2HG0167340.1 HORVU.MOREX.R2.2HG0172730.1 HORVU.MOREX.R2.2HG0173420.1 HORVU.MOREX.R2.3HG0183930.1 HORVU.MOREX.R2.3HG0187780.1 HORVU.MOREX.R2.3HG0238970.1 HORVU.MOREX.R2.3HG0244560.1 HORVU.MOREX.R2.3HG0269080.1 HORVU.MOREX.R2.4HG0276520.1 HORVU.MOREX.R2.4HG0276540.1 HORVU.MOREX.R2.4HG0282400.1 HORVU.MOREX.R2.4HG0287960.1 HORVU.MOREX.R2.4HG0336220.1 HORVU.MOREX.R2.4HG0339440.1 HORVU.MOREX.R2.4HG0346270.1 HORVU.MOREX.R2.5HG0349880.1 HORVU.MOREX.R2.5HG0392950.1 HORVU.MOREX.R2.5HG0393010.1 HORVU.MOREX.R2.5HG0404510.1 HORVU.MOREX.R2.5HG0424170.1 HORVU.MOREX.R2.5HG0444860.1 HORVU.MOREX.R2.5HG0445920.1 HORVU.MOREX.R2.5HG0446810.1 HORVU.MOREX.R2.5HG0446820.1 HORVU.MOREX.R2.6HG0448250.1 HORVU.MOREX.R2.6HG0448330.1 HORVU.MOREX.R2.6HG0451270.1 HORVU.MOREX.R2.6HG0458780.1 HORVU.MOREX.R2.6HG0504970.1 HORVU.MOREX.R2.6HG0514460.1 HORVU.MOREX.R2.6HG0524700.1 HORVU.MOREX.R2.6HG0524710.1 HORVU.MOREX.R2.7HG0527370.1 HORVU.MOREX.R2.7HG0532400.1 HORVU.MOREX.R2.7HG0534210.1 HORVU.MOREX.R2.7HG0537810.1 HORVU.MOREX.R2.7HG0604430.1 HORVU.MOREX.R2.7HG0605030.1 HORVU.MOREX.R2.7HG0607550.1 HORVU.MOREX.R2.7HG0615970.1 HORVU.MOREX.R2.UNG0628380.1 HORVU.MOREX.R2.UNG0630000.1 HORVU.MOREX.R2.UNG0635440.1 |
| 1.75E-34 | 62 | 1520 | 7.970 | GO:0004672 protein kinase activity | HORVU.MOREX.R2.2HG0108680.1 HORVU.MOREX.R2.2HG0112790.1 HORVU.MOREX.R2.2HG0169700.1 |
| 1.53E-32 | 62 | 1663 | 7.285 | GO:0016773 phosphotransferase activity alcohol group as acceptor | HORVU.MOREX.R2.2HG0108680.1 HORVU.MOREX.R2.2HG0112790.1 HORVU.MOREX.R2.2HG0169700.1 HORVU.MOREX.R2.3HG0182560.1 HORVU.MOREX.R2.4HG0282400.1 HORVU.MOREX.R2.6HG0504490.1 HORVU.MOREX.R2.1HG0000640.1 HORVU.MOREX.R2.1HG0001160.1 HORVU.MOREX.R2.1HG0001400.1 HORVU.MOREX.R2.1HG0002140.1 HORVU.MOREX.R2.1HG0007120.1 HORVU.MOREX.R2.1HG0060880.1 HORVU.MOREX.R2.1HG0064720.1 HORVU.MOREX.R2.2HG0080580.1 HORVU.MOREX.R2.2HG0080620.1 HORVU.MOREX.R2.2HG0083080.1 HORVU.MOREX.R2.2HG0085880.1 HORVU.MOREX.R2.2HG0089430.1 HORVU.MOREX.R2.2HG0090920.1 HORVU.MOREX.R2.2HG0131160.1 HORVU.MOREX.R2.2HG0167340.1 HORVU.MOREX.R2.2HG0172730.1 HORVU.MOREX.R2.2HG0173420.1 HORVU.MOREX.R2.3HG0183930.1 HORVU.MOREX.R2.3HG0187780.1 HORVU.MOREX.R2.3HG0238970.1 HORVU.MOREX.R2.3HG0244560.1 HORVU.MOREX.R2.3HG0269080.1 HORVU.MOREX.R2.4HG0276520.1 HORVU.MOREX.R2.4HG0276540.1 HORVU.MOREX.R2.4HG0287960.1 HORVU.MOREX.R2.4HG0336220.1 HORVU.MOREX.R2.4HG0339440.1 HORVU.MOREX.R2.4HG0346270.1 HORVU.MOREX.R2.5HG0349880.1 HORVU.MOREX.R2.5HG0392950.1 HORVU.MOREX.R2.5HG0393010.1 HORVU.MOREX.R2.5HG0404510.1 HORVU.MOREX.R2.5HG0424170.1 HORVU.MOREX.R2.5HG0444860.1 HORVU.MOREX.R2.5HG0445920.1 HORVU.MOREX.R2.5HG0446810.1 HORVU.MOREX.R2.5HG0446820.1 HORVU.MOREX.R2.6HG0448250.1 HORVU.MOREX.R2.6HG0448330.1 HORVU.MOREX.R2.6HG0451270.1 HORVU.MOREX.R2.6HG0458780.1 HORVU.MOREX.R2.6HG0504970.1 HORVU.MOREX.R2.6HG0514460.1 HORVU.MOREX.R2.6HG0524700.1 HORVU.MOREX.R2.6HG0524710.1 HORVU.MOREX.R2.7HG0527370.1 HORVU.MOREX.R2.7HG0532400.1 HORVU.MOREX.R2.7HG0534210.1 HORVU.MOREX.R2.7HG0537810.1 HORVU.MOREX.R2.7HG0604430.1 HORVU.MOREX.R2.7HG0605030.1 HORVU.MOREX.R2.7HG0607550.1 HORVU.MOREX.R2.7HG0615970.1 HORVU.MOREX.R2.UNG0628380.1 HORVU.MOREX.R2.UNG0630000.1 HORVU.MOREX.R2.UNG0635440.1 |

| | | | | | |
|---|---|---|---|---|---|
| 1.53E-32 | 62 | 1665 | 7.276 | GO:0016310 phosphorylation | HORVU.MOREX.R2.2HG0112790.1 HORVU.MOREX.R2.2HG0169700.1 HORVU.MOREX.R2.3HG0182560.1 HORVU.MOREX.R2.6HG0504490.1 HORVU.MOREX.R2.1HG0000640.1 HORVU.MOREX.R2.1HG0001160.1 HORVU.MOREX.R2.1HG0001400.1 HORVU.MOREX.R2.1HG0002140.1 HORVU.MOREX.R2.1HG0007120.1 HORVU.MOREX.R2.1HG0060880.1 HORVU.MOREX.R2.1HG0064720.1 HORVU.MOREX.R2.2HG0080580.1 HORVU.MOREX.R2.2HG0080620.1 HORVU.MOREX.R2.2HG0083080.1 HORVU.MOREX.R2.2HG0085880.1 HORVU.MOREX.R2.2HG0089430.1 HORVU.MOREX.R2.2HG0090920.1 HORVU.MOREX.R2.2HG0108680.1 HORVU.MOREX.R2.2HG0131160.1 HORVU.MOREX.R2.2HG0167340.1 HORVU.MOREX.R2.2HG0172730.1 HORVU.MOREX.R2.2HG0173420.1 HORVU.MOREX.R2.3HG0183930.1 HORVU.MOREX.R2.3HG0187780.1 HORVU.MOREX.R2.3HG0238970.1 HORVU.MOREX.R2.3HG0244560.1 HORVU.MOREX.R2.3HG0269080.1 HORVU.MOREX.R2.4HG0276520.1 HORVU.MOREX.R2.4HG0276540.1 HORVU.MOREX.R2.4HG0282400.1 HORVU.MOREX.R2.4HG0287960.1 HORVU.MOREX.R2.4HG0336220.1 HORVU.MOREX.R2.4HG0339440.1 HORVU.MOREX.R2.4HG0346270.1 HORVU.MOREX.R2.5HG0349880.1 HORVU.MOREX.R2.5HG0392950.1 HORVU.MOREX.R2.5HG0393010.1 HORVU.MOREX.R2.5HG0404510.1 HORVU.MOREX.R2.5HG0424170.1 HORVU.MOREX.R2.5HG0444860.1 HORVU.MOREX.R2.5HG0445920.1 HORVU.MOREX.R2.5HG0446810.1 HORVU.MOREX.R2.5HG0446820.1 HORVU.MOREX.R2.6HG0448250.1 HORVU.MOREX.R2.6HG0448330.1 HORVU.MOREX.R2.6HG0451270.1 HORVU.MOREX.R2.6HG0458780.1 HORVU.MOREX.R2.6HG0504970.1 HORVU.MOREX.R2.6HG0514460.1 HORVU.MOREX.R2.6HG0524700.1 HORVU.MOREX.R2.6HG0524710.1 HORVU.MOREX.R2.7HG0527370.1 HORVU.MOREX.R2.7HG0532400.1 HORVU.MOREX.R2.7HG0534210.1 HORVU.MOREX.R2.7HG0537810.1 HORVU.MOREX.R2.7HG0604430.1 HORVU.MOREX.R2.7HG0605030.1 HORVU.MOREX.R2.7HG0607550.1 HORVU.MOREX.R2.7HG0615970.1 HORVU.MOREX.R2.UNG0628380.1 HORVU.MOREX.R2.UNG0630000.1 HORVU.MOREX.R2.UNG0635440.1 |
| 1.02E-31 | 62 | 1729 | 7.007 | GO:0016301 kinase activity | HORVU.MOREX.R2.2HG0108680.1 HORVU.MOREX.R2.2HG0112790.1 HORVU.MOREX.R2.2HG0169700.1 |
| 8.51E-31 | 64 | 1943 | 6.436 | GO:0016772 transferase activity transferring phosphorus-containing groups | HORVU.MOREX.R2.2HG0108680.1 HORVU.MOREX.R2.2HG0112790.1 HORVU.MOREX.R2.2HG0169700.1 HORVU.MOREX.R2.3HG0182560.1 HORVU.MOREX.R2.4HG0282400.1 HORVU.MOREX.R2.6HG0504490.1 HORVU.MOREX.R2.1HG0000640.1 HORVU.MOREX.R2.1HG0001160.1 HORVU.MOREX.R2.1HG0001400.1 HORVU.MOREX.R2.1HG0002140.1 HORVU.MOREX.R2.1HG0007120.1 HORVU.MOREX.R2.1HG0060880.1 HORVU.MOREX.R2.1HG0064720.1 HORVU.MOREX.R2.2HG0080580.1 HORVU.MOREX.R2.2HG0080620.1 HORVU.MOREX.R2.2HG0083080.1 HORVU.MOREX.R2.2HG0085880.1 HORVU.MOREX.R2.2HG0089430.1 HORVU.MOREX.R2.2HG0090920.1 HORVU.MOREX.R2.2HG0131160.1 HORVU.MOREX.R2.2HG0145800.1 HORVU.MOREX.R2.2HG0167340.1 HORVU.MOREX.R2.2HG0172730.1 HORVU.MOREX.R2.2HG0173420.1 HORVU.MOREX.R2.3HG0183930.1 HORVU.MOREX.R2.3HG0187780.1 HORVU.MOREX.R2.3HG0238970.1 HORVU.MOREX.R2.3HG0244560.1 HORVU.MOREX.R2.3HG0269080.1 HORVU.MOREX.R2.4HG0276520.1 HORVU.MOREX.R2.4HG0276540.1 HORVU.MOREX.R2.4HG0287960.1 HORVU.MOREX.R2.4HG0336220.1 HORVU.MOREX.R2.4HG0339440.1 HORVU.MOREX.R2.4HG0346270.1 HORVU.MOREX.R2.5HG0349880.1 HORVU.MOREX.R2.5HG0392950.1 HORVU.MOREX.R2.5HG0393010.1 HORVU.MOREX.R2.5HG0404510.1 HORVU.MOREX.R2.5HG0424170.1 HORVU.MOREX.R2.5HG0444860.1 HORVU.MOREX.R2.5HG0445920.1 HORVU.MOREX.R2.5HG0446810.1 HORVU.MOREX.R2.5HG0446820.1 HORVU.MOREX.R2.6HG0448250.1 HORVU.MOREX.R2.6HG0448330.1 HORVU.MOREX.R2.6HG0451270.1 HORVU.MOREX.R2.6HG0458780.1 HORVU.MOREX.R2.6HG0504970.1 HORVU.MOREX.R2.6HG0508620.1 HORVU.MOREX.R2.6HG0514460.1 HORVU.MOREX.R2.6HG0524700.1 HORVU.MOREX.R2.6HG0524710.1 HORVU.MOREX.R2.7HG0527370.1 HORVU.MOREX.R2.7HG0532400.1 HORVU.MOREX.R2.7HG0534210.1 HORVU.MOREX.R2.7HG0537810.1 HORVU.MOREX.R2.7HG0604430.1 HORVU.MOREX.R2.7HG0605030.1 HORVU.MOREX.R2.7HG0607550.1 HORVU.MOREX.R2.7HG0615970.1 HORVU.MOREX.R2.UNG0628380.1 HORVU.MOREX.R2.UNG0630000.1 HORVU.MOREX.R2.UNG0635440.1 |
| 4.19E-30 | 72 | 2622 | 5.366 | GO:0005524 ATP binding | HORVU.MOREX.R2.1HG0000640.1 HORVU.MOREX.R2.1HG0001160.1 HORVU.MOREX.R2.1HG0001400.1 |
| 1.93E-29 | 75 | 2946 | 4.975 | GO:0032559 adenyl ribonucleotide | HORVU.MOREX.R2.1HG0000640.1 HORVU.MOREX.R2.1HG0001160.1 HORVU.MOREX.R2.1HG0001400.1 |
| 1.93E-29 | 75 | 2951 | 4.966 | GO:0030554 adenyl nucleotide binding | HORVU.MOREX.R2.1HG0000640.1 HORVU.MOREX.R2.1HG0001160.1 HORVU.MOREX.R2.1HG0001400.1 |

| | | | | | |
|---|---|---|---|---|---|
| 4.42E-28 | 72 | 2843 | 4.949 | GO:0035639 purine ribonucleoside triphosphate binding | HORVU.MOREX.R2.1HG0000640.1 HORVU.MOREX.R2.1HG0001160.1 HORVU.MOREX.R2.1HG0001400.1 HORVU.MOREX.R2.1HG0002140.1 HORVU.MOREX.R2.1HG0007120.1 HORVU.MOREX.R2.1HG0060880.1 HORVU.MOREX.R2.1HG0064720.1 HORVU.MOREX.R2.2HG0080580.1 HORVU.MOREX.R2.2HG0080620.1 HORVU.MOREX.R2.2HG0083080.1 HORVU.MOREX.R2.2HG0085880.1 HORVU.MOREX.R2.2HG0089430.1 HORVU.MOREX.R2.2HG0090920.1 HORVU.MOREX.R2.2HG0108680.1 HORVU.MOREX.R2.2HG0112790.1 HORVU.MOREX.R2.2HG0131160.1 HORVU.MOREX.R2.2HG0145800.1 HORVU.MOREX.R2.2HG0167340.1 HORVU.MOREX.R2.2HG0169700.1 HORVU.MOREX.R2.2HG0172730.1 HORVU.MOREX.R2.2HG0173420.1 HORVU.MOREX.R2.3HG0182560.1 HORVU.MOREX.R2.3HG0183930.1 HORVU.MOREX.R2.3HG0187780.1 HORVU.MOREX.R2.3HG0224440.1 HORVU.MOREX.R2.3HG0238970.1 HORVU.MOREX.R2.3HG0239750.1 HORVU.MOREX.R2.3HG0244560.1 HORVU.MOREX.R2.3HG0269080.1 HORVU.MOREX.R2.4HG0276520.1 HORVU.MOREX.R2.4HG0276540.1 HORVU.MOREX.R2.4HG0282400.1 HORVU.MOREX.R2.4HG0287960.1 HORVU.MOREX.R2.4HG0336220.1 HORVU.MOREX.R2.4HG0339440.1 HORVU.MOREX.R2.4HG0346270.1 HORVU.MOREX.R2.5HG0349880.1 HORVU.MOREX.R2.5HG0364350.1 HORVU.MOREX.R2.5HG0392950.1 HORVU.MOREX.R2.5HG0393010.1 HORVU.MOREX.R2.5HG0404510.1 HORVU.MOREX.R2.5HG0408180.1 HORVU.MOREX.R2.5HG0414110.1 HORVU.MOREX.R2.5HG0424170.1 HORVU.MOREX.R2.5HG0444860.1 HORVU.MOREX.R2.5HG0445920.1 HORVU.MOREX.R2.5HG0446810.1 HORVU.MOREX.R2.5HG0446820.1 HORVU.MOREX.R2.5HG0447820.1 HORVU.MOREX.R2.6HG0448250.1 HORVU.MOREX.R2.6HG0448330.1 HORVU.MOREX.R2.6HG0451270.1 HORVU.MOREX.R2.6HG0458780.1 HORVU.MOREX.R2.6HG0504490.1 HORVU.MOREX.R2.6HG0504970.1 HORVU.MOREX.R2.6HG0514460.1 HORVU.MOREX.R2.6HG0524700.1 HORVU.MOREX.R2.6HG0524710.1 HORVU.MOREX.R2.7HG0527370.1 HORVU.MOREX.R2.7HG0532400.1 HORVU.MOREX.R2.7HG0534210.1 HORVU.MOREX.R2.7HG0537810.1 HORVU.MOREX.R2.7HG0561520.1 HORVU.MOREX.R2.7HG0593370.1 HORVU.MOREX.R2.7HG0599230.1 HORVU.MOREX.R2.7HG0604430.1 HORVU.MOREX.R2.7HG0605030.1 HORVU.MOREX.R2.7HG0607550.1 HORVU.MOREX.R2.7HG0615970.1 HORVU.MOREX.R2.UNG0628380.1 HORVU.MOREX.R2.UNG0630000.1 HORVU.MOREX.R2.UNG0635440.1 |

Table S15

Pathways downregulated in T42 vs C42

| Enrichment FDR | nGenes | Pathway Genes | Fold Enrichment | Pathway | Genes |
|---|---|---|---|---|---|
| | | | | | HORVU.MOREX.R2.1HG0064430.1 |
| | | | | | HORVU.MOREX.R2.1HG0073470.1 |
| | | | | | HORVU.MOREX.R2.1HG0073480.1 |
| | | | | | HORVU.MOREX.R2.2HG0111260.1 |
| | | | | | HORVU.MOREX.R2.2HG0142880.1 |
| | | | | | HORVU.MOREX.R2.5HG0405770.1 |
| | | | | | HORVU.MOREX.R2.5HG0418240.1 |
| | | | | | HORVU.MOREX.R2.5HG0434050.1 |
| | | | | | HORVU.MOREX.R2.5HG0434090.1 |
| | | | | | HORVU.MOREX.R2.6HG0523360.1 |
| | | | | | HORVU.MOREX.R2.7HG0554640.1 |
| 1.69E-23 | 12 | 31 | 206.973607 | GO:0009522 photosystem I | HORVU.MOREX.R2.7HG0554650.1 |
| | | | | | HORVU.MOREX.R2.1HG0073470.1 |
| | | | | | HORVU.MOREX.R2.5HG0418240.1 |
| | | | | | HORVU.MOREX.R2.5HG0434090.1 |
| | | | | | HORVU.MOREX.R2.7HG0554650.1 |
| | | | | | HORVU.MOREX.R2.1HG0064430.1 |
| | | | | | HORVU.MOREX.R2.1HG0073480.1 |
| | | | | | HORVU.MOREX.R2.1HG0073500.1 |
| | | | | | HORVU.MOREX.R2.2HG0111260.1 |
| | | | | | HORVU.MOREX.R2.2HG0142880.1 |
| | | | | | HORVU.MOREX.R2.5HG0434050.1 |
| | | | | | HORVU.MOREX.R2.6HG0460180.1 |
| | | | | GO:0009765 photosynthesis light | HORVU.MOREX.R2.6HG0523360.1 |
| 4.74E-24 | 13 | 40 | 173.7715909 | harvesting | HORVU.MOREX.R2.7HG0554640.1 |

| | | | | | |
|---|---|---|---|---|---|
| 4.27E-23 | 14 | 70 | 106.9363636 | GO:0009535 chloroplast thylakoid membrane | HORVU.MOREX.R2.1HG0073470.1<br>HORVU.MOREX.R2.4HG0321470.1<br>HORVU.MOREX.R2.5HG0418240.1<br>HORVU.MOREX.R2.5HG0434090.1<br>HORVU.MOREX.R2.7HG0554650.1<br>HORVU.MOREX.R2.1HG0064430.1<br>HORVU.MOREX.R2.1HG0073480.1<br>HORVU.MOREX.R2.2HG0111260.1<br>HORVU.MOREX.R2.2HG0142880.1<br>HORVU.MOREX.R2.3HG0214090.1<br>HORVU.MOREX.R2.5HG0405770.1<br>HORVU.MOREX.R2.5HG0434050.1<br>HORVU.MOREX.R2.6HG0523360.1<br>HORVU.MOREX.R2.7HG0554640.1 |
| 4.27E-23 | 14 | 71 | 105.4302177 | GO:0055035 plastid thylakoid membrane | HORVU.MOREX.R2.1HG0073470.1<br>HORVU.MOREX.R2.4HG0321470.1<br>HORVU.MOREX.R2.5HG0418240.1<br>HORVU.MOREX.R2.5HG0434090.1<br>HORVU.MOREX.R2.7HG0554650.1<br>HORVU.MOREX.R2.1HG0064430.1<br>HORVU.MOREX.R2.1HG0073480.1<br>HORVU.MOREX.R2.2HG0111260.1<br>HORVU.MOREX.R2.2HG0142880.1<br>HORVU.MOREX.R2.3HG0214090.1<br>HORVU.MOREX.R2.5HG0405770.1<br>HORVU.MOREX.R2.5HG0434050.1<br>HORVU.MOREX.R2.6HG0523360.1<br>HORVU.MOREX.R2.7HG0554640.1 |

| | | | | | |
|---|---|---|---|---|---|
| | | | | | HORVU.MOREX.R2.1HG0073470.1 |
| | | | | | HORVU.MOREX.R2.4HG0321470.1 |
| | | | | | HORVU.MOREX.R2.5HG0418240.1 |
| | | | | | HORVU.MOREX.R2.5HG0434090.1 |
| | | | | | HORVU.MOREX.R2.7HG0554650.1 |
| | | | | | HORVU.MOREX.R2.1HG0064430.1 |
| | | | | | HORVU.MOREX.R2.1HG0073480.1 |
| | | | | | HORVU.MOREX.R2.2HG0111260.1 |
| | | | | | HORVU.MOREX.R2.2HG0142880.1 |
| | | | | | HORVU.MOREX.R2.3HG0214090.1 |
| | | | | | HORVU.MOREX.R2.5HG0405770.1 |
| | | | | | HORVU.MOREX.R2.5HG0434050.1 |
| | | | | | HORVU.MOREX.R2.6HG0523360.1 |
| 2.23E-22 | 14 | 81 | 92.41414141 | GO:0009534 chloroplast thylakoid | HORVU.MOREX.R2.7HG0554640.1 |
| | | | | | HORVU.MOREX.R2.1HG0073470.1 |
| | | | | | HORVU.MOREX.R2.4HG0321470.1 |
| | | | | | HORVU.MOREX.R2.5HG0418240.1 |
| | | | | | HORVU.MOREX.R2.5HG0434090.1 |
| | | | | | HORVU.MOREX.R2.7HG0554650.1 |
| | | | | | HORVU.MOREX.R2.1HG0064430.1 |
| | | | | | HORVU.MOREX.R2.1HG0073480.1 |
| | | | | | HORVU.MOREX.R2.2HG0111260.1 |
| | | | | | HORVU.MOREX.R2.2HG0142880.1 |
| | | | | | HORVU.MOREX.R2.3HG0214090.1 |
| | | | | | HORVU.MOREX.R2.3HG0270810.1 |
| | | | | | HORVU.MOREX.R2.5HG0405770.1 |
| | | | | | HORVU.MOREX.R2.5HG0434050.1 |
| | | | | | HORVU.MOREX.R2.6HG0523360.1 |
| 7.23E-23 | 15 | 101 | 79.40819082 | GO:0042651 thylakoid membrane | HORVU.MOREX.R2.7HG0554640.1 |

| | | | | | |
|---|---|---|---|---|---|
| | | | | | HORVU.MOREX.R2.1HG0073470.1 |
| | | | | | HORVU.MOREX.R2.4HG0321470.1 |
| | | | | | HORVU.MOREX.R2.5HG0418240.1 |
| | | | | | HORVU.MOREX.R2.5HG0434090.1 |
| | | | | | HORVU.MOREX.R2.7HG0554650.1 |
| | | | | | HORVU.MOREX.R2.1HG0064430.1 |
| | | | | | HORVU.MOREX.R2.1HG0073480.1 |
| | | | | | HORVU.MOREX.R2.2HG0111260.1 |
| | | | | | HORVU.MOREX.R2.2HG0142880.1 |
| | | | | | HORVU.MOREX.R2.2HG0147310.1 |
| | | | | | HORVU.MOREX.R2.3HG0214090.1 |
| | | | | | HORVU.MOREX.R2.3HG0270810.1 |
| | | | | | HORVU.MOREX.R2.5HG0405770.1 |
| | | | | | HORVU.MOREX.R2.5HG0434050.1 |
| | | | | | HORVU.MOREX.R2.6HG0523360.1 |
| 1.78E-22 | 16 | 142 | 60.24583867 | GO:0009526 plastid envelope | HORVU.MOREX.R2.7HG0554640.1 |

| | | | | | |
|---|---|---|---|---|---|
| | | | | | HORVU.MOREX.R2.1HG0073470.1<br>HORVU.MOREX.R2.5HG0418240.1<br>HORVU.MOREX.R2.5HG0434090.1<br>HORVU.MOREX.R2.7HG0554650.1<br>HORVU.MOREX.R2.1HG0029020.1<br>HORVU.MOREX.R2.1HG0064430.1<br>HORVU.MOREX.R2.1HG0073480.1<br>HORVU.MOREX.R2.1HG0073500.1<br>HORVU.MOREX.R2.2HG0086010.1<br>HORVU.MOREX.R2.2HG0086020.1<br>HORVU.MOREX.R2.2HG0086030.1<br>HORVU.MOREX.R2.2HG0086050.1<br>HORVU.MOREX.R2.2HG0111260.1<br>HORVU.MOREX.R2.2HG0142880.1<br>HORVU.MOREX.R2.3HG0214090.1<br>HORVU.MOREX.R2.5HG0405770.1<br>HORVU.MOREX.R2.5HG0434050.1<br>HORVU.MOREX.R2.6HG0460180.1<br>HORVU.MOREX.R2.6HG0523360.1 |
| 9.37E-27 | 20 | 190 | 56.28229665 | GO:0015979 photosynthesis | HORVU.MOREX.R2.7HG0554640.1 |

| | | | | | |
|---|---|---|---|---|---|
| | | | | | HORVU.MOREX.R2.1HG0073470.1 |
| | | | | | HORVU.MOREX.R2.3HG0270810.1 |
| | | | | | HORVU.MOREX.R2.4HG0321470.1 |
| | | | | | HORVU.MOREX.R2.5HG0418240.1 |
| | | | | | HORVU.MOREX.R2.5HG0434090.1 |
| | | | | | HORVU.MOREX.R2.7HG0554650.1 |
| | | | | | HORVU.MOREX.R2.1HG0029020.1 |
| | | | | | HORVU.MOREX.R2.1HG0064430.1 |
| | | | | | HORVU.MOREX.R2.1HG0073480.1 |
| | | | | | HORVU.MOREX.R2.2HG0086010.1 |
| | | | | | HORVU.MOREX.R2.2HG0086020.1 |
| | | | | | HORVU.MOREX.R2.2HG0086030.1 |
| | | | | | HORVU.MOREX.R2.2HG0086050.1 |
| | | | | | HORVU.MOREX.R2.2HG0109810.1 |
| | | | | | HORVU.MOREX.R2.2HG0111260.1 |
| | | | | | HORVU.MOREX.R2.2HG0142880.1 |
| | | | | | HORVU.MOREX.R2.2HG0147310.1 |
| | | | | | HORVU.MOREX.R2.3HG0214090.1 |
| | | | | | HORVU.MOREX.R2.5HG0405770.1 |
| | | | | | HORVU.MOREX.R2.5HG0434050.1 |
| | | | | | HORVU.MOREX.R2.6HG0523360.1 |
| | | | | | HORVU.MOREX.R2.7HG0553370.1 |
| 6.42E-24 | 23 | 464 | 26.50362461 | GO:0009507 chloroplast | HORVU.MOREX.R2.7HG0554640.1 |

| | | | | | |
|---|---|---|---|---|---|
| | | | | | HORVU.MOREX.R2.1HG0073470.1 |
| | | | | | HORVU.MOREX.R2.3HG0270810.1 |
| | | | | | HORVU.MOREX.R2.4HG0321470.1 |
| | | | | | HORVU.MOREX.R2.5HG0418240.1 |
| | | | | | HORVU.MOREX.R2.5HG0434090.1 |
| | | | | | HORVU.MOREX.R2.7HG0554650.1 |
| | | | | | HORVU.MOREX.R2.1HG0029020.1 |
| | | | | | HORVU.MOREX.R2.1HG0064430.1 |
| | | | | | HORVU.MOREX.R2.1HG0073480.1 |
| | | | | | HORVU.MOREX.R2.2HG0086010.1 |
| | | | | | HORVU.MOREX.R2.2HG0086020.1 |
| | | | | | HORVU.MOREX.R2.2HG0086030.1 |
| | | | | | HORVU.MOREX.R2.2HG0086050.1 |
| | | | | | HORVU.MOREX.R2.2HG0109810.1 |
| | | | | | HORVU.MOREX.R2.2HG0111260.1 |
| | | | | | HORVU.MOREX.R2.2HG0142880.1 |
| | | | | | HORVU.MOREX.R2.2HG0147310.1 |
| | | | | | HORVU.MOREX.R2.3HG0214090.1 |
| | | | | | HORVU.MOREX.R2.5HG0405770.1 |
| | | | | | HORVU.MOREX.R2.5HG0434050.1 |
| | | | | | HORVU.MOREX.R2.6HG0523360.1 |
| | | | | | HORVU.MOREX.R2.7HG0553370.1 |
| 4.27E-23 | 23 | 521 | 23.60399581 | GO:0009536 plastid | HORVU.MOREX.R2.7HG0554640.1 |
